# Structure-based discovery of CFTR potentiators and inhibitors

**DOI:** 10.1101/2023.09.09.557002

**Authors:** Fangyu Liu, Anat Levit Kaplan, Jesper Levring, Jürgen Einsiedel, Stephanie Tiedt, Katharina Distler, Natalie S. Omattage, Ivan S. Kondratov, Yurii S. Moroz, Harlan L. Pietz, John J. Irwin, Peter Gmeiner, Brian K. Shoichet, Jue Chen

## Abstract

The cystic fibrosis transmembrane conductance regulator (CFTR) is a crucial ion channel whose loss of function leads to cystic fibrosis, while its hyperactivation leads to secretory diarrhea. Small molecules that improve CFTR folding (correctors) or function (potentiators) are clinically available. However, the only potentiator, ivacaftor, has suboptimal pharmacokinetics and inhibitors have yet to be clinically developed. Here we combine molecular docking, electrophysiology, cryo-EM, and medicinal chemistry to identify novel CFTR modulators. We docked ∼155 million molecules into the potentiator site on CFTR, synthesized 53 test ligands, and used structure-based optimization to identify candidate modulators. This approach uncovered novel mid-nanomolar potentiators as well as inhibitors that bind to the same allosteric site. These molecules represent potential leads for the development of more effective drugs for cystic fibrosis and secretory diarrhea, demonstrating the feasibility of large-scale docking for ion channel drug discovery.

## Introduction

CFTR is an anion channel that is widely expressed in epithelial cells of the lung, intestine, pancreas, and reproductive tract, where it regulates salt and fluid homeostasis^1^. Mutations that disrupt CFTR biosynthesis, folding, trafficking, or ion permeation cause cystic fibrosis (CF), a lethal genetic disease with no cure^2^. In addition, excessive activation of CFTR by bacterial pathogens such as *Vibrio cholerae* and enterotoxigenic *Escherichia coli* leads to secretory diarrhea, a major cause of mortality in children under the age of five^3^. For these reasons, modulators that either up- or downregulate CFTR activity have long been pursued as drug candidates.

Although negative CFTR modulators have not yet been advanced to the clinic, there has been considerable progress in the development of positive modulators, including correctors that increase the abundance of CFTR at the cell surface and potentiators that enhance anion flux^4–6^. Thus far, one potentiator (ivacaftor or VX-770) and three correctors (lumacaftor, tezacaftor, and elexacaftor) have been made available to CF patients^2^. Ivacaftor is prescribed for 178 different CFTR mutations – either singly or in combination with correctors^7^. Although it has certainly improved the health of many CF patients, its physical properties and pharmacokinetics are far from optimal. Ivacaftor is difficult to formulate due to its low water solubility (<0.05 µg/mL; cLogP = 5.6)^8^. Moreover, its bioavailability is highly variable as 99% is bound to plasma proteins^9,10^ and its usage is limited due to side effects, including liver disease and childhood cataracts^11^. Meanwhile, the high expense of the drug (over $200,000 per year) is a strain on public health insurance, and puts the drug out of reach of many. Alternative CFTR potentiators, including those inspired by ivacaftor, would therefore be beneficial for CF patients.

Ivacaftor’s mechanism of action has been studied structurally and functionally. Electrophysiological measurements showed that the drug increases the open probability of many mutants, as well as wild-type (WT), CFTR channels^12,13^. Cryo-EM structures further revealed that ivacaftor binds to CFTR within the lipid membrane, at a hinge region important for gating^14^. The binding-site does not overlap with the positions of disease-causing mutations such as ΔF508 or G551D, indicating that ivacaftor acts as an allosteric modulator. A chemically distinct CFTR potentiator, GLPG1837, binds to the same site on CFTR, indicating that this binding pocket is a hotspot for regulatory ligands. Because the pocket is unique to CFTR, and is not conserved in closely related proteins^15^, it is an excellent target for the discovery of CFTR modulators without off-target effects.

As a result of recent developments in the use of chemical libraries for molecular docking, it is now feasible to computationally screen large and, more recently, ultra-large chemical libraries to identify potential ligands^16–21^. For example, nanomolar and sub-nanomolar ligands have been identified for dopamine D4, melatonin MT1, sigma2, and alpha2a adrenergic receptors from structure-based virtual screening^16,19–21^. Thus far, most of the large-scale library screens have focused on enzymes^16,18^ and G protein-coupled receptors (GPCRs)^17,19–22^. Few studies have been performed on membrane transporters or ion channels. Furthermore, the potentiator-binding site in CFTR is shallow and directly exposed to membrane, posing an extra challenge for virtual screening.

In this study, we sought to identify novel CFTR ligands by iterative molecular docking, electrophysiology, cryo-EM, and medicinal chemistry efforts (Figure 1a). The structure of CFTR in complex with ivacaftor was used to computationally dock a large virtual library of diverse chemical scaffolds. Through iterative optimization, we identified a novel potentiator scaffold with mid-nanomolar affinity that is chemically distinct from known CFTR potentiators and that has favorable physical properties and pharmacokinetics. We also discovered modulators that bind to the potentiator site but inhibit CFTR activity, demonstrating that the membrane-exposed allosteric site can be explored to downregulate CFTR activity.

**Figure 1:**
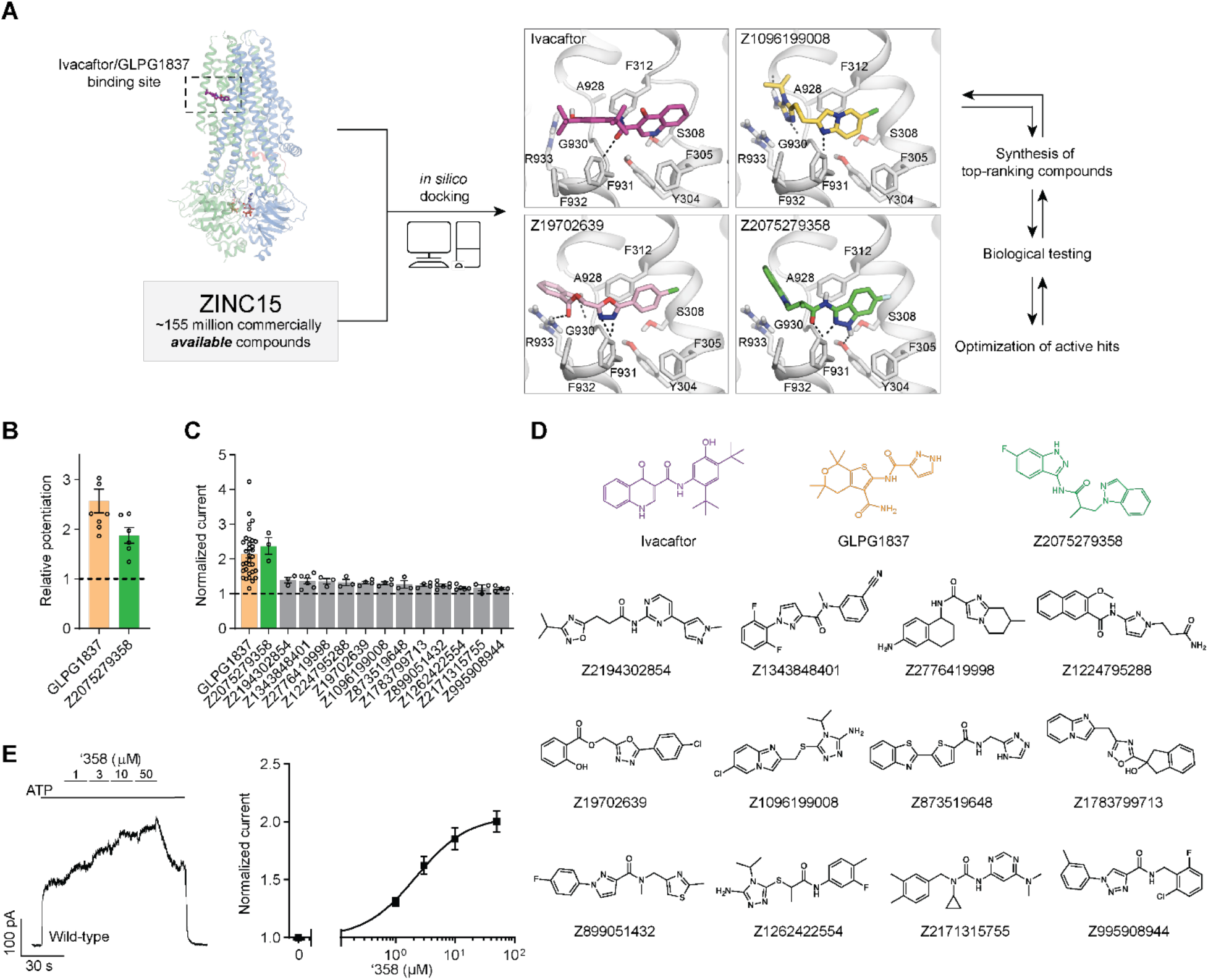
Ultra-large docking screen identifies novel CFTR potentiators. **(a)** The workflow of this study. **(b)** Compound Z2075279358 (’358) potentiates ΔF508 CFTR. CFBE41o^-^ cells homozygous for ΔF508 CFTR were treated with 1 μM lumacaftor and 20 μM forskolin. The relative potentiation was calculated as the ratio of flux rates with and without potentiator. Data points represent the means and standard errors (SEs) of 6 to 8 measurements (each shown as a dot). **(c)** Potentiation activity of 10 µM GLPG1837 or 5 µM compound against WT CFTR fused to a carboxy-terminal GFP tag. Inside-out membrane patches containing WT CFTR were excised from CHO cells and then fully phosphorylated by protein kinase A (PKA) in the presence of 3 mM ATP. The fold stimulation is defined as the ratio of the current in the presence and absence of added compound. Data represent means and SEs of 3-33 patches with individual measurements shown as dots. **(d)** The 2D structures of the potentiators GLPG1837, ivacaftor, and the 13 positive hits from the initial screen. **(e)** Representative macroscopic current trace and dose-response curve of WT CFTR in response to perfusion with ‘358. CFTR-containing membrane patches were fully phosphorylated by PKA. The current in the presence of 3 mM ATP before titration was used to normalize the current potentiated by different concentrations of ‘358. The *EC_50_* is estimated to be 2.2 ± 0.6 μM by fitting the dose-responses with the Hill equation. Data represent means and SEs from 3 patches.

## Results

### Identification of CFTR potentiators

Seeking new CFTR modulators, we docked a virtual library of ∼155 million tangible “drug-like” molecules (molecular weight 300-350 Da; logP ≤ 3.5) from the ZINC database (http://zinc15.docking.org)^23,24^ to the CFTR/ivacaftor cryo-EM structure (PDB: 6O2P) using DOCK3.7 (Figure 1a)^25^. Each molecule was sampled in an average of 421 conformations and 3,727 orientations, resulting in over 63 billion complexes being sampled in 76,365 core hours on an in-house cluster. Seeking diverse chemotypes, we clustered the top 100,000 scoring molecules by 2D similarity using ECFP4 fingerprints and a Tanimoto Coefficient (Tc) cutoff of 0.5. The top ranked molecules from each cluster (a total of 1,000) were visually inspected to remove molecules that were conformationally strained or had unsatisfied hydrogen bond acceptors or donors. Compounds engaging the key potentiator-binding residues S308, Y304, F312, and F931 (PDB: 6O2P and 6O1V) were prioritized and 58 of those, each from a different chemotype family, were selected for experimental evaluation.

Of the 58 prioritized compounds fifty-three were synthesized successfully from the Enamine make-on-demand set, a 91% fulfillment rate; as far as we know, all were new to the planet. They were evaluated for their effects on the CFTR variant found in 90% of CF patients (ΔF508). GLPG1837 was used as a benchmark because ivacaftor has very low solubility^26^, making it difficult to work with, and studies have shown that GLPG1837, although it was not advanced to the clinic, has a higher cellular efficacy than does ivacaftor for CFTR potentiation^27,28^. CFTR-mediated ion flux was measured in a bronchial epithelial cell line derived from a patient homozygous for ΔF508 (CFBE41o^-^) using a well-established halide flux assay^29^. Several of the new compounds potentiated CFTR-mediated ion flux (Extended Figure 1), in particular Z2075279358 (‘358), which increased ion flux to an extent close to that of GLPG1837 (Figure 1b).

Positive hits from this cellular assay were further analyzed electrophysiologically in inside-out membrane patches containing phosphorylated WT CFTR channels (Figure 1c). At a concentration of 5 μM, 13 compounds increased macroscopic currents by 1.2 to 2.4-fold (Figure 1c). The chemotypes of these 13 docking hits were diverse and novel compared to known CFTR potentiators such as GLPG1837 and ivacaftor (Figure 1d). In agreement with our cellular assay, the most efficacious compound was ‘358, whose efficacy was comparable to that of GLPG1837 (increased macroscopic current by 2.2-fold)^14,28,30^ with an EC_50_ of 2.2 ± 0.6 µM. Furthermore, our large-scale docking screen had a 24% hit rate, further supporting the feasibility of large-scale library screens for identifying novel allosteric ion channel modulators and resulted in candidate potentiators that are effective on both WT and ΔF508 CFTR (Figure 1c and Figure E1).

### Analog screen to identify additional potentiators

To identify more potent ligands for CFTR, we screened the ZINC15 library for analogs of ‘358. These were docked into the allosteric potentiator-binding site, and prioritized based on fit. Thirteen high-scoring analogs were selected for synthesis and tested using electrophysiology (Figure 2a). Surprisingly, only one molecule, Z1834339853 (‘853), potentiated WT CFTR currents to the same degree as ‘358 (Figure 2). The other 12 analogs had little or no effect. Concerned by this apparent failure of SAR, we re-examined the structures of the synthesized compounds by detailed NMR spectroscopy. This revealed that both ‘358 and ‘853 are regioisomers of the molecules specified in ZINC15 (Figure 2b). The acyl side chains on the exocyclic nitrogens of the indazole rings in both compounds in the ZINC15 database are instead on the cyclic nitrogens (Figure 2b).

**Figure 2:**
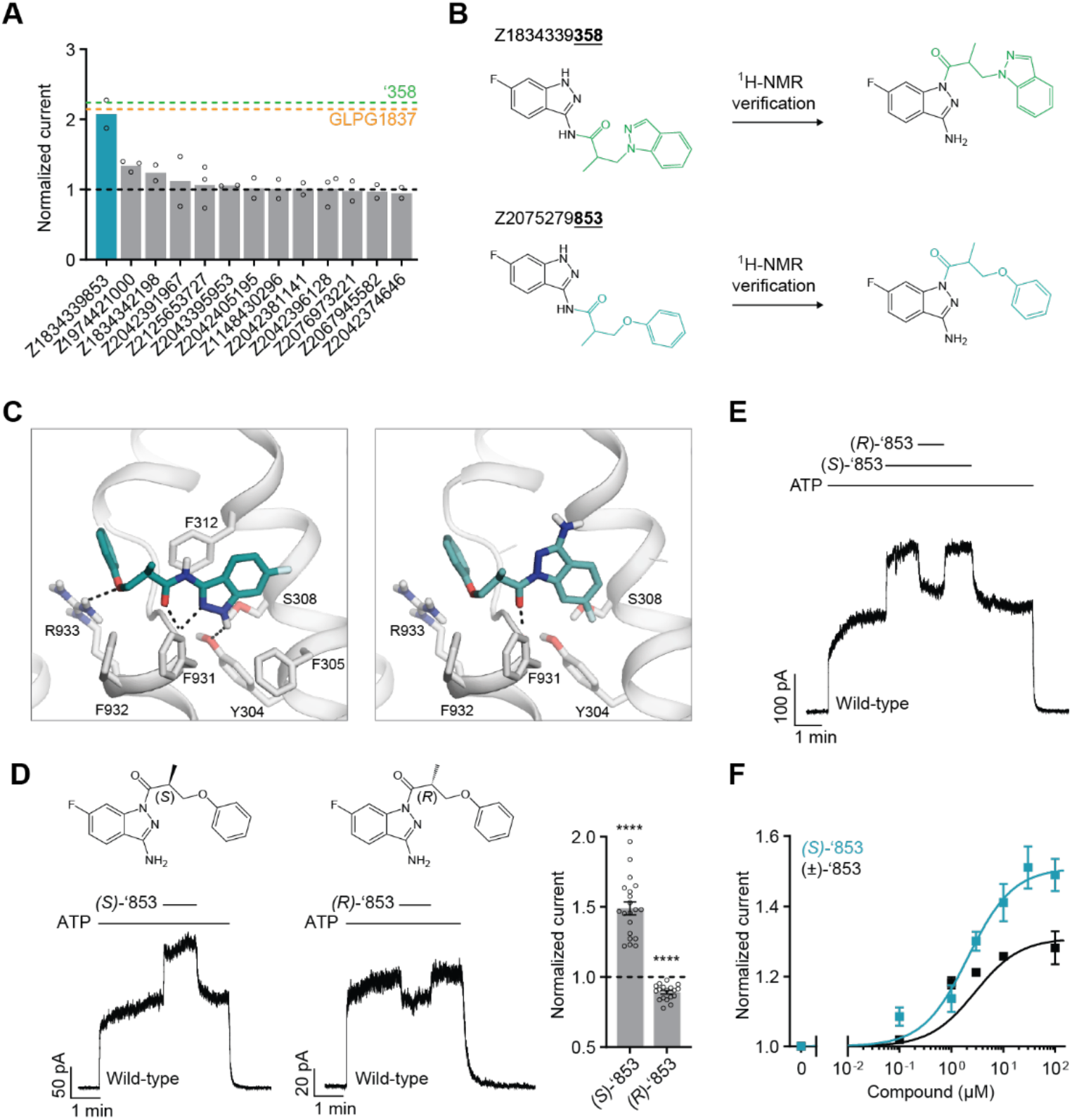
New modulators identified through an analog screen. **(a)** Potentiation activity of ‘358 analogs. All compounds were tested at a single concentration of 10 □M in inside-out membrane patches containing fully phosphorylated WT CFTR. The current stimulation levels of GLPG1837 and ‘358 are indicated as dashed lines. **(b)** Reported structures versus the NMR-determined structures. **(c)** Docked poses of the reported structure of ‘853 (left) versus the NMR-determined structure of ‘853 (right). **(d)** (*S*)-‘853 potentiates WT CFTR currents, while (*R*)-‘853 mildly inhibits them in inside-out patches. Both enantiomers were perfused at 100 □M concentration. Data represent means and SEs of 20 ((*S*)-‘853) or 18 ((*R*)-‘853) patches. Statistical significance relative to no effect was tested by two-tailed Student’s t-test (\*\*\*\**P =* 2.3×10^-9^ for (*S*)-‘853 and *P =* 1.6×10^-7^ for (*R*)-‘853). **(e)** Competition assay showing that the presence of (*R*)-‘853 (100 □M) diminishes the potentiating effect of (*S*)-‘853 (100 □M). **(f)** Dose-response curve of (*S*)-‘853 versus the racemic mixture (±)-‘853, as described for Figure 1e. The *EC_50_* for (*S*)-‘853 was estimated to be 2.1 ± 0.9 μM. Data represent means and SEs of 2-20 patches. 3 mM ATP was used in all panels.

To analyze how these regioisomeric differences affect binding to CFTR, we compared the docking pose of the synthesized ‘853 with its regioisomer in ZINC15 (Figure 2c). Like the ZINC15 regioisomer, the synthesized ‘853 fits well at the potentiator-binding site, forming several identical interactions: its phenyl ring is located in the hydrophobic pocket defined by F312, A436, G438 and R933; the central carbonyl docks to the main chain of F931 via a hydrogen bond; and the indazole ring is poised to stack with F312. The two poses differ in the orientation of the indazole by ∼180° but the docking scores are similar: -34.54 kcal/mol for the synthesized compound and -37.13 kcal/mol for the isomer in the ZINC15 database. This suggests that, despite the side chain difference, the synthesized ‘853 is likely to interact with CFTR at the potentiator-binding site.

Because ‘853 contains a chiral center, the pure enantiomers (*S*)-‘853 and (*R*)-‘853 were synthesized and tested (Figure 2d). Interestingly, although the (*S*)-enantiomer increased the CFTR currents by 1.5-fold, the (*R*)-enantiomer decreased the currents by 0.89-fold (Figure 2d). Furthermore, when tested together, the presence of (*R*)-‘853 diminished the potentiating effect of (*S*)-‘853 in a competition assay (Figure 2e), while dose-response curves confirmed greater current flow through CFTR in the presence of (*S*)-‘853 compared to the racemic mixture (Figure 2f). Together, these data reveal that the two enantiomers of ‘853 have opposite functional effects: while (*S*)-‘853 is a potentiator, (*R*)-‘853 is an inhibitor of CFTR.

### Structural investigation of the *‘*853 binding site

To confirm the docking predicted binding pose of ‘853 and to guide further optimization for affinity and efficacy, we sought to determine the cryo-EM structure of CFTR in complex with ‘853. Using phosphorylated E1371Q CFTR in the presence of saturating ATP-Mg^2+^ (10 mM), we obtained a 3.8 Å reconstruction which revealed clear density at the allosteric potentiator-binding site (Figure 3a and Figure E2). The shape and size of the density are consistent with the molecular structure of ‘853 (Figure 3b). Like ivacaftor and GLPG1837, ‘853 binds to a set of residues near the TM8 hinge region at the protein-membrane interface. Previous mutational scanning experiments identified two polar residues at this location (S308 and Y304) that are critical for ivacaftor and GLPG1837 recognition^14,30^. Alanine substitution of either residue abolished the potentiating effect of (*S*)-‘853 in inside-out patches (Figure 3c), confirming that the modulator discovered by molecular docking specifically engages these residues and interacts with CFTR in the same manner as ivacaftor and GLPG1837.

**Figure 3:**
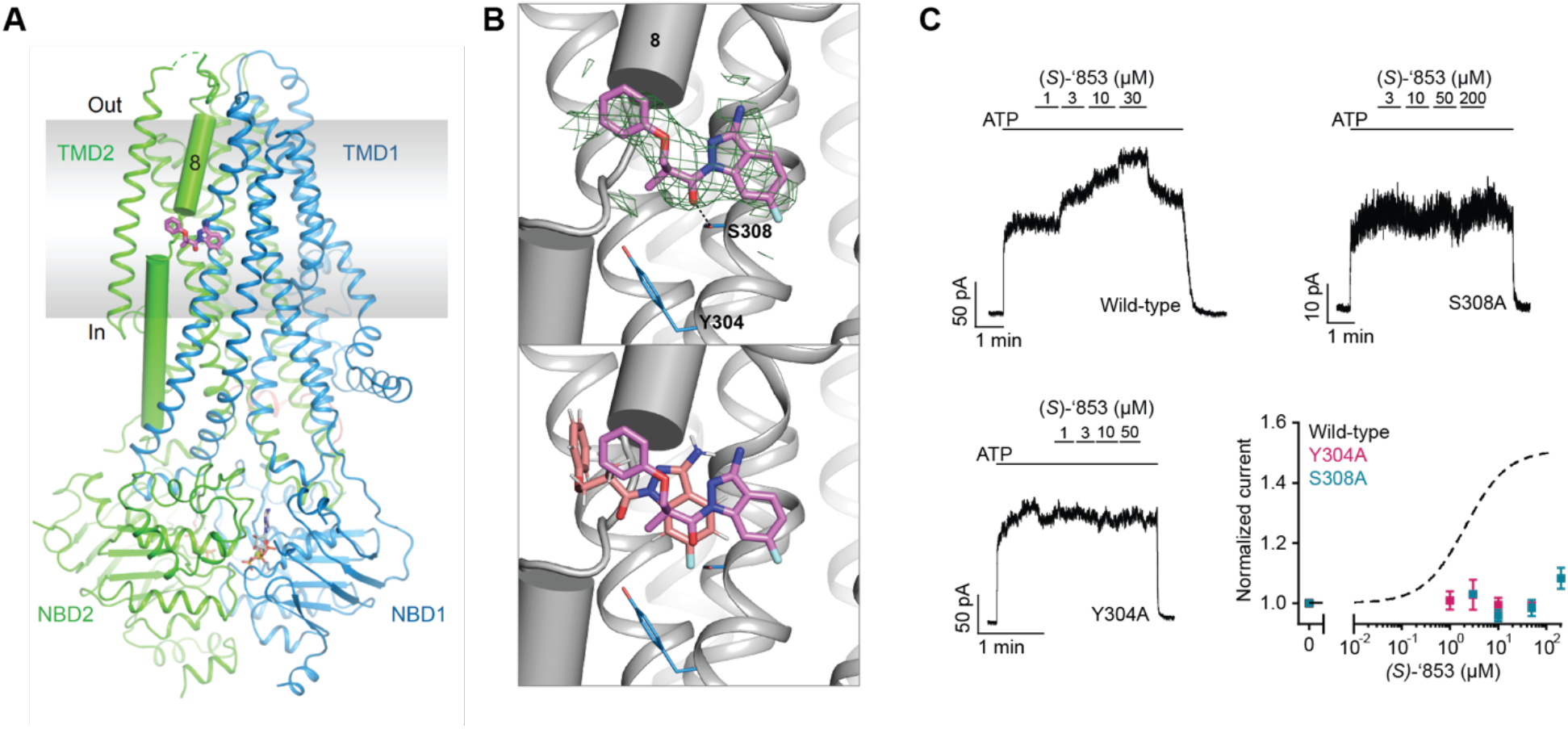
Z1834339853 binds to the same site as ivacaftor and GLPG1837. **(a)** Cryo-EM structure of phosphorylated and ATP-bound CFTR (E1371Q) in complex with ‘853. **(b)** Zoomed-in views of the density of ‘853 (top) and a comparison between the docked pose (salmon) and the cryo-EM pose (magenta). **(c)** Representative macroscopic current traces and dose-response curves of fully phosphorylated WT, S308A, and Y304A CFTR in response to perfusion of (*S*)-‘853 onto inside-out excised membrane patches. 3 mM ATP was used. Each data point represents the mean and SEs determined from 3 to 12 patches.

### Structure-based optimization of the CFTR modulators

With the experimental structure of the CFTR/‘853 complex in hand, and given the intriguing difference between (*S*)*-*‘853 and (*R*)*-*‘853, we carried out further optimization to identify additional potentiators and inhibitors. We used the following medicinal chemistry strategies to synthesize multiple classes of ‘853 analogs (Figure 4a and Extended Figure 3): (1) replacing the fluorine with a larger side chain bearing either hydrogen-bond donor or acceptor functionality (Y); (2) removing the linker methyl or replacing it with a bulkier group (Z), or replacing the linker oxygen with a methylene group (W); (3) removing or modifying the amino group’s hydrogen donor functionality (X); (4) replacing the terminal phenyl ring with aromatic moieties, such as benzofuran or naphthalene, to improve stacking with F312, F316 and F931 or adding side chains to the distal phenyl ring (ARYL).

**Figure 4:**
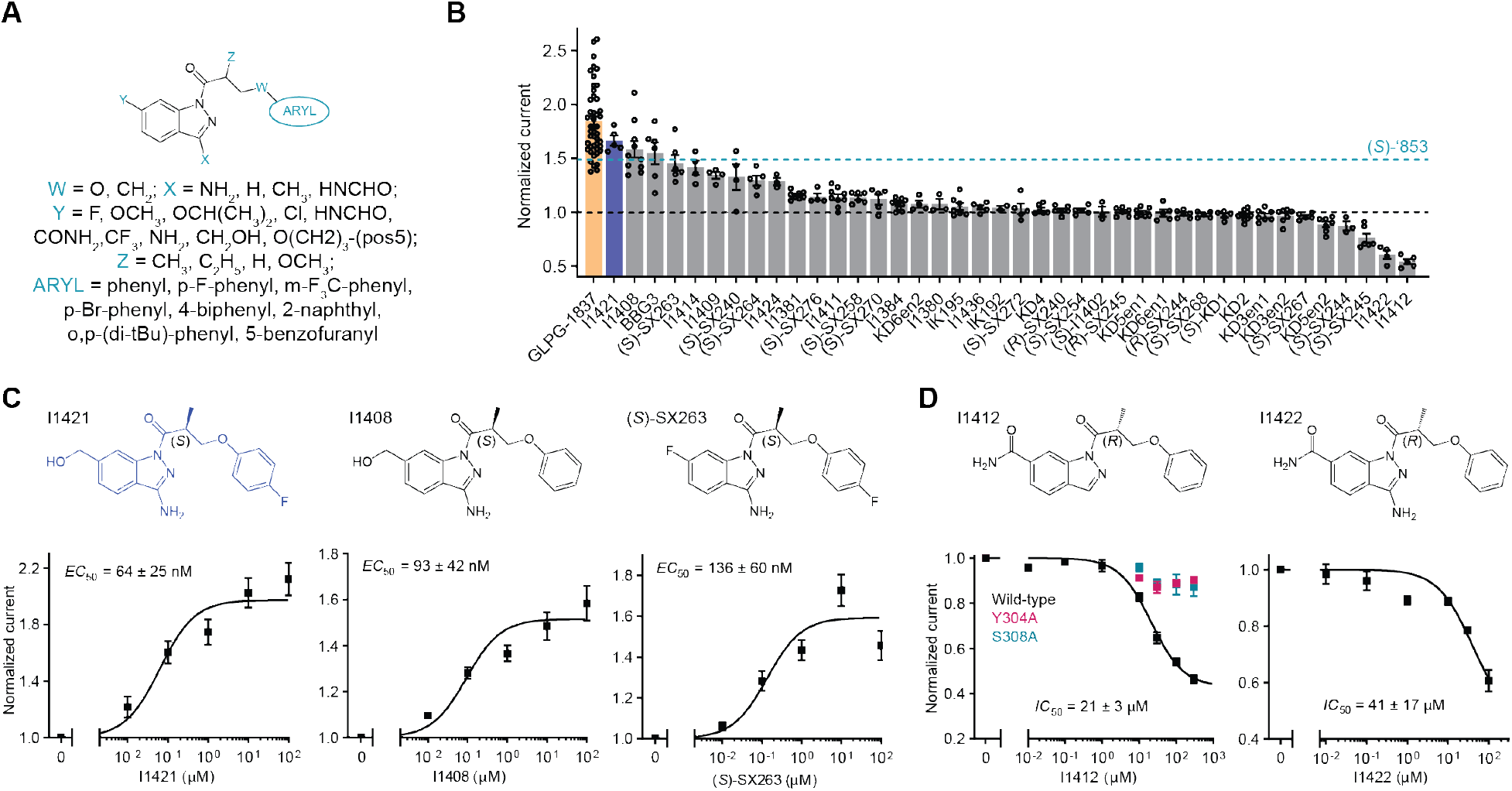
Medicinal chemistry leads to novel CFTR potentiators and inhibitors. **(a)** General formula of newly synthesized ‘853 analogs. **(b)** Effects of ‘853 analogs on currents measured in inside-out excised membrane patches containing fully phosphorylated WT CFTR. Measurements were made with 3 mM ATP. Data represent means and SEs of 2 to 46 patches. **(c)** The structures and dose-response curves of three of the most efficacious potentiators. Data represent means and SEs of 5-11 patches. **(d)** The structures and dose-response curves of the two most efficacious inhibitors. The dose-responses of the Y304A and S308A variants in response to I1412 perfusion were also shown (left). Data represent means and SEs of 2-8 patches.

Each of the designed compounds was docked into the potentiator-binding site *in silico*, and a total of 39 compounds were chosen for synthesis based on their docking score. The efficacies of the synthesized analogs were determined by patch clamp experiments, revealing a continuum of CFTR modulation from inhibition to potentiation (Figure 4b). The most efficacious analogs, which strongly potentiated CFTR, were the ones in which the para-position in the distal phenyl ring was halogenated and/or the indazole fluorine was replaced by a hydroxyl group. The first modification improved the non-polar complementarity in a hydrophobic subsite of the potentiator-binding pocket (Extended Figure 4, right panel), while the replacement of the (*S*)*-*‘853 indazole fluorine with a polar group is predicted to form a hydrogen bond with Q237 and S308 (Figure 4b and Figure E3, Figure 4). Three of the most potent potentiators, I1421, I1408, and (*S*)-SX-263, increased currents through WT CFTR with *EC_50_* values of 64 ± 25 nM, 93 ± 42 nM and 136 ± 60 nM, respectively (Figure 4c).

Of the 39 analogs tested, several were found to inhibit WT CFTR (Figure 4d and Extended Figure 3), including I1412. This compound is an (*R)-*enantiomer of the potentiator I1409, providing a second example of (*S*)- and (*R)-*enantiomers being positive and negative allosteric modulators, respectively (Figure 4b). The most potent inhibitors in this series inhibited WT CFTR with *IC_50_* values of 21 ± 3 µM (I1412) and 41 ± 17 µM (I1422) (Figure 4d). Similar to the potentiator (*S*)-‘853, single alanine substitutions of either of the binding site residues Y304 or S308 eliminated the activity of I1412 (Figure 4d). This medicinal chemistry approach thus led to the identification of a novel potentiator (I1421) with a 30-fold greater potency than (*S*)-‘853, as well as negative modulators that bind to the same site on CFTR.

### I1421 can rescue multiple CF-causing mutants

Finally, we asked whether the strongest potentiator, I1421, could increase the activity of various disease-causing CFTR mutants. Clones carrying ten of these mutations, distributed at different positions in CFTR, were tested using patch-clamp electrophysiology (Figure 5a). The predominant CF mutation, ΔF508, is defective in both folding and gating. Newly synthesized ΔF508 CFTR is largely retained in the endoplasmic reticulum^31^ and the few channels that reach the plasma membrane exhibit little activity^32,33^. I1421 strongly increased currents through ΔF508, indicating that, similar to ivacaftor, it could be used to rescue ΔF508 if used in combination with correctors (Figure 5b, c). Approximately 4% of CF patients carry the G551D mutant, which is expressed on the cell surface but has severe gating defects^34^. In inside-out membrane patches, I1421 increased the activity of G551D by 25-fold (Figure 5b, c). Potentiation of the other eight mutations by I1421 was also comparable to that of GLPG1837 (Figure 5c). These data indicate that I1421 is a strong potentiator that allosterically activates a wide range of CF-causing mutants.

**Figure 5:**
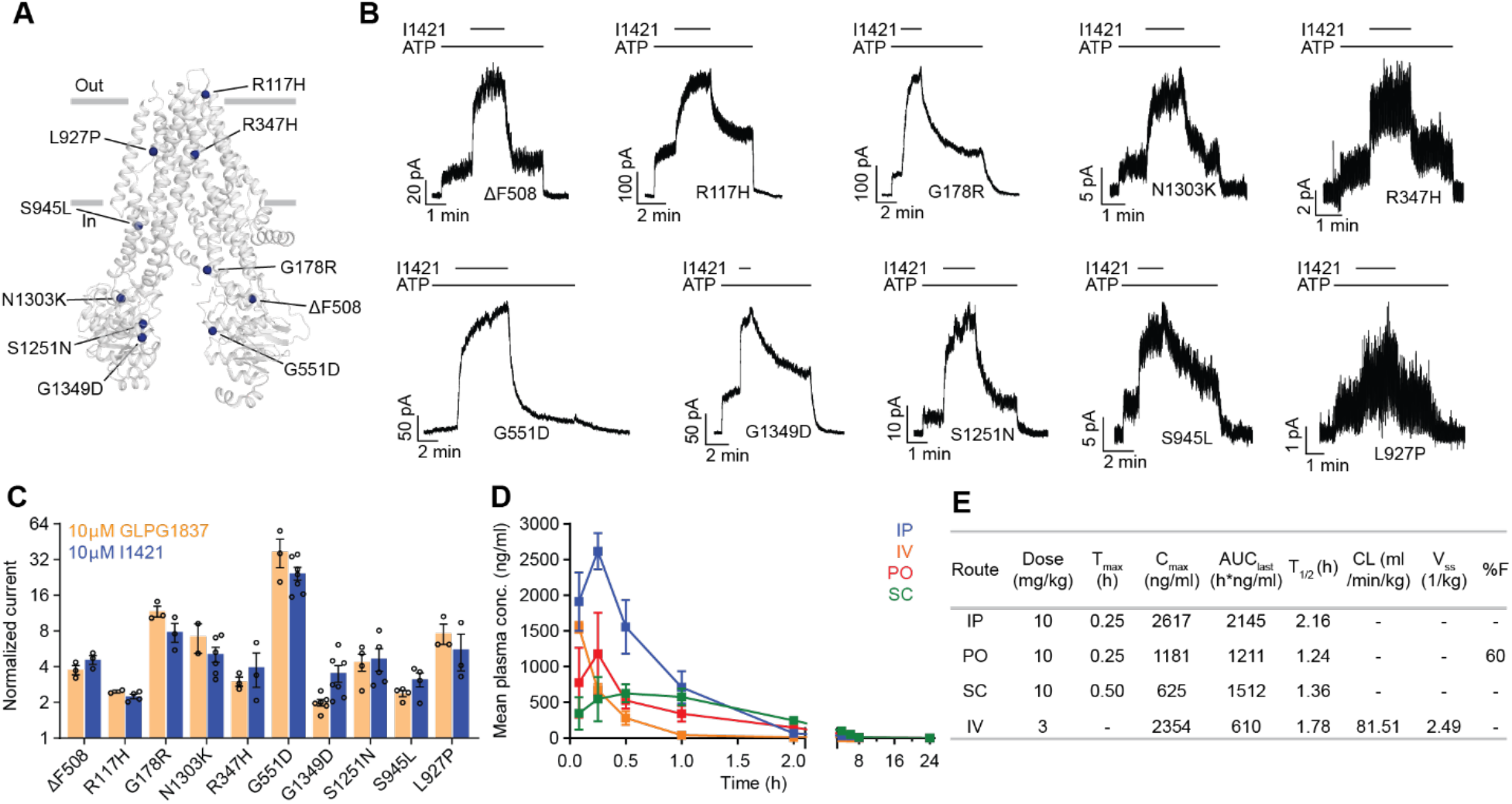
The activity of I1421 against 10 CF-causing mutations. **(a)** The positions of the mutations mapped onto dephosphorylated and ATP-free CFTR (PDB 5UAK). **(b)** Representative macroscopic current traces in response to I1421 (10 μM) perfusion onto inside-out membrane patches excised from CHO cells. 3 mM ATP was used. **(c)** Potentiation activity of I1421 versus GLPG1837. The mean and SE values were determined from 2 to 7 patches. **(d)** Pharmacokinetic analysis of compound I1421. Plasma concentration-time profiles in male C57BL/6N mice following a single subcutaneous (SC), intraperitoneal (IP), per-oral (PO) (dose 10 mg/kg) or intravenous (IV) (3 mg/kg) administration. Data represent means and SDs. **(e)** Selected pharmacokinetic parameters of I1421. C_max_: peak plasma concentration; T_max_: the time when the peak plasma concentration was observed; AUC_last_: the areas under the concentration time curve; T_1/2_: terminal half-life; CL: clearance, V_ss_: steady-state volume of distribution; %F: %bioavailability.

Given these promising results, we investigated the pharmacokinetics of I1421 in C57BL/6 mice following single 10 mg/kg doses administrated by intraperitoneal (IP), oral, or subcutaneous routes (Figure 5d, e). As expected from its cLogP value of 2.5, I1421 was readily formulated in a standard vehicle comprising 10% N-Methyl-Pyrrolidone (NMP), 5% solutol HS-15, and 85% saline. The maximum plasma concentrations (C_max_), observed 15 minutes after IP and oral administration, were 7.4 μM and 3.4 μM, respectively, indicating rapid absorption of the compound. Plasma levels following IP administration were higher than the other two routes of administration, as indicated by the areas under the respective plasma concentration curves (AUC) (Figure 5d, e). To determine oral bioavailability, I1421 was intravenously (IV) injected into a group of nine mice at a dose of 3 mg/kg. Comparison of plasma levels from oral versus IV injection yielded a bioavailability of 60%, an encouraging value given that the compound was not subject to special formulation. In contrast, the low solubility of ivacaftor^26^ has precluded creation of an IV formulation, preventing determination of its oral bioavailability. Low solubility may also affect the non-linear relationship between dose and maximum concentration *in vivo*, causing ivacaftor’s C_max_ to plateau with increasing dose^35,36^, even after extensive formulation optimization. These factors, along with the strong effect of fat-containing food on absorption^9^, may contribute to the variable effects of ivacaftor.

Importantly, I1421 and other molecules in this series are not clinical candidates, have not been optimized for pharmacokinetics, and retain liabilities such as a short oral T_1/2_ of 75 minutes. Nevertheless, these results support the notion that a structure-based approach to CFTR drug discovery can identify molecules with improved physical properties, easier formulation, and improved pharmacokinetics.

## Discussion

Large library docking has been a disruptive innovation in structure-based ligand discovery, in particular for GPCRs^17,19–22^ and enzymes^16,18^. However, such screens, and more broadly structure-based design and discovery, are much less prominent for ion channels. In this study, large library docking against the potentiator-binding site of CFTR revealed novel potentiators and inhibitors with chemotypes unrelated to known modulators and with better physical properties than the widely used drug ivacaftor. Several aspects of this study are worth highlighting. First, despite the shallow, membrane-exposed binding pocket, the docking hit rate (number of molecules active/number experimentally tested) was substantial at 24%. Second, despite their chemical novelty and dissimilarity to previously known modulators, potent potentiators were still found. For example, I1421 had a potency comparable to that of GLPG1837 for all ten CF-causing mutants tested. Third, the molecules that we discovered were substantially more polar than the only FDA-approved potentiating drug, ivacaftor, conferring potential benefits to their behavior *in vivo*. This was borne out in preliminary pharmacokinetic experiments in which I1421 was readily formulated for oral and IV delivery using widely used excipients, resulting in substantial exposure upon oral dosing and a high oral bioavailability of 60%. The good physical properties of this family, and their apparently favorable pharmacokinetics, provide the freedom for further development and optimization. The structure-based approach described here provides multiple points of departure for drug leads for a disease whose treatment remains out of reach for many, and expensive for all. Finally, our study revealed that the allosteric binding site on CFTR can be used to discover not only potentiators but also inhibitors. Such inhibitors may serve as starting points for the development of treatments for secretory diarrhea, the second leading cause of death in children worldwide.

We also wish to draw attention to certain caveats. In particular, we have not tested our potentiators in an animal model, mainly due to the lack of a model system that fully recapitulates the symptoms of human CF^37^. Indeed, CF drug discovery, including the development of ivacaftor, has historically relied on *in vitro* model systems, some of which were used here. In addition, it is important to note that the promising pharmacokinetics of compounds like I1421 are distinct from their pharmacodynamics, so must be viewed cautiously. Moreover, although solubility and oral bioavailability are encouraging, these molecules remain far from optimized and are currently no more than leads. In our docking experiments, the fact that the most potent initial hit represented a different regioisomer to our intended compound tempers the success of the method, even though the two regioisomers occupy similar poses in the binding site and achieve similar docking scores. Finally, although the initial hits sampled a broad range of chemotypes (Figure 1), their potentiating potencies were modest and have yet to be followed up.

Regardless of these caveats, we have made a number of important observations in this study. By docking a large virtual library against the structure of CFTR, we have uncovered 13 diverse chemical scaffolds that are topologically distinct from known ligands and have favorable physical properties. This wide range of chemotypes holds great promise for new drug discovery efforts against this crucial target. Indeed, the chemical novelty of these compounds supported our discovery of CFTR inhibitors that likely bind to the same allosteric site, and which have the potential to become lead molecules for the treatment of secretory diarrhea. By determining the structure of one of the new ligands in complex with CFTR, we have provided a template for further optimization of this series and for other drug discovery efforts. Encouragingly, the beneficial physical properties of I1421 resulted in favorable pharmacokinetics during animal dosing. In conclusion, our study has revealed that combining structural biology with exploration of a large chemical space can reveal new lead molecules for what remains a crucial drug target for CF and secretory diarrhea.

## Methods

### Ultra-large scale virtual ligand screening

The recently determined CFTR/ivacaftor cryo-EM structure (PDB: 6O2P;^14^) was used to prospectively screen ∼155 million “lead-like” molecules (molecular weight 300-350 Da and logP ≤ 3.5), from the ZINC15 database (http://zinc15.docking.org/^24^), using DOCK3.7^25^. DOCK3.7 fits pregenerated flexible ligands into a small molecule binding site by superimposing atoms of each molecule on local hot spots in the site (“matching spheres”), representing favorable positions for individual ligand atoms. Here, 45 matching spheres were used, drawn from the experimentally determined pose of ivacaftor. The resulting docked ligand poses were scored by summing the channel–ligand electrostatics and van der Waals interaction energies and corrected for context-dependent ligand desolvation^38,39^. Channel structures were protonated using Reduce^40^. Partial charges from the united-atom AMBER force field were used for all channel atoms^41^. Potential energy grids for the different energy terms of the scoring function were precalculated based on the AMBER potential^41^ for the van der Waals term and the Poisson–Boltzmann method QNIFFT73^42,43^, for electrostatics. Context-dependent ligand desolvation was calculated using an adaptation of the generalized-Born method^38^. Ligands were protonated with Marvin (version 15.11.23.0, ChemAxon, 2015; https://www.chemaxon.com), at pH 7.4. Each protomer was rendered into 3D using Corina (v.3.6.0026, Molecular Networks GmbH; https://www.mn-am.com/products/corina) and conformationally sampled using Omega (v.2.5.1.4, OpenEye Scientific Software; https://www.eyesopen.com/omega). Ligand atomic charges and initial desolvation energies were calculated as described^24^. In the docking screen, each library molecule was sampled in about 3,727 orientations and, on average, 421 conformations. The best scoring configuration for each docked molecule was relaxed by rigid-body minimization. Overall, over 63 billion complexes were sampled and scored; this took 76,365 core hours– spread over 1000 cores, or slightly more than 3 days.

To identify novel and diverse chemotypes, the top 100,000 scoring molecules were clustered by 2D similarity using ECFP4 fingerprints and a Tanimoto Coefficient (Tc) cutoff of 0.5. The top ranked 1,000 cluster heads were visually inspected in their docked poses to remove molecules that are conformationally strained or with unsatisfied hydrogen-bond acceptors or donors. Topologically diverse molecules that adopted favorable geometries and formed specific interactions with the key potentiator-binding residues S308, Y304, F312, and F931 (PDB: 6O2P and 6O1V), were prioritized from among the top 1000 docking-ranked molecules.

Ultimately, 58 compounds, each from a different chemotype family, were selected for experimental evaluation.

### Synthesis of molecules

Fifty-three molecules prioritized for purchasing were synthesized by Enamine for a total fulfilment rate of 91%. Compounds were sourced from the Enamine REAL database (https://enamine.net/compound-collections/real-compounds). The purities of active molecules synthesized by Enamine were at least 90% and typically above 95%. For compounds synthesized in house purities were at least 95%. The detailed chemical synthesis can be found in the Chemical Synthesis and analytical investigations section.

### Hit Optimization

Potential analogs of the hit compound Z2075279358 were identified through a combination of similarity and substructure searches of the ZINC database^24^. Potential analogs were docked to the CFTR small molecule binding site using DOCK3.7^38^. As was true in the primary screen, the resulting docked poses were manually evaluated for specific interactions and compatibility with the site, and prioritized analogs were acquired and tested experimentally.

### Chemical Synthesis and analytical investigations

The library from Enamine were synthesized taking advantage of synthesis protocols previously published (also see below)^44,45,46^.

The synthetic procedures for pure enantiomers of ‘853 and its analogs were depicted in supplementary information. Furthermore, the ^1^H- and ^13^C-NMR spectra, the HPLC-chromatograms demonstrating purity > 95% and proving optical purity are included.

### Pharmacokinetic Study

Pharmacokinetic experiments of I1421 were performed by Sai Life Sciences (Hyderabad, India) at an AAALAC accredited facility in accordance with the Sai Study Protocol SAIDMPK/PK-22-12-1306 and PK-22-11-1117. International guidelines for animal experiments were followed. Plasma pharmacokinetics of compound I1421 was measured after a single 10 mg/kg dose, administered intraperitoneal (IP), per-orally (PO) or subcutaneously (SC). Plasma samples were also collected from mice after a single 3 mg/kg intravenous (IV) dose injection to determine oral bioavailability. Both doses were formulated in 10% NMP:5% Solutol HS-15: 85% saline (v/v/v). Testing was done in healthy male C57BL/6 mice (8-10 weeks old) weighing between 25 ± 5 g (procured from Global, India). Three mice were housed in each cage. Temperature and humidity were maintained at 22 ± 3 °C and 30-70%, respectively and illumination was controlled to give a sequence of 12 h light and 12 h dark cycle. Temperature and humidity were recorded by an auto-controlled data logger system. All animals were provided laboratory rodent diet (Envigo Research private Ltd, Hyderabad). Reverse osmosis water treated with ultraviolet light was provided *ad libitum*.

For the 10 mg/kg (IP, PO and SC) study, a total of twenty-seven mice were used. Animals in Group 1 (n=9) were administered IP, animals in Group 2 (n=9) were administered PO and animals in Group 3 (n=9) were administered SC with solution formulation of I1421. For the 3 mg/kg (IV) study, animals in Group 1 (n=9) were administered IV with the same solution formulation. Blood samples (approximately 60 μL) were collected from the retro orbital plexus of a set of three mice at 0.083, 0.25, 0.5, 1, 2, 4, 6, 8 and 24 h. Immediately after blood collection, plasma was harvested by centrifugation at 10000 rpm, 10 min at 4 ^0^C and samples were stored at -70±10 °C until bioanalysis. All samples were processed for analysis by protein precipitation method and analyzed with fit-for-purpose LC-MS/MS method (LLOQ = 1.02 ng/mL for plasma). The pharmacokinetic parameters were estimated using non-compartmental analysis tool of Phoenix® WinNonlin software (Ver 8.3).

#### Sample Extraction Procedure

10 µL of study sample plasma or spiked plasma calibration standard was added to individual pre-labeled micro-centrifuge tubes followed by 100 µL of internal standard prepared in Acetonitrile (Cetrizine, 50 ng/mL) except for a blank, where 100 µL of Acetonitrile was added. Samples were vortexed for 5 minutes. Samples were centrifuged for 10 minutes at a speed of 4000 rpm at 4 °C. Following centrifugation, 200 µL of clear supernatant was transferred into 96 well plates and analyzed using LC-MS/MS.

#### Data Analysis

Peak plasma concentration (C_max_) and time for the peak plasma concentration (T_max_) were the observed values. The areas under the concentration time curve (AUC_last_ and AUC_inf_) were calculated by the linear trapezoidal rule. The terminal elimination rate constant, ke was determined by regression analysis of the linear terminal portion of the log plasma concentration-time curve. The terminal half-life (T_1/2_) was estimated at 0.693/ke. Mean, SD and %CV was calculated for each analyte. The bioavailability is calculated by 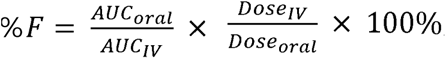, with AUC_oral_ as 1,211 h*ng/ml and AUC_IV_ as 610 h*ng/ml with 10 mg/kg and 3 mg/kg dosing respectively.

### Cell culture

Sf9 cells were cultured in Sf-900 II SFM medium (GIBCO) supplemented with 5% (v/v) fetal bovine serum (FBS) and 1% (v/v) Antibiotic-Antimycotic. HEK293S GnTl^-^ cells were cultured in Freestyle 293 (GIBCO) supplemented with 2% (v/v) FBS and 1% (v/v) Antibiotic-Antimycotic. Chinese hamster ovary (CHO) cells were cultured in DMEM-F12 (ATCC) supplemented with 10% (v/v) FBS and 1% (v/v) Antibiotic-Antimycotic. CFBE41o-cells expressing F508del-CFTR and the fluorescent protein eYFP-H148Q/I152L/F46L were cultured in MEM-alpha with 10% (v/v) FBS, 1% (v/v) Pen-Strep, 2 mg/mL Puromycin and 0.75 mg/mL G418.

### Mutagenesis

All mutations were introduced using QuikChange Site-Directed Mutagenesis System (Stratagene).

### Protein expression and purification

CFTR E1317Q was expressed and purified as described ^15,47^. In summary, bacmids carrying CFTR E1317Q construct were generated in *E. Coli* DH10Bac cells (Invitrogen). Recombinant baculoviruses were produced and amplified in Sf9 cells. Proteins were expressed in HEK293S GnTl^-^ cells infected with 10% (v/v) baculovirus at a density of 3×10^6^ cells/ml. Cells were induced with 10 mM sodium butyrate 12 hours after infection and cultured at 30 °C for another 48 hours before harvesting.

For protein purification, cell membranes were solubilized in buffer containing 1.25% (w/v) 2,2-didecylpropane-1,3-bis-β-D-maltopyranoside (LMNG) and 0.25% (w/v) cholesteryl hemisuccinate (CHS). Protein was purified via its C-terminal green fluorescence protein (GFP) tag using GFP nanobody-coupled Sepharose beads (GE Healthcare) and eluted by removing the GFP tag with PreScission Protease. The sample was phosphorylated using protein kinase A (NEB) and then further purified on size exclusion chromatography.

### EM data acquisition

The phosphorylated E1371Q sample (5.5 mg/mL in 0.06% (w/v) digitonin) was incubated with 10 mM ATP and MgCl_2_ plus 200 µM Z1834339853 on ice for 15 min. 3 mM fluorinated Fos-choline-8 was added to the sample immediately before freezing on to Quantifoil R1.2/1.3 400 mesh Au grids using a Vitrobot Mark IV (FEI). EM images were collected on a 300 kV Titian Krios (FEI) with a K2 Summit detector (Gatan) in super-resolution mode using SerialEM. The defocus ranged from 1 to 2.5 μm and the dose rate was 8 e-/pixel/sec. The data were collected in two sessions, with 3,088 and 3,879 movies collected.

### EM data processing

The images of the CFTR/Z1834339853 dataset were first corrected for gain reference and binned by 2 to obtain a physical pixel size of 1.03 Å. Beam-induced sample motion was corrected using MotionCor2 ^48^. CTF estimation was performed using Gctf ^49^. Particles were automatically picked by Relion ^50^.

Two datasets were collected. For the first dataset, a total number of 1,346,123 particles were extracted from 3,879 movies and were subjected to two rounds of 2D classification. 267,967 particles and 190,253 particles were selected and combined. Duplicated molecules were removed to yield 313,972 particles for 3D classification. The best classes from the last 4 iterations of 3D classification were combined; duplicated particles were removed to yield 254,123 particles. For further classification, the best map from 3D classification was low pass filtered to 8, 16, 24, 32, 40, 48 Å and used as reference maps. Local searches were performed at 7.5 and 3.75 degrees for 25 iterations respectively. Thorough searches were done by setting–maxsig 5. Such classification led to 6 classes, with the best class including 40% of the 254,123 particles and having a resolution of 7.92 Å. The best class (102,891 particles) was polished and refined to yield a reconstruction of 4.1 Å. Postprocessing did not improve the resolution further. In the second dataset, a total number of 801,032 particles from 3,088 movies were used for 2D classification, and subsequently 602,374 particles were used for 3D classification in RELION2^51^. 3D classification yielded a best class with 200,815 particles. These particles were further classified with the same six reference models as the previous dataset. The best class contained 106,212 particles and were polished and refined to a 4.9 Å map. The polished particles from the two datasets were combined and imported to cryosparc for non-uniform refinement to yield a final map of 3.8 Å.

### Model building and refinement

The model building and refinement of CFTR were carried out as described^15^. In brief, the data was randomly split into two halves, one half for model building and refinement (working set) and the other half for validation (free set). The model was built in Coot^52^ and refined in reciprocal space with Refmac^53,54^. MolProbity^55,56^ was used for geometry validation and Blocres^57^ was used for local resolution estimation. Final structures contain residues 1–390 of transmembrane domain 1 (TMD1); 391–409, 437–637 of nucleotide binding domain (NBD1); 845–889, 900–1173 of transmembrane domain 2 (TMD2); 1202–1451 of nucleotide binding domain 2 (NBD2); and 17 residues of R domain (built as alanines). Figures were generated with PyMOL and Chimera^58^.

### Inside-out patch clamp recording

All CFTR constructs used for electrophysiology were cloned into the BacMam expression vector. A GFP tag was fused to the C-terminus of CFTR for visualization of transfected cells. Chinese hamster ovary (CHO) cells were plated at 0.4×10^6^ cells per 35-mm dish (Falcon) 12 hours before the transfection. For each dish, cells were transfected with 1 μg BacMam plasmids using Lipofectamine 3000 (Invitrogen) in Opti-MEM media (Gibco). After 12 hours of transfection, media was exchanged to DMEM: F12 (ATCC) supplemented with 2% (v/v) FBS and the cells were incubated at 30 °C for 2 days before recording.

Macroscopic currents were recorded in inside-out membrane patches excised from CHO cells, using buffer compositions, and recording parameters as described ^28^. The culture media were first changed to a bath solution consisting of 145 mM NaCl, 2 mM MgCl_2_, 5 mM KCl, 1 mM CaCl_2_, 5 mM glucose, 5 mM HEPES, and 20 mM sucrose, pH 7.4 with NaOH. The pipette solution contained 140 mM *N*-Methyl-D-glucamine (NMDG), 5 mM CaCl_2_, 2 mM MgCl_2_, and 10 mM HEPES (pH 7.4 with HCl). The perfusion solution contained 150 mM NMDG, 2 mM MgCl_2_, 1 mM CaCl_2_, 10 mM EGTA, and 8 mM Tris (pH 7.4 with HCl). Borosilicate micropipettes (OD 1.5 mm, ID 0.86 mm, Sutter) were pulled to 2-5 MΩ resistance. After a gigaseal was formed, inside-out patches were excised and exposed to 25 units/ml bovine heart PKA (Sigma-Aldrich, P2645) and 3 mM ATP to phosphorylate CFTR. Currents were recorded at -30 mV and 25 LC using an Axopatch 200B amplifier, a Digidata 1550 digitizer and pCLAMP software (Molecular Devices). The recordings were low-pass filtered at 1 kHz and sampled at 20 kHz. All displayed recordings were further low-pass filtered at 100 Hz. Data were analyzed with Clampfit, GraphPad Prism, and OriginPro. The effect of the compunds were reported as normalized current which was defined as the current level after application of the tested compound divided by the current level before application of the compound.

### YFP fluorescent assay

The experiments were performed as described in the literature^59^. In brief, 50,000 cells (not cultured beyond passage 10) were seeded per well in 96-well plates (GBO 655866) 48 hours before the start of assay, and incubated at 37 °C. 24 hours before the start of the assay, the CFTR corrector lumacaftor was added at 1 µM in a final culture volume of 200 µL. The plates were incubated at 30 °C for 24 hours. On the day of experiments, the cells were first washed three times with 200 µL PBS (HyClone SH30264.02), and then incubated with 20 µM forskolin and 1 µM test compound for 25 minutes in a 60 µL total volume.The plate was then transferred to a microplate reader for a 14 second fluorescence reading, split into a 2 second baseline reading, rapid injection of 165 µL iodide containing solution (PBS with 137 mM Cl^-^ replaced by I^-^), and thena 12 second reading. Data were normalized to initial background-subtracted fluorescence. I^-^ influx rate was determined by fitting the final 11 seconds of data for each well with an exponential function to extrapolate the initial rate of fluorescence quenching (dF/dt). Relative potentiation is defined as the initial rate of fluorescence decay with test compound divided by the initial rate of fluorescence decay without test compound.

### Quantification and Statistical Analysis

GraphPad Prism 9 was used to fit the dose response curves of CFTR potentiators and inhibitors and to calculate *EC_50_*s. Statistical significance was calculated by two-tailed Student’s t-test in Prism 9.

## Data availability

All data are available in the main text, the supplementary materials, the listed Protein Data Bank (PDB) file and the Electron Microscopy Data Bank (EMDB) files. The 3D cryo-EM density map of CFTR-‘853 complex generated in this study has been deposited with accession code EMD-40207. The coordinate of CFTR-**‘**853 complex has been deposited with PDB accession code 8GLS. The identities of compounds docked in this study are freely available from the ZINC15 and ZINC20 databases (https://zinc15.docking.org/ and https://zinc20.docking.org/), and active compounds may be purchased from Enamine. DOCK3.7 is freely available for noncommercial research (https://dock.compbio.ucsf.edu/DOCK3.7/). A web-based version is freely available to all (https://blaster.docking.org/).

## Acknowledgments

We thank M. Ebrahim and J. Sotiris at Rockefeller’s Evelyn Gruss Lipper Cryo-Electron Microscopy Resource Center for assistance in data collection, Yiming Niu and Chen Zhao for help with cryo-EM data processing, and Iris Torres and Bilge Bebek for technical assistance. We would also like to thank Luis J. Galietta for sharing the CFBE41o^-^ cells expressing F508del-CFTR and the fluorescent protein EYFP-H148Q/I152L/F46L.

## Funding

This work is supported by the Howard Hughes Medical Institute (to J.C.), R35GM122481 (to B.K.S.), GM71896 (to J.J.I.) and the Deutsche Forschungsgemeinschaft (to P.G.).

## Author Contributions

A.L.K. performed molecular docking; F.L., N.S.O, and H.L.P. tested compounds from the initial screen; J.L. carried out majority of the patch-clamp experiments; N.S.O. prepared sample and collected data of the CFTR/’853 complex; F.L. determined the structure; I.S.K., Y.S.M., and J.J.I. synthesized the first panel of ligands; S.T., K.D., and J.E. synthesized and chemically characterized new CFTR potentiators and inhibitors. P.G., B.K.S., and J.C. supervised the study. F.L., A.L.K., J.L., J. E., B.K.S., and J.C. wrote the manuscript with input from all authors.

## Competing interests

B.K.S. and P.G. are founders of Epiodyne. B.K.S. is a co-founder of BlueDolphin and of Deep Apple Therapeutics, as is J.J.I., and serves on the SRB of Genentech and on the SABs of Vilya Therapeutics and Umbra Therapeutics, and consults for Great Point Ventures and for Levator Therapeutics. A patent on the discovery of positive and negative allosteric regulators for CFTR has been filed. The authors declare no other competing interests.

## Additional information

Extended data and supplementary information are available for this manuscript.

---

**Extended Data Figure 1.**
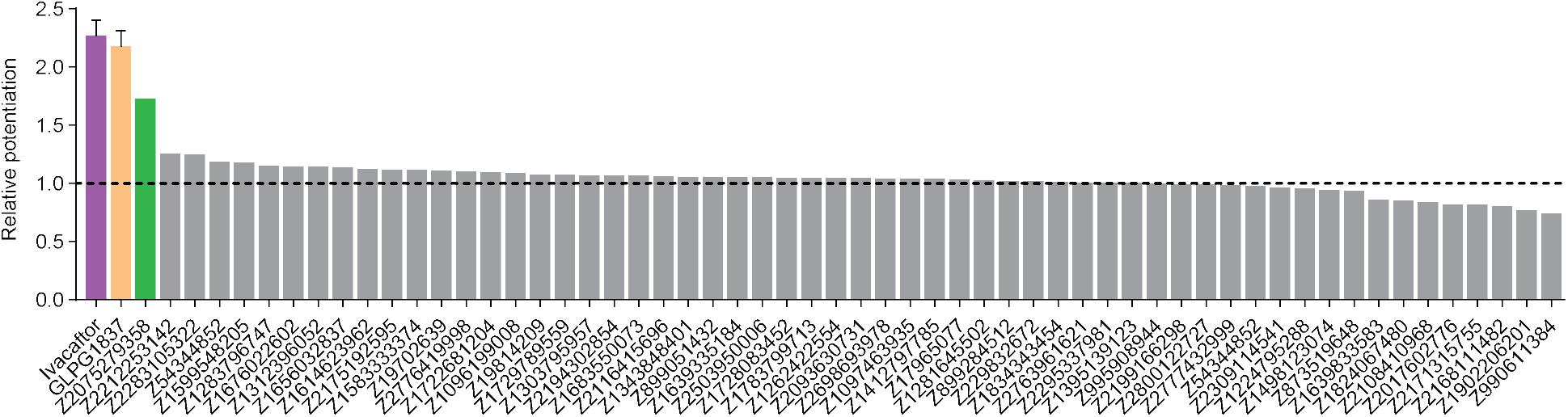
Z2075279358 increased the ion flux rate in a bronchial epithelial cell line derived from a patient homozygous for ΔF508 (CFBE41o^-^ cells expressing ΔF508-CFTR and the fluorescent protein eYFP-H148Q/I152L/F46L^59^). Cells were treated with 20 μM forskolin +/-1 µM test compound. The relative potentiation was calculated as the ratio of flux rates with and without compound. Data points represent a single measurement for each test compound. For ivacaftor and GLPG1837 data represent means and SEs for 9 or 2 measurements, respectively.

**Extended Data Figure 2.**
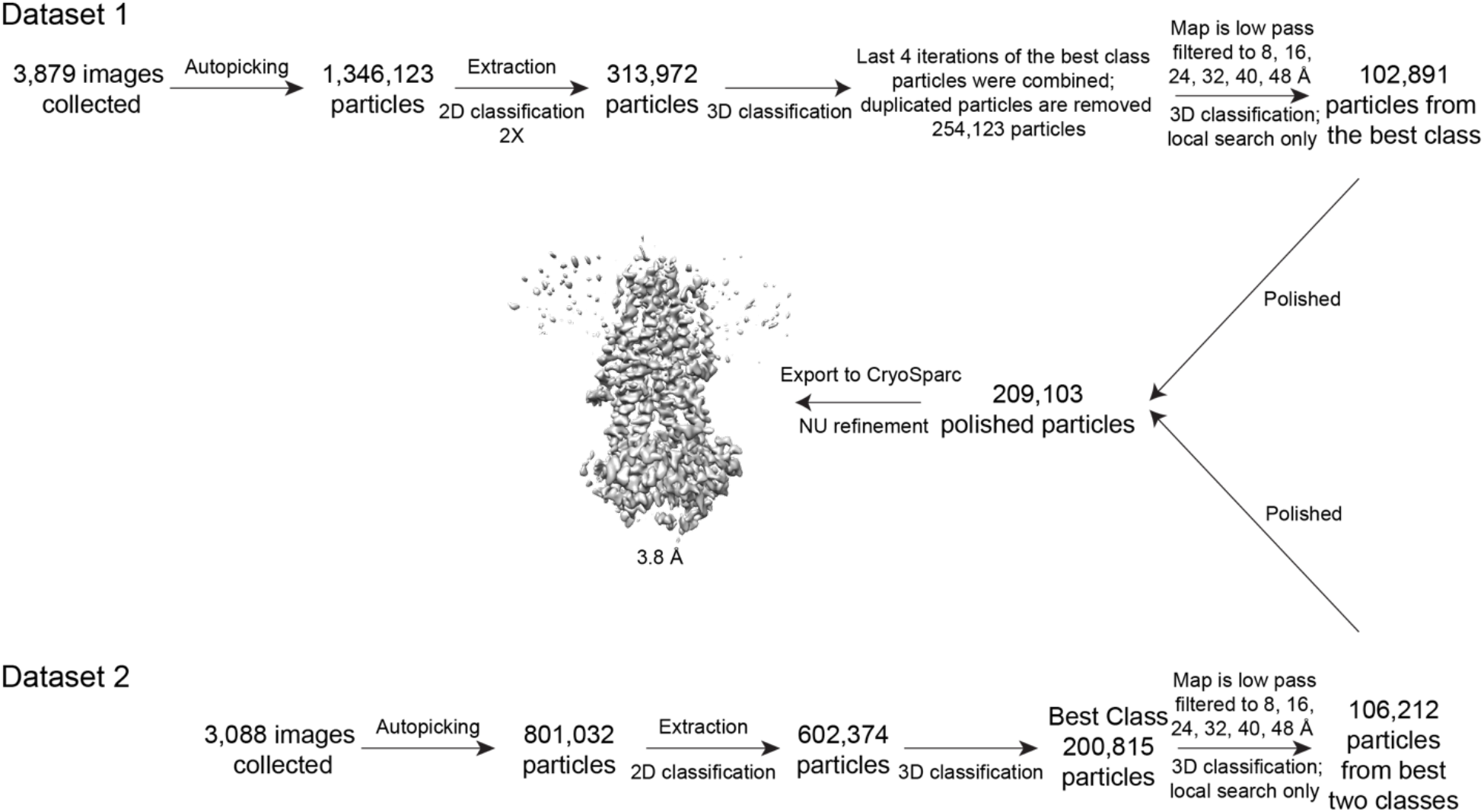
Cryo-EM reconstructions of the CFTR-‘853 complex. Summary of the image processing procedure.

**Extended Data Figure 3.**
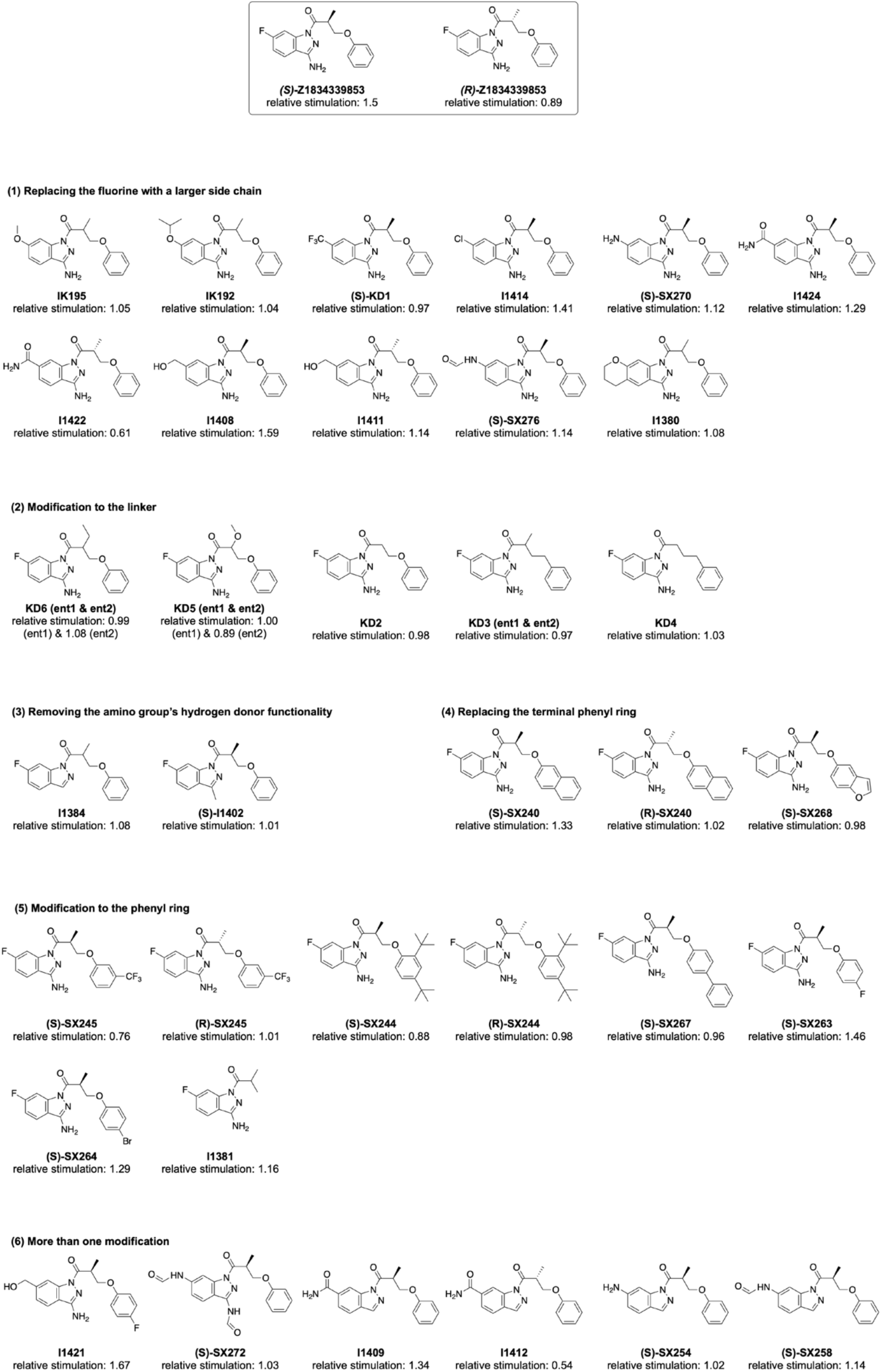
SAR for optimization of compound ‘853.

**Extended Data Figure 4.**
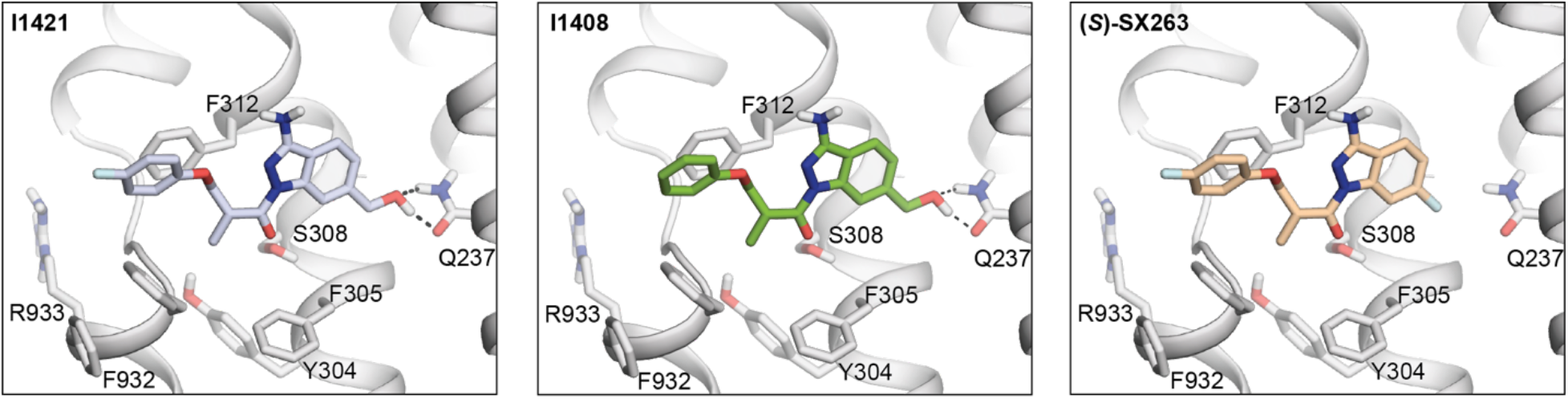
Docked poses of I1421, I1408, and *(S)*-SX263.

**Extended Data Table 1.**
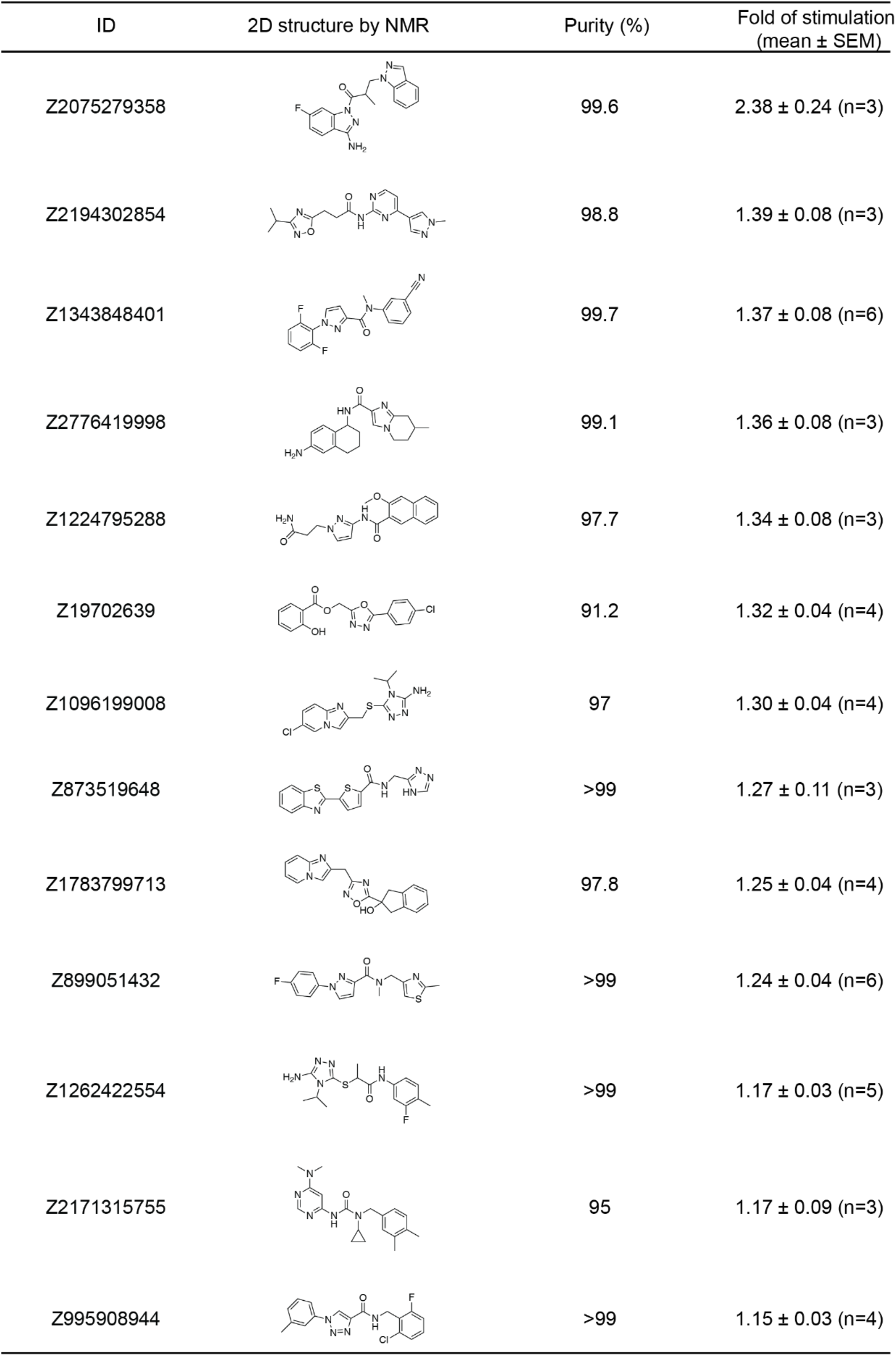
Potentiating activities of the 13 CFTR primary hits.

| ID | 2D structure by NMR | Purity (%) | Fold of stimulation<br>(mean ± SEM) |
| --- | --- | --- | --- |
| Z2075279358 | | 99.6 | 2.38 ± 0.24 (n=3) |
| Z2194302854 | | 98.8 | 1.39 ± 0.08 (n=3) |
| Z1343848401 | | 99.7 | 1.37 ± 0.08 (n=6) |
| Z2776419998 | | 99.1 | 1.36 ± 0.08 (n=3) |
| Z1224795288 | | 97.7 | 1.34 ± 0.08 (n=3) |
| Z19702639 | | 91.2 | 1.32 ± 0.04 (n=4) |
| Z1096199008 | | 97 | 1.30 ± 0.04 (n=4) |
| Z873519648 | | >99 | 1.27 ± 0.11 (n=3) |
| Z1783799713 | | 97.8 | 1.25 ± 0.04 (n=4) |
| Z899051432 | | >99 | 1.24 ± 0.04 (n=6) |
| Z1262422554 | | >99 | 1.17 ± 0.03 (n=5) |
| Z2171315755 | | 95 | 1.17 ± 0.09 (n=3) |
| Z995908944 | | >99 | 1.15 ± 0.03 (n=4) |

**Extended Data Table 2.**
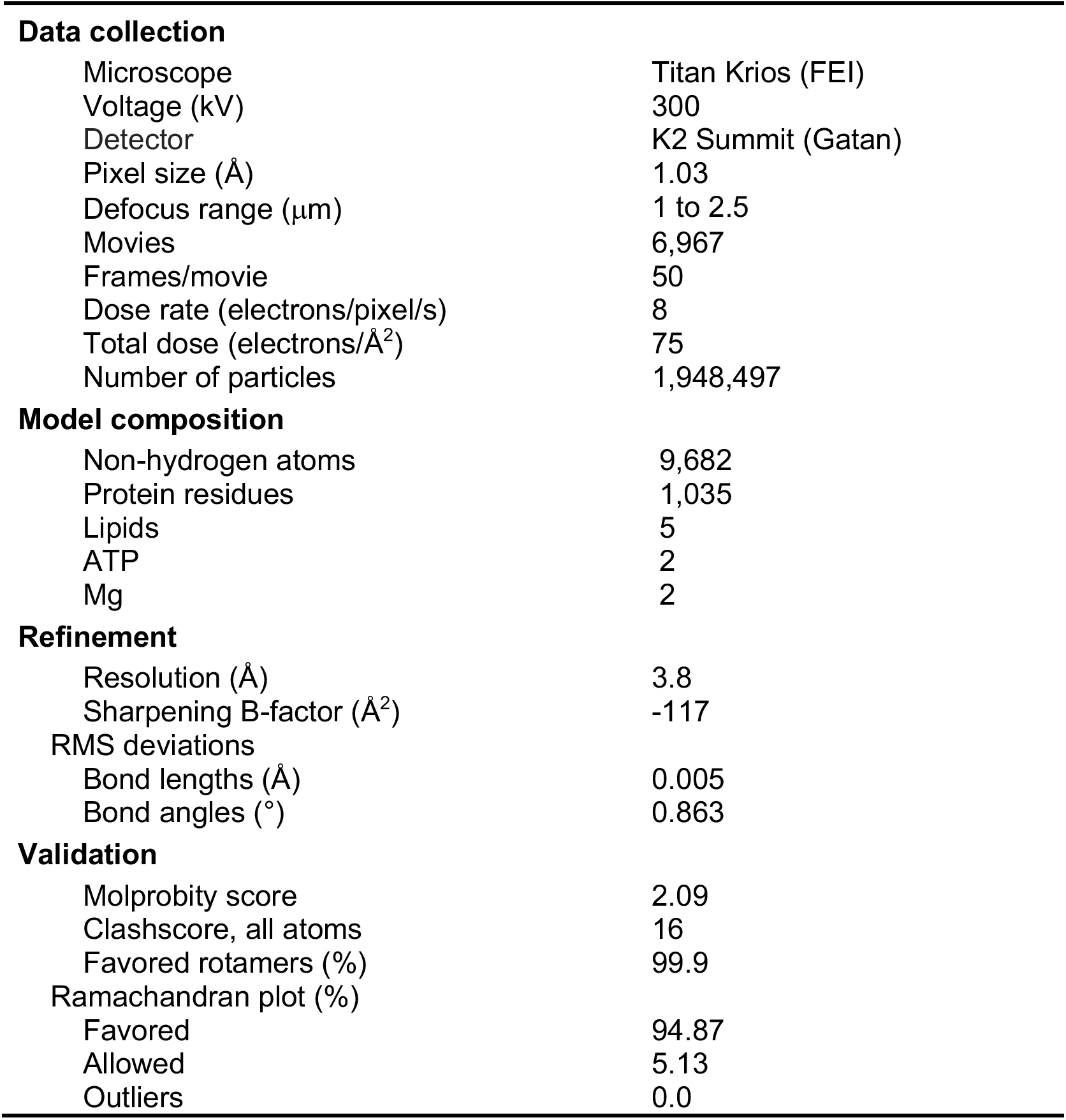
Summary of EM data and structure refinement statistics for CFTR in complex with ‘853.

## Notes

### Summary of Updates

The acknowledgment is modified to reflect changes on the funding information

