## supplemental information for "Structure-based discovery of CFTR potentiators and inhibitors"

**Supplementary Information**

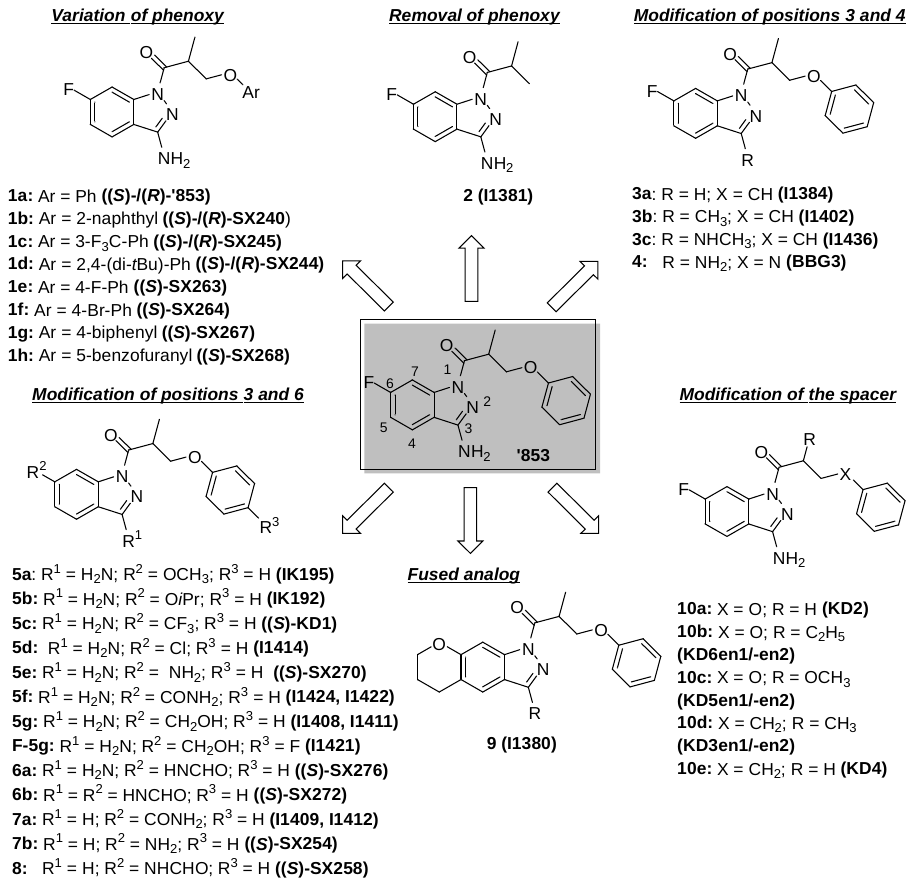

Overview over the synthesized **‘853**-derivatives

The syntheses of the 1-acyl-3-aminoindazole derivatives of type 1 were performed as shown in scheme S1. Starting from the corresponding phenol derivatives, a recently reported protocol comprising deprotonation with NaH and subsequent S_N_2-reaction with enantiopure (*R*)- and (*S*)-3-bromo-2-methyl propanol, ^1^ respectively, allowed to obtain the enantiopure (*R*)- and (*S*)-3-aryloxy-2-methylpropanol derivatives **11a-h** and **ent11a-d**, respectively. Both phenoxy-derivatives **11a/ent11a** ^1^ and the fluoro derivative **11e** ^2^ have been described in the literature.

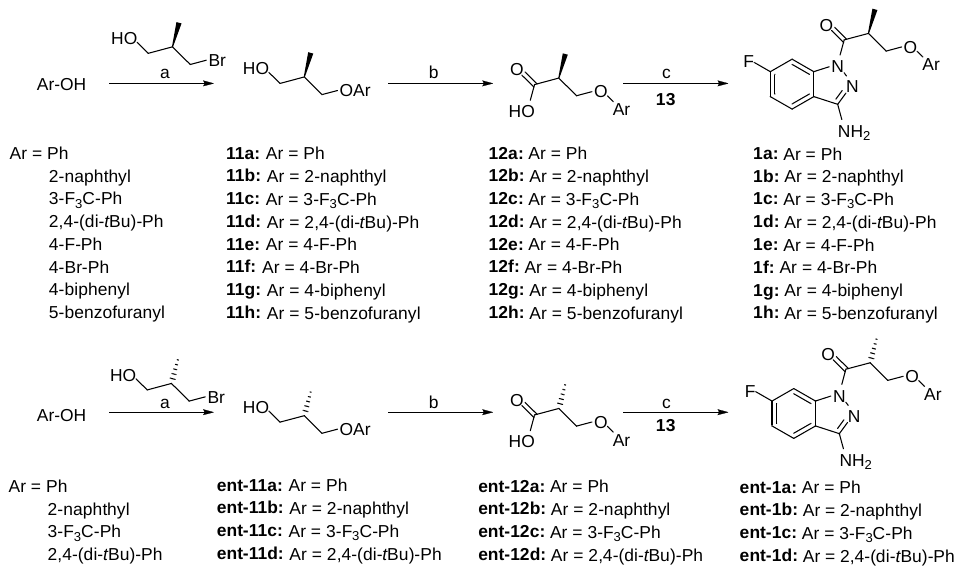

**Scheme S1.** Synthesis of the enantiopure derivatives of **‘853**, modified at the phenoxy site, reagents and conditions: (a) 1. NaH, DMF, 0 °C, 30 min, 2. (*R*)-3-bromo-2-methylpropan-1-ol or (*S*)-3-bromo-2-methylpropan-1-ol, DMF, rt, 18-31 h, (50-72% crude); (b) 1. CrO_3_, H_2_SO_4_, H_2_O, acetone, 0 °C, 3-5 h, 2. *i*PrOH, 0 °C -> rt (48-64%, crude); (c) 1. EDC × HCl, HOAt, DMF, rt, 2. 6‑fluoro-1*H*-indazole-3-amine (**13**), rt, 1-3 h (15-53%).

Jones oxidation gave the corresponding carboxylic acid derivatives **12a-h** and **ent12a-d**. It is worthy of note, that the phenoxy derivatives **12a/ent12a** ^1^, the racemic naphthyl derivative of **12b** ^3^ and the biphenyl derivative **12g** ^4^ have been previously published.

The coupling reactions using 1-ethyl-3-(3-dimethylaminopropyl)carbodiimide hydrochloride (EDC × HCl) and 1-hydroxy-7-azabenzotriazole (HOAt) with 6-fluoro-1*H*-indazole-3-amine (**13**) proceeded with high regioselectivity to give access to the 1-acyl-3-aminoindazole derivatives **1a-h** and **ent1a-d** in 15-53% yield. Investigation of the alcohol derivatives **11a-h** / **ent11a-d** and the target compounds **1a-h** / **ent1a-d** by chiral HPLC proved, that the synthetic pathway allowed to obtain test compounds with high enantiopurity.

The fluoroindazole derivative renouncing the phenoxy moiety **2** was obtained in 90% yield by reacting isobutyryl chloride with HOAt, followed by the addition of 6-fluoro-1*H*-indazole-3-amine (**13**) (scheme S2).

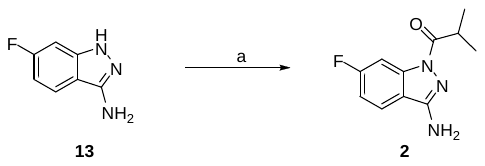

**Scheme S2.** Synthesis of the **‘853** derivative renouncing the phenoxy site, reagents and conditions: (a) 1. isobutyryl chloride, HOAt, DIPEA, DMF, 0 °C, 1 h, 2. 6‑fluoro-1*H*-indazole-3-amine (**13**), microwave irradiation (90%).

The derivatives of **‘853** modified in position 3 of the indazole scaffold were prepared as displayed in scheme S3. The 3-desamino derivative **3a** was synthesized as a racemate starting from 6-fluoro-1*H*-indazole (**14**) and the commercially available racemic carboxylic acid derivative **rac-12a**. 6-Fluoro-3-methylindazole **(15)** gave access to the enantiopure target compound **3b**.

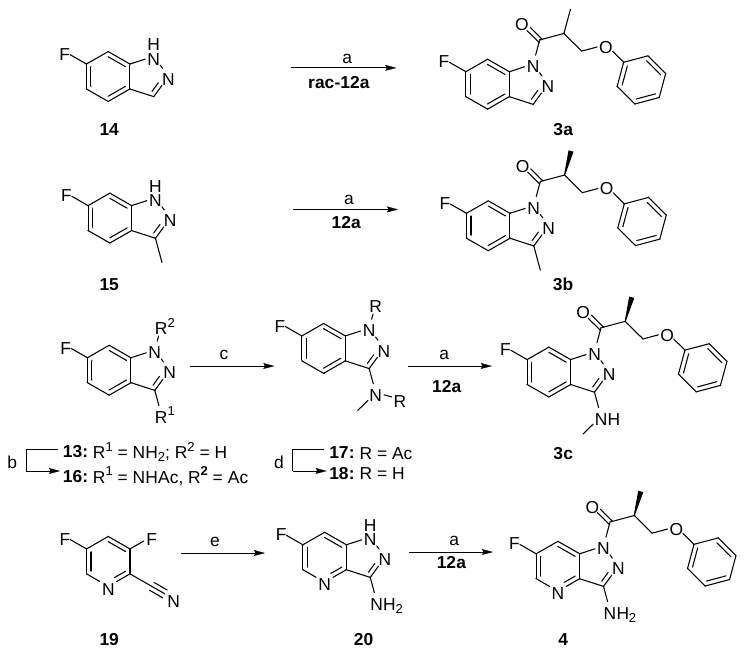

**Scheme S3.** Synthesis of the ‘853 derivatives with modifications in positions 3 or 4, reagents, and conditions: (a) EDC × HCl, HOAt, DMF, rt (**3a:** 39%, **3b:** 57%, **3c:** 29%, **4:** 31%); (b) AcCl, pyridine, DMAP, 0 °C to rt, 2 h (72%), (c) NaH, MeI, DMF, rt, 1 h (77%); (d) 1.25 M HCl in MeOH, 115 °C, 2 h (100% crude); (e) N_2_H_4_, EtOH, 70 °C, 17 h (15%).

For the synthesis of the *N*-methyl derivative **3c**, a novel, efficient pathway for the preparation of 3-(*N*-methylamino)indazole derivatives had to be established. In detail, the diacetyl derivative **16** was prepared from 6-fluoro-1*H*-indazole-3-amine (**13**) using acetylchloride in the presence of pyridine and 4-dimethylaminopyridine (DMAP). Deprotonation with sodium hydride (NaH) and subsequent reaction with methyl iodide (MeI) gave rise to the *N*-methyl derivative **17**, which was deprotected with hydrogen chloride (HCl) in methanol (MeOH). The resulting 6-fluoro-*N*-methyl-1*H*-indazol-3-amine (**18**) was acylated with the (*S*)-carboxylic acid derivative **12a** to give the target compound **3c**. The aza analog of **‘853** **4** was synthesized starting from 6-fluoro-1*H*-pyrazolo[4,3-*b*]pyridin-3-amine (**20**), which was available by a ring closing reaction of the pyridine derivative **19** with hydrazine hydrate according to a previously reported protocol. ^5^ It is noteworthy, that the cyclization reaction also led to the nucleophilic attack of hydrazine in position 5 of the pyridine nucleus. Thus, the respective aryl hydrazine side product was formed, which had to be separated by preparative HPLC. Coupling with the (*S*)-carboxylic acid derivative **12a** resulted in the formation of target compound **4**.

Furthermore, the fluorine in position 6 of the indazole scaffold was replaced by a variety of substituents (scheme S4).

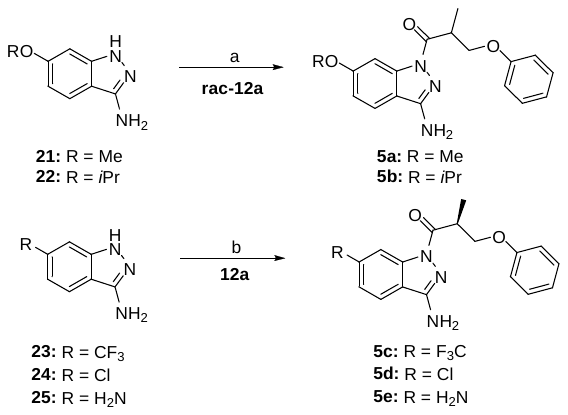

**Scheme S4.** Synthesis of the **‘853** derivatives with alkoxy substituents in positions 6, reagents and conditions: (a) PyBOP, HOAt, DMF, microwave irradiation (**5a:** 56%, **5b:** 54%); (b) EDC × HCl, HOAt, DMF, rt (**5c:** 23%, **5d:** 54%, **5e:** 44%).

The alkoxy substituted compounds **5a,b** were synthesized as racemates starting from **rac-12a**, employing the respective, commercially available 6-alkoxy-indazole derivatives **21** and **22**. Benzotriazol-1-yloxytripyrrolidinophosphonium hexafluorophosphate (PyBOP) in presence of HOAt and *N,N*-diisopropylethylamine (DIPEA) were used for the coupling reaction under microwave promoted conditions. Starting from the buyable 6-trifluoromethyl-, 6-chloro- or 6-aminoindazole analogs **23**, **24** and **25**, the EDC/HOAt promoted coupling with the (*S*)-carboxylic acid derivative **12a** furnished the respective target compounds **5c**, **5d** and **5e**, respectively. The syntheses of the **‘853**-congeners bearing an aminocarbonyl or a hydroxymethyl group in position 6 are outlined in scheme S5. The primary amide derivative **27** was obtained from the commercially available carboxylic acid derivative **26** by a one pot coupling protocol employing PyBOP in presence of NH_4_Cl and DIPEA.

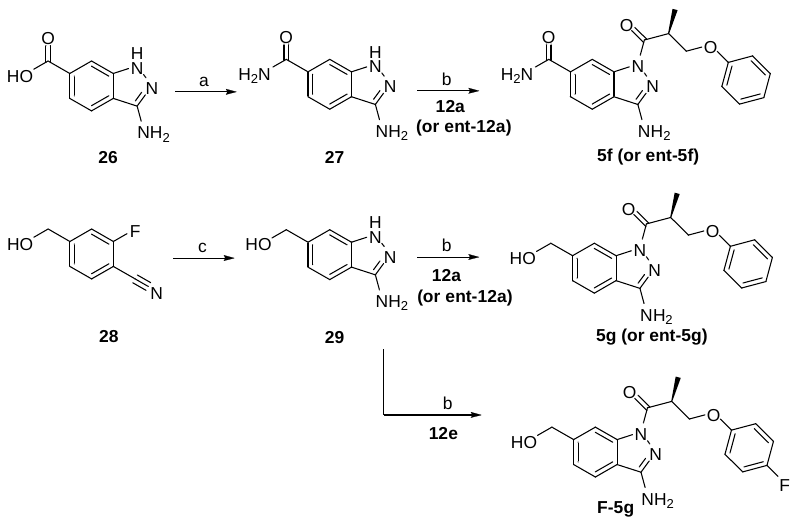

**Scheme S5.** Synthesis of novel indazole scaffolds and the corresponding acyl derivatives, reagents and conditions: (a) NH_4_Cl, PyBOP, DIPEA, DMF, rt, 12 h (83%); (b) EDC × HCl, HOAt, DMF, rt, 3 h (**5f:** 64%, **5g:** 53%, **F-5g:** 45%); (c) N_2_H_4_, BuOH, 120 °C, 4 h (69%).

The carbinol derivative **29** was synthesized by the cyclization reaction of the commercially available o-fluorobenzonitrile precursor **28** with hydrazine hydrate according to an established method.^6^ Both scaffolds were coupled with the (*S*)-carboxylic acid derivative **12a** to furnish the both target compounds **5f** and **5g**. Starting from the (*R*)-carboxylic acid derivative **ent-12a**, the corresponding enantiomers **ent-5f** and **ent-5g** were prepared as well. Furthermore, the (*S*)-4-fluorophenoxy analog **F-5g** was obtained in a similar fashion by the acylation with the (*S*)-carboxylic acid derivative **12e**.

The 6-carboxamide and 6-amino derivatives **7a** and **7b** renouncing the amino group in position 3 were obtained analogously starting from the respective purchasable precursors **30** and **31** (scheme S6). The enantiomer **ent-7a** was synthesized employing (*R*)-carboxylic acid derivative **ent-12a**.

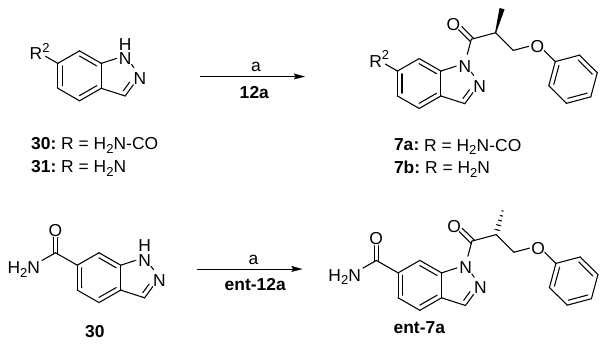

**Scheme S6.** Synthesis of the **‘853** derivatives with modifications in the positions 3 and 6, reagents and conditions: (a) EDC × HCl, HOAt, DMF, rt (**7a:** 64%, **7b:** 14%).

The *N*-formyl congeners **6a**, **6b** and **8** were prepared starting from the corresponding amine derivatives **5e** and **7b** taking advantage of activated formic acid (scheme S7).

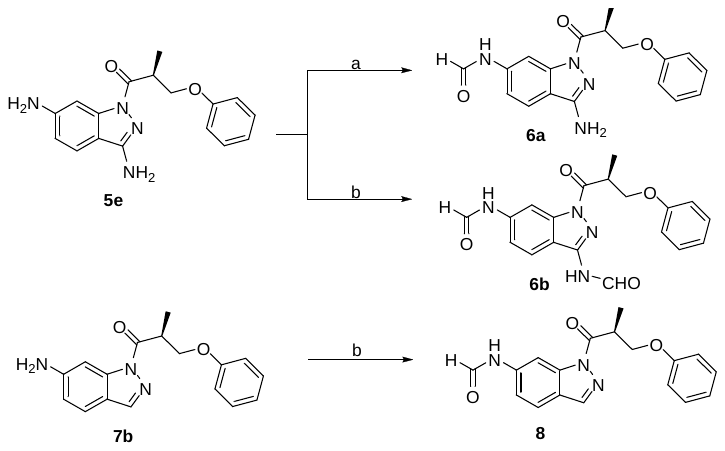

**Scheme S7.** Synthesis of the ‘853 derivatives further modified with a formyl group, reagents and conditions: (a) 1. HCO_2_H, Ac_2_O, THF, 60 °, 2 h, 2. **5e**, 0 °C, 30 min (42%); (b) 1. HCO_2_H, Ac_2_O, THF, 60 °C, 2 h, 2. **5e** or **7b**, 0 °C to rt, for **6b:** 60 min, for **8:** 90 min (**6b:** 37%, **8:** 28%).

The preparation of the fused derivative **9** (scheme S8) was achieved starting from the alkine derivative **32**, performing a gold(I) promoted hydroarylation reaction to give the chromene derivative **33**.^7^ Since the original cyclization protocol did not work with satisfying yield and purity, we preferred to apply more regioselective conditions for ring closure employing toluene as the solvent and working at 0 °C, ^8^ thus leading to a cleaner formation of only 14% regioisomer instead of 25% reported in the literature. The mixture thus obtained was subjected to hydrogenation to give the chroman derivative **34** (and the respective regioisomer). Reaction with *N*-bromosuccinimide furnished the 6-brominated derivative **35**, when chromatography allowed to reduce the amount of the regioisomer to 6%. Transnitrilation reaction employing *n*-butyllithium and dimethylmalononitrile allowed to cleanly prepare the cyclization precursor **36**, ^9^ which could be subjected to hydrazine hydrate to obtain the fused indazole-amine derivative **37**. Acylation with the racemic carboxylic acid derivative **rac-12a** led to the formation of the target compound **9**.

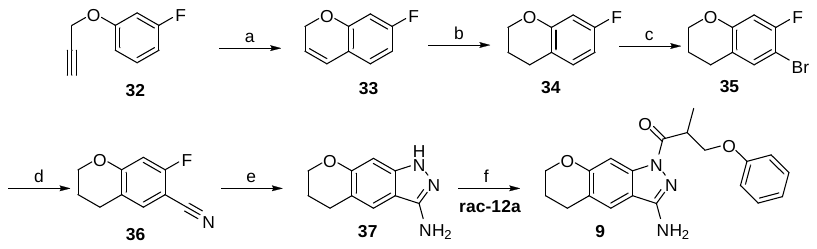

**Scheme S8.** Synthesis of the fused acylindazole derivative (a) (acetonitrile)[2-biphenyl)di-*tert*-butylphosphine]gold(I) hexafluoroantimonate, toluene, 0 °C (crude); (b) H_2_/Pd(OH)_2_/C, MeOH, rt, 2 h (crude); (c) *N*-bromosuccinimide, CH_3_CN, 0 °C (23% over three steps); (d) 1. BuLi, THF, -79 °C. 2. dimethylmalononitrile, THF, -79 °C (54%); (e) N_2_H_4_, BuOH, 120 °C, 22 h (76%); (f) EDC × HCl, HOAt, DMF, microwave irradiation (61%).

The preparation of the compounds incorporating a modified spacer was performed as shown in scheme S9. Starting from the commercially available carboxylic acids **39-42** or the respective sodium carboxylate **38**, coupling with EDC × HCl and HOAt allowed to obtain the target compounds **10a-e**. Preparative chiral HPLC gave rise to the separated enantiomers of **10b-d**.

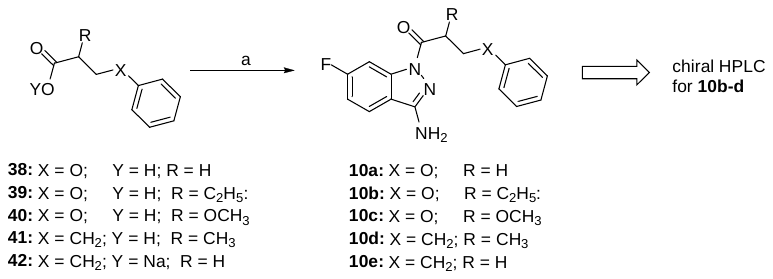

**Scheme S9.** Synthesis of the ‘853 derivatives with a modified spacer, reagents and conditions: (a) 1. EDC × HCl, HOAt, DMF, rt, 10 min, 2. 6‑fluoro-1*H*-indazole-3-amine (**13**), rt, **10a** and **10b:** 3 h, **10c:** 2 h, **10d:** 1 h, **10e:** 1.5 h (**10a:** 27%, **10b:** 45%, **10c:** 43%, **10d:** 30%, **10e:** 42%).

**General materials and methods for organic synthesis of the library compounds Z2075279358** - **Z995908944**

**Method 1:** A solution of amine (100 mg), the carboxylic acid (1.1 mol. eq. to the amine), and DMSO (0.5 mL) were placed into a capped glass vial and the mixture was stirred for 30 min at rt. 1-Ethyl-3-(3-dimethylaminopropyl)-carbodiimide (EDC, 1.2 mol. eq. to the amine) was added and stirring was continued for 1 h. In case of obtaining a clear solution, the mixture was left overnight at rt. In case of the solution remained opalescent, the vial was placed in an ultrasound bath overnight. The mixture was filtered and the solvent was evaporated to give the crude product, which was further purified by preparative HPLC.

**Method 2:** A solution of amine (100 mg), the carboxylic acid (1.1 mol. eq. to the amine), and DMF (0.5 mL) was stirred for 30 min, and subsequently carbonyldiimidazole (CDI, 1.1 mol. eq. to the amine) was added. The mixture was stirred in a sealed vial for 24 hours at rt. Chloroform (3 mL) was added to the reaction mixture and a washing step with water (2×1 mL) was performed. The solvent was evaporated under reduced pressure and trifluoroacetic acid (0.6 mL) was added to the residue. The vial was left in a shaker for 12 h at rt, then chloroform (3 mL) was added and evaporation under reduced pressure was performed to give the crude product, which was purified by HPLC.

**Method 3:** A solution of amine (100 mg) and DIPEA (1.1 mol. eq. to the amine; additional equivalents were added when the amine used was in a salt form) in DMSO (0.5 mL) was shaken for 20 min at rt and subsequently, the respective alkyl chloride (1 mol. eq. to the amine) was added. The vial was sealed and stirred for 1 h and then heated for 8 h at 100 °C. After cooling down, evaporation under reduced pressure was performed to give the crude product, which was purified by HPLC.

**Method 4:** According to a previously published procedure.^10^

**Method 5:** According to a previously published procedure.^11^

**Method 6:** According to a previously published procedure.^12^

^1^H- and ^13^C-NMR (DEPTQ) and IR spectra, ESI-MS, high-resolution mass spectra, and analytical HPLC were acquired as described for the target compounds of type **1-10** (see below).

**General materials and methods for organic synthesis of the target compounds of type 1-10**

Reagents and solvents were purchased in their purest grade from abcr, Acros, Alfa Aesar, Chemspace, Enamine, Manchester Organics, Sigma Aldrich, TCI, VWR and were used without further purification. Unless otherwise noted, reactions were performed under nitrogen atmosphere employing dry solvents of commercial quality, used as purchased. All reactions were carried out using a magnetic stirrer with optional aluminum heating block or ice bath for sealed microwave vials and oil or ice bath for round bottom flasks, respectively. Solvents were evaporated by a rotation evaporator equipped with a membrane vacuum pump. Microwave assisted (Discover^®^ microwave oven, CEM Corp.) synthesis was carried out by 20 × irradiating with microwaves (50 W, irradiation time: 10 s). In between each irradiation step, intermittent cooling of the reaction mixture to a temperature of -10°C was achieved by sufficient agitation in an ethanol-ice bath. TLC analyses were performed using Merck 60 F254 aluminum sheets and analyzed by UV light (254 nm). Purification by flash column chromatography was conducted using silica gel 60 (40-63 µm mesh, Merck) and eluents as binary mixtures with the volume ratios indicated. Preparative HPLC was performed on an Agilent 1200 preparative series HPLC system or on an Agilent HPLC 1260 Infinity system combined with an MWD detector and fraction collector, applying a linear gradient and a flow rate as indicated below. As HPLC column, a Zorbax-Eclipse XDB-C8 PrepHT (21.2 mm × 150 mm, 5 µm) was used. Products purified by preparative HPLC using aqueous solvents were lyophilized. For the separation of enantiomers, a Waters preparative HPLC system consisting of the 2545 binary gradient module, 2707 autosampler, 2998 photodiode array detector and the fraction collector III was employed, using a Chiralpak IC column (250 mm × 30 mm, 5 µm), a flow rate of 45 mL/min and an isocratic binary solvent system specified below. Compounds were characterized by NMR-, IR- and high-resolution mass spectra (HRMS). Purity was assessed by RP-HPLC. All assayed compounds were >95% pure. ESI-mass spectra were recorded using LC-MS: Thermo Scientific Dionex Ultimate 3000 UHPLC quarternary pump, autosampler, and RS-diode array detector, column: Zorbax-Eclipse XDB-C8 analytical column, 3.0 mm × 100 mm, 3.5 μm, flow rate 0.4 mL/min using DAD detection (230 nm; 254 nm), coupled to a Bruker Daltonics Amazon mass spectrometer using ESI as the ionization source. High mass accuracy and resolution experiments were performed on a Bruker Daltonics timsTOF Pro spectrometer using electrospray ionization (ESI) as an ionization source. NMR spectra were obtained either on a Bruker Avance III 400 (400 MHz for ^1^H and 101 MHz for ^13^C) or a Bruker Avance III 600 (600 MHz for ^1^H and 151 MHz for ^13^C) spectrometer, the latter equipped with a Prodigy nitrogen-cooled probe at 297 K, using the deuterated solvents indicated below. For the spectra recorded in organic solvents, the chemical shifts are reported in ppm (*δ*) relative to TMS. For measurements in D_2_O, the water peak was used for calibration. IR spectra were performed on a Jasco FT/IR 4100 spectrometer using a KBr pellet or with substance film on a NaCl crystal plate, as specified. Substance purities were assessed by analytical HPLC (Agilent 1100 analytical series, equipped with a quarternary pump and variable wavelength detector; column Zorbax Eclipse XDB-C8 analytical column, 4.6 mm × 150 mm, 5 μm, flow rate 0.5 mL / min, detection wavelengths: 220 nm, 254 nm; system 1: methanol / 0.1% aq. HCOOH, linear gradient: 10% methanol for 3 min, 10% to 100% methanol in 15 min, 100% methanol for 6 min; system 2: acetonitrile / 0.1% aq. HCOOH, linear gradient: 5% acetonitrile for 3 min, 5% to 95% acetonitrile in 15 min, 95% acetonitrile for 6 min; system 3: acetonitrile / 0.1% aq. TFA, linear gradient: 3% to 85% acetonitrile in 26 min, 85% to 95% acetonitrile in 2 min, 95% acetonitrile for 2 min; system 4: acetonitrile / 0.1% aq. TFA, linear gradient: 3% to 25% acetonitrile in 26 min) system 5: acetonitrile / 0.1% aq. HCOOH, linear gradient: 3% to 85% acetonitrile in 26 min, 85% to 95% acetonitrile in 2 min, 95% acetonitrile for 2 min;. Chiral analytical HPLC was run on an AGILENT series 1100 system equipped with a VWD and detection at 254 nm. As a chiral column, a DAICEL Chiralpak IC column (4.6 mm × 250 mm, 5 µM) was used at 20 °C and a flow rate 1.0 mL / min with the solvent system as indicated. Specific optical rotation values (°× mL × dm^-1^ × g^-1^) were obtained from a Jasco P2000 polarimeter with the solvents indicated. Melting points were determined in open capillaries using a Büchi 510 melting point apparatus and are given uncorrected.

**General experimental procedures**

General procedure A (GPA): preparation of 1.26 M Jones reagent

To a stirred solution of CrO_3_ (126 mg) in 500 µL water at 0 °C was slowly added conc. H_2_SO_4_ (126 µL). The volume was adjusted to 1 mL using water.

**Synthesis procedures of the library compounds Z2075279358 - Z995908944**

**(*R*,*S*)-*N*-(6-Fluoro-1*H*-indazol-3-yl)-3-(1*H*-indazol-1-yl)-2-methylpropanamide (Z2075279358)**

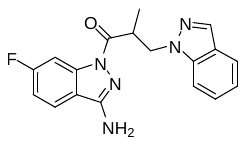

The synthesis was done according to method 1, starting from 3-(1*H*-indazol-1-yl)-2-methylpropanoic acid (EN300-206691) and 6-fluoro-1*H*-indazol-3-amine (EN300-100294), to obtain 62% yield.

ESI-MS:	*m/z* 338.3 [M+H]^+^.

HR-ESI-MS:	*m/z* [M+H]^+^ calcd. 338.1412 for C_18_H_17_FN_5_O, found 338.1411.

IR (NaCl):	3418, 1676, 1636, 1439, 1425 cm^-1^.

^1^H-NMR:	(600 MHz, DMSO-d_6_) *δ* 1.17 (d, *J* = 7.0 Hz, 3H), 4.31 (ddq, *J* = 7.0, 7.0, 7.0  Hz, 1H), 4.53 (dd, *J* = 14.2, 7.0 Hz, 1H), 4.86 (dd, *J* = 14.2, 7.0 Hz, 1H), 6.59 (s, 2H), 7.10 – 7.14 (m, 1H), 7.23 (ddd, *J* = 9.0, 8.8, 2.4 Hz, 1H), 7.36 – 7.40 (m, 1H), 7.71 – 7.74 (m, 2H), 7.89 (dd, *J* = 9.8, 2.4 Hz, 1H), 7.92 (dd, *J* = 8.8, 5.2 Hz, 1H), 8.02 – 8.03 (m, 1H).

^13^C-NMR:	(DEPTQ, 151 MHz, DMSO-d_6_) *δ* 15.0, 38.4, 50.0, 101.7 (d, *J* = 29 Hz), 109.8, 112.2 (d, *J* = 25 Hz), 117.0, 120.4, 120.7, 122.6 (d, *J* = 11 Hz), 123.4, 126.1, 132.9, 139.6, 139.8 (d, *J* = 18 Hz), 152.6, 163.2 (d, *J* = 244 Hz), 172.5.

HPLC:	system 3: λ = 220 nm,	*t*_R_ = 20.6 min, purity: 99.6%.

**3-(3-Isopropyl-1,2,4-oxadiazol-5-yl)-*N*-(4-(1-methyl-1*H*-pyrazol-4-yl)pyrimidin-2-yl) propanamide (Z2194302854)**

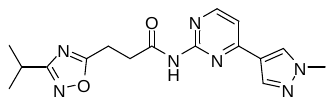

The synthesis was done according to method 1, starting from 3-(3-isopropyl-1,2,4-oxadiazol-5-yl)propanoic acid (EN300-27664) and 4-(1-methyl-1*H*-pyrazol-4-yl)pyrimidin-2-amine (EN300-187048), to obtain 47% yield.

ESI-MS:	*m/z* 342.2 [M+H]^+^.

HR-ESI-MS:	*m/z* [M+H]^+^ calcd. 342.1673 for C_16_H_20_N_7_O_2_, found 342.1672.

IR (KBr):	3434, 3150, 3104, 2968, 2934, 1681, 1596, 1581 cm^-1^.

^1^H-NMR:	(600 MHz, DMSO-d_6_) *δ* 1.24 (d, *J* = 6.9 Hz, 6H), 3.01 (sept, *J* = 6.9 Hz, 1H), 3.15 – 3.19 (m, 2H), 3.19 – 3.24 (m, 2H), 3.91 (s, 3H), 7.37 (d, *J* = 5.4 Hz, 1H), 8.11 (s, 1H), 8.41 (s, 1H), 8.53 (d, *J* = 5.4 Hz, 1H), 10.55 (s, 1H).

^13^C-NMR:	(DEPTQ, 151 MHz, DMSO-d_6_) *δ* 20.3, 21.3, 26.0, 32.8, 38.9, 111.1, 120.5, 131.3, 138.2, 157.7, 158.5, 159.4, 170.4, 174.3, 179.1.

HPLC:	system 3: λ = 220 nm,	*t*_R_ = 13.4 min, purity: 98.8%.

***N*-(3-Cyanophenyl)-1-(2,6-difluorophenyl)-*N*-methyl-1*H*-pyrazole-3-carboxamide (Z1343848401)**

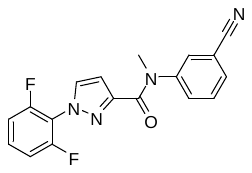

The synthesis was done according to method 1, starting from 1-(2,6-difluorophenyl)-1*H*-pyrazole-3-carboxylic acid (EN300-91144) and 3-(methylamino)benzonitrile (EN300-55410), to obtain 26% yield.

ESI-MS:	*m/z* 339.2 [M+H]^+^.

HR-ESI-MS:	*m/z* [M+H]^+^ calcd. 339.1052 for C_18_H_12_F_2_N_4_O, found 339.1052.

IR (KBr):	3434, 2923, 2227, 1648 cm^-1^.

^1^H-NMR:	(600 MHz, DMSO-d_6_) *δ* 3.43 (s, 3H), 6.67 (s, 1H), 7.32 (dd, *J* = 8.4, 8.4 Hz, 2H), 7.50 (dd, *J* = 7.8, 7.8 Hz, 2H), 7.55 – 7.62 (m, 2H), 7.68 (ddd, *J* = 7.8, 1.3, 1.3 Hz, 1H), 7.74 (s, 1H), 8.08 (s, 1H).

^13^C-NMR:	(DEPTQ, 151 MHz, DMSO-d_6_) *δ* 37.6, 108.8, 111.6, 112.6 (dd, *J* = 19, 4 Hz), 117.3 (t, *J* = 15 Hz), 118.2, 130.11, 130.14, 130.5, 131.4 (t, *J* = 10 Hz), 131.9, 134.2, 145.0, 148.2, 156.8 (dd, *J* = 254, 5 Hz), 162.6.

HPLC:	system 3: λ = 220 nm,	*t*_R_ = 18.8 min, purity: 99.7%.

**(*R,S*)-*N*-((*S,R*)-6-amino-1,2,3,4-tetrahydronaphthalen-1-yl)-7-methyl-5,6,7,8-tetrahydro-imidazo[1,2-a]pyridine-2-carboxamide** and **(*R,S*)-*N*-((*R,S*)-6-amino-1,2,3,4-tetrahydro-naphthalen-1-yl)-7-methyl-5,6,7,8-tetrahydroimidazo[1,2-a]pyridine-2-carboxamide (Z2776419998)**

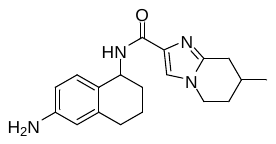

The synthesis was done according to method 2, starting from (*R,S*)-7-methyl-5,6,7,8-tetrahydroimidazo[1,2-a]pyridine-2-carboxylic acid (EN300-309016) and (*R,S*)-*tert*-butyl (5-amino-5,6,7,8-tetrahydronaphthalen-2-yl)carbamate (EN300-142125). The substance was obtained as a mixture of two diastereomers (like and unlike, each racemates), which were not separated to obtain 26% yield.

ESI-MS:	*m/z* 325.3 [M+H]^+^.

HR-ESI-MS:	*m/z* [M+H]^+^ calcd. 325.2023 for C_19_H_25_N_4_O, found 325. 2023

IR (KBr):	3427, 3348, 3170, 1690, 1664, 1567 cm^-1^.

^1^H-NMR:	(600 MHz, DMSO-d_6_, diastereomers) *δ* 1.04 (d, *J* = 6.6 Hz, 3H), 1.53 – 1.62 (m, 1H), 1.63 – 1.70 (m, 1H), 1.73 – 1.81 (m, 2H), 1.81 – 1.87 (m, 1H), 1.91 – 2.02 (m, 2H), 2.27 (dd, *J* = 16.4, 10.5 Hz, 1H), 2.51 – 2.57 (m, 1H), 2.59 – 2.66 (m, 1H), 2.80 and 2.81 (2× ddd, each *J* = 16.4, 4.3, 2.1 Hz, 1H), 3.88 (ddd, *J* = 12.5, 12.0, 4.7 Hz, 1H), 4.08 (ddd, *J* = 12.5, 5.6, 3.3 Hz, 1H), 4.90 (brs, 2H), 4.91 – 4.96 (m, 1H), 6.27 (d, *J* = 2.2 Hz, 1H), 6.350 and 6.354 (2× dd, each *J* = 8.3, 2.2 Hz, 1H), 6.785 and 6.792 (2× d, each *J* = 8.3 Hz, 1H), 7.33 and 7.34 (2× d, each *J* = 9.0 Hz, 1H), 7.52 (s, 1H).

^13^C-NMR:	(DEPTQ, 151 MHz, DMSO-d_6_, diastereomers) *δ* 20.05 and 20.09, 20.8, 27.1, 29.0, 30.0, 30.2, 31.91 and 31.92. 43.8, 45.65 and 45.66, 112.5, 113.2, 120.9, 124.61 and 124.63, 128.68 and 128.72, 135.1. 137.4, 144.3, 147.3, 161.3.

HPLC:	system 3: λ = 220 nm,	*t*_R1_ = 10.0 min, *t*_R2_ = 10.1 min, purity: 99.1% (both diastereomers).

***N*-(1-(3-Amino-3-oxopropyl)-1*H*-pyrazol-3-yl)-3-methoxy-2-naphthamide (Z1224795288)**

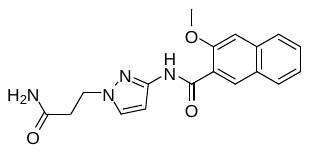

The synthesis was done according to method 1, starting from 3-methoxy-2-naphthoic acid (EN300-00001) and 3-(3-amino-1*H*-pyrazol-1-yl)propanamide (EN300-80390), to obtain 75% yield.

ESI-MS:	*m/z* 339.2 [M+H]^+^.

HR-ESI-MS:	*m/z* [M+H]^+^ calcd. 339.1452 for C_18_H_19_N_4_O_3_, found 339. 1452.

IR (KBr):	3426, 3348, 3170, 1690, 1664, 1568, 1500 cm^-1^.

^1^H-NMR:	(600 MHz, DMSO-d_6_) *δ* 2.60 (t, *J* = 7.0 Hz, 2H), 4.01 (s, 3H), 4.23 (t, *J* = 7.0 Hz, 2H), 6.61 (d, *J* = 2.1 Hz, 1H), 6.90 (s, 1H), 7.40 (s, 1H), 7.42 (ddd, *J* = 8.2, 6.9, 1.1 Hz, 1H), 7.51 (s, 1H), 7.56 (ddd, *J* = 8.0, 6.9, 1.2 Hz, 1H), 7.88 (brd, *J* = 8.0 Hz, 1H), 7.96 (brd, *J* = 8.2 Hz, 1H), 8.29 (s, 1H), 10.49 (s, 1H).

^13^C-NMR:	(DEPTQ, 151 MHz, DMSO-d_6_) *δ* 35.5, 47.4, 56.0, 96.9, 106.7, 124.3, 125.3, 126.4, 127.5, 127.9, 128.4, 130.5, 130.6, 135.1, 146.6, 154.2, 162.9, 171.5.

HPLC:	system 3: λ = 220 nm,	*t*_R_ = 16.9 min, purity: 97.7%.

**(5-(4-Chlorophenyl)-1,3,4-oxadiazol-2-yl)methyl 2-hydroxybenzoate (Z19702639)**

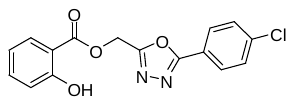

The synthesis was done according to method 3, starting from salicylic acid (EN300-16723) and 2-(chloromethyl)-5-(4-chlorophenyl)-1,3,4-oxadiazole (EN300-08138), to obtain 87% yield.

ESI-MS:	*m/z* 331.2 [M+H]^+^.

HR-ESI-MS:	*m/z* [M+H]^+^ calcd. 331.0480 for C_16_H_12_ClN_2_O_4_, found 331.0480.

IR (KBr):	3448, 3240, 3089, 1690, 1484, 1252 cm^-1^.

^1^H-NMR:	(600 MHz, DMSO-d_6_) *δ* 5.69 (s, 2H), 6.96 (ddd, *J* = 8.1, 7.1, 0.9 Hz, 1H), 7.01 (dd, *J* = 8.5, 0.9 Hz, 1H), 7.54 (ddd, *J* = 8.5, 7.1, 1.7 Hz, 1H), 7.67 – 7.71 (m, 2H), 7.84 (dd, *J* = 8.1, 1.7 Hz, 1H), 8.00 – 8.05 (m, 2H), 10.22 (s, 1H).

^13^C-NMR:	(DEPTQ, 151 MHz, DMSO-d_6_) *δ* 56.0, 113.0, 117.6, 119.4, 121.9, 128.5, 129.7, 130.5, 135.9, 137.1, 159.7, 162.2, 164.0, 167.0.

HPLC:	system 3: λ = 220 nm, *t*_R_ = 22.7 min, purity: 91.2%.

**5-(((6-Chloroimidazo[1,2-*a*]pyridin-2-yl)methyl)thio)-4-isopropyl-4*H*-1,2,4-triazol-3-amine (Z1096199008)**

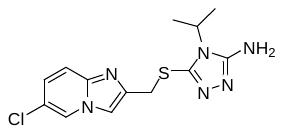

The synthesis was done according to method 5, starting from 5-amino-4-isopropyl-4*H*-1,2,4-triazole-3-thiol (EN300-2493721) and 6-chloro-2-(chloromethyl)imidazo[1,2-*a*]pyridine (EN300-33589), to obtain 35% yield.

ESI-MS:	*m/z* 323.2 [M+H]^+^.

HR-ESI-MS:	*m/z* [M+H]^+^ calcd. 323.0840 for C_13_H_16_ClN_6_S, found 323.0839.

IR (KBr):	3434, 3103, 1654, 1566 cm^-1^.

^1^H-NMR:	(600 MHz, DMSO-d_6_) *δ* 1.25 (d, *J* = 7.0 Hz, 3H), 4.30 (s, 2H), 4.34 (sept, *J* = 7.0 Hz, 1H), 5.73 (s, 2H), 7.26 (dd, *J* = 9.6, 2.1 Hz, 1H), 7.54 (d, *J* = 9.6 Hz, 1H), 7.78 (s, 1H), 8.78 (dd, *J* = 2.1, 0.9 Hz, 1H).

^13^C-NMR:	(DEPTQ, 151 MHz, DMSO-d_6_) *δ* 20.1, 32.3, 46.2, 111.8, 117.2, 118.8, 124.7, 125.6, 141.9, 142.6, 143.1, 155.4.

HPLC:	system 3: λ = 220 nm,	*t*_R_ = 10.0 min, purity: 97.0%.

***N*-((4*H*-1,2,4-Triazol-3-yl)methyl)-5-(benzo[*d*]thiazol-2-yl)thiophene-2-carboxamide (Z873519648)**

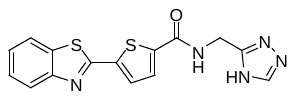

The synthesis was done according to method 1, starting from 5-(benzo[*d*]thiazol-2-yl)thiophene-2-carboxylic acid (EN300-00707) and (4*H*-1,2,4-triazol-3-yl)methanamine (EN300-1264530), to obtain 33% yield.

ESI-MS:	*m/z* 342.2 [M+H]^+^.

HR-ESI-MS:	*m/z* [M+H]^+^ calcd. 342.0478 for C_15_H_12_N_5_OS_2_, found 342.0477.

IR (KBr):	3431, 3379, 3276, 1653, 1624, 1546, 1529 cm^-1^.

^1^H-NMR:	(600 MHz, DMSO-d_6_, two isomers were observed) *δ* 4.57 (brs, 2H), 7.48 (ddd, *J* = 8.0, 7.3, 1.2 Hz, 1H), 7.56 (ddd, *J* = 8.0, 7.3, 1.2 Hz, 1H), 7.88 (d, *J* = 4.2 Hz, 1H), 7.90 (d, *J* = 4.2 Hz, 1H), 8.05 (ddd, *J* = 8.0, 1.2, 0.7 Hz, 1H), 8.15 (ddd, *J* = 8.0, 1.2, 0.7 Hz, 1H), 8.49 (s, 1H), 9.29 (s, 1H), 13.90 (brs, 1H).

^13^C-NMR:	(DEPTQ, 151 MHz, DMSO-d_6_, two isomers were observed) *δ* 35.3 and 36.9, 122.5, 122.8, 125.9, 126.9, 129.1, 129.5, 134.5, 139.7, 142.3 and 143.0, 144.0, 148.8 (?), 151.3, 153.0, 160.3, 160.4 and 160.8.

HPLC:	system 3: λ = 220 nm,	*t*_R_ = 16.3 min, purity: >99%.

**2-(3-(Imidazo[1,2-*a*]pyridin-2-ylmethyl)-1,2,4-oxadiazol-5-yl)-2,3-dihydro-1*H*-inden-2-ol (Z1783799713)**

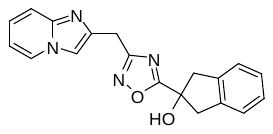

The synthesis was done according to method 4, starting from 2-(imidazo[1,2-*a*]pyridin-2-yl)acetonitrile (EN300-25598) and 2-hydroxy-2,3-dihydro-1*H*-indene-2-carboxylic acid (EN300-82840), to obtain 18% yield.

ESI-MS:	*m/z* 333.2 [M+H]^+^.

HR-ESI-MS:	*m/z* [M+H]^+^ calcd. 333.1348 for C_19_H_17_N_4_O_2_, found 333.1347.

IR (KBr):	3434, 2794, 1637, 1576, 1505 cm^-1^.

^1^H-NMR:	(600 MHz, DMSO-d_6_) *δ* 3.27 (d, *J* = 16.3 Hz, 2H), 3.53 (d, *J* = 16.3 Hz, 2H), 4.20 (s, 2H), 6.30 (s, 1H), 6.86 (ddd, *J* = 6.8, 6.8, 1.2 Hz, 1H), 7.20 (ddd, *J* = 9.1, 6.8, 1.2 Hz, 1H), 7.16 – 7.20 (m, 2H), 7.23 – 7.27 (m, 2H), 7.47 (dddd, *J* = 9.1, 1.2, 1.2 , 1.2 Hz, 1H), 7.84 (dd, *J* = 1.2, 0.7 Hz, 1H), 8.49 (ddd, *J* = 6.8, 1.2, 1.2 Hz, 1H).

^13^C-NMR:	(DEPTQ, 151 MHz, DMSO-d_6_) *δ* 25.8, 46.1, 77.0, 110.9, 111.9, 116.3, 124.56, 124.58, 126.7, 139.8, 140.6, 144.1, 168.3, 182.5.

HPLC:	system 3: λ = 220 nm,	*t*_R_ = 14.0 min, purity: 97.8%.

**1-(4-Fluorophenyl)-*N*-methyl-*N*-((2-methylthiazol-4-yl)methyl)-1*H*-pyrazole-3-carboxamide (Z899051432)**

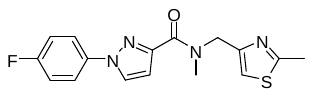

The synthesis was done according to method 1, starting from 1-(4-fluorophenyl)-1*H*-pyrazole-3-carboxylic acid (EN300-36826) and *N*-methyl-1-(2-methylthiazol-4-yl)methanamine (EN300-27495), to obtain 76% yield.

ESI-MS:	*m/z* 331.2 [M+H]^+^.

HR-ESI-MS:	*m/z* [M+H]^+^ calcd. 331.1023 for C_16_H_16_FN_4_OS, found 331.1023.

IR (KBr):	3443, 3114, 2926, 1625, 1523, 1509 cm^-1^.

^1^H-NMR:	(600 MHz, DMSO-d_6_, two isomers were observed) *δ* 2.63 and 2.65 (2× s, 3H), 3.01 and 3.07 (2× s, 3H), 4.72 and 5.06 (2× s, 2H), 6.86 and 6.87 (2× d, each *J* = 2.5 Hz, 1H), 7.25 and 7.29 (2× s, 1H), 7.33 – 7.41 (m, 2H), 7.81 – 7.86 and 7.89 – 7.94 (2× m, 2H), 8.53 and 8.55 (2× d, each *J* = 2.5 Hz, 1H).

^13^C-NMR:	(DEPTQ, 151 MHz, DMSO-d_6_, two isomers were observed) *δ* 18.8, 33.9 and 36.9, 47.2 and 50.2, 110.1 and 110.2, 115.3 and 115.7, 116.4 (d, *J* = 23 Hz), 120.8 and 121.0 (2× d, each *J* = 9 Hz), 128.77 and 128.82, 135.8 and 135. 9 (2× d, each *J* = 3 Hz), 148.1 and 148.2, 151.6 and 152.5, 160.57 (d, *J* = 246 Hz) and 160.62 (d, *J* = 242 Hz), 162.5 and 162.9, 165.6 and 165.7.

HPLC:	system 3: λ = 220 nm,	*t*_R_ = 17.8 min, purity: >99%.

**(*R*,*S*)-2-((5-Amino-4-isopropyl-4*H*-1,2,4-triazol-3-yl)thio)-*N*-(3-fluoro-4-methylphenyl) propanamide (Z1262422554)**

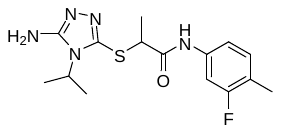

The synthesis was done according to method 5, starting from 5-amino-4-isopropyl-4*H*-1,2,4-triazole-3-thiol (EN300-2493721) and 2-chloro-*N*-(3-fluoro-4-methylphenyl)propanamide (EN300-23356), to obtain 37% yield.

ESI-MS:	*m/z* 338.2 [M+H]^+^.

HR-ESI-MS:	*m/z* [M+H]^+^ calcd. 338.1445 for C_15_H_21_FN_5_OS, found 338.1446.

IR (KBr):	3316, 3190, 2980, 2933, 1684, 1627, 1551, 1511 cm^-1^.

^1^H-NMR:	(600 MHz, DMSO-d_6_) *δ* 1.34 and 1.35 (2× d, each *J* = 7.1 Hz, 6H), 1.46 (d, *J* = 7.0 Hz, 3H), 2.17 (d, *J* = 1.5 Hz, 3H), 4.14 (q, *J* = 7.0 Hz, 3H), 4.43 (qq, *J* = 7.1 Hz, 3H), 5.80 (s, 2H), 7.16 (dd, *J* = 8.3, 1.9 Hz, 1H), 7.20 (dd, *J* = 8.3, 8.3 Hz, 1H), 7.49 (dd, *J* = 12.8, 1.9 Hz, 1H), 10.36 (s, 1H).

^13^C-NMR:	(DEPTQ, 151 MHz, DMSO-d_6_) *δ* 13.6 (d, *J* = 2 Hz), 18.1, 20.18, 20.21, 46.4, 47.2, 105.9 (d, *J* = 27 Hz), 114.8 (d, *J* = 2 Hz), 118.8 (d, *J* = 17 Hz), 131.4 (d, *J* = 7 Hz), 138.1 (d, *J* = 13 Hz), 140.6, 155.6, 160.2, (d, *J* = 241 Hz), 169.5.

HPLC:	system 3: λ = 220 nm,	*t*_R_ = 16.3 min, purity: >99%.

**1-Cyclopropyl-3-(6-(dimethylamino)pyrimidin-4-yl)-1-(3,4-dimethylbenzyl)urea (Z2171315755)**

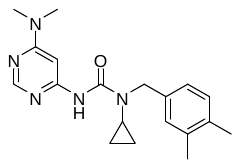

The synthesis was done according to method 6, starting from *N^4^,N^4^*-dimethylpyrimidine-4,6-diamine (EN300-187848) and *N*-(3,4-dimethylbenzyl)cyclopropanamine (N300-83837), to obtain 24% yield.

ESI-MS:	*m/z* 340.3 [M+H]^+^.

HR-ESI-MS:	*m/z* [M+H]^+^ calcd. 340.2132 for C_19_H_26_N_5_O, found 340.2130.

IR (KBr):	3422, 2936, 1684, 1608, 1521, 1498 cm^-1^.

^1^H-NMR:	(600 MHz, DMSO-d_6_) *δ* 0.73 – 0.80 (m, 2H), 0.84 – 0.89 (m, 2H), 2.18 (s, 3H), 2.20 (s, 3H), 2.65 – 2.70 (m, 1H), 3.03 (s, 6H), 4.45 (s, 2H), 6.96 (dd, *J* = 7.8, 1.5 Hz, 1H), 7.02 (brs, 1H), 7.08 (d, *J* = 7.8 Hz, 1H), 7.16 (d, *J* = 1.1 Hz, 1H), 8.19 (d, *J* = 1.1 Hz, 1H), 8.34 (brs, 1H).

^13^C-NMR:	(DEPTQ, 151 MHz, DMSO-d_6_) *δ* 8.7, 19.0, 19.4, 28.2, 36.8, 49.2, 86.9, 124.7, 128.4, 129.5, 134.7, 135.8, 136.1, 155.0, 156.9, 157.5, 162.8.

HPLC:	system 3: λ = 220 nm,	*t*_R_ = 18.5 min, purity: 95.0%.

***N*-(2-Chloro-6-fluorobenzyl)-1-(*m*-tolyl)-1*H*-1,2,3-triazole-4-carboxamide (Z995908944)**

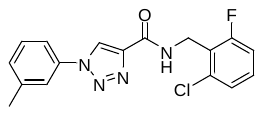

The synthesis was done according to method 1, starting from 1-(*m*-tolyl)-1*H*-1,2,3-triazole-4-carboxylic acid (EN300-61349) and (2-chloro-6-fluorophenyl)methanamine (EN300-33074), to obtain 50% yield.

ESI-MS:	*m/z* 345.1 [M+H]^+^.

HR-ESI-MS:	*m/z* [M+H]^+^ calcd. 345.0913 for C_17_H_15_ClFN_4_O, found 345.0912.

IR (NaCl):	3415, 3117, 2921, 1685 cm^-1^.

^1^H-NMR:	(600 MHz, DMSO-d_6_) *δ* 2.42 (s, 3H), 4.64 (dd, *J* = 5.3, 1.0 Hz, 3H), 6.61 (s, 2H), 7.24 (ddd, *J* = 9.6, 8.3, 1.3 Hz, 1H), 7.32 – 7.35 (m, 1H), 7.34 (dd, *J* = 8.1, 1.3 Hz, 1H), 7.40 (ddd, *J* = 8.3, 8.1, 6.1 Hz, 1H), 7.48 (dd, *J* = 7.8, 7.8 Hz, 1H), 7.71 – 7.74 (m, 1H), 7.78 – 7.80 (m, 1H), 8.87 (t, *J* = 5.3 Hz, 1H), 9.22 (s, 2H).

^13^C-NMR:	(DEPTQ, 151 MHz, DMSO-d_6_) *δ* 20.9, 34.4 (d, *J* = 4 Hz), 114.5 (d, *J* = 23 Hz), 117.5, 120.9, 123.7 (d, *J* = 15 Hz), 124.7, 125.4 (d, *J* = 3 Hz), 129.7, 130.1 (d, *J* = 10 Hz), 134.8 (d, *J* = 8 Hz), 136.2, 139.7, 143.2, 159.1, 61.5 (d, *J* = 249 Hz).

HPLC:	system 3: λ = 220 nm,	*t*_R_ = 21.7 min, purity: >99%.

**Detailed synthesis procedures of the compounds of type 1-10**

**(*R*)-2-Methyl-3-phenoxypropan-1-ol (11a)** ^1^

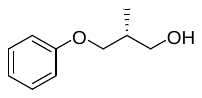

To a stirred solution of phenol distilled from molecular sieve (187 µL, 2.13 mmol) in DMF (4 mL) in a flame-dried flask, sodium hydride (60% in mineral oil; 97.4 mg, 2.44 mmol) was added. After stirring the suspension for 30 min at rt, (*R*)-3-bromo-2-methylpropan-1-ol (213 µL, 2.03 mmol) was added. The mixture was stirred for additional 18 h, then 2N NaOH was added and extraction with *tert*-butyl methyl ether (3×) was performed. The combined organic layers were washed with water and brine, dried over Na_2_SO_4_ and concentrated. After purification by flash-chromatography (isohexane / *tert*-butyl methyl ether, 7:3) compound **11a** (202 mg, 1.22 mmol, 60%) was obtained as colorless liquid.

ESI-MS:	*m/z* 189.3 [M+Na]^+^.

^1^H-NMR:	(400 MHz, CDCl_3_) *δ* 1.04 (d, *J* = 7.0 Hz, 3H), 1.90 (dd, *J* = 5.7, 5.7 Hz, 1H), 2.21 (m, 1H), 3.73 – 3.70 (m, 2H), 3.93 (dd, *J* = 9.1, 7.0 Hz, 1H), 3.98 (dd, *J* = 9.1, 5.3 Hz, 1H), 6.89 – 6.93 (m, 2H), 6.93 – 6.98 (m, 1H), 7.32 – 7.26 (m, 2H).

^13^C-NMR:	(DEPTQ, 101 MHz, CDCl_3_) *δ* 13.6, 35.7, 66.2, 71.2, 114.5, 120.9, 129.5, 158.8.

chiral HPLC:	isocratic elution with *n*-hexane / isopropanol, 95:5: *t*_R_ = 6.6 min, 99% *ee* (254 nm).

[*α*]*_D_^24^*:	+2.4 (*c* = 0.9, chloroform).

The analytical data of this compound are in accordance with those reported in the literature.

**(*S*)-2-Methyl-3-phenoxypropan-1-ol (ent-11a)** ^1^

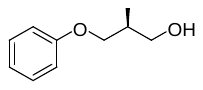

(*S*)-2-Methyl-3-phenoxypropan-1-ol **(ent-11a)** was synthesized analogously as described for **(*R*)-SX224 11a**, starting from (*S*)-3-bromo-2-methylpropan-1-ol.

chiral HPLC:	isocratic elution with *n*-hexane / isopropanol, 95:5: *t*_R_ = 6.9 min, 99% *ee* (254 nm).

[*α*]*_D_^25^*:	-2.9 (*c* = 1.6, chloroform).

The analytical data of this compound are in accordance with those reported in the literature.

**(*R*)-2-Methyl-3-(naphthalen-2-yloxy)propan-1-ol (11b)**

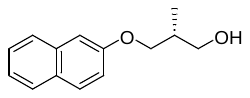

To a stirred solution of 2-naphthol (200 mg, 1.39 mmol) in DMF (4 mL) in a flame-dried flask sodium hydride (60% in mineral oil; 64.0 mg, 1.60 mmol) was added. After stirring the suspension for 30 min at rt, (*R*)-3-bromo-2-methylpropan-1-ol (138 µL, 1.32 mmol) was added. The mixture was stirred for additional 22 h, then 2N NaOH was added and extraction with *tert*-butyl methyl ether (3×) was performed. The combined organic layers were washed with water and brine, dried over Na_2_SO_4_ and concentrated. After purification by flash-chromatography (isohexane / *tert*-butyl methyl ether, 7:3) compound **11b** (206 mg, 0.95 mmol, 72%) was obtained as pale yellow liquid, which solidified upon standing (white wax).

ESI-MS:	*m/z* 238.9 [M+Na]^+^.

HR-ESI-MS:	*m/z* [M+H]^+^ calcd. 239.1043 for C_14_H_16_NaO_2_, found 239.1044.

^1^H-NMR:	(400 MHz, CDCl_3_) *δ* 1.09 (d, *J* = 7.0 Hz, 3H), 2.22 – 2.34 (m, 1H), 3.74 – 3.78 (m, 2H), 4.06 (dd, *J* = 9.1, 6.7 Hz, 1H), 4.09 (dd, *J* = 9.1, 5.5 Hz, 1H), 7.18 – 7.10 (m, 2H), 7.34 (ddd, *J* = 8.1, 6.9, 1.3 Hz, 1H), 7.44 (ddd, *J* = 8.2, 6.9, 1.3 Hz, 1H), 7.68 – 7.80 (m, 3H).

^13^C-NMR:	(DEPTQ, 101 MHz, CDCl_3_) *δ* 13.7, 35.7, 66.2, 71.2, 106.7, 118.8, 123.7, 126.4, 126.7, 127.6, 129.0, 129.4, 134.5, 156.7.

chiral HPLC:	isocratic elution with *n*-hexane / isopropanol, 98:2: *t*_R_ = 17.5 min, >99% *ee* (254 nm).

[*α*]*_D_^24^*:	+1.8 (*c* = 0.7, chloroform).

**(*S*)-2-Methyl-3-(naphthalen-2-yloxy)propan-1-ol (ent-11b)**

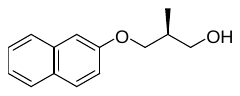

Methyl-3-(naphthalen-2-yloxy)propan-1-ol **(ent-11b)** was synthesized analogously as described for **11b**, starting from (*S*)-3-bromo-2-methylpropan-1-ol. The analytical data were in accordance.

chiral HPLC:	isocratic elution with *n*-hexane / isopropanol, 98:2: *t*_R_ = 18.2 min, >99% *ee* (254 nm).

[*α*]*_D_^23^*:	-3.0 (*c* = 0.8, chloroform).

**(*R*)-2-Methyl-3-(3-(trifluoromethyl)phenoxy)propan-1-ol (11c)**

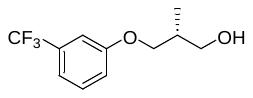

To a stirred solution of 3-(trifluormethyl)phenol (150 µL, 1.23 mmol) in DMF (4 mL) in a flame-dried flask sodium hydride (60% in mineral oil; 56.2 mg, 1.40 mmol) was added. After stirring the suspension for 30 min at rt, (*R*)-3-bromo-2-methylpropan-1-ol (123 µL, 1.17 mmol) was added. The mixture was stirred for additional 22 h, then 2N NaOH was added and extraction with *tert*-butyl methyl ether (3×) was performed. The combined organic layers were washed with water and brine, dried over Na_2_SO_4_ and concentrated. Purification by flash-chromatography (isohexane / *tert*-butyl methyl ether, 7:3) afforded **11c** as pale yellow liquid (148 mg, 0.63 mmol) still containing minor amounts of (*R*)-3-bromo-2-methylpropan-1-ol (confirmed by NMR). The crude product was used for the next step without further purification.

ESI-MS:	*m/z* 257.5 [M+Na]^+^.

HR-ESI-MS:	*m/z* [M+H]^+^ calcd. 257.0760 for C_11_H_13_F_3_NaO_2_, found 257.0761.

^1^H-NMR:	(400 MHz, CDCl_3_) *δ* 1.07 (d, *J* = 7.0 Hz, 3H), 1.66 – 1.73 (m, 1H), 2.16 – 2.27 (m, 1H), 3.70 – 3.75 (m, 2H), 3.97 – 4.00 (m, 2H), 7.05 – 7.10 (m, 1H), 7.12 – 7.16 (m, 1H), 7.18 – 7.23 (m, 1H), 7.35 – 7.42 (m, 1H).

^13^C-NMR:	(DEPTQ, 101 MHz, CDCl_3_) *δ* 13.8, 35.8, 65.7, 71.1, 111.4 (q, *J* = 4 Hz), 117.6 (q, *J* = 4 Hz), 118.1, 124.1 (q, *J* = 272 Hz), 130.1, 132.0 (q, *J* = 32 Hz), 159.1.

chiral HPLC:	isocratic elution with *n*-hexane/isopropanol, 98:2: *t*_R_ = 7.0 min, 99% *ee* (254 nm).

**(*S*)-2-Methyl-3-(3-(trifluoromethyl)phenoxy)propan-1-ol (ent-11c)**

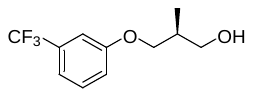

(*S*)-2-Methyl-3-(3-(trifluoromethyl)phenoxy)propan-1-ol **(ent-11c)** was synthesized analogously as described for **11c**, starting from (*S*)-3-bromo-2-methylpropan-1-ol. The analytical data were in accordance.

chiral HPLC:	isocratic elution with *n*-hexane / isopropanol, 98:2: *t*_R_ = 7.7 min, >99% *ee* (254 nm).

**(*R*)-3-(2,4-Di-*tert*-butylphenoxy)-2-methylpropan-1-ol (11d)**

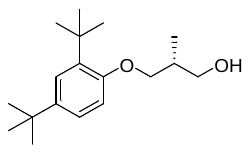

To a stirred solution of 2,4-di-*tert*-butylphenol (215 mg, 1.04 mmol) in DMF (4 mL) in a flame-dried flask sodium hydride (60% in mineral oil; 47.5 mg, 1.19 mmol) was added. After stirring the suspension for 30 min at rt, (*R*)-3-bromo-2-methylpropan-1-ol (104 µL, 0.99 mmol) was added. The mixture was stirred for additional 22 h, then 2N NaOH was added and extraction with *tert*-butyl methyl ether (3×) was performed. The combined organic layers were washed with water and brine, dried over Na_2_SO_4_ and concentrated. After purification by flash-chromatography (isohexane / *tert*-butyl methyl ether, 7:3) compound **11d** (174 mg, 0.62 mmol, 63%) was obtained as colorless liquid, which solidified upon standing (white wax).

ESI-MS:	*m/z* 301.2 [M+Na]^+^.

HR-ESI-MS:	*m/z* [M+H]^+^ calcd. 301.2138 for C_18_H_30_NaO_2_, found 301.2138.

^1^H-NMR	(400 MHz, CDCl_3_) *δ* 1.11 (d, *J* = 6.9 Hz, 3H), 1.30 (s, 9H), 1.40 (s, 9H), 1.57 (dd, *J* = 5.6, 5.6 Hz, 1H), 2.13 – 2.32 (m, 1H), 3.69 – 3.81 (m, 2H), 3.94 (dd, *J* = 9.1, 5.3 Hz, 1H), 3.98 (dd, *J* = 9.1, 6.1 Hz, 1H), 6.83 (d, *J* = 8.5 Hz, 1H), 7.17 (dd, *J* = 8.5, 2.5 Hz, 1H), 7.34 (d, *J* = 2.5 Hz, 1H).

^13^C-NMR	(DEPTQ, 101 MHz, CDCl_3_) *δ* 14.4, 30.1, 31.7, 34.4, 35.2, 36.2, 49.6, 65.8, 70.3, 111.4, 123.5, 124.1, 137.2, 142.7, 155.5.

chiral HPLC:	isocratic elution with *n*-hexane / isopropanol, 98:2: *t*_R_ = 5.1 min, >99% *ee* (254 nm).

[*α*]*_D_^23^*:	-0.9 (*c* = 1.1, methanol).

**(*S*)-3-(2,4-Di-*tert*-butylphenoxy)-2-methylpropan-1-ol (ent-11d)**

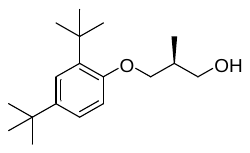

(*S*)-3-(2,4-Di-tert-butylphenoxy)-2-methylpropan-1-ol **(ent-11d)** was synthesized analogously as described for **11d**, starting from (*S*)-3-bromo-2-methylpropan-1-ol. The analytical data were in accordance.

chiral HPLC:	isocratic elution with *n*-hexane / isopropanol, 98:2: *t*_R_ = 5.4 min, 98% *ee* (254 nm).

[*α*]*_D_^24^*:	+1.6 (*c* = 1.6, methanol).

**(*R*)-3-(4-Fluorophenoxy)-2-methylpropan-1-ol (11e)** ^2^

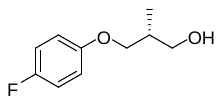

To a stirred solution of 4-fluorophenol (150 mg, 1.34 mmol) in DMF (3 mL) in a flame-dried flask sodium hydride (60% in mineral oil; 61.0 mg, 1.52 mmol) was added. After stirring the suspension for 30 min at rt, (*R*)-3-bromo-2-methylpropan-1-ol (134 µL, 1.27 mmol) was added. The mixture was stirred for additional 31 h, then 2N NaOH was added and extraction with *tert*-butyl methyl ether (3×) was performed. The combined organic layers were washed with water and brine, dried over Na_2_SO_4_ and concentrated. After purification by flash-chromatography (isohexane / *tert*-butyl methyl ether, 7:3) compound **11e** (125 mg, 0.68 mmol, 53%) was obtained as pale yellow liquid.

ESI-MS:	*m/z* 207.5 [M+Na]^+^.

HR-ESI-MS:	*m/z* [M+H]^+^ calcd. 207.0797 for C_10_H_13_FNaO_2_, found 207.0794.

^1^H-NMR:	(400 MHz, CDCl_3_) *δ* 1.04 (d, *J* = 7.0 Hz, 3H), 1.83 (dd, *J* = 5.6, 5.6 Hz, 1H), 2.13 – 2.26 (m, 1H), 3.69 – 3.73 (m, 2H), 3.90 (dd, *J* = 9.0, 6.6 Hz, 1H), 3.93 (dd, *J* = 9.0, 5.5 Hz, 1H), 6.81 – 6.88 (m, 2H), 6.93 – 7.01 (m, 2H).

^13^C-NMR:	(DEPTQ, 101 MHz, CDCl_3_) *δ* 13.7, 35.8, 66.2, 71.9, 115.6 (d, *J* = 8 Hz), 115.9 (d, *J* = 23 Hz), 155.1 (d, *J* = 2 Hz), 157.4 (d, *J* = 238 Hz).

chiral HPLC:	isocratic elution with *n*-hexane / isopropanol, 98:2: *t*_R_ = 10.9 min, >99% *ee* (254 nm).

[*α*]*_D_^25^*:	+3.2 (*c* = 0.9, chloroform).

**(*R*)-3-(4-Bromophenoxy)-2-methylpropan-1-ol (11f)**

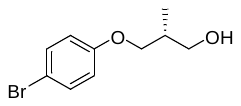

To a stirred solution of 4-bromophenol (150 mg, 0.87 mmol) in DMF (3 mL) in a flame-dried flask sodium hydride (60% in mineral oil; 39.8 mg, 1.00 mmol) was added. After stirring the suspension for 30 min at rt, (*R*)-3-bromo-2-methylpropan-1-ol (86.5 µL, 0.83 mmol) was added. The mixture was stirred for additional 31 h, then 2N NaOH was added and extraction with *tert*-butyl methyl ether (3×) was performed. The combined organic layers were washed with water and brine, dried over Na_2_SO_4_ and concentrated. After purification by flash-chromatography (isohexane / *tert*-butyl methyl ether, 7:3) compound **11f** (112 mg, 0.46 mmol, 55%) was obtained as pale yellow liquid.

ESI-MS:	*m/z* 274.3 [M+Na]^+^.

HR-ESI-MS:	*m/z* [M+H]^+^ calcd. 266.9991 for C_10_H_13_BrNaO_2_, found 266.9994.

^1^H-NMR:	(400 MHz, CDCl_3_) *δ* 1.04 (d, *J* = 7.0 Hz, 3H), 1.73 (dd, *J* = 5.6, 5.6 Hz, 1H), 2.13 – 2.25 (m, 1H), 3.68 – 3.72 (m, 2H), 3.90 – 3.93 (m, 2H), 6.76 – 6.82 (m, 2H), 7.34 – 7.40 (m, 2H).

^13^C-NMR:	(DEPTQ, 101 MHz, CDCl_3_) *δ* 13.6, 35.6, 65.8, 71.2, 113.0, 116.3, 132.2, 158.0.

chiral HPLC:	isocratic elution with *n*-hexane / isopropanol, 98:2: *t*_R_ = 11.8 min, >99% *ee* (254 nm).

[*α*]*_D_^25^*:	+1.7 (*c* = 0.6, chloroform).

**(*R*)-3-([1,1'-Biphenyl]-4-yloxy)-2-methylpropan-1-ol (11g)**

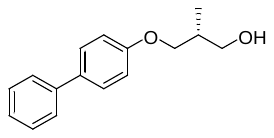

To a stirred solution of [1,1'-biphenyl]-4-ol (150 mg, 0.88 mmol) in DMF (4 mL) in a flame-dried flask sodium hydride (60% in mineral oil; 41.0 mg, 1.01 mmol) was added. After stirring the suspension for 30 min at rt, (*R*)-3-bromo-2-methylpropan-1-ol (88.0 µL, 0.84 mmol) was added. The mixture was stirred for additional 31 h, then 2N NaOH was added and extraction with *tert*-butyl methyl ether (3×) was performed. The combined organic layers were washed with water and brine, dried over Na_2_SO_4_ and concentrated. After purification by flash-chromatography (isohexane / *tert*-butyl methyl ether, 7:3) compound **11g** (104 mg, 0.42 mmol, 50%) was obtained as white resin.

ESI-MS:	*m/z* 243.1 [M+H]^+^.

HR-ESI-MS:	*m/z* [M+H]^+^ calcd. 265.1199 for C_16_H_18_NaO_2_, found 265.1199.

^1^H-NMR:	(400 MHz, CDCl_3_) *δ* 1.06 (d, *J* = 7.0 Hz, 3H), 1.85 (dd, *J* = 5.6, 5.6 Hz, 1H), 2.18 – 2.30 (m, 1H), 3.70 – 3.76 (m, 2H), 3.98 (dd, *J* = 9.1, 6.8 Hz, 1H), 4.02 (dd, *J* = 9.1, 5.5 Hz, 1H), 6.96 – 7.02 (m, 2H), 7.31 (m, 1H), 7.45 – 7.39 (m, 2H), 7.58 – 7.50 (m, 4H).

^13^C-NMR:	(DEPTQ, 101 MHz, CDCl_3_) *δ* 13.6, 35.7, 66.2, 71.3, 114.7, 126.7, 126.7, 128.2, 128.7, 134.0, 140.8, 158.4.

chiral HPLC:	isocratic elution with *n*-hexane / isopropanol, 98:2: *t*_R_ = 24.3 min, 99% *ee* (254 nm).

[*α*]*_D_^25^*:	+2.1 (*c* = 0.6, chloroform).

**(*R*)-3-(Benzofuran-5-yloxy)-2-methylpropan-1-ol (11h)**

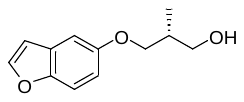

To a stirred solution of benzofuran-5-ol (150 mg, 1.12 mmol) in DMF (3 mL) in a flame-dried flask sodium hydride (60% in mineral oil; 50.9 mg, 1.27 mmol) was added. After stirring the suspension for 30 min at rt, (*R*)-3-bromo-2-methylpropan-1-ol (112 µL, 1.06 mmol) was added. The mixture was stirred for additional 18 h, then 2N NaOH was added and extraction with *tert*-butyl methyl ether (3×) was performed. The combined organic layers were washed with water and, brine, dried over Na_2_SO_4_ and concentrated. After purification by flash-chromatography (isohexane / *tert*-butyl methyl ether, 7:3) compound **11h** (117 mg, 0.57 mmol, 54%) was obtained as yellow resin.

ESI-MS:	*m/z* 207.0 [M+H]^+^.

HR-ESI-MS:	*m/z* [M+H]^+^ calcd. 229.0835 for C_12_H_14_NaO_3_, found 229.0837.

^1^H-NMR:	(400 MHz, CDCl_3_) *δ* 1.05 (d, *J* = 7.0 Hz, 3H), 2.00 (dd, *J* = 5.6, 5.6 Hz, 1H), 2.24 (m, 1H), 3.73 (m, 2H), 3.96 (dd, *J* = 9.1, 7.0 Hz, 1H), 4.01 (dd, *J* = 9.1, 5.6 Hz, 1H), 6.70 (dd, *J* = 2.2, 0.9 Hz, 1H), 6.90 (dd, *J* = 8.9, 2.6 Hz, 1H), 7.08 (d, *J* = 2.6 Hz, 1H), 7.39 (ddd, *J* = 8.9, 0.9, 0.5 Hz, 1H), 7.59 (d, *J* = 2.2 Hz, 1H).

^13^C-NMR	(DEPTQ, 101 MHz, CDCl_3_) *δ* 13.6, 35.7, 66.4, 72.4, 104.6, 106.7, 111.8, 113.5, 128.0, 145.8, 150.0, 155.1.

chiral HPLC:	isocratic elution with *n*-hexane / isopropanol, 98:2: *t*_R_ = 18.5 min, >99% *ee* (254 nm).

[*α*]*_D_^24^*:	+3.8 (*c* = 1.5, chloroform).

**(*S*)-2-Methyl-3-phenoxypropanoic acid (12a)** ^1^

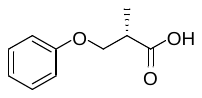

**11a** (171 mg, 1.03 mmol) was dissolved in acetone (9 mL, 0.11 M). Jones reagent (816 µL, 1.26 M, 1.03 mmol, **GPA**) was added at 0 °C and the mixture was stirred for 3 h at 0 °C. After terminating the reaction by the addition of 2-propanol, the mixture was filtered through a pad of celite, which was further washed with ethyl acetate. The combined organic layers were extracted with 1N NaOH. After acidifying the aqueous layer using 2N HCl and extraction with ethyl acetate (2×), the combined organic layers were washed with water and brine, dried over Na_2_SO_4_ and concentrated. Purification by flash-chromatography (CH_2_Cl_2_ / MeOH + 0.1% HCOOH, 98:2) afforded compound **12a** (105 mg, 0.58 mmol, 57%) as pale yellow liquid, which solidified upon standing (beige resin).

ESI-MS:	*m/z* 203.2 [M+Na]^+^.

^1^H-NMR:	(400 MHz, CDCl_3_) *δ* 1.35 (d, *J* = 7.1 Hz, 3H), 3.00 (ddq, *J* = 7.1, 6.7, 5.9 Hz, 1H), 4.03 (dd, *J* = 9.1, 5.9 Hz, 1H), 4.20 (dd, *J* = 9.1, 6.7 Hz, 1H), 6.89 – 6.93 (m, 2H), 6.94 – 6.98 (m, 1H), 7.26 – 7.31 (m, 2H).

^13^C-NMR:	(DEPTQ, 101 MHz, CDCl_3_) *δ* 13.8, 39.7, 69.1, 114.7, 121.1, 129.5, 158.5, 180.2.

[*α*]*_D_^22^*:	+2.0 (*c* = 2.5, chloroform).

The analytical data of this compound are in accordance with those reported in the literature.

**(*R*)-2-Methyl-3-phenoxypropanoic acid (ent-12a)** ^1^

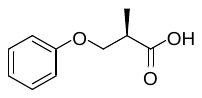

(*R*)-2-Methyl-3-phenoxypropanoic acid **(ent-12a)** was synthesized analogously as described for **12a**, starting from **ent-11a**. The analytical data of this compound are in accordance with those reported in the literature.

**(*S*)-2-Methyl-3-(naphthalen-2-yloxy)propanoic acid (12b)** ^3^

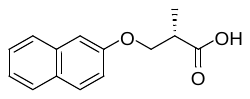

**11b** (191 mg, 0.88 mmol) was dissolved in acetone (8 mL, 0.11 M). Jones reagent (701 µL, 1.26 M, 0.88 mmol, **GPA**) was added at 0 °C and the mixture was stirred for 5 h at 0 °C. After terminating the reaction by the addition of 2-propanol, the mixture was filtered through a pad of celite, which was further washed with ethyl acetate. The combined organic layers were extracted with 1N NaOH. After acidifying the aqueous layer using 2N HCl and extraction with ethyl acetate (2×), the combined organic layers were washed with water and brine, dried over Na_2_SO_4_ and concentrated. Compound **12b** (130 mg, 0.56 mmol, 64%) was obtained as an orange solid, which was used for the next step without further purification.

ESI-MS:	*m/z* 252.9 [M+Na]^+^.

HR-ESI-MS:	*m/z* [M+H]^+^ calcd. 253.0835 for C_14_H_14_NaO_3_, found 253.0837.

^1^H-NMR:	(400 MHz, CDCl_3_) *δ* 1.39 (d, *J* = 7.1 Hz, 3H), 3.06 (ddq, *J* = 7.1, 6.7, 5.9 Hz, 1H), 4.14 (dd, *J* = 9.1, 5.9 Hz, 1H), 4.32 (dd, *J* = 9.1, 6.7 Hz, 1H), 7.12 – 7.18 (m, 2H), 7.34 (ddd, *J* = 8.2, 6.9, 1.3 Hz, 1H), 7.43 (ddd, *J* = 8.2, 6.9, 1.3 Hz, 1H), 7.70 – 7.79 (m, 3H).

^13^C-NMR:	(DEPTQ, 151 MHz, CDCl_3_) *δ* 13.9, 39.7, 69.2, 107.0, 118.8, 123.8, 126.4, 126.8, 127.6, 129.1, 129.4, 134.5, 156.5, 179.9.

[*α*]*_D_^24^*:	-3.5 (*c* = 0.6, chloroform).

**(*R*)-2-Methyl-3-(naphthalen-2-yloxy)propanoic acid (ent-12b)**

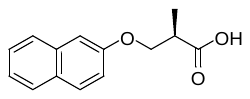

(*R*)-2-Methyl-3-(naphthalen-2-yloxy)propanoic acid **(ent-12b)** was synthesized analogously as described for **12b**, starting from **ent-11b**. The analytical data were in accordance.

[*α*]*_D_^24^*:	+2.5 (*c* = 0.7, chloroform).

**(*S*)-2-Methyl-3-(3-(trifluoromethyl)phenoxy)propanoic acid (12c)**

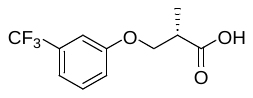

Crude **11c** (141 mg, 0.60 mmol) was dissolved in acetone (6 mL, 0.10 M). Jones reagent (476 µL, 1.26 M, 0.60 mmol, **GPA**) was added at 0 °C and the mixture was stirred for 4.5 h at 0 °C. After terminating the reaction by the addition of 2-propanol, the mixture was filtered through a pad of celite, which was further washed with ethyl acetate. The combined organic layers were extracted with 1N NaOH. After acidifying the aqueous layer using 2N HCl and extraction with ethyl acetate (2×), the combined organic layers were washed with water and brine, dried over Na_2_SO_4_ and concentrated. Compound **12c** (65.2 mg, 0.26 mmol) was obtained as yellow liquid, containing a minor amount of a residual impurity (confirmed by NMR). The product was used for the next step without further purification.

ESI-MS:	*m/z* 271.0 [M+Na]^+^.

HR-ESI-MS:	*m/z* [M+H]^+^ calcd. 271.0552 for C_11_H_11_F_3_NaO_3_, found 271.0553.

^1^H-NMR:	(400 MHz, CDCl_3_) *δ* 1.36 (d, *J* = 7.2 Hz, 3H), 3.01 (ddq, *J* = 7.2, 6.8, 5.6 Hz, 1H), 4.07 (dd, *J* = 9.0, 5.6 Hz, 1H), 4.22 (dd, *J* = 9.0, 6.8 Hz, 1H), 7.05 – 7.10 (m, 1H), 7.12 – 7.15 (m, 1H), 7.20 – 7.24 (m, 1H), 7.36 – 7.43 (m, 1H).

^13^C-NMR:	(DEPTQ, 151 MHz, CDCl_3_) *δ* 13.7, 39.6, 69.4, 111.4 (q, *J* = 4 Hz), 117.81 (q, *J* = 4 Hz), 118.1, 123.9 (q, *J* = 272 Hz), 130.0, 131.9 (q, *J* = 32 Hz), 158.6, 179.4.

**(*R*)-2-Methyl-3-(3-(trifluoromethyl)phenoxy)propanoic acid (ent-12c)**

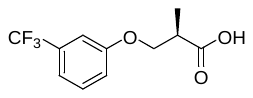

(*R*)-2-Methyl-3-(3-(trifluoromethyl)phenoxy)propanoic acid **(ent-12c)** was synthesized analogously as described for **12c**, starting from crude **ent-11c**. The analytical data were in accordance.

**(*S*)-3-(2,4-Di-*tert*-butylphenoxy)-2-methylpropanoic acid (12d)**

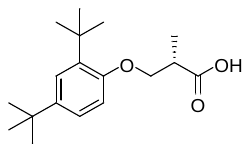

**11d** (82.1 mg, 0.29 mmol) was dissolved in acetone (2.5 mL, 0.12 M). Jones reagent (230 µL, 1.26 M, 0.29 mmol, **GPA**) was added at 0 °C and the mixture was stirred for 4 h at 0 °C. After terminating the reaction by the addition of 2-propanol, the mixture was filtered through a pad of celite, which was further washed with ethyl acetate. The combined organic layers were extracted with 1N NaOH. After acidifying the aqueous layer using 2N HCl and extraction with ethyl acetate (2×), the combined organic layers were washed with water and brine, dried over Na_2_SO_4_ and concentrated. Compound **12d** (50.8 mg, 0.17 mmol, 60%) was obtained as a yellow oil, which solidified upon standing (beige wax) and which was used for the next step without further purification.

ESI-MS:	*m/z* 315.1 [M+Na]^+^.

HR-ESI-MS:	*m/z* [M+H]^+^ calcd. 315.1931 for C_18_H_28_NaO_3_, found 315.1932.

^1^H-NMR:	(400 MHz, CDCl_3_) *δ* 1.30 (s, 9H), 1.35 (s, 9H), 1.37 (d, *J* = 7.2 Hz, 3H), 3.04 (ddq, *J* = 7.2, 6.5, 5.5 Hz, 1H), 4.09 (dd, *J* = 8.8, 5.5 Hz, 1H), 4.18 (dd, *J* = 8.8, 6.5 Hz, 1H), 6.79 (d, *J* = 8.5 Hz, 1H), 7.17 (dd, *J* = 8.5, 2.5 Hz, 1H), 7.33 (d, *J* = 2.5 Hz, 1H).

^13^C-NMR:	(DEPTQ, 101 MHz, CDCl_3_) *δ* 14.3, 30.0, 31.7, 34.4, 35.1, 40.1, 69.5, 111.1, 123.4, 124.2, 137.3, 143.0, 155.0, 180.1.

**(*R*)-3-(2,4-Di-tert-butylphenoxy)-2-methylpropanoic acid (ent-12d)**

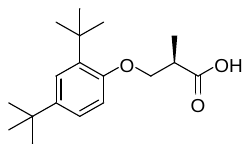

(*R*)-3-(2,4-Di-tert-butylphenoxy)-2-methylpropanoic acid **(ent-12d)** was synthesized analogously as described for **12d**, starting from **ent-11d**. The analytical data were in accordance.

**(*S*)-3-(4-Fluorophenoxy)-2-methylpropanoic acid (12e)**

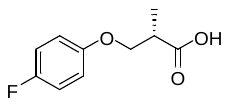

**11e** (120 mg, 0.65 mmol) was dissolved in acetone (6 mL, 0.11 M). Jones reagent (516 µL, 1.26 M, 0.65 mmol, **GPA**) was added at 0 °C and the mixture was stirred for 4.5 h at 0 °C. After terminating the reaction by the addition of 2-propanol, the mixture was filtered through a pad of celite, which was further washed with ethyl acetate. The combined organic layers were extracted with 1N NaOH. After acidifying the aqueous layer using 2N HCl and extraction with ethyl acetate (2×), the combined organic layers were washed with water and brine, dried over Na_2_SO_4_ and concentrated. Compound **12e** (76.5 mg, 0.39 mmol, 59%) was obtained as a yellow oil. The compound was used for the next step without further purification.

ESI-MS:	*m/z* 221.4 [M+Na]^+^.

HR-ESI-MS:	*m/z* [M+H]^+^ calcd. 221.0584 for C_10_H_11_FNaO_3_, found 221.0586.

^1^H-NMR:	(400 MHz, CDCl_3_) *δ* 1.33 (d, *J* = 7.1 Hz, 3H), 2.97 (ddq, *J* = 7.1, 6.8, 5.7 Hz, 1H), 3.99 (dd, *J* = 9.0, 5.7 Hz, 1H), 4.15 (dd, *J* = 9.0, 6.8 Hz, 1H), 6.81 – 6.88 (m, 2H), 6.93 – 7.00 (m, 2H).

^13^C-NMR:	(DEPTQ, 101 MHz, CDCl_3_) *δ* 13.7, 39.7, 69.9, 115.7 (d, *J* = 2 Hz), 115.9 (d, *J* = 17 Hz), 154.6 (d, *J* = 2 Hz), 157.5 (d, *J* = 239 Hz), 180.2.

[*α*]*_D_^24^*:	+3.1 (*c* = 0.9, chloroform).

**(*S*)-3-(4-Bromophenoxy)-2-methylpropanoic acid (12f)**

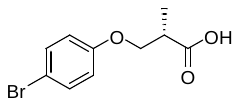

**11f** (103 mg, 0.42 mmol) was dissolved in acetone (4 mL, 0.11 M). Freshly prepared Jones reagent (333 µL, 1.26 M, 0.42 mmol, **GPA**) was added at 0 °C and the mixture was stirred for 5 h at 0 °C. After terminating the reaction by the addition of 2-propanol, the mixture was filtered through a pad of celite, which was further washed with ethyl acetate. The combined organic layers were extracted with 1N NaOH. After acidifying the aqueous layer using 2N HCl and extraction with ethyl acetate (2×), the combined organic layers were washed with water and brine, dried over Na_2_SO_4_ and concentrated. Compound **12f** (56.4 mg, 0.22 mmol, 52%) was obtained as brown oil, which was used for the next step without further purification.

ESI-MS:	*m/z* 281.2 [M+Na]^+^.

HR-ESI-MS:	*m/z* [M+H]^+^ calcd. 280.9784 for C_10_H_11_BrNaO_3_, found 280.9784.

^1^H-NMR:	(400 MHz, CDCl_3_) *δ* 1.33 (d, *J* = 7.2 Hz, 3H), 2.97 (ddq, *J* = 7.2, 6.8, 5.8 Hz, 1H), 3.99 (dd, *J* = 9.0, 5.7 Hz, 1H), 4.15 (dd, *J* = 9.0, 6.8 Hz, 1H), 6.76 – 6.81 (m, 2H), 7.34 – 7.40 (m, 2H).

^13^C-NMR:	(DEPTQ, 101 MHz, CDCl_3_) *δ* 13.7, 39.6, 69.4, 113.3, 116.4, 132.3, 157.6, 180.0.

[*α*]*_D_^24^*:	+1.4 (*c* = 1.0, chloroform).

**(*S*)-3-([1,1'-Biphenyl]-4-yloxy)-2-methylpropanoic acid (12g)** ^4^

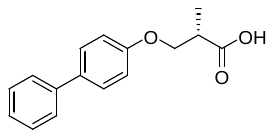

**11g** (120 mg, 0.50 mmol) was dissolved in acetone (5 mL, 0.10 M). Freshly prepared Jones reagent (393 µL, 1.26 M, 0.50 mmol, **GPA**) was added at 0 °C and the mixture was stirred for 4.5 h at 0 °C. After terminating the reaction by the addition of 2-propanol, the mixture was filtered through pad of celite, which was further washed with ethyl acetate. The combined organic layers were extracted with 1N NaOH. After acidifying the aqueous layer using 2N HCl and extraction with ethyl acetate (2x), the combined organic layers were washed with water and brine, dried over Na_2_SO_4_ and concentrated. Compound **12g** (61.4 mg, 0.24 mmol, 48%) was obtained as a white solid, which was used for the next step without further purification.

ESI-MS:	*m/z* 279.1 [M+Na]^+^.

HR-ESI-MS:	*m/z* [M+H]^+^ calcd. 279.0992 for C_16_H_16_NaO_3_, found 279.0994.

^1^H-NMR:	(400 MHz, CDCl_3_) *δ* 1.37 (d, *J* = 7.1 Hz, 3H), 3.03 (ddq, *J* = 7.1, 6.8, 5.9 Hz, 1H), 4.08 (dd, *J* = 9.1, 5.9 Hz, 1H), 4.24 (dd, *J* = 9.1, 6.8 Hz, 1H), 6.96 – 7.02 (m, 2H), 7.28 – 7.34 (m, 1H), 7.38 – 7.45 (m, 2H), 7.49 – 7.58 (m, 4H).

^13^C-NMR:	(DEPTQ, 101 MHz, CDCl_3_) *δ* 13.8, 39.6, 69.3, 114.9, 126.7, 126.7, 128.2, 128.7, 134.2, 140.7, 158.0, 179.2.

[*α*]*_D_^24^*:	+1.1 (*c* = 0.5, chloroform).

**(*S*)-3-(Benzofuran-5-yloxy)-2-methylpropanoic acid (12h)**

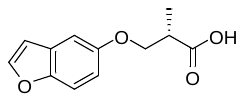

**11h** (47.0 mg, 0.23 mmol) was dissolved in acetone (2.3 mL, 0.10 M). Freshly prepared Jones reagent (181 µL, 1.26 M, 0.23 mmol, **GPA**) was added at 0 °C and the mixture was stirred for 4.5 h at 0 °C. After terminating the reaction by the addition of 2-propanol, the mixture was filtered through a pad of celite, which was further washed with ethyl acetate. The combined organic layers were extracted with 1N NaOH. After acidifying the aqueous layer using 2N HCl and extraction with ethyl acetate (2×), the combined organic layers were washed with water and brine, dried over Na_2_SO_4_ and concentrated. Compound **12h** (26.4 mg, 0.12 mmol) was obtained as an orange oil, still containing a minor amount of residual contaminations (confirmed by NMR). The product was used crude for the next step without further purification.

ESI-MS:	*m/z* 243.0 [M+Na]^+^.

^1^H-NMR:	(400 MHz, CDCl_3_) *δ* 1.35 (d, *J* = 7.1 Hz, 3H), 3.01 (ddq, *J* = 7.1, 6.8, 5.8 Hz, 1H), 4.06 (dd, *J* = 9.1, 5.8 Hz, 1H), 4.23 (dd, *J* = 9.1, 6.8 Hz, 1H), 6.70 (dd, *J* = 2.2, 0.9 Hz, 1H), 6.91 (dd, *J* = 8.9, 2.6 Hz, 1H), 7.08 (d, *J* = 2.6 Hz, 1H), 7.38 (d, *J* = 8.9 Hz, 1H), 7.59 (d, *J* = 2.2 Hz, 1H).

^13^C-NMR:	(DEPTQ, 101 MHz, CDCl_3_) *δ* 13.8, 39.8, 70.3, 105.0, 106.7, 111.8, 113.7, 127.9, 145.8, 150.2, 154.8, 180.3.

**(*S*)-1-(3-Amino-6-fluoro-1*H*-indazol-1-yl)-2-methyl-3-phenoxypropan-1-one ((*S*)-‘853, 1a)**

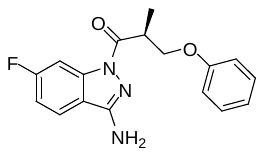

To a solution of compound **12a** (25.0 mg, 0.14 mmol) in DMF (2 mL) were added HOAt (19.1 mg, 0.14 mmol) and EDC × HCl (27.1 mg, 0.14 mmol) at rt. The resulting solution was stirred for 10 min at rt and then 6‑fluoro-1*H*-indazole-3-amine (**13**, 17.5 mg, 0.12 mmol) in DMF (0.5 mL) was added. After stirring the reaction mixture at rt for 1 h, the solvent was removed by lyophilization. Purification by preparative HPLC using a solvent system of CH_3_OH / 0.1% aq. HCOOH, a flow rate of 10 mL /min and a gradient of 50% to 80% CH_3_OH in 15 min, 80% to 95% CH_3_OH in 2 min (*t*_R_ = 11.5 min, λ = 254 nm) afforded **(*S*)-‘853 (1a)** (18.1 mg, 57.9 μmol, 48%) as an off-white solid.

ESI-MS:	*m/z* 314.1 [M+H]^+^.

HR-ESI-MS:	*m/z* [M+H]^+^ calcd. 314.1299 for C_17_H_17_FN_3_O_2_, found 314.1302.

IR (NaCl):	3452, 3350, 3226, 1686, 1628, 1436, 1426 cm^-1^.

m.p.:	45 °C.

^1^H-NMR:	(600 MHz, DMSO-d_6_) *δ* 1.29 (d, *J* = 7.1 Hz, 3H), 4.04 – 4.11 (m, 2H), 4.36 – 4.41 (m, 1H), 6.61 (s, 2H), 6.90 – 6.96 (m, 3H), 7.25 (ddd, *J* = 9.2, 8.7, 2.2 Hz, 1H), 7.25 – 7.29 (m, 2H), 7.94 (dd, *J* = 8.7, 5.4 Hz, 1H), 7.96 (dd, *J* = 9.7, 2.2 Hz, 1H).

^13^C-NMR:	(DEPTQ, 151 MHz, DMSO-d_6_) *δ* 13.9, 37.7, 68.8, 101.8 (d, *J* = 28 Hz), 112.2 (d, *J* = 24 Hz), 114.5, 117.0, 120.7, 122.7 (d, *J* = 11 Hz), 129.5, 139.8 (d, *J* = 13 Hz), 152.6, 158.3, 163.2 (d, *J* = 244 Hz), 172.4.

HPLC:	system 2: λ = 254 nm, *t*_R_ = 18.2 min, purity: 99%.

chiral HPLC:	isocratic elution with *n*-hexane / isopropanol + 0.1% ethylene diamine, 9:1: *t*_R_ = 7.4 min, 99% *ee* (254 nm).

[*α*]*_D_^23^*:	-4.1 (*c* = 0.2, methanol).

**(*R*)-1-(3-Amino-6-fluoro-1*H*-indazol-1-yl)-2-methyl-3-phenoxypropan-1-one ((*R*)-‘853, ent-1a)**

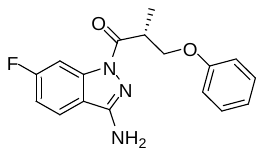

(*R*)-1-(3-Amino-6-fluoro-1H-indazol-1-yl)-2-methyl-3-phenoxypropan-1-one **((*S*)-‘853, ent-1a)** was synthesized analogously as described for **1a**, starting from **ent-12a**. The analytical data were in accordance.

HR-ESI-MS:	*m/z* [M+H]^+^ calcd. 314.1299 for C_17_H_17_FN_3_O_2_, found 314.1300.

HPLC:	system 2: λ = 254 nm,	*t*_R_ = 18.4 min, purity: >99%.

chiral HPLC:	isocratic elution with *n*-hexane / isopropanol + 0.1% ethylene diamine, 9:1: *t*_R_ = 6.4 min, 99% *ee* (254 nm).

[*α*]*_D_^24^*:	+4.7 (*c* = 0.3, methanol).

**(*S*)-1-(3-Amino-6-fluoro-1*H*-indazol-1-yl)-2-methyl-3-(naphthalen-2-yloxy)propan-1-one ((*S*)‑SX240, 1b)**

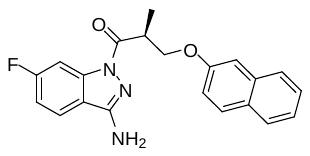

To a solution of compound **12b** (25.2 mg, 0.11 mmol) in DMF (2 mL) were added HOAt (15.3 mg, 0.11 mmol) and EDC × HCl (21.3 mg, 0.11 mmol) at rt. The resulting solution was stirred for 10 min at rt and then 6‑fluoro-1*H*-indazole-3-amine (**13**, 13.8 mg, 91.2 µmol) in DMF (0.5 mL) were added. After stirring the mixture at rt for 1.5 h, the solvent was removed by lyophilization. Purification by preparative HPLC using a solvent system of CH_3_OH / 0.1% aq. TFA, a flow rate of 10 mL / min and a gradient of 50% to 80% CH_3_OH in 15 min, 80% to 95% CH_3_OH in 2 min, 95% CH_3_OH for 5 min (*t*_R_ = 21.0 min, λ = 254 nm) afforded **(*S*)-SX240** (**1b**, 11.4 mg, 31.3 μmol, 34%) as a white solid.

ESI-MS	*m/z* 364.2 [M+H]^+^.

HR-ESI-MS:	*m/z* [M+H]^+^ calcd. 364.1456 for C_21_H_19_FN_3_O_2_, found 364.1460.

IR (NaCl):	3458, 2924, 2851, 1629, 1424 cm^-1^.

m.p.:	59 °C.

^1^H-NMR:	(600 MHz, DMSO-d_6_) *δ* 1.34 (d, *J* = 7.0 Hz, 1H), 4.16 (ddq, *J* = 7.8, 7.0, 5.5 Hz, 1H), 4.21 (dd, *J* = 9.1, 5.5 Hz, 1H), 4.52 (dd, *J* = 9.1, 7.8 Hz, 1H), 6.62 (s, 2H), 7.11 (dd, *J* = 9.0, 2.5 Hz, 1H), 7.26 (ddd, *J* = 9.0, 8.9, 2.5 Hz, 1H), 7.34 (ddd, *J* = 8.1, 6.9, 1.2 Hz, 1H), 7.39 (d, *J* = 2.5 Hz, 1H), 7.45 (ddd, *J* = 8.1, 6.9, 1.2 Hz, 1H), 7.78 – 7.83 (m, 3H), 7.94 – 7.99 (m, 2H).

^13^C-NMR:	(DEPTQ, 151 MHz, DMSO-d_6_) *δ* 14.0, 37.7, 69.0, 101.8 (d, *J* = 28 Hz), 106.9, 112.2 (d, *J* = 25 Hz), 117.0, 118.6, 122.7 (d, *J* = 11 Hz), 123.6, 126.4, 126.7, 127.5, 128.5, 129.3, 134.2, 139.8 (d, *J* = 13 Hz), 152.7, 156.2, 163.2 (d, *J* = 245 Hz), 172.5.

^19^F-NMR:	(377 MHz, DMSO-d_6_) *δ* -110.5.

HPLC:	system 1: λ = 254 nm, *t*_R_ = 20.5 min, purity: 95%

chiral HPLC:	isocratic elution with *n*-hexane / isopropanol, 9:1: *t*_R_ = 9.7 min, >99% *ee* (254 nm).

[*α*]*_D_^23^*:	-15.8 (*c* = 0.4, methanol).

**(*R*)-1-(3-Amino-6-fluoro-1*H*-indazol-1-yl)-2-methyl-3-(naphthalen-2-yloxy)propan-1-one ((*R*)‑SX240, ent-1b)**

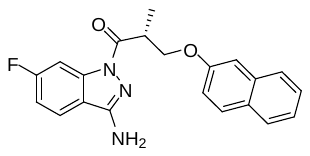

(*R*)-1-(3-Amino-6-fluoro-1H-indazol-1-yl)-2-methyl-3-(naphthalen-2-yloxy)propan-1-one **((*R*)‑SX240, ent-1b)** was synthesized analogously as described for **(*S*)-SX240 (1b)**, starting from **ent-12b**. The analytical data were in accordance.

HR-ESI-MS:	*m/z* [M+H]^+^ calcd. 364.1456 for C_21_H_19_FN_3_O_2_, found 364.1456.

HPLC:	system 1: λ = 254 nm, *t*_R_ = 20.9 min, purity: >99%.

chiral HPLC:	isocratic elution with *n*-hexane / isopropanol, 9:1: *t*_R_ = 8.2 min, 98% *ee* (254 nm)

[*α*]*_D_^23^*:	+21.0 (*c* = 0.4, methanol).

**(*S*)-1-(3-Amino-6-fluoro-1*H*-indazol-1-yl)-2-methyl-3-(3-(trifluoromethyl)phenoxy)propan-1-one ((*S*)-SX245, 1c)**

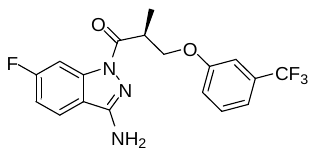

To a solution of crude **12c** (14.5 mg, 58.4 µmol) in DMF (1.5 mL) were added HOAt (7.9 mg, 58.4 µmol) and EDC × HCl (11.2 mg, 58.4 µmol) at rt. The resulting solution was stirred for 10 min at rt and then 6‑fluoro-1*H*-indazole-3-amine (**13**, 10.6 mg, 70.1 µmol) in DMF (0.5 mL) was added. After stirring the reaction mixture at rt for 2 h, the solvent was removed by lyophilization. Purification by preparative HPLC using a solvent system of CH_3_OH / 0.1% aq. HCOOH, a flow rate of 10 mL / min and a gradient of 50% to 80% CH_3_OH in 15 min, 80% to 95% CH_3_OH in 2 min, 95% CH_3_OH for 3 min (*t*_R_ = 18.5 min, λ = 254 nm) yielded **(*S*)-SX245** (**1c**, 8.2 mg, 21.5 μmol, 37%) as a white solid.

ESI-MS:	m/z 382.1 [M+H]^+^.

HR-ESI-MS:	*m/z* [M+H]^+^ calcd. 382.1173 for C_18_H_16_F_4_N_3_O_2_, found 382.1172.

IR (NaCl):	3401, 3350, 2922, 2850, 1672, 1625, 1423, 1317, 1125 cm^-1^.

m.p.:	121 °C.

^1^H-NMR:	(600 MHz, DMSO-d_6_) *δ* 1.30 (d, *J* = 7.0 Hz, 3H), 4.10 (ddq, *J* = 8.0, 7.0, 5.4 Hz, 1H), 4.17 (dd, *J* = 9.3, 5.4 Hz, 1H), 4.47 (dd, *J* = 9.3, 8.0 Hz, 1H), 6.61 (s, 2H), 7.21 – 7.30 (m, 4H), 7.48 – 7.53 (m, 1H), 7.89 – 7.99 (m, 2H).

^13^C-NMR:	(DEPTQ, 151 MHz, DMSO-d_6_) *δ* 13.9, 37.7, 69.4, 101.8 (d, *J* = 28 Hz), 111.0 (q, *J* = 4 Hz), 112.3 (d, *J* = 25 Hz), 117.0, 117.2 (q, *J* = 4 Hz), 118.8, 122.7 (d, *J* = 11 Hz), 123.9 (q, J = 273 Hz), 130.3 (q, *J* = 32 Hz), 130.7, 139.8 (d, *J* = 14 Hz), 152.7, 158.6, 163.2 (d, *J* = 245 Hz), 172.2.

^19^F-NMR:	(377 MHz, DMSO-d_6_) *δ* -60.6, -110.5.

HPLC:	system 2: λ = 254 nm, *t*_R_ = 19.6 min, purity: 98%.

chiral HPLC:	isocratic elution with *n*-hexane / isopropanol, 98:2: *t_R_* = 13.4 min, >99% ee (254 nm).

[*α*]*_D_^25^*:	-5.1 (c = 0.1, methanol).

**(*R*)-1-(3-Amino-6-fluoro-1*H*-indazol-1-yl)-2-methyl-3-(3-(trifluoromethyl)phenoxy)propan-1-one ((*R*)-SX245, ent-1c)**

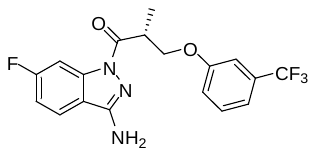

(*R*)-1-(3-Amino-6-fluoro-1H-indazol-1-yl)-2-methyl-3-(3-(trifluoromethyl)phenoxy)propan-1-one **((*R*)-SX245, ent-1c)** was synthesized analogously as described for **(*S*)-SX245 (1c)**, starting from crude **ent-12c**. The analytical data were in accordance.

HR-ESI-MS:	*m/z* [M+H]^+^ calcd. 382.1173 for C_18_H_16_F_4_N_3_O_2_, found 382.1176.

HPLC:	system 2: λ = 254 nm, *t_R_* = 19.2 min, purity: >99%.

chiral HPLC:	isocratic elution with *n*-hexane / isopropanol, 98:2: *t_R_* = 12.1 min, >99% ee (254 nm).

[*α*]*_D_^26^*:	+5.8 (c = 0.2, methanol).

**(*S*)-1-(3-Amino-6-fluoro-1*H*-indazol-1-yl)-3-(2,4-di-tert-butylphenoxy)-2-methylpropan-1-one ((*S*)-SX244, 1d)**

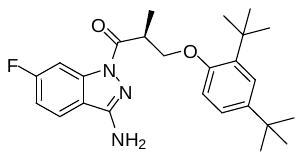

To a solution of crude **12d** (13.6 mg, 46.5 µmol) in DMF (1.5 mL) were added HOAt (6.5 mg, 47.8 µmol), followed by EDC × HCl (9.3 mg, 48.5 µmol) at rt. The resulting solution was stirred for 10 min at rt before adding 6‑fluoro-1*H*-indazole-3-amine (**13**, 5.9 mg, 38.8 µmol) dissolved in anhydrous DMF (0.5 mL). After stirring the reaction mixture at rt for 2 h, the solvent was removed by lyophilization. Purification by preparative HPLC using a solvent system of CH_3_OH / 0.1% aq. HCOOH, a flow rate of 10 mL / min and a gradient of 50% to 80% CH_3_OH in 15 min, 80% to 95% CH_3_OH in 2 min, 95% CH_3_OH for 6 min (*t*_R_ = 20.5 min, λ = 254 nm) afforded **(*S*)-SX244** (**1d**, 2.4 mg, 5.64 μmol, 15%) as a white solid.

ESI-MS:	*m/z* 426.5 [M+H]^+^.

HR-ESI-MS:	*m/z* [M+H]^+^ calcd. 426.2551 for C_25_H_33_FN_3_O_2_, found 426.2558.

IR (NaCl):	3463, 3350, 2960, 2864, 1676, 1624, 1423, 1231 cm^-1^.

m.p.:	53 °C.

^1^H-NMR:	(600 MHz, DMSO-d_6_) *δ* 1.13 (s, 9H), 1.23 (s, 9H), 1.33 (d, *J* = 6.7 Hz, 3H), 4.17 (dd, *J* = 8.0, 5.0 Hz, 1H), 4.23 – 4.17 (m, 1H), 4.27 (dd, *J* = 8.0, 7.7 Hz, 1H), 6.57 (s, 2H), 6.89 (d, *J* = 8.5 Hz, 1H), 7.14 (dd, *J* = 8.5, 2.5 Hz, 1H), 7.16 (d, *J* = 2.5 Hz, 1H), 7.24 (ddd, *J* = 9.0, 8.9, 2.3 Hz, 1H), 7.91 – 7.96 (m, 2H).

^13^C-NMR:	(DEPTQ, 151 MHz, DMSO-d_6_) *δ* 14.0, 29.5, 31.4, 33.9, 34.3, 37.6, 69.4, 101.7 (d, *J* = 28 Hz), 111.3, 112.1 (d, *J* = 25 Hz), 117.1, 122.6 (d, *J* = 11 Hz), 122.9, 123.4, 136.0, 139.9 (d, *J* = 13 Hz), 141.8, 152.5, 154.6, 163.1 (d, *J* = 244 Hz), 172.9.

^19^F-NMR:	(377 MHz, DMSO-d_6_) *δ* -110.8.

HPLC:	system 2: λ = 254 nm, *t*_R_ = 22.0 min, purity: >99%.

chiral HPLC:	isocratic elution with *n*-hexane / isopropanol, 99:1: *t*_R_ = 11.4 min, >99% *ee* (254 nm).

[*α*]*_D_^23^*:	-5.7 (*c* = 0.1, methanol).

**(*R*)-1-(3-Amino-6-fluoro-1*H*-indazol-1-yl)-3-(2,4-di-tert-butylphenoxy)-2-methylpropan-1-one ((*R*)-SX244, ent-1d)**

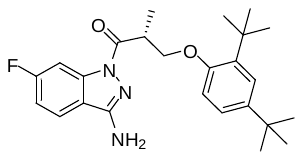

(*R*)-1-(3-Amino-6-fluoro-1H-indazol-1-yl)-3-(2,4-di-tert-butylphenoxy)-2-methylpropan-1-one **((*R*)-SX244, ent-1d)** was synthesized analogously as described for **(*S*)-SX244 (1d)**, starting from crude **ent-12d**. The analytical data were in accordance.

HR-ESI-MS:	*m/z* [M+H]^+^ calcd. 426.2551 for C_25_H_33_FN_3_O_2_, found 426.2552.

HPLC:	system 2: λ = 254 nm, *t*_R_ = 22.0 min, purity: >99%.

chiral HPLC:	isocratic elution with *n*-hexane / isopropanol, 99:1: *t*_R_ = 10.5 min, 98% *ee* (254 nm).

[*α*]*_D_^26^*:	+7.1 (*c* = 0.1, methanol).

**(*S*)-1-(3-Amino-6-fluoro-1*H*-indazol-1-yl)-3-(4-fluorophenoxy)-2-methylpropan-1-one ((*S*)‑SX263, 1e)**

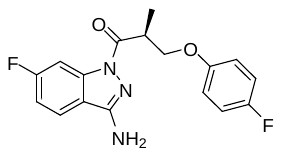

To a solution of **12e** (20.7 mg, 0.10 mmol) in DMF (2.5 mL) were added HOAt (13.6 mg, 0.10 mmol) and EDC × HCl (19.2 mg, 0.10 mmol) at rt. The resulting solution was stirred for 10 min at rt and then 6‑fluoro-1*H*-indazol-3-amine (**13**, 19.6 mg, 0.13 mmol) in DMF (0.5 mL) was added. After stirring the reaction mixture at rt for 1.5 h, the solvent was removed by lyophilization. Purification by preparative HPLC using a solvent system of CH_3_OH / 0.1% aq. HCOOH, a flow rate of 10 mL / min and a gradient of 50% to 80% CH_3_OH in 15 min, 80% to 95% CH_3_OH in 2 min, 95% CH_3_OH for 2 min (*t*_R_ = 16.0 min, λ = 254 nm) afforded **(*S*)-SX263** (**1e**, 16.9 mg, 51.0 μmol, 51%) as a white solid.

ESI-MS:	*m/z* 332.2 [M+H]^+^.

HR-ESI-MS:	*m/z* [M+H]^+^ calcd. 332.1205 for C_17_H_16_F_2_N_3_O_2_, found 332.1203.

IR (NaCl):	3463, 3357, 2925, 2849, 1683, 1625, 1509, 1419 cm^-1^.

m.p.:	148 °C.

^1^H-NMR:	(600 MHz, DMSO-d_6_) *δ* 1.28 (d, *J* = 6.9 Hz, 3H), 4.02 – 4.11 (m, 2H), 4.33 – 4.40 (m, 1H), 6.62 (s, 2H), 6.93 – 6.99 (m, 2H), 7.06 – 7.13 (m, 2H), 7.26 (ddd, *J* = 9.0, 8.9, 2.3 Hz, 1H), 7.92 – 7.99 (m, 2H).

^13^C-NMR:	(DEPTQ, 151 MHz, DMSO-d_6_) *δ* 13.9, 37.7, 69.5, 101.8 (d, *J* = 28 Hz), 112.2 (d, *J* = 25 Hz), 115.7 (d, *J* = 7 Hz), 115.8 (d, *J* = 8 Hz), 117.0, 122.7 (d, *J* = 11 Hz), 139.8 (d, *J* = 13 Hz), 152.7, 155.1 (d, *J* = 2 Hz), 156.5 (d, *J* = 236 Hz), 163.2 (d, *J* = 245 Hz), 172.4.

^19^F-NMR:	(377 MHz, DMSO-d_6_) *δ* -110.5, -123.3.

HPLC:	system 1: λ = 254 nm, *t*_R_ = 20.1 min, purity: 97%.

chiral HPLC:	isocratic elution with *n*-hexane / isopropanol, 95:5: *t*_R_ = 11.4 min, 99% *ee* (254 nm).

[*α*]*_D_^23^*:	-2.0 (*c* = 0.9, methanol).

**(*S*)-1-(3-Amino-6-fluoro-1*H*-indazol-1-yl)-3-(4-bromophenoxy)-2-methylpropan-1-one ((*S*)‑SX264, 1f)**

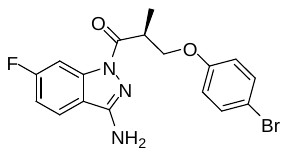

To a solution of **12f** (19.2 mg, 74.1 μmol) in DMF (2.5 mL) were added HOAt (10.1 mg, 74.1 μmol and EDC × HCl (14.2 mg, 74.1 μmol) at rt. The resulting solution was stirred for 10 min at rt and then 6‑fluoro-1*H*-indazol-3-amine (**13**, 13.4 mg, 88.9 μmol) in DMF (0.5 mL) was added. After stirring the reaction mixture at rt for 1.5 h, the solvent was removed by lyophilization. Purification by preparative HPLC using a solvent system of CH_3_OH / 0.1% aq. HCOOH, a flow rate of 10 mL / min and a gradient of 50% to 80% CH_3_OH in 15 min, 80% to 95% CH_3_OH in 2 min, 95% CH_3_OH for 4 min (*t*_R_ = 18.0 min, λ = 254 nm) afforded **(*S*)-SX264** (**1f**, 8.22 mg, 21.0 μmol, 28%) as a white solid.

ESI-MS:	*m/z* 392.3 [M+H]^+^.

HR-ESI-MS:	*m/z* [M+H]^+^ calcd. 392.0404 for C_17_H_16_BrFN_3_O_2_, found 392.0402

IR (NaCl):	3466, 3346, 2922, 1680, 1625, 1560, 1423, 1238, 1074 cm^-1^.

m.p.:	112 °C.

^1^H-NMR:	(600 MHz, DMSO-d_6_) *δ* 1.28 (d, *J* = 6.6 Hz, 3H), 4.03 – 4.10 (m, 2H), 4.34 – 4.43 (m, 1H), 6.62 (s, 2H), 6.89 – 6.97 (m, 2H), 7.26 (ddd, *J* = 9.0, 8.9, 2.3 Hz, 1H), 7.40 – 7.46 (m, 2H), 7.92 – 7.97 (m, 2H).

^13^C-NMR:	(DEPTQ, 151 MHz, DMSO-d_6_) *δ* 13.8, 37.6, 69.2, 101.8 (d, *J* = 28 Hz), 112.1, 112.3 (d, *J* = 25 Hz), 116.8, 117.0, 122.7 (d, *J* = 11 Hz), 132.1, 139.8 (d, *J* = 13 Hz), 152.7, 157.6, 163.2 (d, *J* = 245 Hz), 172.3.

^19^F-NMR:	(377 MHz, DMSO-d_6_) *δ* -110.4.

HPLC:	system 1: λ = 254 nm, *t*_R_ = 20.8 min, purity: >99%.

chiral HPLC:	isocratic elution with *n*-hexane / isopropanol, 95:5: *t*_R_ = 12.0 min, 98% *ee* (254 nm).

[*α*]*_D_^26^*:	-6.1 (*c* = 0.3, methanol).

**(*S*)-3-([1,1'-Biphenyl]-4-oxy)-1-(3-amino-6-fluoro-1*H*-indazol-1-yl)-2-methylpropan-1-one ((*S*)-SX267, 1g)**

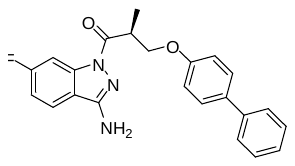

To a solution of compound **12g** (21.5 mg, 83.9 μmol) in DMF (2.5 mL) were added HOAt (11.4 mg, 83.9 μmol) and EDC × HCl (16.1 mg, 83.9 μmol) at rt. The resulting solution was stirred for 10 min at rt and then 6‑fluoro-1*H*-indazol-3-amine (**13**, 15.2 mg, 101 μmol) in DMF (0.5 mL) was added. After stirring the reaction mixture at rt for 3 h, the solvent was removed by lyophilization. Purification by preparative HPLC using a solvent system of CH_3_OH / 0.1% aq. HCOOH, a flow rate of 10 mL / min and a gradient of 50% to 80% CH_3_OH in 15 min, 80% to 95% CH_3_OH in 2 min, 95% CH_3_OH for 4 min (*t*_R_ = 19.0 min, λ = 254 nm) afforded **(*S*)-SX267 (1g**, 17.3 mg, 44.3 μmol, 53%) as a white solid.

ESI-MS:	*m/z* 390.3 [M+H]^+^.

HR-ESI-MS:	*m/z* [M+H]^+^ calcd. 390.1612 for C_23_H_21_FN_3_O_2_, found 390.1611.

IR (NaCl):	3456, 3343, 2925m 1686, 1625, 1440, 1422, 1320, 762 cm^-1^.

m.p.:	101 °C.

^1^H-NMR:	(600 MHz, DMSO-d_6_) *δ* 1.31 (d, *J* = 6.7 Hz, 3H), 4.08 – 4.15 (m, 2H), 4.42 – 4.48 (m, 1H), 6.63 (s, 2H), 7.00 – 7.08 (m, 2H), 7.26 (ddd, *J* = 9.0, 8.9, 2.4 Hz, 1H), 7.28 – 7.32 (m, 1H), 7.40 – 7.45 (m, 2H), 7.52 – 7.64 (m, 4H), 7.92 – 8.01 (m, 2H).

^13^C-NMR:	(DEPTQ, 151 MHz, DMSO-d_6_) *δ* 13.9, 37.7, 69.0, 101.8 (d, *J* = 28 Hz), 112.2 (d, *J* = 25 Hz), 115.0, 117.0, 122.7 (d, *J* = 11 Hz), 126.1, 126.7, 127.7, 128.8, 132.7, 139.8, 139.8 (d, *J* = 13 Hz), 152.7, 157.9, 163.2 (d, *J* = 245 Hz), 172.4.

^19^F-NMR:	(377 MHz, DMSO-d_6_) *δ* -110.4.

HPLC:	system 2: λ = 254 nm, *t*_R_ = 19.6 min, purity: >99%.

chiral HPLC:	isocratic elution with *n*-hexane / isopropanol, 9:1: *t*_R_ = 11.5 min, 99% *ee* (254 nm).

[*α*]*_D_^25^*:	-11.0 (*c* = 1.1, methanol).

**(*S*)-1-(3-Amino-6-fluoro-1*H*-indazol-1-yl)-3-(benzofuran-5-yloxy)-2-methylpropan-1-one ((*S*)-SX268, 1h)**

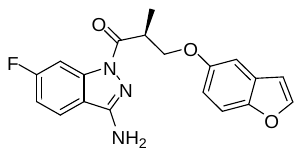

To a solution of **12h** (11.5 mg, 52.2 μmol) in DMF (2.5 mL) were added HOAt (7.1 mg, 52.2 μmol) and EDC × HCl (10.0 mg, 52.2 μmol) at rt. The resulting solution was stirred for 10 min at rt and then 6‑fluoro-1*H*-indazol-3-amine (**13**, 9.5 mg, 62.7 μmol) in DMF (0.5 mL) was added. After stirring the reaction mixture at rt for 2 h, the solvent was removed by lyophilization. Purification by preparative HPLC using a solvent system of CH_3_OH / 0.1% aq. HCOOH, a flow rate of 10 mL / min and a gradient of 50% to 80% CH_3_OH in 15 min, 80% to 95% CH_3_OH in 2 min, 95% CH_3_OH for 4 min (*t*_R_ = 16.0 min, λ = 254 nm) afforded **(*S*)-SX268** (**1h**, 3.55 mg, 10.0 μmol, 19%) as a white solid.

ESI-MS:	*m/z* 354.2 [M+H]^+^.

HR-ESI-MS:	*m/z* [M+H]^+^ calcd. 354.1248 for C_19_H_17_FN_3_O_3_, found 354.1247.

IR (NaCl):	3446, 3350, 2922, 1686, 1628, 1436, 1197 cm^-1^.

m.p.:	53 °C.

^1^H-NMR:	(600 MHz, DMSO-d_6_) *δ* 1.30 (d, *J* = 6.6 Hz, 3H), 4.07 – 4.13 (m, 2H), 4.36 – 4.45 (m, 1H), 6.60 (s, 2H), 6.86 (dd, *J* = 2.2, 0.9 Hz, 1H), 6.87 (dd, *J* = 7.5, 2.6 Hz, 1H), 7.20 (d, *J* = 2.6 Hz, 1H), 7.25 (ddd, *J* = 9.0, 8.9, 2.4 Hz, 1H), 7.45 (dd, *J* = 7.5, 0.9 Hz, 1H), 7.93 (d, *J* = 2.2 Hz, 1H), 7.93 – 7.98 (m, 2H).

^13^C-NMR:	(DEPTQ, 151 MHz, DMSO-d_6_) *δ* 14.0, 37.8, 69.7, 101.8 (d, *J* = 28 Hz), 104.8, 106.9, 111.7, 112.2 (d, *J* = 25 Hz), 113.4, 117.0, 122.7 (d, *J* = 11 Hz), 127.8, 139.8 (d, *J* = 14 Hz), 146.7, 149.3, 152.6, 154.6, 163.2 (d, *J* = 245 Hz), 172.5.

HPLC:	system 1: λ = 254 nm*, t*_R_ = 20.1 min, purity: 97%.

chiral HPLC:	isocratic elution with *n*-hexane / isopropanol, 95:5: *t*_R_ = 18.4 min, 99% *ee* (254 nm).

[*α*]*_D_^25^*:	-9.5 (*c* = 0.1, methanol).

**1-(3-Amino-6-methoxy-1*H*-indazol-1-yl)-2-methylpropan-1-one (I1381, 2)**

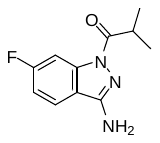

To a solution of HOAt (54.0 mg, 0.40 mmol) in DMF (1 mL) was added DIPEA (96.5 mg, 130 μL, 0.74 mmol) and then isobutyryl chloride (40.7 mg, 40 μL, 0.38 mmol) at 0 °C. After 1 h of stirring, a solution of 6-fluoro-1*H*-indazol-3-amine (**13**, 50.0 mg, 0.33 mmol) in DMF (0.5 mL) at rt was added. Microwave irradiation (described in the general part) in a sealed tube was performed, then the solvent was removed by lyophilization and the residue was purified by preparative HPLC using a solvent system of CH_3_CN / 0.1% aq. TFA, a flow rate of 10 mL / min and a gradient of 40 to 60% in 15 min (*t*_R_: 12.6 min, λ = 220 nm) to afford **I1381** (**2**, 65.6 mg, 90%) as a white solid.

ESI-MS:	*m/z* 222.0 [M+H]^+^.

HR-ESI-MS:	*m/z* [M+H]^+^ calcd. 222.1037 for C_11_H_13_FN_3_O, found 222.1038.

IR (KBr):	3460, 3382, 3350, 3211, 1672 cm^-1^.

m.p.:	103 °C.

^1^H-NMR:	(400 MHz, DMSO-d_6_) *δ* 1.19 (d, *J* = 6.9 Hz, 6H), 3.65 (sept, *J* = 6.9 Hz, 1H), 6.56 (s, 2H), 7.23 (ddd, *J* = 9.3, 8.8, 2.3 Hz, 1H), 7.92 (dd, *J* = 8.8, 5.4 Hz, 1H), 7.94 (dd, *J* = 10.1, 2.3 Hz, 1H).

^13^C-NMR:	(DEPTQ, 151 MHz, DMSO-d_6_) *δ* 18.8, 31.8, 101.7 (d, *J* = 28 Hz), 111.9 (d, *J* = 24 Hz), 116.8, 122.5 (d, *J* = 11 Hz), 139.9 (d, *J* = 10 Hz), 152.4, 163.2 (d, *J* = 245 Hz), 175.5.

HPLC:	system 3: λ = 220 nm*, t*_R_ = 21.2 min, purity: >99%.

**(*R*,*S*)-1-(6-Fluoro-1*H*-indazol-1-yl)-2-methyl-3-phenoxypropan-1-one (I1384, 3a)**

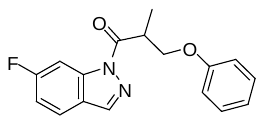

To a solution of commercially available (*R*,*S*)-2-methyl-3-phenoxypropionic acid (**rac-12a**, 79.4 mg, 0.44 mmol) and HOAt (60.0 mg, 0.44 mmol) in DMF (1 mL) was added EDC × HCl (84.5 mg, 0.44 mmol). After stirring for 10 min at rt, a solution of commercially available 6-fluoro-1*H*-indazole (**14**, 50.0 mg, 0.37 mmol) in DMF (0.5 mL) was added. After stirring the reaction mixture for 4 h, the solvent was removed by lyophilization and the residue was purified by preparative HPLC using a solvent system of CH_3_OH / 0.1% aq. HCOOH, a flow rate of 10 mL / min and a gradient of 20% to 90% CH_3_OH in 25 min, 90% to 95% CH_3_OH in 2 min, 95% CH_3_OH for 2 min (*t*_R_: 27.5 min, λ = 220 nm) to afford **I1384** (**3a**, 43.0 mg, 0.14 mmol, 39%) as a yellow-brownish resin.

ESI-MS:	*m/z* 299.1 [M+H]^+^.

HR-ESI-MS:	*m/z* [M+H]^+^ calcd. 299.1190 for C_17_H_16_FN_2_O_2_, found 299.1191.

IR (KBr):	3424, 1715 cm^-1^.

^1^H-NMR:	(600 MHz, DMSO-d_6_) *δ* 1.36 (d, *J* = 7.1 Hz, 3H), 4.19 (dd, *J* = 9.6, 5.1 Hz, 1H), 4.31 (ddq, *J* = 7.4, 5.1, 7.1 Hz, 1H), 4.41 (dd, *J* = 9.6, 7.4 Hz, 1H), 6.90 – 6.95 (m, 3H), 7.24 – 7.29 (m, 2H), 7.36 (ddd, *J* = 9.2, 8.8, 2.1 Hz, 1H), 8.00 (dd, *J* = 8.8, 5.3 Hz, 1H), 8.06 (dd, *J* = 9.2, 2.1 Hz, 1H), 8.55 (brs, 1H).

^13^C-NMR:	(DEPTQ, 151 MHz, DMSO-d_6_) *δ* 14.0, 38.2, 68.9, 101.3 (d, *J* = 28 Hz), 113.8 (d, *J* = 24 Hz), 114.5, 120.8, 123.0, 123.8 (d, *J* = 11 Hz), 129.5, 139.9 (d, *J* = 13 Hz), 140.8, 158.2, 163.0 (d, *J* = 245 Hz), 174.3.

HPLC:	system 3: λ = 220 nm*, t*_R_ = 27.5 min, purity: 96%.

**(*S*)-1-(6-Fluoro-3-methyl-1*H*-indazol-1-yl)-2-methyl-3-phenoxypropan-1-one (I1402, 3b)**

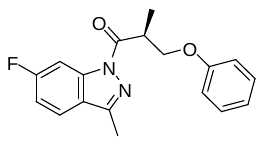

To a solution of **12a** (15.0 mg, 0.08 mmol) and HOAt (11.3 mg, 0.08 mmol) in DMF (0.5 mL) was added EDC × HCl (16.0 mg, 0.08 mmol). After stirring for 10 min at rt, a solution of commercially available 6-fluoro-3-methyl-1*H*-indazole (**15**, 15.0 mg, 0.10 mmol) in DMF (0.5 mL) was added. After stirring the reaction mixture for 4 h, the solvent was removed by lyophilization and the residue was purified by preparative HPLC using a solvent system of CH_3_OH / 0.1% aq. HCOOH, a flow rate of 10 mL / min and a gradient of 20% to 90% CH_3_OH in 22 min, 90% to 95% CH_3_OH in 2 min, 95% CH_3_OH for 2 min (*t*_R_: 24.0 min, λ = 220 nm) to afford **I1402** (**3b**, 14.9 mg, 47.7 µmol, 57%) as a white solid.

ESI-MS:	*m/z* 313.1 [M+H]^+^.

HR-ESI-MS:	*m/z* [M+H]^+^ calcd. 313.1347 for C_18_H_18_FN_2_O_2_, found 313.1348.

IR (NaCl):	3447, 1707 cm^-1^.

m.p.:	51 °C.

^1^H-NMR:	(600 MHz, DMSO-d_6_) *δ* 1.34 (d, *J* = 7.1 Hz, 3H), 2.57 (s, 3H), 4.15 (dd, *J* = 9.4, 5.4 Hz, 1H), 4.27 (ddq, *J* = 8.0, 5.4, 7.1 Hz, 1H), 4.39 (dd, *J* = 9.4, 8.0 Hz, 1H), 6.91 – 6.94 (m, 3H), 7.24 – 7.29 (m, 2H), 7.35 (ddd, *J* = 9.2, 8.8, 2.2 Hz, 1H), 7.95 (ddd, *J* = 8.8, 5.1, 0.3 Hz, 1H), 8.02 (ddd, *J* = 9.5, 2.2, 0.3 Hz, 1H).

^13^C-NMR:	(DEPTQ, 151 MHz, DMSO-d_6_) *δ* 11.8, 13.9, 37.9, 68.8, 101.3 (d, *J* = 29 Hz), 113.3 (d, *J* = 25 Hz), 114.4, 120.7, 122.0, 123.0 (d, *J* = 11 Hz), 129.4, 139.4 (d, *J* = 17 Hz), 149.1, 158.1, 163.0 (d, *J* = 256 Hz), 173.8.

HPLC:	system 3: λ = 220 nm*, t*_R_ = 28.7 min, purity: 98%.

chiral HPLC:	isocratic elution with *n*-hexane / isopropanol, 99:1: *t*_R_ = 9.6 min, 99% *ee* (254 nm).

[*α*]*_D_^20^*:	+33.3 (*c* = 0.5, chloroform).

***N*-(1-Acetyl-6-fluoro-1*H*-indazol-3-yl)acetamide (16)**

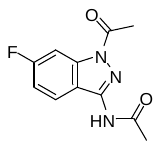

To a solution of 6-fluoro-1*H*-indazol-3-amine (**13**, 100.0 mg, 0.66 mmol) and DMAP in pyridine (1.5 mL) was slowly added acetyl chloride (207.7 mg, 189 μL, 2.65 mmol) at 0 °C. After 5 min of stirring at rt, dioxane (3 mL) was added and stirring was continued for 1 h. The solvent was removed *in vacuo* and flash chromatography was performed (isohexane / ethyl acetate, 9:1 🡪 8:2) to afford **16** (11.4 mg, 72%) as a white solid.

ESI-MS:	*m/z* 258.1 [M+Na]^+^.

HR-ESI-MS:	*m/z* [M+H]^+^ calcd. 236.0830 for C_11_H_11_FN_3_O_2_, found 236.0828.

IR (KBr):	3409, 3273, 3232, 1711, 1670 cm^-1^.

m.p.:	193-194 °C.

^1^H-NMR:	(600 MHz, DMSO-d_6_) *δ* 2.19 (s, 3H), 2.64 (s, 3H), 7.29 (ddd, *J* = 9.0, 9.0, 2.3 Hz, 1H), 8.00 (dd, *J* = 9.7, 2.3 Hz, 1H), 8.08 (dd, *J* = 9.0, 5.5 Hz, 1H), 10.93 (s, 1H).

^13^C-NMR:	(DEPTQ, 151 MHz, DMSO-d_6_) *δ* 22.7, 23.8, 101.0 (d, *J* = 29 Hz), 112.8 (d, *J* = 25 Hz), 117.0, 125.6 (brs), 139.7 (d, *J* = 14 Hz), 145.1, 163.0 (d, *J* = 246 Hz), 169.1 (brs), 170.2.

***N*-(1-Acetyl-6-fluoro-1H-indazol-3-yl)-*N*-methylacetamide (17)**

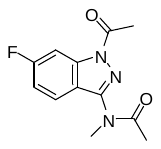

To a solution of *N*-(1-acetyl-6-fluoro-1*H*-indazol-3-yl)acetamide (**16**, 68.1 mg, 0.29 mmol) in DMF (1 mL) was added sodium hydride (8.3 mg, 0.35 mmol) and after 10 min methyl iodide (164.4 mg, 72 μL, 1.16 mmol) at rt. After 1 h of stirring, the solvent was removed *in vacuo* and flash chromatography was performed (isohexane / ethyl acetate, 9:1 🡪 8:2) to afford **17** (55.7 mg, 77%) as a white solid.

ESI-MS:	*m/z* 250.1 [M+H]^+^.

HR-ESI-MS:	*m/z* [M+Na]^+^ calcd. 272.0806 for C_12_H_12_FN_3_NaO_2_, found 272.0802.

IR (KBr):	1728, 1690, 1679 cm^-1^.

m.p.:	98-100 °C.

^1^H-NMR:	(600 MHz, DMSO-d_6_) *δ* 2.10 (brs, 3H), 2.68 (s, 3H), 3.37 (brs, 3H), 7.36 (brdd, *J* = 8.5, 8.5 Hz, 1H), 7.90 (brs, 1H), 8.05 (brd, *J* = 9.7, 1H).

^13^C-NMR:	(DEPTQ, 151 MHz, DMSO-d_6_) *δ* 22.4, 22.6, 35.2 (detected by HSQC), 101.4 (d, *J* = 28 Hz), 113.8 (brs), 117.9, 124.3 (d, *J* = 12 Hz), 139.8 (d, *J* = 18 Hz), 149.0, 163.2 (d, *J* = 246 Hz), 170.6.

**6-Fluoro-*N*-methyl-1*H*-indazol-3-amine (18)**

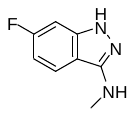

A solution of *N*-(1-acetyl-6-fluoro-1*H*-indazol-3-yl)-*N*-methylacetamide (**17**, 30.0 mg, 0.12 mmol) in 1.25 M HCl in methanol (2 mL) was heated at 115 °C in a sealed tube. After 2 h of stirring, the solvent was removed *in vacuo* to give crude **18** (21.9 mg, crude) as a brownish oil, which was used for the next reaction without further purification. For analytical purposes, a small sample of the substance was purified by preparative HPLC using a solvent system of CH_3_OH / 0.1% aq. TFA, a flow rate of 12 mL/min and a gradient of 10 to 50% in 7 min (*t*_R_: 8.6 min, λ = 220 nm) to furnish a yellow oil.

ESI-MS:	*m/z* 166.1 [M+H]^+^.

HR-ESI-MS:	*m/z* [M+H]^+^ calcd. 166.0775 for C_8_H_9_FN_3_, found 166.0775.

IR (KBr):	3418, 3215, 1651 cm^-1^.

^1^H-NMR:	(600 MHz, DMSO-d_6_) *δ* 2.89 (s, 3H), 6.83-6.89 (m, 1H), 7.06-7.10 (m, 1H), 7.76-7.81 (m, 1H) 11.80 (brs, 1H).

^13^C-NMR:	(DEPTQ, 151 MHz, DMSO-d_6_) *δ* 29.8, 95.4 (d, *J* = 27 Hz), 117.8 (d, *J* = 27 Hz), 110.0, 122.6 (d, *J* = 10 Hz), 142.5 (d, *J* = 13 Hz), 150.2, 162.9 (d, *J* = 242 Hz).

**(*S*)-1-(6-fluoro-3-(methylamino)-1*H*-indazol-1-yl)-2-methyl-3-phenoxypropan-1-one (I1436, 3c)**

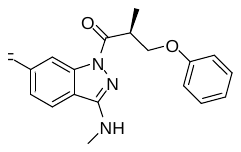

To a solution of **12a** (10.0 mg, 0.055 mmol) in DMF (0.5 mL) was added HOAt (7.6 mg, 0.055 mmol) and EDC × HCl (10.6 mg, 0.055 mmol) at rt. The solution was stirred for 10 min at rt and then crude 6-fluoro-*N*-methyl-1*H*-indazol-3-amine (**18**, 11.0 mg, 0.12 mmol) in DMF (0.5 mL) was added. After stirring the reaction mixture at rt for 14 h, the solvent was removed by lyophilization. Purification by preparative HPLC using a solvent system of CH_3_OH / 0.1% aq. TFA, a flow rate of 12 mL / min and a gradient of 50% to 80% CH_3_OH in 15 min, 80% to 95% CH_3_OH in 2 min (*t*_R_ = 16.0 min, λ = 220 nm) afforded **I1436** (**3c**, 5.2 mg, 57.9 μmol, 29%) as a white solid.

ESI-MS:	*m/z* 328.5 [M+H]^+^.

HR-ESI-MS:	*m/z* [M+H]^+^ calcd. 328.1456 for C_18_H_19_FN_3_O_2_, found 328.1455.

IR (NaCl):	3376, 1682, cm^-1^.

m.p.:	80 °C.

^1^H-NMR:	(600 MHz, DMSO-d_6_) *δ* 1.30 (d, *J* = 7.0 Hz, 3H), 2.93 (d, *J* = 4.9 Hz, 3H), 4.09 (dd, *J* = 9.4, 5.8 Hz, 1H), 4.16 (ddq, *J* = 7.7, 5.8, 7.0 Hz, 1H), 4.42 (dd, *J* = 9.4, 7.7 Hz, 1H), 6.91 – 6.95 (m, 1H), 7.18 (q, *J* = 4.9 Hz, 1H), 7.25 (ddd, *J* = 8.9, 8.5, 2.3 Hz, 1H), 7.25 – 7.30 (m, 2H), 7.91 (dd, *J* = 8.5, 5.4 Hz, 1H), 7.97 (dd, *J* = 9.8, 2.3 Hz, 1H).

^13^C-NMR:	(DEPTQ, 151 MHz, DMSO-d_6_) *δ* 13.6, 28.8, 37.6, 68.7, 101.8 (d, *J* = 28 Hz), 112.3 (d, *J* = 24 Hz), 114.5, 116.6, 120.6, 122.1 (d, *J* = 11 Hz), 129.4, 140.0 (d, *J* = 13 Hz), 153.0, 158.2, 163.1 (d, *J* = 244 Hz), 172.4.

HPLC:	system 3: λ = 220 nm,	*t*_R_ = 26.3 min, purity: >99%.

[*α*]*_D_^23^*:	-5.7 (*c* = 0.17, methanol).

**6-Fluoro-1*H*-pyrazolo[4,3-*b*]pyridin-3-amine (20)** ^5^

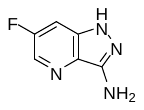

A solution of 3,5-difluoro-2-pyridinecarbonitrile (**19** 70 mg, 0.50 mmol) and hydrazine hydrate (37.6 mg, 36 µL, 0.75 mmol) in ethanol (1.3 mL) was heated at 70 °C in a sealed tube for 17 h. After cooling to rt, the solvent was removed *in vacuo*, and the residue was purified by preparative HPLC using a solvent system of CH_3_CN / 0.1% aq. TFA, a flow rate of 10 mL / min and a gradient of 3% to 18% CH_3_CN in 12 min (*t*_R_: 9.6 min, λ = 220 nm) to afford **20** (11.4 mg, 41.4 µmol, 15%) as a white solid.

ESI-MS:	*m/z* 153.0 [M+H]^+^.

IR (KBr):	3444, 3315, 3210 cm^-1^.

^1^H-NMR:	(600 MHz, DMSO-d_6_) *δ* 5.44 (s, 2H), 7.57 (dd, *J* = 9.6, 2.6 Hz, 1H), 8.26 (d, *J* = 2.6, 1.2 Hz, 1H), 11.73 (brs, 1H).

^13^C-NMR:	(DEPTQ, 151 MHz, DMSO-d_6_) *δ* 102.7 (d, *J* = 22 Hz), 131.3 (d, *J* = 29 Hz), 133.5 (d, *J* = 9 Hz), 148.9, 157.9 (d, *J* = 249 Hz).

The analytical data of this compound are in accordance with those reported in the literature.

**(*S*)-1-(3-amino-6-fluoro-1H-pyrazolo[4,3-*b*]pyridin-1-yl)-2-methyl-3-phenoxypropan-1-one (BBG3, 4)**

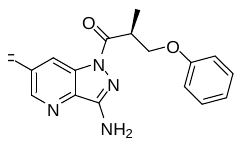

To a solution of **12a** (13.5 mg, 0.075 mmol) and HOAt (10.2 mg, 0.075 mmol) in DMF (0.5 mL) was added EDC × HCl (14.4 mg, 0.075 mmol). After 10 min at rt, a solution of **20** (11.4 mg, 0.09 mmol) in DMF (0.5 mL) was added. After stirring for 3 h, the solvent was removed by lyophilization and the residue was purified by preparative HPLC using a solvent system of CH_3_CN / 0.1% aq. TFA, a flow rate of 10 mL / min and a gradient of, 5% to 76% CH_3_CN in 23 min, 76% to 95% in 2 min (*t*_R_: 23.4 min, λ = 220 nm) to afford **BBG3** (**4**, 7.2 mg, 22.9 µmol, 31%) as a white solid.

ESI-MS:	*m/z* 315.1 [M+H]^+^.

HR-ESI-MS:	*m/z* [M+H]^+^ calcd. 315.1252 for C_16_H_16_FN_4_O_2_, found 315.1254

IR (NaCl):	3450, 3344, 1694, 1630 cm^-1^.

m.p.:	101 °C.

^1^H-NMR:	(600 MHz, DMSO-d_6_) *δ* 1.29 (m, 3H), 4.06 – 4.13 (m, 2H), 4.34 – 4.42 (m, 1H), 6.87 (s, 2H), 6.90 – 6.95 (m, 3H), 7.24 – 7.29 (m, 2H), 8.33 (dd, *J* = 9.0, 2.5 Hz, 1H), 8.67 (dd, *J* = 2.5, 0.9 Hz, 1H).

^13^C-NMR:	(DEPTQ, 151 MHz, DMSO-d_6_) *δ* 13.7, 37.3, 68.6, 109.4 (d, *J* = 25 Hz), 114.4, 120.6, 129.4, 132.7 (d, *J* = 12 Hz), 134.2, 136.1 (d, *J* = 29 Hz), 152.2, 159.2 (d, *J* = 254 Hz), 172.9.

HPLC:	system 3: λ = 220 nm,	*t*_R_ = 23.6 min, purity: >99%.

[*α*]*_D_^23^*:	-3.6 (*c* = 0.3, methanol).

**(*R*,*S*)-1-(3-Amino-6-methoxy-1*H*-indazol-1-yl)-2-methyl-3-phenoxypropan-1-one (IK195, 5a)**

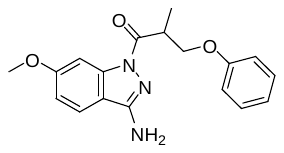

To a solution of commercially available (*R*,*S*)-2-methyl-3-phenoxypropionic acid (**rac-12**, 66.0 mg, 0.37 mmol), PyBOP (194 mg, 0.37 mmol) and HOAt (76 mg, 0.56 mmol) in anhydrous DMF (3 mL) was added DIPEA (64 µL, 0.37 mmol) and then a solution of commercially available 6-methoxy-1*H*-indazol-3-amine (**21**, 50.0 mg, 0.26 mmol) in DMF (0.5 mL) at rt. Microwave irradiation (described in the general part) in a sealed tube was performed, then the solvent was removed by lyophilization and the residue was purified by preparative HPLC using a solvent system of CH_3_CN / 0.1% aq. TFA, a flow rate of 20 mL / min and a gradient of 40% CH_3_CN for 1 min, 40% to 65% CH_3_CN in 10 min (*t*_R_: 10.2 min, λ = 254 nm) to afford **IK195** (**5a**, 56.3 mg, 0.17 mmol, 56%) as a white solid.

ESI-MS:	*m/z* 326.2 [M+H]^+^.

HR-ESI-MS:	*m/z* [M+H]^+^ calcd. 326.1499 for C_18_H_20_N_3_O_3_, found 326.1503.

IR (KBr):	3446, 3339, 3229, 1676 cm^-1^.

m.p.:	160-163 °C.

^1^H-NMR:	(400 MHz, DMSO-d_6_) *δ* 1.28 (d, *J* = 6.8 Hz, 3H), 3.83 (s, 3H), 4.00 – 4.14 (m, 2H), 4.34 – 4.43 (m, 1H), 6.46 (s, 2H), 6.89 – 6.95 (m, 3H), 6.69 (dd, *J* = 8.7, 2.2 Hz, 1H), 7.24 – 7.31 (m, 2H), 7.77 (d, *J* = 8.7 Hz, 1H), 7.79 (d, *J* = 2.2 Hz, 1H).

^13^C-NMR:	(DEPTQ, 151 MHz, DMSO-d_6_) *δ* 14.0, 37.7, 55.5, 68.9, 98.3, 113.2, 114.0, 114.4, 120.6, 121.5, 129.5, 140.9, 152.9, 158.3, 161.2, 172.3.

HPLC:	system 3: λ = 220 nm*, t*_R_ = 24.6 min, purity: >99%.

**(*R*,*S*)-1-(3-Amino-6-isopropoxy-1*H*-indazol-1-yl)-2-methyl-3-phenoxypropan-1-one (IK192, 5b)**

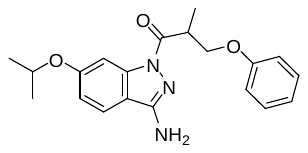

To a solution of commercially available (*R*,*S*)-2-methyl-3-phenoxypropionic acid (**rac-12**, 56.5 mg, 0.31 mmol), PyBOP (163 mg, 0.31 mmol) and HOAt (64 mg, 0.47 mmol) in anhydrous DMF (3 mL) was added DIPEA (54 µL, 0.31 mmol) and then a solution of commercially available 6-isopropoxy-1*H*-indazol-3-amine (**22**, 50.0 mg, 0.26 mmol) in DMF (0.5 mL) at ambient temperature. Microwave irradiation (described in the general part) in a sealed tube was performed, then the solvent was removed by lyophilization and the residue was purified by preparative HPLC using a solvent system of CH_3_CN / 0.1% aq. TFA, a flow rate of 20 mL / min and a gradient of 40% CH_3_CN for 1 min, 40% to 65% CH_3_CN in 10 min, 65% to 95% CH_3_CN in 2 min (*t*_R_: 12.9 min, λ = 250 nm) to afford **IK192** (**5b**, 50.0 mg, 0.14 mmol, 54%) as a white solid.

ESI-MS:	*m/z* 354.2 [M+H]^+^.

HR-ESI-MS:	*m/z* [M+H]^+^ calcd. 354.1812 for C_18_H_20_N_3_O_3_, found 354.1814.

IR (KBr):	3438, 3343, 3233, 1679 cm^-1^.

m.p.:	138 °C.

^1^H-NMR:	(400 MHz, DMSO-d_6_) *δ* 1.27 (d, *J* = 6.8 Hz, 3H), 1.30 (d, *J* = 6.0 Hz, 6H), 4.00 – 4.13 (m, 2H), 4.33 – 4.43 (m, 1H), 4.66 (sept, *J* = 6.0 Hz, 1H), 6.48 (brs, 2H), 6.90 – 6.96 (m, 3H), 6.92 (dd, *J* = 8.8, 2.2 Hz, 1H), 7.23 – 7.32 (m, 2H), 7.75 (d, *J* = 8.8 Hz, 1H), 7.77 (d, *J* = 2.2 Hz, 1H).

^13^C-NMR:	(DEPTQ, 101 MHz, DMSO-d_6_) *δ* 14.1, 21.7, 21.8, 37.8, 68.9, 70.0, 100.2, 113.9, 114.3, 114.5, 120.7, 121.7, 129.5, 140.9, 152.9, 158.3, 159.4, 172.4.

HPLC:	system 3: λ = 220 nm*, t*_R_ = 26.7 min, purity: 99%.

**(*S*)-1-(3-Amino-6-(trifluoromethyl)-1*H*-indazol-1-yl)-2-methyl-3-phenoxypropan-1-one ((*S*)-KD1, 5c)**

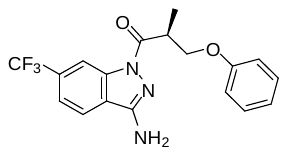

To a solution of **12a** (10.0 mg, 0.06 mmol) and HOAt (7.7 mg, 0.06 mmol) in anhydrous DMF (1 mL) was added EDC × HCl (11.1 mg, 0.06 mmol). After stirring for 10 min at ambient temperature, a solution of commercially available 6-(trifluoromethyl)-1*H*-indazol-3-amine (**23**, 16.7 mg, 0.10 mmol) in DMF (0.5 mL) was added. After stirring the reaction mixture for 2 h, the solvent was removed by lyophilization and the residue was purified by preparative HPLC using a solvent system of CH_3_OH / 0.1% aq. HCOOH, a flow rate of 10 mL / min and a gradient of 50% to 80% CH_3_OH in 15 min, 80% to 95% CH_3_OH in 2 min, 95% CH_3_OH for 4 min (*t*_R_: 20.6 min, λ = 254 nm) to afford **(*S*)-KD1** (**5c**, 4.0 mg, 11.0 µmol, 23%) as a white resin.

ESI-MS:	*m/z* 364.1 [M+H]^+^.

HR-ESI-MS:	*m/z* [M+H]^+^ calcd. 364.1267 for C_18_H_17_F_3_N_3_O_2_, found 364.1272.

IR (KBr):	3441, 3339, 3254, 3229, 1685 cm^-1^.

^1^H-NMR:	(600 MHz, DMSO-d_6_) *δ* 1.28 – 1.34 (m, 3H), 4.05 – 4.17 (m, 2H), 4.35 – 4.44 (m, 1H), 6.82 (s, 2H), 6.90 – 6.97 (m, 3H), 7.24 – 7.31 (m, 2H), 7.75 (dd, *J* = 8.4, 1.2 Hz, 1H), 8.17 (d, *J* = 8.4 Hz, 1H), 8.56 (brs, 1H).

^13^C-NMR:	(DEPTQ, 151 MHz, DMSO-d_6_) *δ* 13.8, 37.7, 68.7, 111.9 (q, *J* = 5 Hz), 114.4, 120.3 (q, *J* = 3 Hz), 120.6, 122.3, 122.7, 124.1 (q, *J* = 273 Hz), 129.3, 129.7 (q, *J* = 32 Hz), 138.4, 152.5, 158.2, 172.7.

HPLC:	system 3: λ = 220 nm*, t*_R_ = 27.0 min, purity: >99%.

chiral HPLC:	isocratic elution with *n*-hexane / isopropanol, 98:2: *t*_R_ = 15.3 min, 99% *ee* (254 nm).

[*α*]*_D_^25^*:	+2.2 (*c* = 0.3, methanol).

**(*S*)-1-(3-Amino-6-chloro-1*H*-indazol-1-yl)-2-methyl-3-phenoxypropan-1-one (I1414, 5d)**

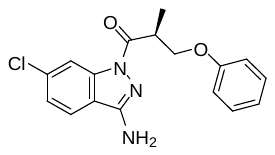

To a solution **12a** (15.0 mg, 0.08 mmol) and HOAt (11.3 mg, 0.08 mmol) in DMF (1 mL) was added EDC × HCl (15.9 mg, 0.08 mmol). After stirring for 10 min at rt, a solution of 6-chloro-1*H*-indazol-3-amine (**24**, 16.7 mg, 0.10 mmol) in DMF (0.5 mL) was added. After stirring the reaction mixture for 4 h, the solvent was removed by lyophilization and the residue was purified by preparative HPLC using a solvent system of CH_3_OH / 0.1% aq. HCOOH, a flow rate of 10 mL / min and a gradient of 50% to 80% CH_3_OH in 15 min, 80% to 95% CH_3_OH in 2 min, 95% CH_3_OH for 2 min, 95% to 50% CH_3_OH in 2 min (*t*_R_: 20.2 min, λ = 220 nm) to afford **I1414** (**5d**, 14.7 mg, 44.7 µmol, 54%) as a grey-white resin.

ESI-MS:	*m/z* 330.2 [M+H]^+^.

HR-ESI-MS:	*m/z* [M+H]^+^ calcd. 330.1004 for C_17_H_17_ClN_3_O_2_, found 330.1005.

IR (KBr):	3429, 3335, 3247, 3223, 1685 cm^-1^.

^1^H-NMR:	(600 MHz, DMSO-d_6_) *δ* 1.27 – 1.39 (m, 3H), 4.03 – 4.11 (m, 2H), 4.35 – 4.41 (m, 1H), 6.65 (s, 2H), 6.90 – 6.95 (m, 3H), 7.24 – 7.30 (m, 2H), 7.43 (dd, *J* = 8.4, 2.0 Hz, 1H), 7.93 (d, *J* = 8.4 Hz, 1H), 8.26 (d, *J* = 2.0 Hz, 1H).

^13^C-NMR:	(DEPTQ, 151 MHz, DMSO-d_6_) *δ* 13.9, 37.8, 68.8, 114.5, 114.7, 119.1, 120.7, 122.4, 124.2, 129.5, 139.7, 152.6, 158.3, 172.5.

HPLC:	system 3: λ = 220 nm*, t*_R_ = 25.8 min, purity: 96%.

chiral HPLC:	isocratic elution with *n*-hexane / isopropanol, 95:5: *t*_R_ = 13.0 min, 99% *ee* (254 nm).

[*α*]*_D_^20^*:	+45.3 (*c* = 0.5, chloroform).

**(*S*)-1-(3,6-Diamino-1*H*-indazol-1-yl)-2-methyl-3-phenoxypropan-1-one ((*S*)-SX270, 5e)**

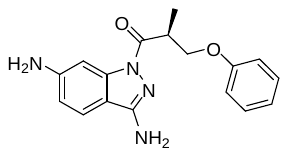

To a solution of **12a** (34.8 mg, 0.19 mmol) in DMF (2.5 mL) were added HOAt (25.9 mg, 0.19 mmol) and EDC × HCl (36.4 mg, 0.19 mmol) at rt. The resulting solution was stirred for 10 min at rt and then 1*H*‑indazole-3,6-diamine (**25**, 34.3 mg, 0.23 mmol) in DMF (0.5 mL) was added. After stirring the reaction mixture at rt for 2 h, the solvent was removed by lyophilization. Purification by preparative HPLC using a solvent system of CH_3_OH / 0.1% aq. HCOOH, a flow rate of 12 mL / min and a gradient of 50% to 71% CH_3_OH in 11 min, 71% to 95% CH_3_OH in 2 min, 95% CH_3_OH for 2 min (*t*_R_ = 8.0 min, λ = 254 nm) afforded **(*S*)-SX270** (**5e**, 25.7 mg, 82.9 μmol, 44%) as an off-white solid.

ESI-MS:	*m/z* 311.1 [M+H]^+^.

HR-ESI-MS:	*m/z* [M+H]^+^ calcd. 311.1503 for C_17_H_19_N_4_O_2_, found 311.1505.

IR (NaCl):	3452, 3353, 2925, 1676, 1625, 1409, 1238 cm^-1^.

m.p.:	75 °C.

^1^H-NMR:	(600 MHz, DMSO-d_6_) *δ* 1.25 (d, *J* = 6.9 Hz, 3H), 4.00 (dd, *J* = 8.7, 5.6 Hz, 1H), 4.04 (ddq, *J* = 7.4, 6.9, 5.6 Hz, 1H), 4.36 (dd, *J* = 8.7, 7.4 Hz, 1H), 5.66 (s, 2H), 6.15 (s, 2H), 6.55 (dd, *J* = 8.5, 1.9 Hz, 1H), 6.90 – 6.95 (m, 3H), 7.25 – 7.29 (m, 2H), 7.42 (d, *J* = 1.9 Hz, 1H), 7.46 (d, *J* = 8.5 Hz, 1H).

^13^C-NMR:	(DEPTQ, 151 MHz, DMSO-d_6_) *δ* 14.1, 37.6, 69.0, 98.1, 110.3, 112.0, 114.5, 120.6, 121.0, 129.5, 141.6, 151.2, 153.1, 158.4, 171.7.

HPLC:	system 1: λ = 254 nm*, t*_R_ = 17.3 min, purity: 98%.

chiral HPLC:	isocratic elution with *n*-hexane / isopropanol + 0.1% ethylene diamine, 9:1: *t*_R_ = 41.4 min, >99% *ee* (254 nm).

[*α*]*_D_^27^*:	+64.3 (*c* = 0.5, methanol).

**3-Amino-1*H*-indazole-6-carboxamide (27)**

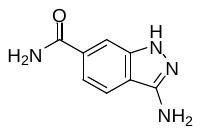

In a solution of 3-amino-1*H*-indazole-6-carboxylic acid (**26**, 75.0 mg, 0.42 mmol) and DIPEA (1.62  mL, 9.31 mmol) in DMF (2 mL) was suspended ammonium chloride (452 mg, 0.10 mmol) and then PyBOP (229 mg, 0.42 mmol) was added. After stirring the reaction mixture for 12 h at rt, the solvent was removed by lyophilization. The residue was purified by preparative HPLC using a solvent system of CH_3_CN/0.1% aq. TFA, a flow rate of 10 mL / min and a gradient of 2% to 17% CH_3_CN in 17 min (*t*_R_: 7.1 min, λ = 220 nm). Subsequent flash chromatography on silica gel (CH_2_Cl_2_ / MeOH 95:5 🡪 9:1 🡪 8:2) afforded **27** (62.0 mg, 0.35 mmol, 83%) as a white solid.

ESI-MS:	*m/z* 176.9 [M+H]^+^.

HR-ESI-MS:	*m/z* [M+H]^+^ calcd. 177.0771 for C_8_H_9_N_4_O, found 177.0770.

IR (KBr):	3418, 3387, 3300, 3188, 1670 cm^-1^.

m.p.:	225 °C (decomposition).

^1^H-NMR:	(600 MHz, DMSO-d_6_) *δ* 5.40 (s, 2H), 7.29 (brs, 1H), 7.39 (dd, *J* = 8.3, 1.3 Hz, 1H), 7.70 (d, *J* = 8.3 Hz, 1H), 7.76 (brs, 1H), 7.97 (brs, 1H), 11.64 (brs, 1H).

^13^C-NMR:	(DEPTQ, 151 MHz, DMSO-d_6_) *δ* 109.2, 115.3, 116.6, 119.8, 132.2, 138.2, 140.9, 149.1, 168.5.

HPLC:	system 4: λ = 220 nm*, t*_R_ = 8.5 min, purity: 99%.

**(*S*)-3-Amino-1-(2-methyl-3-phenoxypropanoyl)-1*H*-indazole-6-carboxamide (I1424, 5f)**

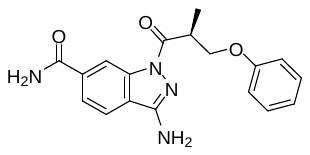

To a solution of **12a** (12.3 mg, 0.07 mmol) and HOAt (9.3 mg, 0.07 mmol) in DMF (0.5 mL) was added EDC × HCl (13.1 mg, 0.09 mmol). After for 10 min at rt, this solution was added to a suspension of 3-amino-1*H*-indazole-6-carboxamide (**27**, 12.0 mg, 0.09 mmol) in DMF (0.5 mL). After stirring the reaction mixture for 3 h, the solvent was removed by lyophilization and the residue was purified by preparative HPLC using a solvent system of CH_3_OH / 0.1% aq. HCOOH, a flow rate of 10 mL / min and a gradient of 10% to 45% CH_3_OH in 22 min (*t*_R_: 11.7 min, λ = 220 nm) to afford **I1424** (**5f**, 17.1 mg, 50.6 µmol, 64%) as a white resin.

ESI-MS:	*m/z* 339.2 [M+H]^+^.

HR-ESI-MS:	*m/z* [M+Na]^+^ calcd. 361.1271 for C_18_H_18_N_4_NaO_3_, found 361.1268.

IR (KBr):	3447, 3354, 3310, 3242, 1685 cm^-1^.

^1^H-NMR:	(600 MHz, DMSO-d_6_) *δ* 1.30 (d, *J* = 7.0 Hz, 3H), 4.08 (dd, *J* = 9.3, 5.4 Hz, 1H), 4.12 (ddq, *J* = 7.2, 5.4, 7.0 Hz, 1H), 4.40 (dd, *J* = 9.3, 7.2 Hz, 1H), 6.63 (brs, 1H), 6.89 – 6.96 (m, 3H), 7.24 – 7.30 (m, 2H), 7.46 (brs, 1H), 7.81 (dd, *J* = 8.3, 1.5 Hz, 1H), 7.94 (d, *J* = 8.3 Hz, 1H), 8.14 (brs, 1H), 8.74 (brs, 1H).

^13^C-NMR:	(DEPTQ, 151 MHz, DMSO-d_6_) *δ* 14.0, 37.8, 68.9, 114.5, 114.7, 120.4, 120.7, 121.9, 122.9, 129.5, 135.9, 139.0, 152.7, 158.3, 167.8, 172.3.

HPLC:	system 3: λ = 220 nm*, t*_R_ = 18.9 min, purity: >99%.

[*α*]*_D_^22^*:	+44.5 (*c* = 0.25, methanol).

**(*R*)-3-Amino-1-(2-methyl-3-phenoxypropanoyl)-1*H*-indazole-6-carboxamide (I1422, ent-5f)**

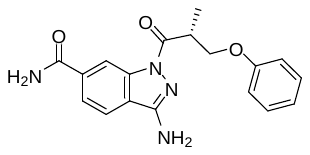

**I1422 (ent-5f)** was synthesized analogously as described for **I1424 (5f)**, starting from **ent-12a**. The analytical data were in accordance.

HPLC:	system 3: λ = 220 nm*, t*_R_ = 18.9 min, purity: 98%.

[*α*]*_D_^22^*:	-44.5 (*c* = 0.7, methanol).

**(3-Amino-1*H*-indazol-6-yl)methanol (29)**

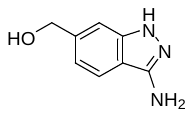

A solution of commercially available 2-fluoro-4-(hydroxymethyl)benzonitrile (**28**, 100 mg, 0.66 mmol) and hydrazine hydrate (319 µL, 6.62 mmol) in *n*-butanol (3 mL) was heated at 120 °C in a sealed tube for 4 h. After cooling to rt, the resulting solution was diluted with a 20% aqueous Na_2_CO_3_-solution, extracted with ethyl acetate and dried over MgSO_4_. After evaporation of the solvent, flash chromatography on silica gel (CH_2_Cl_2_ / MeOH 95:5 🡪 9:1) was performed to afford **29** (74 mg, 0.45 mmol, 69%) as a white solid.

ESI-MS:	*m/z* 163.8 [M+H]^+^.

HR-ESI-MS:	*m/z* [M+H]^+^ calcd. 164.0818 for C_8_H_10_N_3_O, found 164.0819.

IR (KBr):	3438, 3343, 3199 cm^-1^.

m.p.:	214-215 °C.

^1^H-NMR:	(600 MHz, DMSO-d_6_) *δ* 4.56 (s, 2H), 5.20 (s, 1H), 5.24 (s, 2H), 6.83 (d, *J* = 8.3 Hz, 1H), 7.18 (s, 1H), 7.59 (d, *J* = 8.3 Hz, 1H), 11.28 (brs, 1H).

^13^C-NMR:	(DEPTQ, 151 MHz, DMSO-d_6_) *δ* 63.2, 106.5, 113.0, 116.6, 119.7, 140.9, 141.8, 148.9.

**(*S*)-1-(3-Amino-6-(hydroxymethyl)-1*H*-indazol-1-yl)-2-methyl-3-phenoxypropan-1-one (I1408, 5g)**

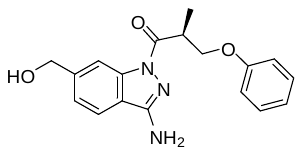

To a solution of (**12a**,15.0 mg, 0.08 mmol) and HOAt (13.6 mg, 0.10 mmol) in DMF (0.5 mL) was added EDC × HCl (19.2 mg, 0.10 mmol). After stirring for 10 min at rt, a solution of **29** (16.3 mg, 0.10 mmol) in DMF (0.5 mL) was added. After stirring the reaction mixture for 3 h, the solvent was removed by lyophilization and the residue was purified by preparative HPLC using a solvent system of CH_3_OH / 0.1% aq. HCOOH, a flow rate of 10 mL / min and a gradient of 50% to 80% CH_3_OH in 15 min (*t*_R_: 12.5 min, λ = 220 nm) to afford **I1408** (**5g**, 14.4 mg, 44.3 µmol, 53%) as a white solid.

ESI-MS:	*m/z* 326.1 [M+H]^+^.

HR-ESI-MS:	*m/z* [M+H]^+^ calcd. 326.1499 for C_18_H_19_N_3_O_3_, found 326.1499.

IR (KBr):	3478, 3357, 1636 cm^-1^.

m.p.:	132 °C.

^1^H-NMR:	(600 MHz, DMSO-d_6_) *δ* 1.28 (d, *J* = 7.0 Hz, 3H), 4.05 (dd, *J* = 8.8, 5.5 Hz, 1H), 4.11 (ddq, *J* = 8.0, 5.5, 7.0 Hz, 1H), 4.39 (dd, *J* = 8.8, 8.0 Hz, 1H), 4.64 (d, *J* = 5.8 Hz, 2H), 5.37 (t, *J* = 5.8 Hz, 1H), 6.49 (s, 2H), 6.90 – 6.95 (m, 3H), 7.25 – 7.30 (m, 3H), 7.82 (d, *J* = 8.0 Hz, 1H), 8.27 (brs, 1H).

^13^C-NMR:	(DEPTQ, 151 MHz, DMSO-d_6_) *δ* 14.1, 37.8, 63.0, 69.0, 112.8, 114.5, 119.2, 120.3, 122.4, 129.5, 139.7, 145.0, 152.9, 158.4, 172.2.

HPLC:	system 3: λ = 220 nm*, t*_R_ = 20.2 min, purity: 99%.

chiral HPLC:	isocratic elution with *n*-hexane / ethanol, 9:1: *t*_R_ = 16.7 min, 98% *ee* (254 nm).

[*α*]*_D_^22^*:	+27.9 (*c* = 0.5, methanol).

**(*R*)-1-(3-Amino-6-(hydroxymethyl)-1*H*-indazol-1-yl)-2-methyl-3-phenoxypropan-1-one (I1411, ent-5g)**

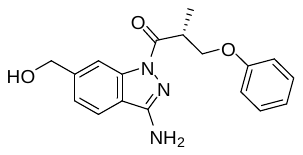

**I1411** **(ent-5g)** was synthesized analogously as described for **I1408 (5g)**, starting from **ent-12a**. The analytical data were in accordance.

HPLC:	system 3: λ = 220 nm*, t*_R_ = 20.2 min, purity: 99%.

chiral HPLC:	isocratic elution with *n*-hexane / isopropanol, 99:1: *t*_R_ = 13.5 min, 98% *ee* (254 nm).

[*α*]*_D_^22^*:	-28.2 (*c* = 0.4, methanol).

**(*S*)-1-(3-Amino-6-(hydroxymethyl)-1*H*-indazol-1-yl)-3-(4-fluorphenoxy)-2-methylpropan-1-one (I1421, F-5g)**

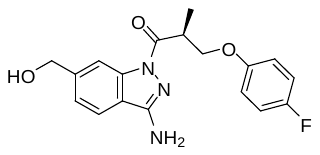

To a solution of **12e** (18.0 mg, 0.09 mmol) and HOAt (12.4 mg, 0.09 mmol) in DMF (0.5 mL) was added EDC × HCl (17.4 mg, 0.09 mmol). After stirring for 10 min at rt, a solution of **29** (14.8 mg, 0.09 mmol) in DMF (0.5 mL) was added. After stirring the reaction mixture for 3 h, the solvent was removed by lyophilization and the residue was purified by preparative HPLC using a solvent system of CH_3_OH / 0.1% aq. HCOOH, a flow rate of 10 mL / min and a gradient of 50% to 80% CH_3_OH in 15 min (*t*_R_: 10.2 min, λ = 220 nm) to afford **I1421** (**F-5g**, 14.2 mg, 41.4 µmol, 45%) as a white solid.

ESI-MS:	*m/z* 344.3 [M+H]^+^.

HR-ESI-MS:	*m/z* [M+H]^+^ calcd. 344.1404 for C_18_H_19_FN_3_O_3_, found 344.1404.

IR (KBr):	3473, 3357, 3314, 1636 cm^-1^.

m.p.:	136 °C.

^1^H-NMR:	(600 MHz, DMSO-d_6_) *δ* 1.27 (d, *J* = 7.3 Hz, 3H), 4.03 (dd, *J* = 9.1, 5.6 Hz, 1H), 4.09 (ddq, *J* = 7.9, 5.6, 7.3 Hz, 1H), 4.36 (dd, *J* = 9.1, 7.9 Hz, 1H), 4.63 (d, *J* = 5.6 Hz, 2H), 5.36 (t, *J* = 5.6 Hz, 1H), 6.94 – 6.98 (m, 2H), 7.07 – 7.12 (m, 2H), 7.28 (dd, *J* = 8.3, 1.1 Hz, 1H), 7.82 (d, *J* = 8.3 Hz, 1H), 8.26 (brs, 1H).

^13^C-NMR:	(DEPTQ, 151 MHz, DMSO-d_6_) *δ* 14.0, 37.7, 62.9, 69.7, 112.7, 115.77 (d, *J* = 12 Hz), 115.82 (d, *J* = 19 Hz), 119.1, 120.2, 122.4, 139.6, 145.0, 152.9, 154.7, 156.5 (d, *J* = 239 Hz), 172.1.

HPLC:	system 3: λ = 220 nm*, t*_R_ = 20.4 min, purity: >99%.

chiral HPLC:	isocratic elution with *n*-hexane / ethanol, 9:1: *t*_R_ = 18.7 min, 99% *ee* (254 nm).

[*α*]*_D_^22^*:	+31.8 (*c* = 0.5, methanol).

**(*S*)-*N*-(3-Amino-1-(2-methyl-3-phenoxypropanoyl)-1*H*-indazol-6-yl)formamide ((*S*)-SX276, 6a)**

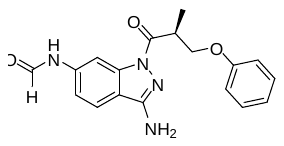

A mixture of formic acid (15.1 µL, 18.5 mg, 401 µmol) and acetic anhydride (25.2 µL, 27.3 mg, 268 µmol) was stirred at 60 °C for 2 h. After cooling to rt, the mixed anhydride was added dropwise to a solution of compound **(*S*)-SX270** (**5e**, 16.6 mg, 53.5 μmol) in anhydrous THF (1 mL) at 0 °C. After stirring the reaction mixture for 30 min at rt, the mixture was *immediately* treated with CH_3_OH / water (1:2) in order to hydrolyze the excess of mixed anhydride and then lyophilization was performed. Purification by preparative HPLC using a solvent system of CH_3_OH / 0.1% aq. HCOOH, a flow rate of 12 mL / min and a gradient of 50% to 75% CH_3_OH in 12.5 min, 75% to 95% CH_3_OH in 0.5 min, 95% CH_3_OH for 1 min (*t*_R_ = 9.7 min, λ = 254 nm) afforded **(*S*)-SX276** (**6a**, (7.64 mg, 22.6 μmol, 42%) as a white solid.

ESI-MS:	*m/z* 339.2 [M+H]^+^.

HR-ESI-MS:	*m/z* [M+H]^+^ calcd. 339.1452 for C_18_H_19_N_4_O_3_, found 339.1452.

IR (NaCl):	3442, 3336, 2925, 1683, 1618, 1436, 1235 cm^-1^.

m.p.:	77 °C.

^1^H-NMR:	(600 MHz, DMSO-d_6_, two sets of resonances were observed, rotamers, assignment was confirmed by 2D spectroscopy) *δ* 1.28 (d, *J* = 6.8 Hz, 3H), 4.05 (dd, *J* = 8.8, 5.5 Hz, 1H), 4.07 (ddq, *J* = 7.4, 6.8, 5.5 Hz 1H), 4.38 (dd, *J* = 8.8, 7.4 Hz, 1H), 6.47 (s, 1.6 H), 6.49 (s, 0.4 H), 6.90 – 6.96 (m, 3H), 7.24 – 7.29 (m, 2 H and 0.2 H), 7.50 (dd, *J* = 8.5, 1.8 Hz, 0.8 H), 7.81 (d, *J* = 8.4 Hz, 0.8 H), 7.82 (d, *J* = 8.6 Hz, 0.8 H),, 8.04 (d, *J* = 1.9 Hz, 0.2 H), 8.34 (d, *J* = 1.8 Hz, 0.8 H), 8.68 (d, *J* = 1.7 Hz, 0.8 H), 8.89 (d, *J* = 11.0 Hz, 0.2 H), 10.43 (d, *J* = 11.0 Hz, 0.2 H), 10.48 (d, *J* = 1.9 Hz, 0.8 H).

^13^C-NMR:	(DEPTQ, 151 MHz, DMSO-d_6_, two sets of resonances were observed, rotamers) *δ* 14.0 (minor), 14.0 (major), 37.7, 68.9 (minor), 68.9 (major), 103.4 (minor), 105.3 (major), 113.8 (minor), 114.5, 115.7 (major), 116.1 (major), 116.3 (minor), 120.6, 121.1 (major), 121.8 (minor), 129.5, 139.7 (major), 139.9 (major), 140.1 (minor), 140.2 (minor), 152.7 (major), 152.8 (minor), 158.3, 159.8 (major), 162.6 (minor), 172.1 (major), 172.2 (minor).

HPLC:	system 2: λ = 254 nm, *t*_R_ = 16.1 min, purity: >99%.

chiral HPLC:	isocratic elution with *n*-hexane / isopropanol, 9:1: *t*_R_ = 40.7 min, >99% *ee* (254 nm).

[*α*]*_D_^26^*:	+78.1 (*c* = 0.3, methanol).

**(*S*)-*N,N'*-(1-(2-Methyl-3-phenoxypropanoyl)-1*H*-indazole-3,6-diyl)diformamide ((*S*)-SX272, 6b)**

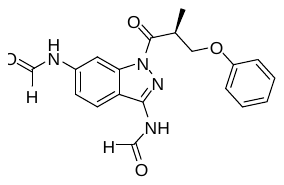

A mixture of formic acid (14.9 µL, 18.1 mg, 394 µmol) and acetic anhydride (24.8 µL, 26.8 mg, 263 µmol) was stirred at 60 °C for 2 h. After cooling to rt, the mixed anhydride was added dropwise to a solution of compound **(*S*)-SX270** (**5e**, 16.3 mg, 52.5 mmol) in anhydrous THF (2 mL) at 0 °C. After stirring the reaction mixture for 1 h at rt, the solvent was removed under reduced pressure. The residue was treated with a mixture of CH_3_OH / water (1:2) and lyophilized. After purification by preparative HPLC using a solvent system of CH_3_OH / 0.1% aq. HCOOH, a flow rate of 12 mL / min and a gradient of 50% to 75% CH_3_OH in 12.5 min, 75% to 95% CH_3_OH in 0.5 min, 95% CH_3_OH for 1 min (*t*_R_ = 11.0 min, λ = 254 nm), **(*S*)-SX272** (**6b**, 6.52 mg, 19.3 μmol, 37%) was obtained as a white solid.

ESI-MS:	*m/z* 367.2 [M+H]^+^.

HR-ESI-MS:	*m/z* [M+H]^+^ calcd. 367.1401 for C_19_H_19_N_4_O_4_, found 367.1404.

IR (NaCl):	3354, 2921, 1689, 1632, 1240 cm^-1^.

^1^H-NMR:	(400 MHz, DMSO-d_6_, two major sets of resonances, rotamers) *δ* 1.33 (d, *J* = 6.7 Hz, 3 H), 4.11 – 4.27 (m, 2H), 4.32 – 4.43 (m, 1H), 6.89 – 6.97 (m, 3H), 7.23 – 7.31 (m, 2H), 7.40 (brd, *J* = 8.9 Hz, 0.2 H), 7.53 (brd, *J* = 8.8 Hz, 0.8 H), 7.98 (bd, *J* = 8.5 Hz, 1H), 8.16 – 8.08 (m, 0.2 H), 8.38 (d, *J* = 1.7 Hz, 0.8 H), 8.55 (s, 0.2 H), 8.87 (brs, 0.8 H), 8.97 (d, *J* = 10.1 Hz, 0.2 H), 9.10 (brs, 0.8 H), 10.58 (d, *J* = 10.1 Hz, 0.2 H), 10.68 (brs, 0.8 H), 11.31 (brs, 1H).

^13^C-NMR:	(DEPTQ, 151 MHz, DMSO-d_6_, two sets of resonances were observed, rotamers) *δ* 13.9 (major), 14.0 (minor), 37.7 (minor), 38.0 (major), 68.9, 102.6 (minor), 104.7 (major), 114.5 (minor), 114.5 (major), 114.8 (minor), 116.9 (major), 120.7 (major), 121.2 (minor), 129.4, 139.7, 139.7, 140.6, 145.1, 158.2, 160.1, 162.2, 162.7, 173.3 (major), 173.4 (minor).

HPLC:	system 1: λ = 254 nm*, t*_R_ = 18.1 min, purity: 96%.

chiral HPLC:	isocratic elution with *n*-hexane / isopropanol + 0.1% ethylene diamine, 8:2: *t*_R_ = 28.3 min, 97% *ee* (254 nm).

[*α*]*_D_^26^*:	+18.3 (*c* = 0.1, methanol).

**(*S*)-1-(2-Methyl-3-phenoxypropanoyl)-1*H*-indazole-6-carboxamide (I1409, 7a)**

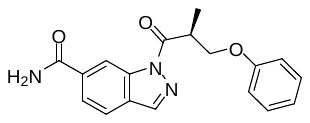

To a solution of **12a** (15.0 mg, 0.08 mmol) and HOAt (13.6 mg, 0.10 mmol) in DMF (0.5 mL) was added EDC × HCl (19.2 mg, 0.10 mmol). After stirring for 10 min at rt, a solution of commercially available 1*H*-indazole-6-carboxamide (**30**, 16.1 mg, 0.10 mmol) in DMF (0.5 mL) was added. After stirring the reaction mixture for 3 h, the solvent was removed by lyophilization and the residue was purified by preparative HPLC using a solvent system of CH_3_OH / 0.1% aq. HCOOH, a flow rate of 10 mL / min and a gradient of 50% to 80% CH_3_OH in 15 min, 80% to 95% CH_3_OH in 2 min (*t*_R_: 15.5 min, λ = 220 nm) to afford **I1409** (**7a**, 17.1 mg, 52.9 µmol, 64%) as a white solid.

ESI-MS:	*m/z* 324.1 [M+H]^+^.

HR-ESI-MS:	*m/z* [M+H]^+^ calcd. 324.1343 for C_18_H_17_N_3_O_3_, found 324.1345.

IR (KBr):	3482, 3437, 3397, 3357, 3197, 1718, 1707, 1662 cm^-1^.

m.p.:	141 °C.

^1^H-NMR:	(600 MHz, DMSO-d_6_) *δ* 1.38 (d, *J* = 7.2 Hz, 3H), 4.21 (dd, *J* = 10.0, 5.3 Hz, 1H), 4.35 (ddq, *J* = 6.8, 5.3, 7.2 Hz, 1H), 4.43 (dd, *J* = 10.0, 6.8 Hz, 1H), 6.90-6.94 (m, 3H), 7.24 – 7.28 (m, 2H), 7.53 (brs, 1H), 7.91 (dd, *J* = 8.3, 1.6 Hz, 1H), 7.98 (dd, *J* = 8.3, 0.6 Hz, 1H), 8.24 (brs, 1H), 8.60 (d, *J* = 1.0 Hz, 1H), 8.85 (ddd, *J* = 1.6, 1.0, 0.6 Hz, 1H).

^13^C-NMR:	(DEPTQ, 151 MHz, DMSO-d_6_) *δ* 13.9, 38.2, 68.9, 114.2, 114.4, 120.7, 121.4, 123.9, 129.4, 135.7, 138.2, 140.5, 158.1, 167.6, 174.1.

HPLC:	system 3: λ = 220 nm*, t*_R_ = 21.7 min, purity: 98%.

chiral HPLC:	isocratic elution with *n*-hexane / isopropanol, 85:5: *t*_R_ = 21.6 min, 98% *ee* (254 nm).

[*α*]*_D_^22^*:	+111.4 (*c* = 0.4, methanol).

**(*R*)-1-(2-Methyl-3-phenoxypropanoyl)-1*H*-indazole-6-carboxamide (I1412, ent-7a)**

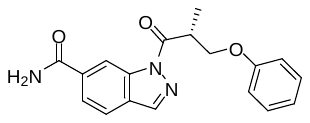

**I1412 (ent-7a)** was synthesized analogously as described for **I1409 (7a)**, starting from **ent-12a**. The analytical data were in accordance.

HPLC:	system 3: λ = 220 nm*, t*_R_ = 21.7 min, purity: 99%.

chiral HPLC:	isocratic elution with *n*-hexane / isopropanol, 85:5: *t*_R_ = 22.7 min, 98% *ee* (254 nm).

[*α*]*_D_^22^*:	-111.0 (*c* = 0.65, methanol).

**(*S*)-1-(6-Amino-1*H*-indazol-1-yl)-2-methyl-3-phenoxypropan-1-one ((*S*)-SX254, 7b)**

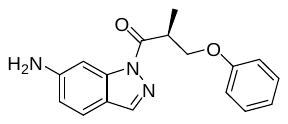

To a stirred solution of compound **12a** (39.8 mg, 0.22 mmol) in anhydrous DMF (1.5 mL) was added HOAt (29.9 mg, 0.22 mmol), followed by EDC × HCl (42.2 mg, 0.22 mmol) at rt. The resulting solution was stirred for 10 min at rt before adding commercially available 6‑amino-1*H*-indazole (**31**, 35.9 mg, 0.27 mmol) dissolved in anhydrous DMF (0.5 mL). After stirring the reaction mixture at rt for 40 min., the solvent was removed by lyophilization. Purification by preparative HPLC using a solvent system of CH_3_OH / 0.1% aq. HCOOH, a flow rate of 10 mL / min and a gradient of 50% to 80% CH_3_OH in 15 min, 80% to 95% CH_3_OH in 2 min, 95% CH_3_OH for 4 min (*t*_R_ = 15.0 min, λ = 254 nm) yielded **(*S*)-SX254** (**7b**, 8.79 mg, 29.8 μmol, 14%) as a colorless resin.

ESI-MS:	*m/z* 296.1 [M+H]^+^.

HR-ESI-MS:	*m/z* [M+H]^+^ calcd. 296.1394 for C_17_H_18_N_3_O_2_, found 296.1392 [M+H]^+^.

IR (NaCl):	3469, 3368, 2920, 2851, 1698, 1618, 1493, 1424, 1406, 1241 cm^-1^.

^1^H-NMR:	(400 MHz, DMSO-d_6_) δ 1.31 (d, *J* = 7.0 Hz, 1H), 4.12 (dd, *J* = 9.1, 5.2 Hz, 1H), 4.25 (ddq, *J* = 7.9, 7.0, 5.2 Hz, 1H), 4.38 (dd, *J* = 9.1, 7.9 Hz, 1H), 5.86 (s, 2H), 6.68 (dd, *J* = 8.5, 2.0 Hz, 1H), 6.87 – 6.98 (m, 3H), 7.23 – 7.30 (m, 2H), 7.46 – 7.51 (m, 2H), 8.15 (d, *J* = 0.7 Hz, 1H).

^13^C-NMR:	(DEPTQ, 101 MHz, DMSO-d_6_) *δ* 14.1, 38.1, 69.0, 96.8, 114.0, 114.5, 116.8, 120.8, 121.9, 129.5, 140.8, 140.9, 151.4, 158.3, 174.0.

HPLC:	system 1: λ = 254 nm*, t*_R_ = 19.2 min, purity: 97%.

chiral HPLC:	isocratic elution with *n*-hexane / isopropanol, 9:1: *t*_R_ = 12.3 min, >99% *ee*  (254 nm).

[*α*]*_D_^23^*:	+139.1 (*c* = 0.1, methanol).

**(*S*)-*N*-(1-(2-Methyl-3-phenoxypropanoyl)-1*H*-indazol-6-yl)formamide [(*S*)-SX258, 8]**

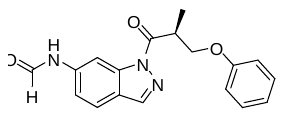

A mixture of formic acid (5.9 µL, 7.25 mg, 157 µmol) and acetic anhydride (9.9 µL, 10.7 mg, 105 µmol) was stirred at 60 °C for 2 h. After cooling to rt, the mixed anhydride was added dropwise to a solution of compound **(*S*)-SX254** **7b** (6.2 mg, 21.0 µmol) in anhydrous THF (1 mL) at 0 °C. After stirring the reaction mixture for 1.5 h at rt, the solvent was removed under reduced pressure. The residue was treated with CH_3_OH / water (1:2) and lyophilized. Purification by preparative HPLC using a solvent system of CH_3_OH / 0.1% aq. HCOOH, a flow rate of 10 mL / min and a gradient of 50% to 80% CH_3_OH in 15 min, 80% to 95% CH_3_OH in 2 min, 95% CH_3_OH for 1 min (*t*_R_ = 15.0 min, λ = 254 nm) yielded **(*S*)-SX258** **8** (1.87 mg, 5.78 μmol, 28%) as a white solid.

ESI-MS:	*m/z* 324.2 [M+H]^+^.

HR-ESI-MS:	*m/z* [M+H]^+^ calcd. 324.1343 for C_18_H_18_N_3_O_3_, found 324.1343.

IR (NaCl):	3357, 3275, 2922, 2850, 1700, 1618, 1597, 1491, 1409, 1241, 1183 cm^-1^.

^1^H-NMR:	(600 MHz, DMSO-d_6_, two sets of signals were observed, rotamers) *δ* 1.35 (d, *J* = 7.0 Hz, 3H), 4.17 (dd, *J* = 9.3, 5.2 Hz, 1H), 4.31 (ddq, *J* = 7.9, 7.0, 5.2 Hz, 1H), 4.40 (dd, *J* = 9.3, 7.9 Hz, 1H), 6.89 – 6.95 (m, 3H), 7.23 – 7.30 (m, 2H), 7.38 (dd, *J* = 8.4, 1.7 Hz, 0.2 H), 7.54 (dd, *J* = 8.4, 1.7 Hz, 0.8 H), 7.85 (d, *J* = 8.4 Hz, 0.8 H), 7.86 – 7.87 (m, 0.2 H), 8.12 – 8.14 (m, 0.2 H), 8.37 (d, *J* = 1.7 Hz, 0.8 H), 8.44 (d, *J* = 0.9 Hz, 0.8 H), 8.55 – 8.56 (m, 0.2 H), 8.84 – 8.88 (m, 0.8 H), 8.94 (d, *J* = 10.6 Hz, 0.2 H), 10.51 (d, *J* = 10.6 Hz, 0.2 H), 10.57 – 10.61 (m, 0.8 H).

^13^C-NMR:	(DEPTQ, 151 MHz, DMSO-d_6_, two sets of signals were observed, rotamers) *δ* 14.0, 38.2, 69.0, 102.5 (minor), 104.5 (major), 114.6 (major), 115.4 (minor), 117.2, 120.8, 122.1 (major), 122.2 (major), 122.4 (minor), 122.8 (minor), 129.5, 139.1 (major), 139.3 (minor), 139.7 (major), 140.2 (minor), 140.6 (major), 140.7 (minor), 158.2, 160.0, 162.7 (major), 165.0 (minor), 174.1 (major), 174.2 (minor).

HPLC:	system 1: λ = 254 nm, *t*_R_ = 19.6 min, purity: 97%.

chiral HPLC:	isocratic elution with *n*-hexane / isopropanol, 9:1: *t*_R_ = 28.4 min, 99% *ee* (254 nm).

[*α*]*_D_^23^*:	+74.8 (*c* = 0.1, methanol).

**7-Fluoro-2*H*-chromene (33)** ^7^

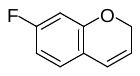

(Acetonitrile)[2-biphenyl)di-*tert*-butylphosphine]gold(I) hexafluoroantimonate (0.50 g, 0.66 mmol) was added slowly to an ice cooled solution of **32** ^7^ (6.00 g, 39.9 mmol) in toluene (10 mL). After stirring for 90 min the resulting solution was directly subjected to flash chromatography on silica gel (isohexane 100%) to afford crude **33** (5.80 g, 97%) as a volatile, colourless oil, contaminated with residual catalyst and the other ring closing isomer 5-fluoro-2*H*-chromene (14%, determined by ^1^H NMR). This crude material was used for the following reaction without further purification.

^1^H NMR:	(600 MHz, CDCl_3_) *δ* 4.81 (dd, *J* = 3.6, 2.0 Hz, 2H), 5.70 (dd, *J* = 10.0, 3.6 Hz, 2H), 6.38 (dd, *J* = 10.0, 2.0 Hz, 2H), 6.50 (dd, *J* = 10.1, 2.7 Hz, 1H), 6.55 (ddd, *J* = 8.5, 8.4, 2.7 Hz, 1H), 6.89 (dd, *J* = 8.5, 6.3 Hz, 1H).

The NMR data of this compound are in accordance with those reported in the literature.

**7-Fluorochromane (34)** ^7^

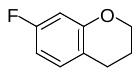

To a solution of crude **33** (5.80 g) in methanol / ethyl acetate 1:1 (200 mL) was added palladium hydroxide 20% on charcoal (0.50 g). Hydrogenation by stirring for 2 h under a balloon filled with H_2_ was performed at rt. The resulting solution was filtered through a pad of celite and the solvent was removed *in vacuo*, not lower than 200 mbar to afford crude **34** (5.80 g) as a volatile, colourless liquid, still contaminated with residual catalyst and the other ring closing isomer 5-fluorochromane formed by the hydrogenation of the 5-fluoro-2*H*-chromene byproduct. This crude material was used for the following reaction without further purification.

^1^H NMR:	(600 MHz, CDCl_3_) *δ* 1.99 (tt, *J* = 6.4, 5.2 Hz, 2H), 2.74 (t, *J* = 6.4 Hz, 2H), 4.17 (t, *J* = 5.2 Hz, 2H), 6.51 (dd, *J* = 10.4, 2.5 Hz, 1H), 6.55 (ddd, *J* = 8.4, 8.3, 2.5 Hz, 1H), 6.96 (ddd, *J* = 8.3, 6.8, 0.8 Hz, 1H).

The NMR data of this compound are in accordance with those reported in the literature.

**6-Bromo-7-fluorochromane (35)** ^7^

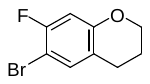

To a solution of the crude **34** (5.80 g, 38.1 mmol) in acetonitrile (40 mL) was added slowly *N*-bromosuccinimide (5.43 g, 30.5 mmol) at 0 °C. After stirring for 2 h, another portion of *N*-bromosuccinimide (0.68 g, 4,54 mmol) was added. After stirring at 0 °C for 1 h, allowing a full conversion of the educt according to TLC, water was added and the mixture was allowed to warm to ambient temperature. Extraction with ethyl acetate (4x) was performed and the organic layer was washed with water and dried over MgSO_4_. Gradual evaporation of the solvent was done and the precipitate, which was formed (succinimide), was removed by filtration. Flash chromatography (isohexane 100%) afforded **35** (2.20 g, 9.51 mmol, 23% over the last three steps) as a colourless oil, containing another regioisomer (6% according to NMR) resulting from the bromination of the 5-fluorochromane impurity. This compound was employed for the next reaction without further purification.

^1^H NMR:	(400 MHz, CDCl_3_) *δ* 1.98 (tt, *J* = 6.4, 5.2 Hz, 2H), 2.73 (t, *J* = 6.4 Hz, 2H), 4.16 (t, *J* = 5.2 Hz, 2H), 6.58 (d, *J* = 9.7 Hz, 1H), 7.17 (dd, *J* = 7.2, 1.0 Hz, 1H).

The NMR data of this compound are in accordance with those reported in the literature.

**7-Fluorochromane-6-carbonitrile (36)**

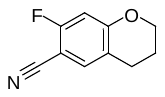

To a solution of **35** (700 mg, 3.03 mmol) in THF (14 mL) at -79 °C was added slowly a 2.5 M solution of butyllithium in hexane (1.33 mL, 3.33 mmol). After stirring for 20 min, a solution of dimethylmalononitrile (428 mg, 4.54 mmol) in THF (2 mL) was added slowly. After stirring at ‑79 °C for additional 30 min, the reaction was stopped by the addition of a saturated aqueous NH_4_Cl-solution, warmed to rt, extracted with *tert*-butyl methyl ether and dried over MgSO_4_. After evaporation of the solvent, flash chromatography on silica gel (isohexane / ethyl acetate, 99:1 🡪 98:2) was performed to afford **36** (290 mg, 1.47 mmol, 54%) as a white solid.

ESI-MS:	*m/z* 178.1 [M+H]^+^.

HR-ESI-MS:	*m/z* [M+H]^+^ calcd. 178.0663 for C_10_H_9_FNO, found 178.0663.

IR (NaCl):	2229 cm^-1^.

m.p.:	102 °C.

^1^H-NMR:	(400 MHz, CDCl_3_) *δ* 2.02 (tt, *J* = 6.4, 5.2 Hz, 2H), 2.76 (t, *J* = 6.4 Hz, 2H), 4.25 (t, *J* = 5.2 Hz, 2H), 6.59 (d, *J* = 5.2 Hz, 1H), 7.27 (dt, *J* = 7.2, 1.0 Hz, 1H).

^13^C-NMR:	(DEPTQ, 151 MHz, DMSO-d_6_) *δ* 21.5, 24.1, 67.2, 92.5 (d, *J* = 15 Hz), 104.8 (d, *J* = 22 Hz), 114.7, 119.4 (d, *J* = 3 Hz), 134.2 (d, *J* = 2 Hz), 160.1 (d, *J* = 12 Hz), 162.6 (d, *J* = 255 Hz).

**1,5,6,7-Tetrahydropyrano[3,2-*f*]indazol-3-amine (37)**

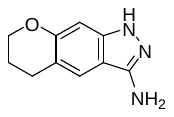

A solution of **36** (200 mg, 1.13 mmol) and hydrazine hydrate (550 µL, 11.3 mmol) in *n*-butanol (5 mL) was heated at 120 °C in a sealed tube. After 20 h, another portion of hydrazine hydrate (280 µL, 6.64 mmol) was added and heating was continued for additional 2 h. After cooling to ambient temperature, the resulting solution was diluted with a 20% aqueous Na_2_CO_3_-solution, extracted with ethyl acetate and dried over MgSO_4_. After evaporation of the solvent, flash chromatography (CH_2_Cl_2_ / CH_3_OH, 100:0 🡪 99:1 🡪 98:2) was performed to afford **37** (162 mg, 0.86 mmol, 76%) as a white solid.

ESI-MS:	*m/z* 189.8 [M+H]^+^.

HR-ESI-MS:	*m/z* [M+H]^+^ calcd. 190.0975 for C_10_H_12_N_3_O, found 190.0975.

IR (KBr):	3350, 3275, 3172, 1636 cm^-1^.

m.p.:	158-159 °C.

^1^H-NMR:	(600 MHz, DMSO-d_6_) *δ* 2.02 (tt, *J* = 6.3, 5.3 Hz, 2H), 2.82 (t, *J* = 6.3 Hz, 2H), 4.12 (t, *J* = 5.3 Hz, 2H), 5.23 (s, 2H), 6.46 (s, 1H), 7.33 (s, 1H), 10.84 (s, 1H).

^13^C-NMR:	(DEPTQ, 151 MHz, DMSO-d_6_) *δ* *δ* 24.2, 27.3, 69.4, 94.1, 109.3, 114.4, 120.0, 141.5, 148.6, 154.1.

**(*R*,*S*)-1-(3-Amino-6,7-dihydropyrano[3,2-*f*]indazol-1(5*H*)-yl)-2-methyl-3-phenoxypropan-1-one (I1380, 9)**

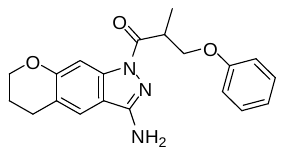

To a solution of commercially available (*R*,*S*)-2-methyl-3-phenoxypropionic acid (**rac-12a**, 57.1 mg, 0.32 mmol), PyBOP (165 mg, 0.32 mmol) and HOAt (60.0 mg, 0.48 mmol) in DMF (1.5 mL) was added DIPEA (55 µL, 0.44 mmol) and then a solution of **32** (50.0 mg, 0.26 mmol) in DMF (0.5 mL) at rt. Microwave irradiation (described in the general part) in a sealed tube was performed, then the solvent was removed by lyophilization and the residue was purified by preparative HPLC using a solvent system of CH_3_CN / 0.1% aq. TFA, a flow rate of 10 mL / min and a gradient of 40% to 65% CH_3_CN in 19 min, 65% to 95% CH_3_CN in 2 min (*t*_R_: 19.0 min, λ = 220 nm) to afford **I1380** (**9**, 56.3 mg, 0.16 mmol, 61%) as a white solid.

ESI-MS:	*m/z* 352.1 [M+H]^+^.

HR-ESI-MS:	*m/z* [M+H]^+^ calcd. 352.1656 for C_20_H_22_N_3_O_3_, found 352.1660.

IR (KBr):	3424, 3353, 1647 cm^-1^.

m.p.:	185 °C.

^1^H-NMR:	(600 MHz, DMSO-d_6_) *δ* 1.24 – 1.27 (m, 3H), 1.95 (tt, *J* = 6.3, 5.1 Hz, 1H), 2.86 (t, *J* = 6.3 Hz, 2H), 4.00 – 4.07 (m, 2H), 4.20 (t, *J* = 5.1 Hz, 2H), 4.33 – 4.39 (m, 1H), 6.37 (s, 2H), 6.90 – 6.95 (m, 3H), 7.25 – 7.29 (m, 2H), 7.55 (s, 1H), 7.57 (s, 1H).

^13^C-NMR:	(DEPTQ, 151 MHz, DMSO-d_6_) *δ* 13.9, 21.5, 24.7, 37.5, 66.4, 68.8, 101.8, 113.8, 114.4, 119.7, 120.5, 121.1, 129.4, 138.8, 152.7, 156.4, 158.2, 171.7.

HPLC:	system 3: λ = 220 nm*, t*_R_ = 24.5 min, purity: >99%.

**1-(3-Amino-6-fluoro-1*H*-indazol-1-yl)-3-phenoxypropan-1-one (KD2, 10a)**

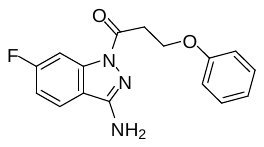

To a solution of 3-phenoxypropionic acid (**38**, 52.8 mg, 0.32 mmol) and HOAt (46.1 mg, 0.34 mmol) in DMF (1 mL) was added EDC × HCl (61.6 mg, 0.32 mmol). After stirring for 10 min at rt, a solution of 6-fluoro-1*H*-indazol-3-amine (**13**, 40.3 mg, 0.27 mmol) in DMF (0.5 mL) was added. After stirring the reaction mixture for 3 h, the solvent was removed by lyophilization and the residue was recrystallized from methanol. The mother liquor was purified by preparative HPLC using a solvent system of CH_3_OH / 0.1% aq. HCOOH, a flow rate of 10 mL / min and a gradient of 50% to 80% CH_3_OH in 15 min, 80% to 95% CH_3_OH in 2 min, 95% CH_3_OH for 4 min (*t*_R_: 20.4 min, λ = 254 nm). Combining both fractions afforded **KD2** (**10a**, 21.4 mg, 0.07 mmol, 27%) as a white solid.

ESI-MS:	*m/z* 300.1 [M+H]^+^.

HR-ESI-MS:	*m/z* [M+H]^+^ calcd. 300.1143 for C_18_H_17_F_3_N_3_O_2_, found 300.1147.

IR (KBr):	3406, 3334, 3209, 1685 cm^-1^.

m.p.:	181 °C.

^1^H-NMR:	(600 MHz, DMSO-d_6_) *δ* 3.43 (t, *J* = 6.0 Hz, 2H), 4.38 (t, *J* = 6.0 Hz, 2H), 6.59 (s, 2H), 6.92 – 6.97 (m, 3H), 7.25 (ddd, *J* = 9.1, 8.6, 2.4 Hz, 1H), 7.28 – 7.31 (m, 2H), 7.82 – 7.86 (m, 2H).

^13^C-NMR:	(DEPTQ, 151 MHz, DMSO-d_6_) *δ* 34.3, 62.7, 101.5 (d, *J* = 30 Hz), 112.1 (d, *J* = 25 Hz), 114.4, 116.9, 120.6, 122.6 (d, *J* = 11 Hz), 129.4, 139.6 (d, *J* = 12 Hz), 152.5, 158.2, 163.1 (d, *J* = 245 Hz), 169.0.

HPLC:	system 3: λ = 220 nm*,* *t*_R_ = 23.6 min, purity: >99%.

**1-(3-Amino-6-fluoro-1*H*-indazol-1-yl)-2-(phenoxymethyl)butan-1-one (KD6, 10b)**

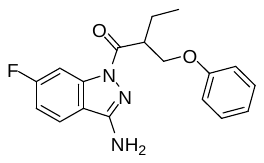

To a solution of commercially available 2-(phenoxymethyl)butanoic acid (**39**, 61.9 mg, 0.32 mmol) and HOAt (43.2 mg, 0.32 mmol) in DMF (1 mL) was added EDC × HCl (61.0 mg, 0.32 mmol). After stirring for 10 min at rt, a solution of 6-fluoro-1*H*-indazol-3-amine (**13**, 40.6 mg, 0.27 mmol) in DMF (0.5 mL) was added. After stirring the reaction mixture for 3 h, the solvent was removed by lyophilization and the residue was purified by preparative HPLC using a solvent system of CH_3_OH / 0.1% aq. HCOOH, a flow rate of 10 mL / min and a gradient of 50% to 80% CH_3_OH in 15 min, 80% to 95% CH_3_OH in 2 min, 95% CH_3_OH for 4 min (*t*_R_: 19.7 min, λ = 254 nm) to afford **KD6** (**10b**, (45.5 mg, 0.14 mol, 45%) as a white solid. The enantiomers were separated by performing chiral preparative HPLC (isocratic solvent system: *n*-hexane / isopropanol, 98:2) to afford the two enantiomers **KD6en1 (10b-ent1)** (*t*_R_: 16.5 min) and **KD6en2 (10b-ent2)** (*t*_R_: 25.1 min).

ESI-MS:	*m/z* 328.2 [M+H]^+^.

HR-ESI-MS:	*m/z* [M+H]^+^ calcd. 328.1456 for C_18_H_19_FN_3_O_2_, found 328.1456.

IR (KBr):	3437, 3340, 3231, 1688, 1642, 1624 cm^-1^.

m.p.:	97 °C.

^1^H-NMR:	(600 MHz, DMSO-d_6_) *δ* 0.93 (dd, *J* = 7.7, 7.3 Hz, 3 H), 1.77 (ddq, *J* = 14.5, 8.5, 7.3 Hz, 1H), 1.82 (ddq, *J* = 14.5, 7.7, 5.5 Hz, 1H), 4.05 (dddd, *J* = 8.5, 8.0, 5.8, 5.5 Hz, 1H), 4.13 (dd, *J* = 9.3, 5.5 Hz, 1H), 4.37 (dd, *J* = 9.3, 8.0 Hz, 1H), 6.60 (s, 2H), 6.93 – 6.91 (m, 3H), 7.28 – 7.24 (m, 3H), 7.95 (dd, *J* = 8.6, 5.3, 1H), 7.98 (dd, *J* = 9.8, 2.3 Hz).

^13^C-NMR:	(DEPTQ, 151 MHz, DMSO-d_6_) *δ* 11.2, 21.7, 44.0, 67.7, 101.8 (d, *J* = 28 Hz), 112.2 (d, *J* = 24 Hz), 114.4, 117.0, 120.6, 122.6 (d, *J* = 11 Hz), 129.4, 139.6 (d, *J* = 19 Hz), 152.5, 158.2, 163.1 (d, *J* = 245 Hz), 171.9.

HPLC:	system 5: λ = 220 nm*,* for both enantiomers *t*_R_ = 25.9 min, **KD6en1 (10b-ent1)**: purity: >99%, **KD6en2 (10b-ent2)**: purity: >99%.

chiral HPLC:	isocratic elution with *n*-hexane / isopropanol, 98:2: **KD6en1 (10b-ent1)**: *t*_R_ = 17.2 min, >99% *ee* (254 nm); **KD6en2 (10b-ent2)**: *t*_R_ = 23.2 min, 98% *ee* (254 nm).

[*α*]*_D_^25^*:	**KD6en1 (10b-ent1)**: +23.6 (*c* = 0.4, methanol); **KD6en2 (10b-ent2)**: -23.6 (*c* = 0.2, methanol).

**1-(3-Amino-6-fluoro-1*H*-indazol-1-yl)-2-methoxy-3-phenoxypropan-1-one (KD5, 10c)**

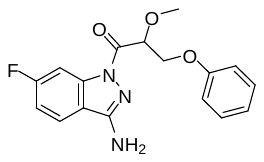

To a solution of commercially available 2-methoxy-3-phenoxypropanoic acid (**40**, 62.4 mg, 0.32 mmol) and HOAt (43.4 mg, 0.32 mmol) in DMF (1 mL) was added EDC × HCl (61.4 mg, 0.32 mmol). After stirring for 10 min at rt, a solution of 6-fluoro-1*H*-indazol-3-amine (**13**, 40.3 mg, 0.27 mmol) in DMF (0.5 mL) was added. After stirring the reaction mixture for 2 h, the solvent was removed by lyophilization and the residue was purified by preparative HPLC using a solvent system of CH_3_OH / 0.1% aq. HCOOH, a flow rate of 10 mL / min and a gradient of 50% to 80% CH_3_OH in 15 min, 80% to 95% CH_3_OH in 2 min (*t*_R_: 15.3 min, λ = 254 nm) to afford **KD5** (**10c**, 37.6 mg, 0.11 mmol, 43%) as a white solid. The enantiomers were separated by 2× performing chiral preparative HPLC (isocratic solvent system: *n*-hexane / isopropanol, 93:7) to afford the two enantiomers **KD5en1 (10c-ent1)** (*t*_R_: 32.6 min) and **KD5en2 (10c-ent2)** (*t*_R_: 35.1 min).

ESI-MS:	*m/z* 330.2 [M+H]^+^.

HR-ESI-MS:	*m/z* [M+H]^+^ calcd. 330.1248 for C_17_H_17_FN_3_O_3_, found 330.1249.

IR (KBr):	3457, 3349, 3227, 1695, 1632 cm^-1^.

m.p.:	134 °C.

^1^H-NMR:	(600 MHz, DMSO-d_6_) *δ* 3.41 (s, 3H), 4.26 (dd, *J* = 10.7, 6.9 Hz), 4.41 (dd, *J* = 10.7, 2.9 Hz, 1H), 5.21 (dd, *J* = 6.9, 2.9 Hz, 1H), 6.70 (s, 2H), 6.91 – 6.98 (m, 3H), 7.25 – 7.31 (m, 3H), 7.94 – 7.99 (m, 2H).

^13^C-NMR:	(DEPTQ, 151 MHz, DMSO-d_6_) *δ* 57.6, 67.8, 78.1, 101.6 (d, *J* = 28 Hz), 112.5 (d, *J* = 25 Hz), 114.4, 116.8, 122.7 (d, *J* = 12 Hz), 129.4, 139.8 (d, *J* = 12 Hz), 152.9, 158.0, 163.2 (d, *J* = 246 Hz), 166.8.

HPLC:	system 3: λ = 220 nm*,* for both enantiomers *t*_R_ = 22.5 min, **KD5en1 (10c-ent1)**: purity: 97%, **KD5en2 (10c-ent2)**: purity: >99%.

chiral HPLC:	isocratic elution with *n*-hexane / isopropanol, 93:7: **KD5en1 (10c-ent1)**: *t*_R_ = 29.6 min, >99% *ee* (254 nm); **KD5en2 (10c-ent2)**: *t*_R_ = 33.7 min, 91% *ee* (254 nm).

[*α*]*_D_^25^*:	**KD5en1 (10c-ent1)**: +33.6 (*c* = 0.9, methanol); **KD5en2 (10c-ent2)**: n.d.

**1-(3-Amino-6-fluoro-1*H*-indazol-1-yl)-2-methyl-4-phenylbutan-1-one (KD3, 10d)**

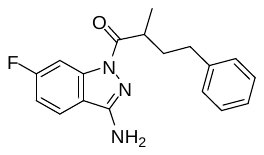

To a solution of 2-methyl-4-phenylbutyric acid (**41**, 56.7 mg, 0.32 mmol) and HOAt (43.8 mg, 0.32 mmol) in DMF (1 mL) was added EDC x HCl (61.1 mg, 0.32 mmol). After stirring for 10 min at rt, a solution of 6-fluoro-1*H*-indazol-3-amine (**13**, 40.2 mg, 0.27 mmol) in DMF (0.5 mL) was added. After stirring the reaction mixture for 1 h, the solvent was removed by lyophilization and the residue was purified by preparative HPLC (Nucleodur C18 HTec, 32 mm × 250 mm, 5 µm) using a solvent system of CH_3_OH / 0.1% aq. HCOOH, a flow rate of 25 mL / min and a gradient of 50% to 80% CH_3_OH in 15 min, 80% to 95% CH_3_OH in 2 min, 95% CH_3_OH for 7 min (*t*_R_: 23.0 min, λ = 254 nm) to afford **KD3** (**10d**, 25.1 mg, 0.08 mmol, 30%) as a colorless oil. The enantiomers were separated by 2× performing chiral preparative HPLC (isocratic solvent system: *n*‑hexane / isopropanol, 98:2) to afford the two enantiomers **KD3en1 (10d-ent1)** (*t*_R_: 14.7 min) and **KD3en2 (10d-ent2)** (*t*_R_: 17.2 min).

ESI-MS:	*m/z* 312.1 [M+H]^+^.

HR-ESI-MS:	*m/z* [M+H]^+^ calcd. 312.1507 for C_18_H_19_FN_3_O, found 312.1509.

IR (KBr):	3459, 3350, 3227, 1678, 1626 cm^-1^.

^1^H-NMR:	(600 MHz, DMSO-d_6_) *δ* 1.22 (d, *J* = 6.9 Hz, 3H), 1.76 (dddd, *J* = 13.3, 9.8, 6.5, 6.4 Hz, 1H), 2.07 (dddd, *J* = 13.3, 9.8, 7.6, 6.0 Hz, 1H), 2.57 (ddd, *J* = 13.6, 9.8, 6.0 Hz, 1H), 2.62 (ddd, *J* = 13.6, 9.8, 6.4 Hz, 1H), 3.64 (ddq, *J* = 7.6, 6.5, 6.9, 1H), 6.53 (s, 2H), 7.14 – 7.18 (m, 3H), 7.22 – 7.26 (m, 3H), 7.93 (dd, *J* = 8.7, 5.3, 1H), 7.97 (dd, *J* = 9.8, 2.3 Hz).

^13^C-NMR:	(DEPTQ, 151 MHz, DMSO-d_6_) *δ* 17.2, 32.7, 34.3, 36.7, 101.7 (d, *J* = 28 Hz), 111.9 (d, *J* = 24 Hz), 116.8, 122.5 (d, *J* = 12 Hz), 125.7, 128.1, 128.2, 139.8 (d, *J* = 15 Hz), 141.5, 152.3, 163.6 (d, *J* = 245 Hz), 174.7.

HPLC:	system 3: λ = 220 nm*,* for both enantiomers *t*_R_ = 26.0 min, **KD3 10d-ent1**: purity: 96%, **KD3en2 (10d-ent2)**: purity: 97%.

chiral HPLC:	isocratic elution with *n*-hexane / isopropanol, 98:2: **KD3en1 (10d-ent1)**: *t*_R_ = 14.3 min, >99% *ee* (254 nm); **KD3en2 (10d-ent2)**: *t*_R_ = 16.6 min, 99% *ee* (254 nm).

[*α*]*_D_^25^*:	**KD3en1 (10d-ent1)**: -13.9 (*c* = 0.9, methanol); **KD3en2 (10d-ent2)**: +14.6 (*c* = 0.1, methanol).

**1-(3-Amino-6-fluoro-1*H*-indazol-1-yl)-4-phenylbutan-1-one (KD4, 10e)**

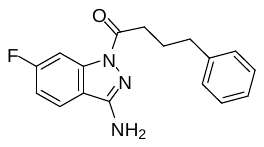

To a solution of sodium 4-phenylbutyrate (**42**, 52.5 mg, 0.32 mmol) and HOAt (43.2 mg, 0.32 mmol) in DMF (1 mL) was added EDC × HCl (61.0 mg, 0.32 mmol). After stirring for 10 min at rt, a solution of 6-fluoro-1*H*-indazol-3-amine (**13**, 40.7 mg, 0.27 mmol) in DMF (0.5 mL) was added. After stirring the reaction mixture for 1.5 h, the solvent was removed by lyophilization and the residue was purified by preparative HPLC (Nucleodur C18 HTec, 32 mm × 250 mm, 5 µm) using a solvent system of CH_3_OH / 0.1% aq. HCOOH, a flow rate of 25 mL / min and a gradient of 50% to 80% CH_3_OH in 15 min, 80% to 95% CH_3_OH in 2 min, 95% CH_3_OH for 7 min (*t*_R_: 22.8 min, λ = 254 nm) to afford **KD4** (**10a**, 33.9 mg, 0.11 mmol, 42%) as a white solid

ESI-MS:	*m/z* 298.1 [M+H]^+^.

HR-ESI-MS:	*m/z* [M+H]^+^ calcd. 298.1350 for C_18_H_17_F_3_N_3_O_2_, found 298.1353.

IR (KBr):	3429, 3341, 3233, 3197, 1674, 1658, 1644 cm^-1^.

m.p.:	82 °C.

^1^H-NMR:	(600 MHz, DMSO-d_6_) *δ* 1.99 (tt, *J* = 7.5, 7.5 Hz, 2H), 2.68 (t, *J* = 7.5 Hz, 2H), 2.95 (t, *J* = 7.5 Hz, 2H), 6.52 (s, 2H), 7.16 – 7.24 (m, 4H), 7.27 – 7.31 (m, 2H), 7.91 (dd, *J* = 8.7, 5.2 Hz, 1H), 7.93 (dd, *J* = 10.0, 2.5 Hz, 1H).

^13^C-NMR:	(DEPTQ, 151 MHz, DMSO-d_6_) *δ* 25.5, 33.5, 34.4, 101.4 (d, *J* = 27 Hz), 111.8 (d, *J* = 24 Hz), 116.7, 122.5 (d, *J* = 11 Hz), 125.7, 128.2, 139.6 (d, *J* = 13 Hz), 141.5, 152.2, 158.2, 163.1 (d, *J* = 249 Hz), 171.4.

HPLC:	system 3: λ = 220 nm*,* *t*_R_ = 25.1 min, purity: 97%.

**^1^H and ^13^C NMR Spectra of the library compounds Z2075279358 - Z995908944**

Compound **Z2075279358**, ^1^H-NMR, DMSO-d_6_, 600 MHz

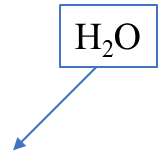

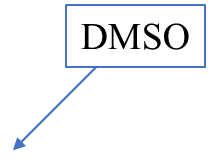

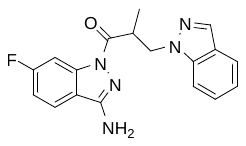

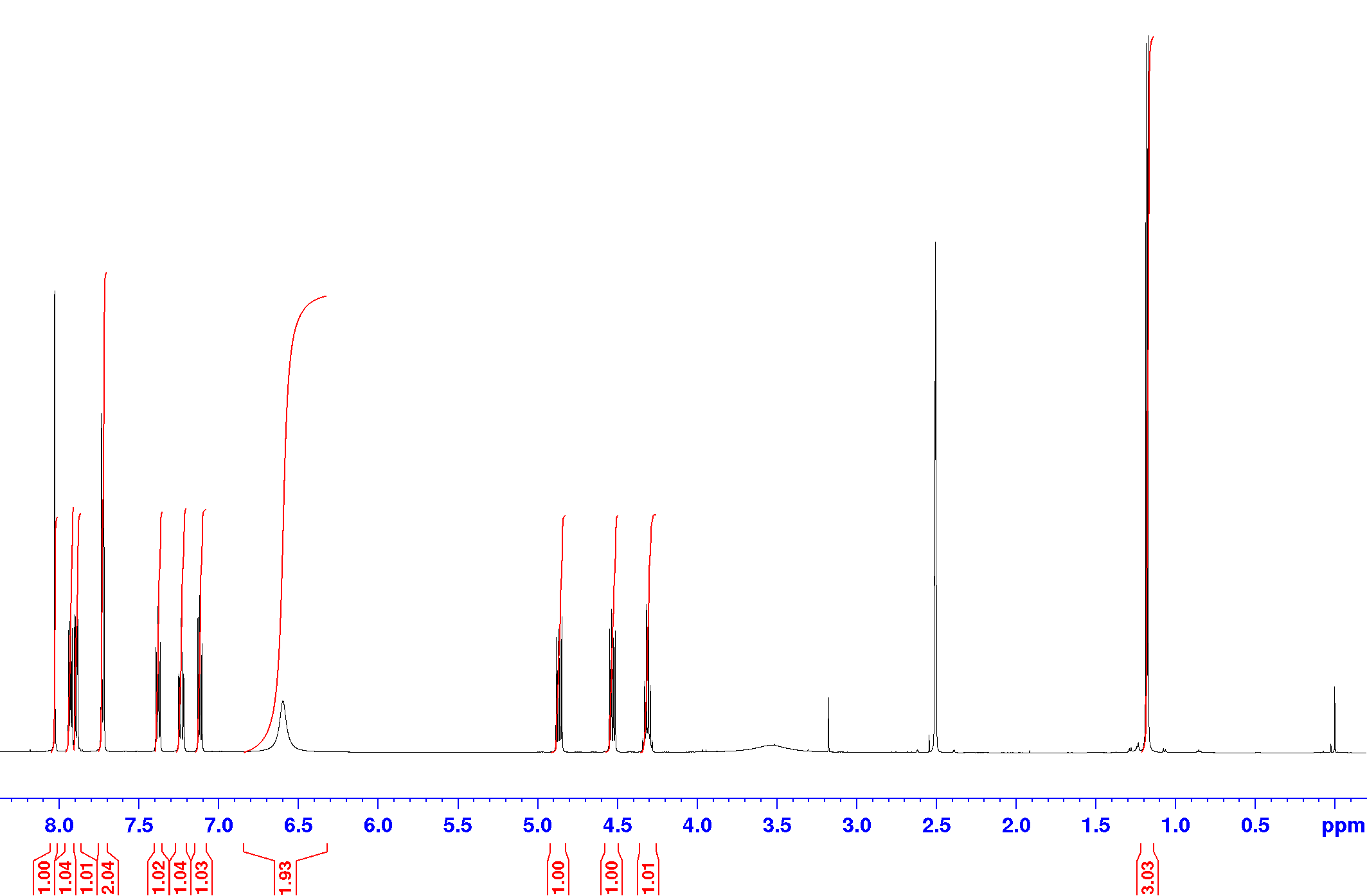

Compound **Z2075279358**, ^13^C-NMR (DEPTQ), DMSO-d_6_, 151 MHz

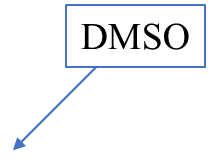

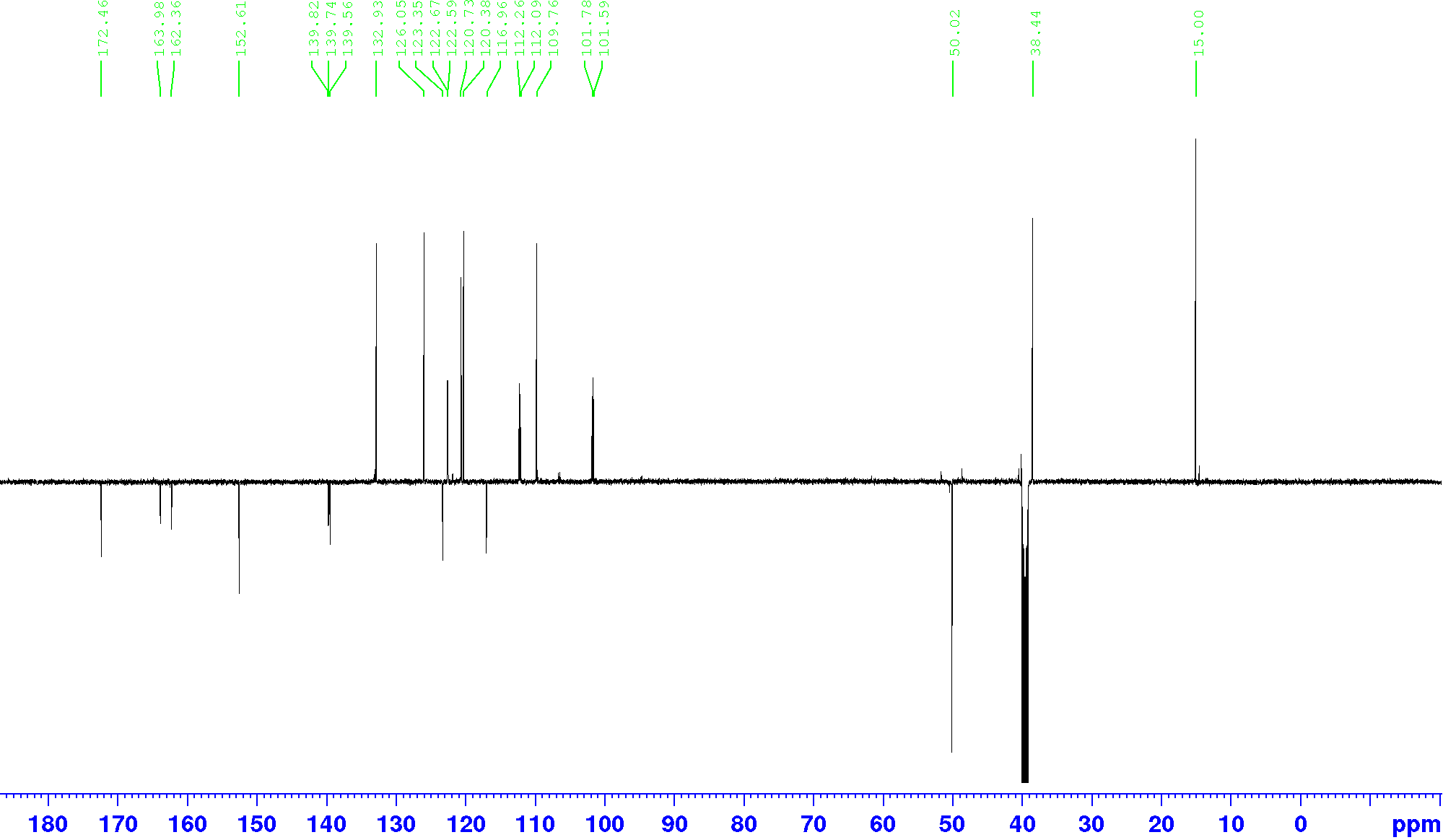

Compound **Z2194302854**, ^1^H-NMR, DMSO-d_6_, 600 MHz

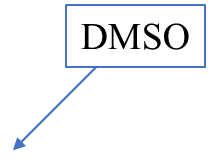

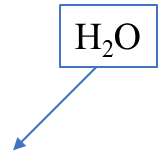

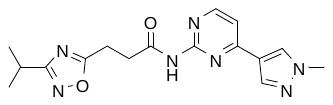

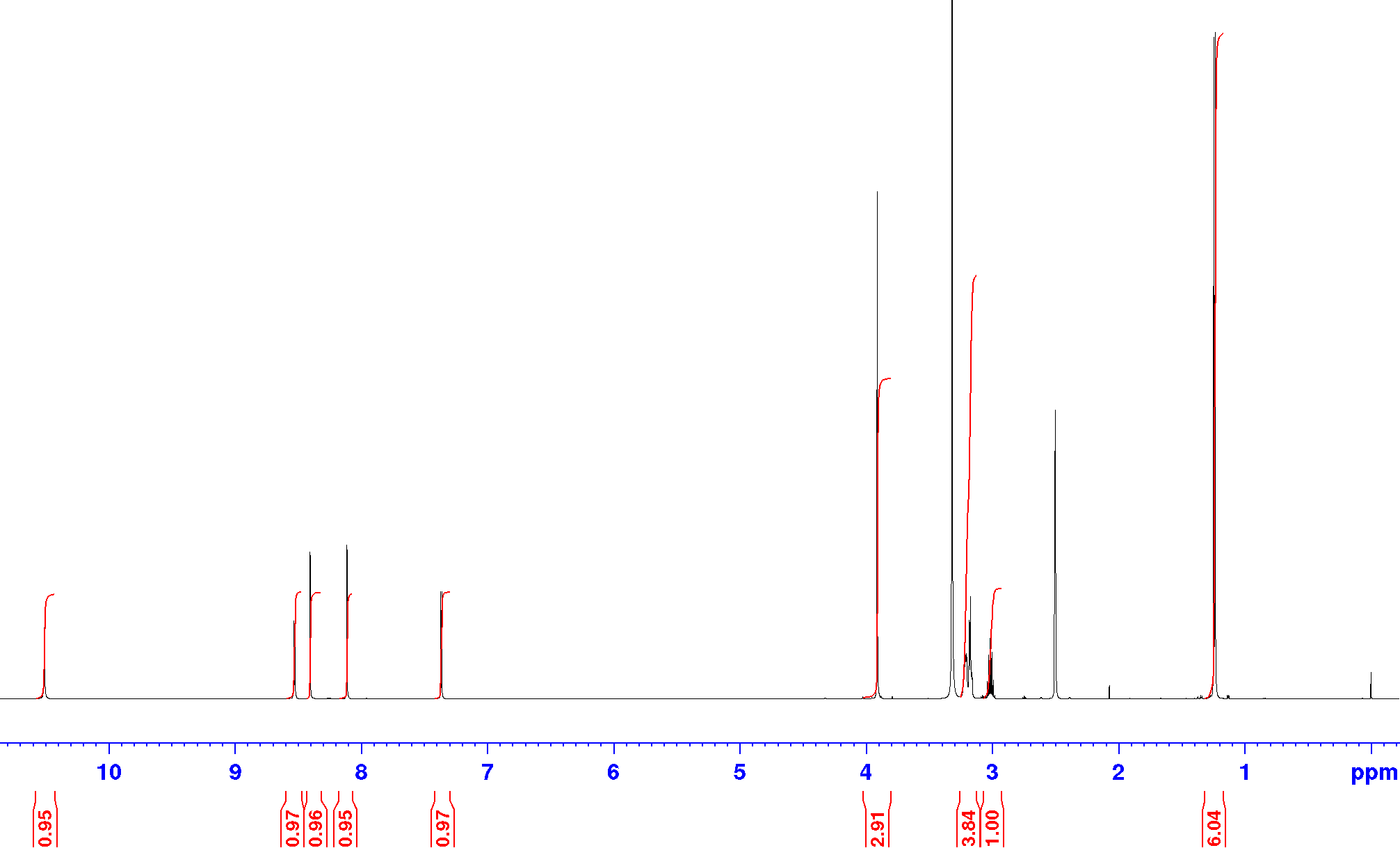

Compound **Z2194302854**, ^13^C-NMR (DEPTQ), DMSO-d_6_, 151 MHz

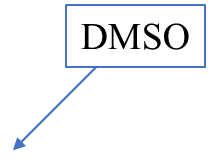

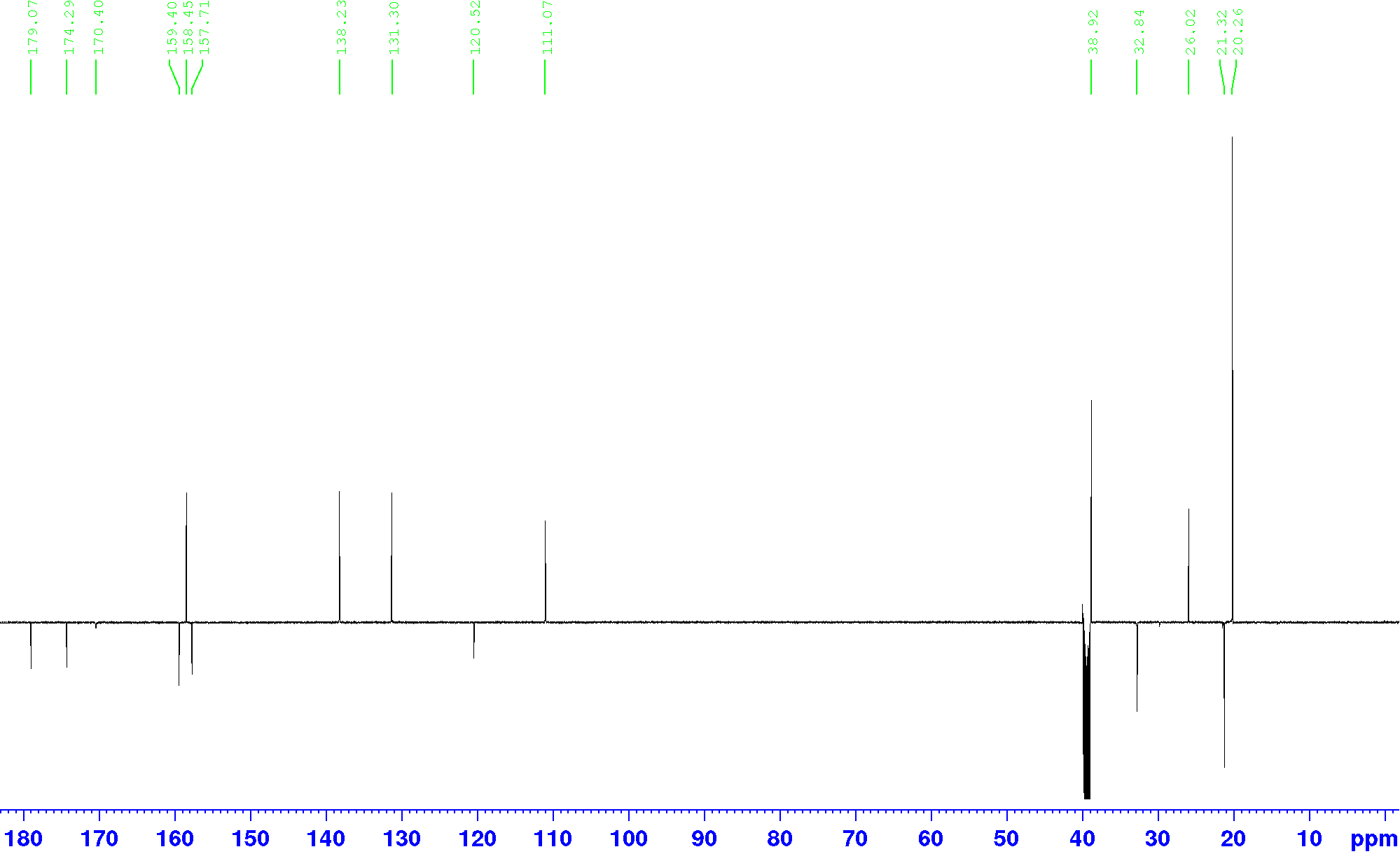

Compound **Z1343848401**, ^1^H-NMR, DMSO-d_6_, 600 MHz

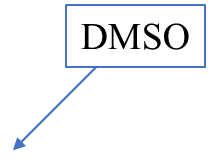

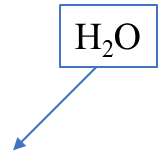

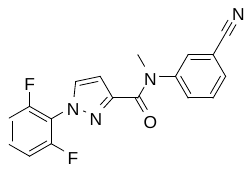

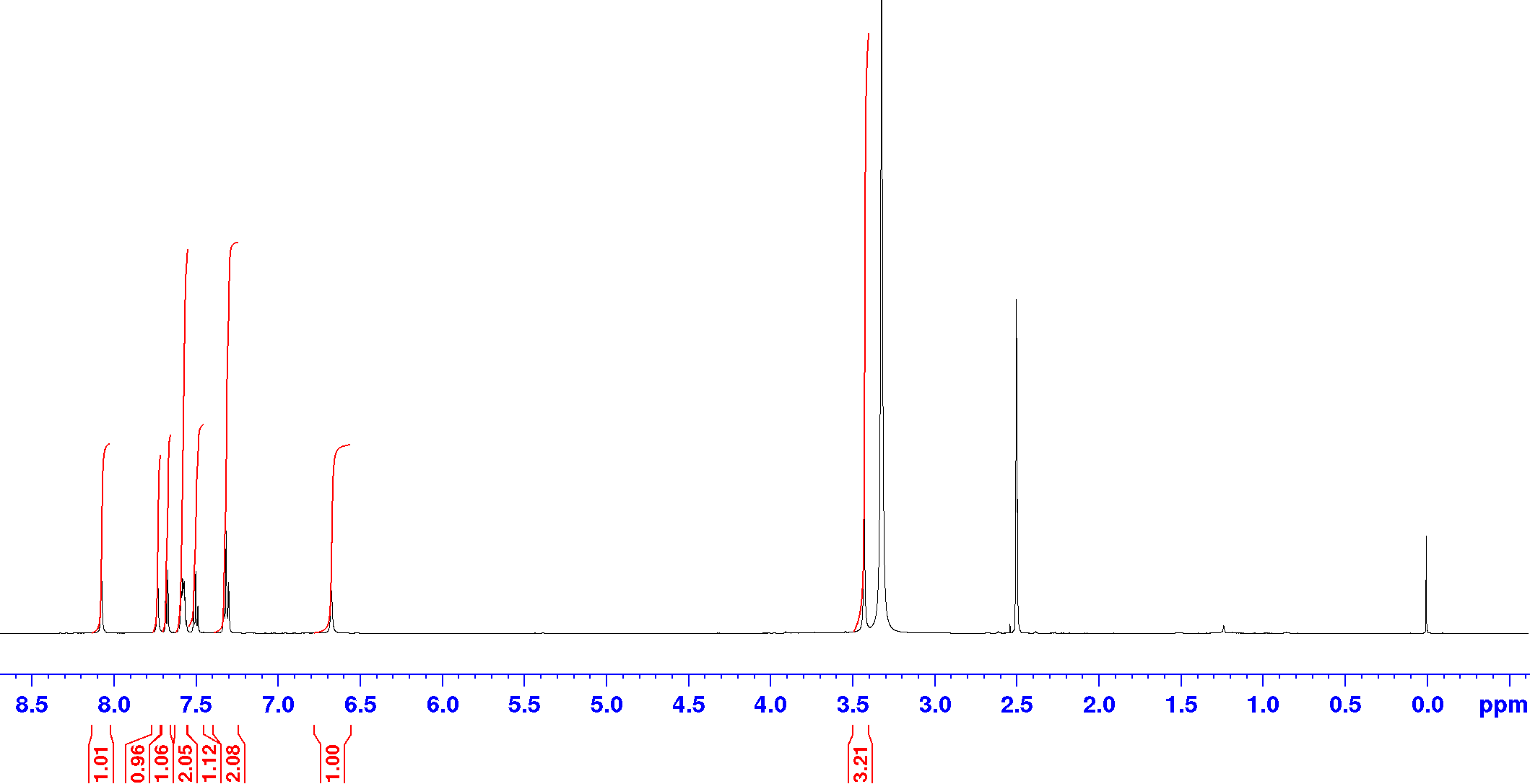

Compound **Z1343848401**, ^13^C-NMR (DEPTQ), DMSO-d_6_, 151 MHz

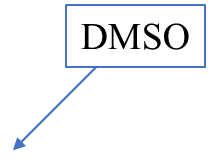

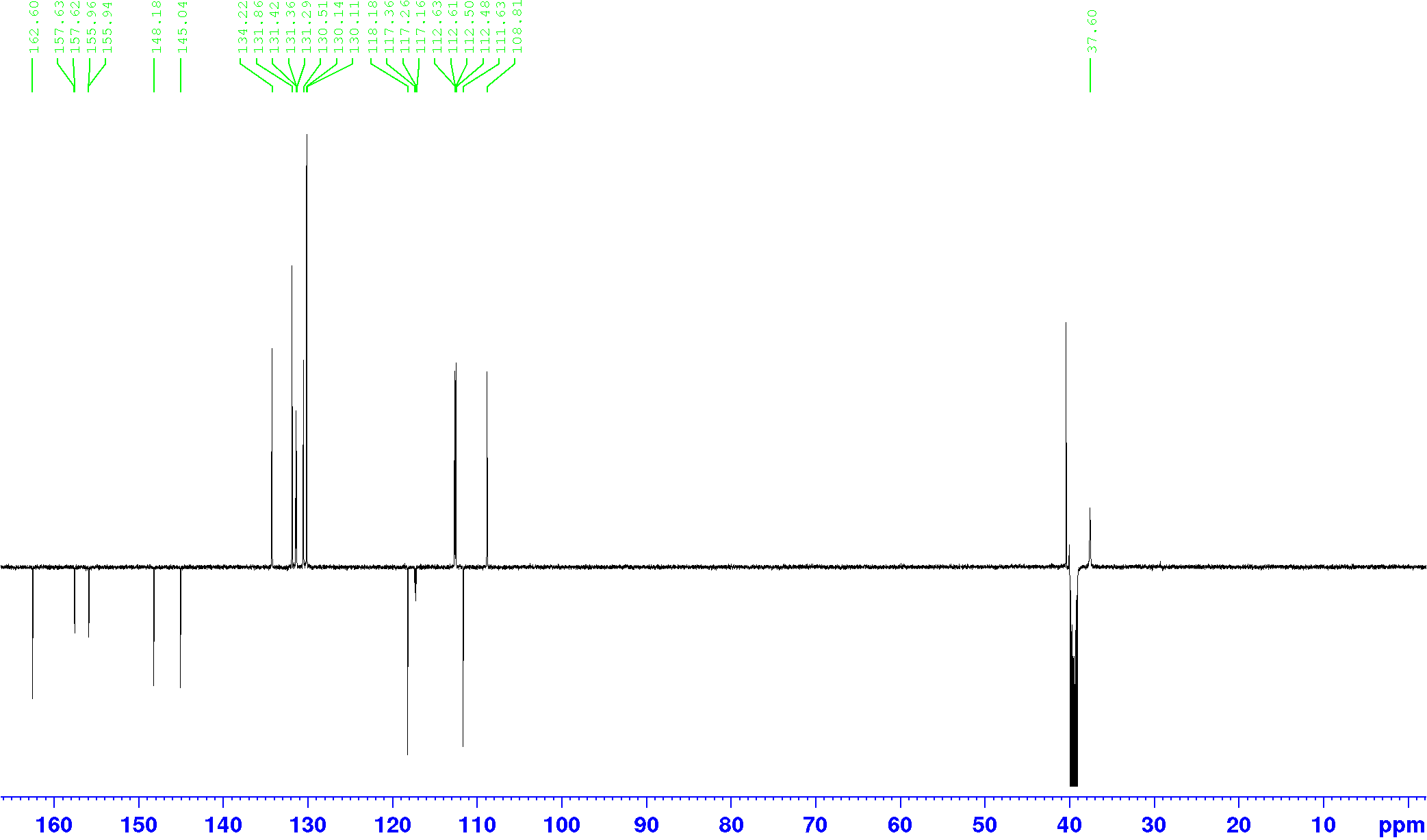

Compound **Z277641998**, ^1^H-NMR, DMSO-d_6_, 600 MHz

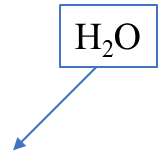

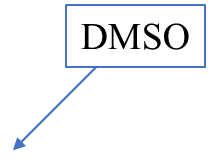

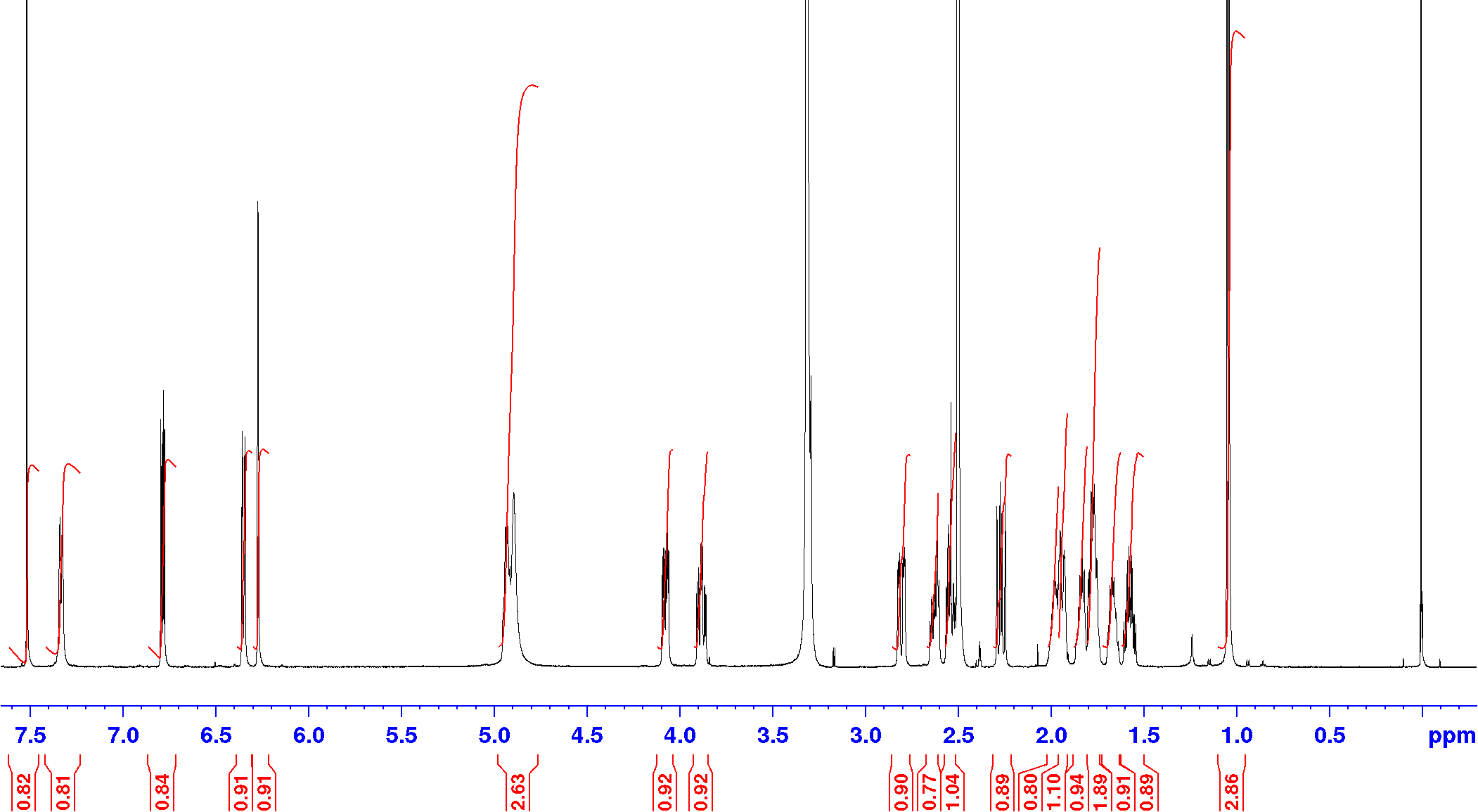

Compound **Z277641998**, ^13^C-NMR (DEPTQ), DMSO-d_6_, 151 MHz

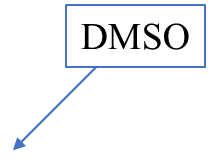

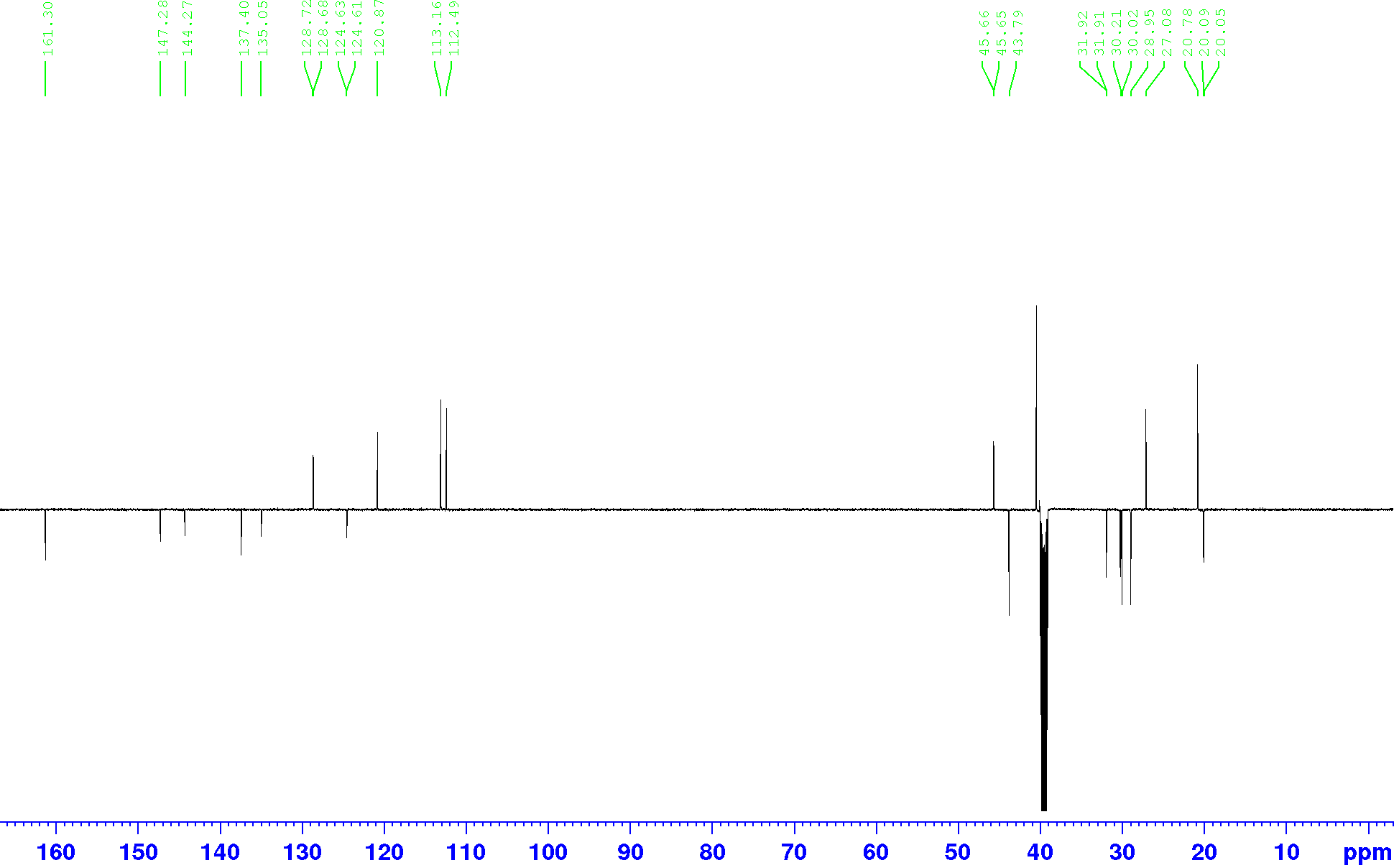

Compound **Z1224795288**, ^1^H-NMR, DMSO-d_6_, 600 MHz

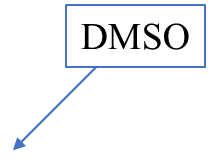

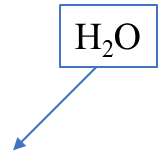

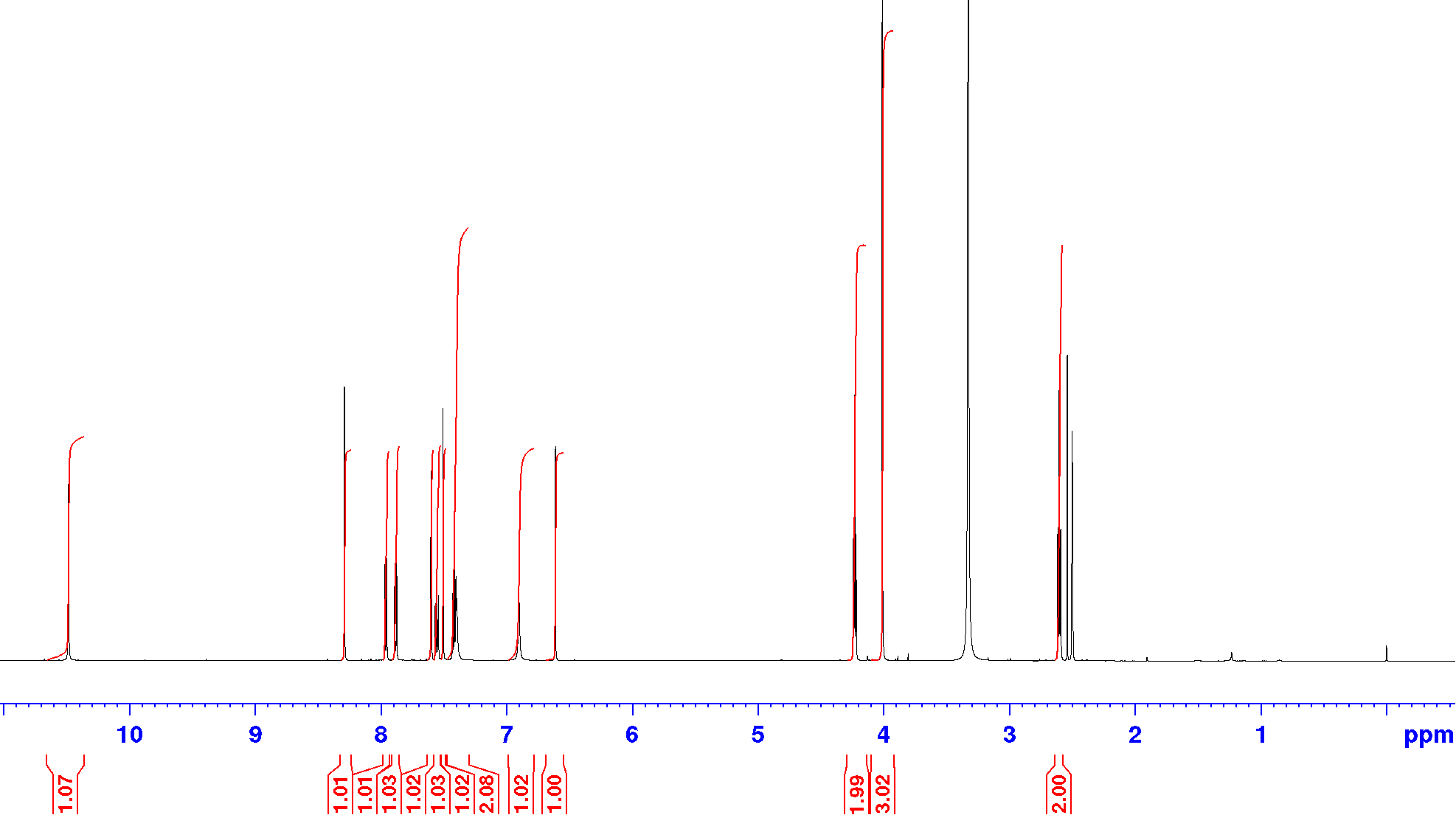

Compound **Z1224795288**, ^13^C-NMR (DEPTQ), DMSO-d_6_, 151 MHz

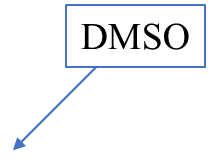

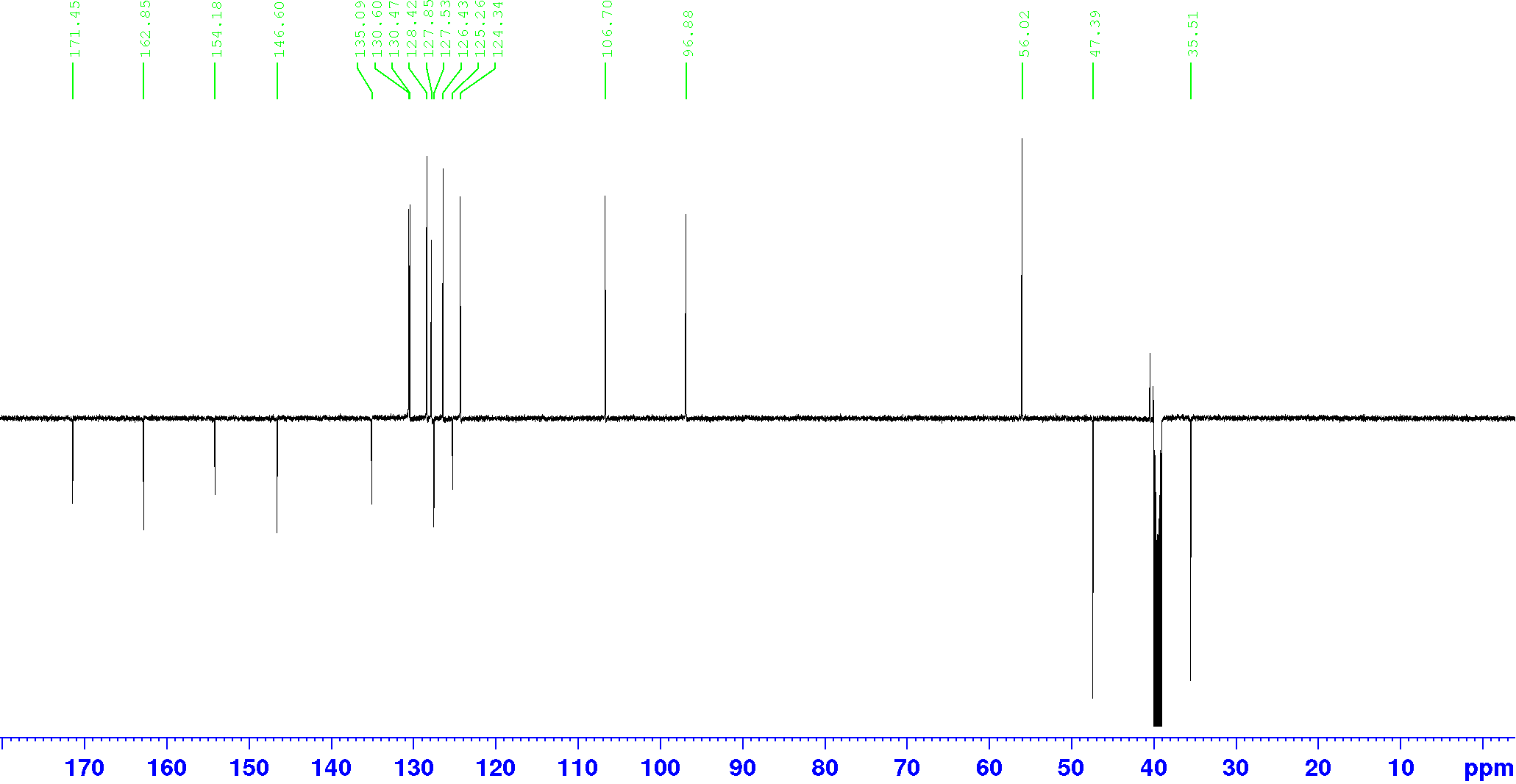

Compound **Z19702639**, ^1^H-NMR, DMSO-d_6_, 600 MHz

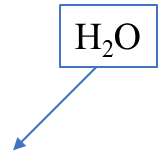

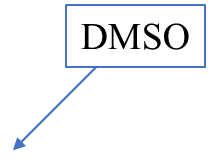

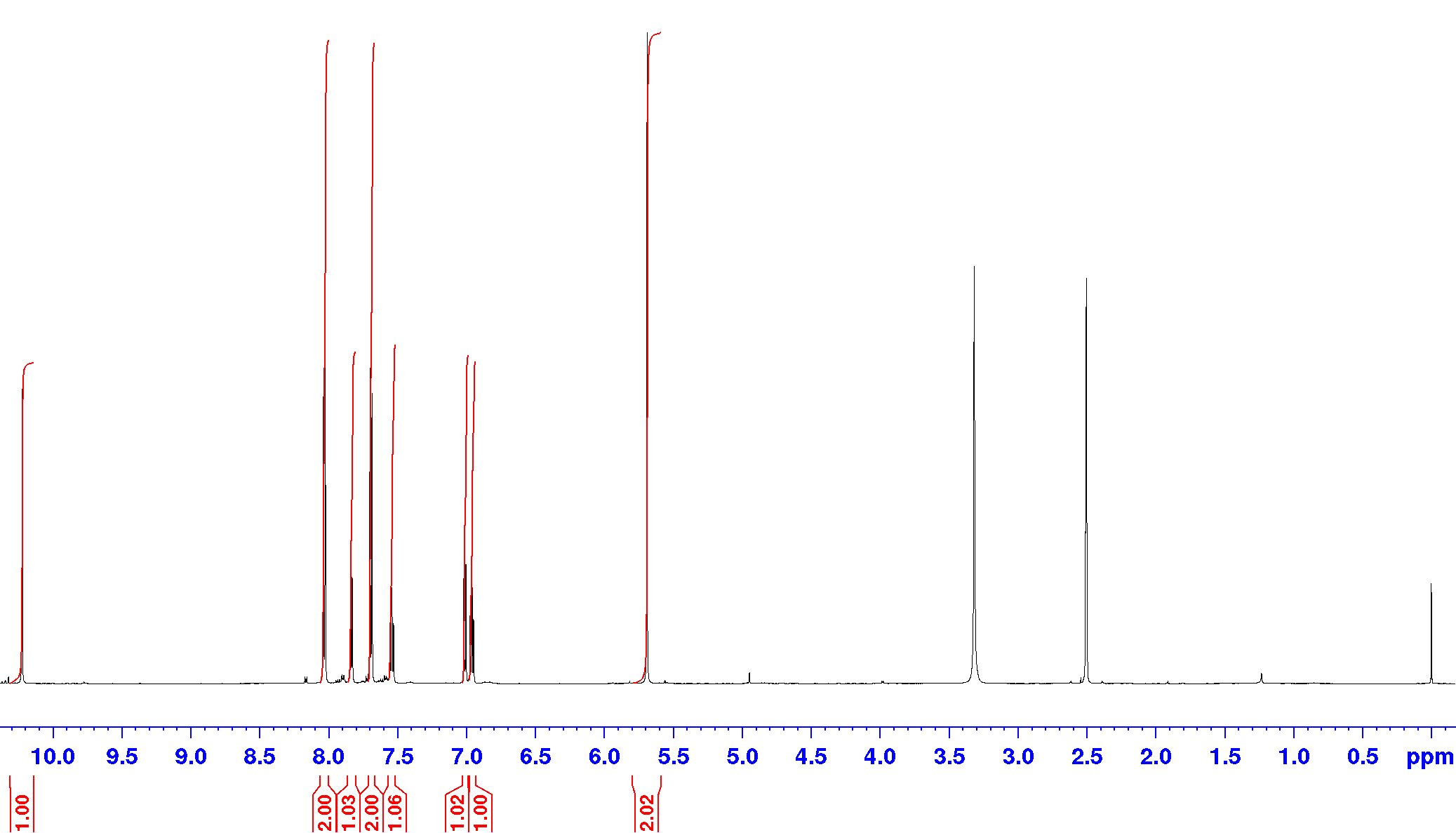

Compound **Z19702639**, ^13^C-NMR (DEPTQ), DMSO-d_6_, 151 MHz

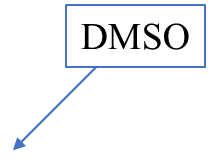

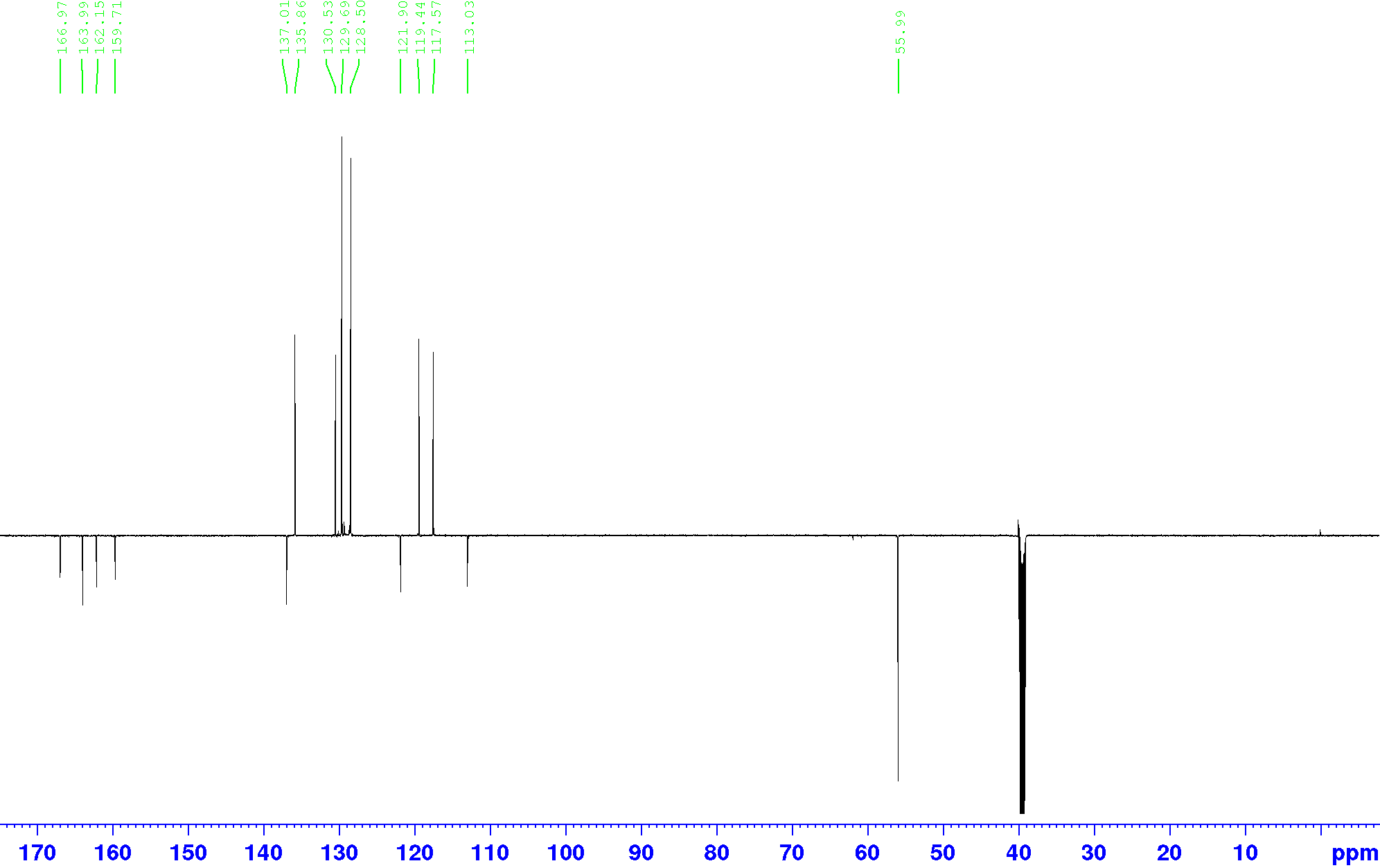

Compound **Z1096199008**, ^1^H-NMR, DMSO-d_6_, 600 MHz

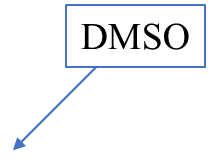

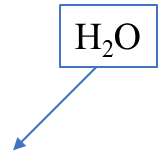

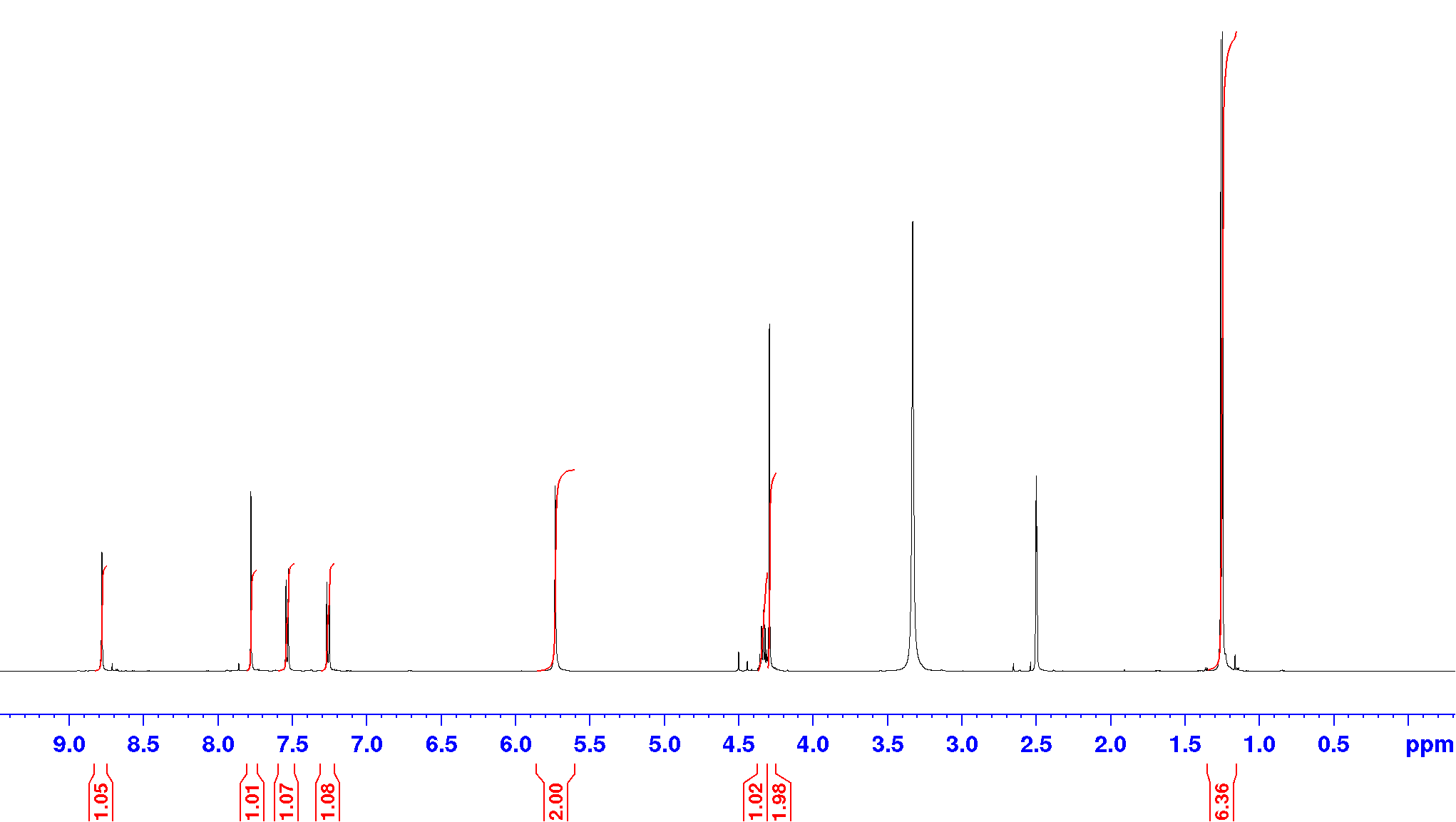

Compound **Z1096199008**, ^13^C-NMR (DEPTQ), DMSO-d_6_, 151 MHz

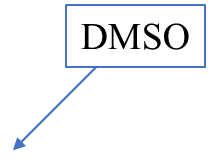

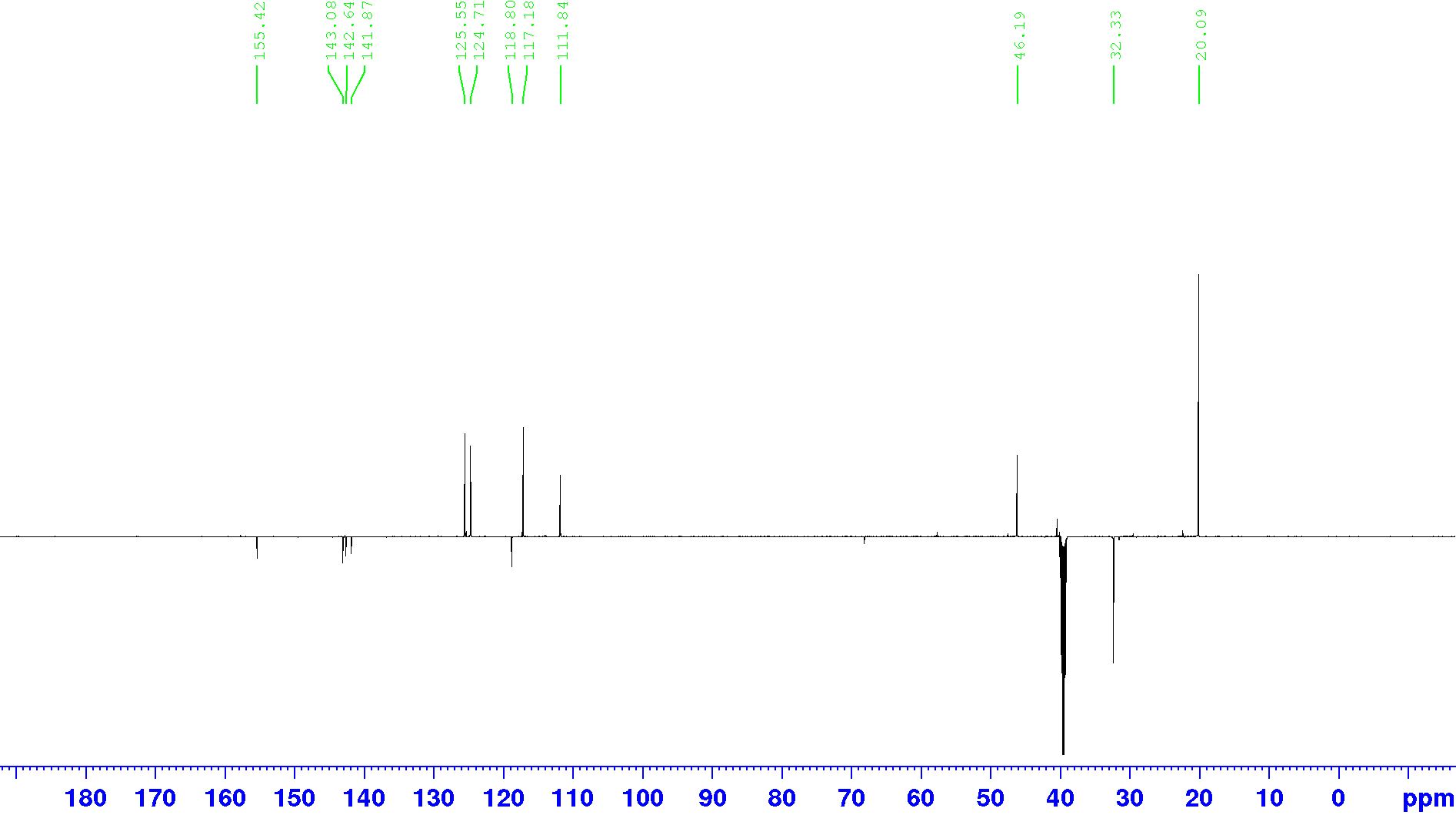

Compound **Z873519648**, ^1^H-NMR, DMSO-d_6_, 600 MHz

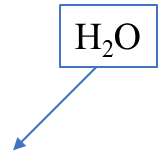

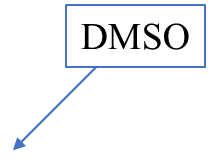

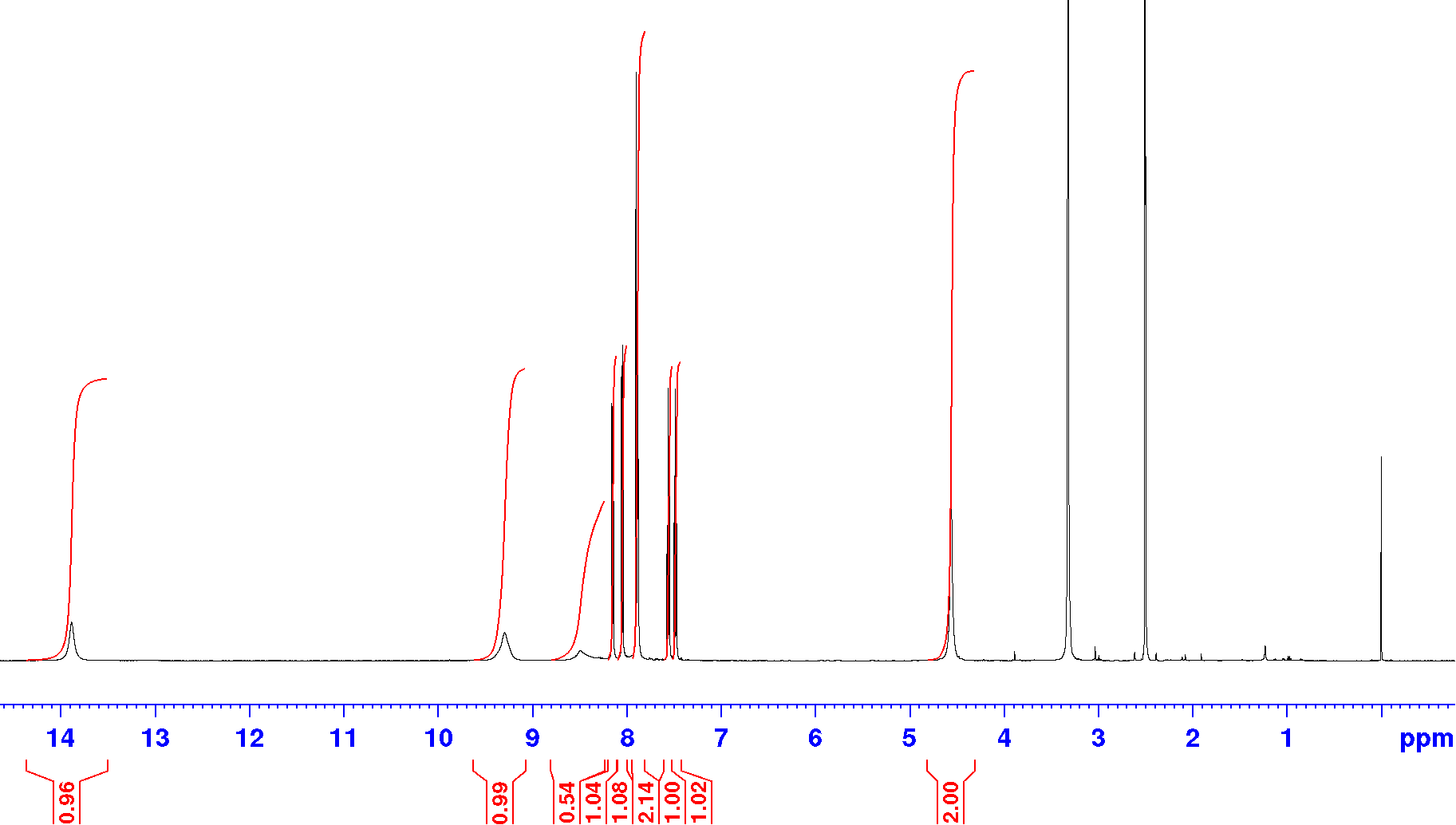

Compound **Z873519648**, ^13^C-NMR (DEPTQ), DMSO-d_6_, 151 MHz

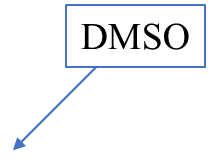

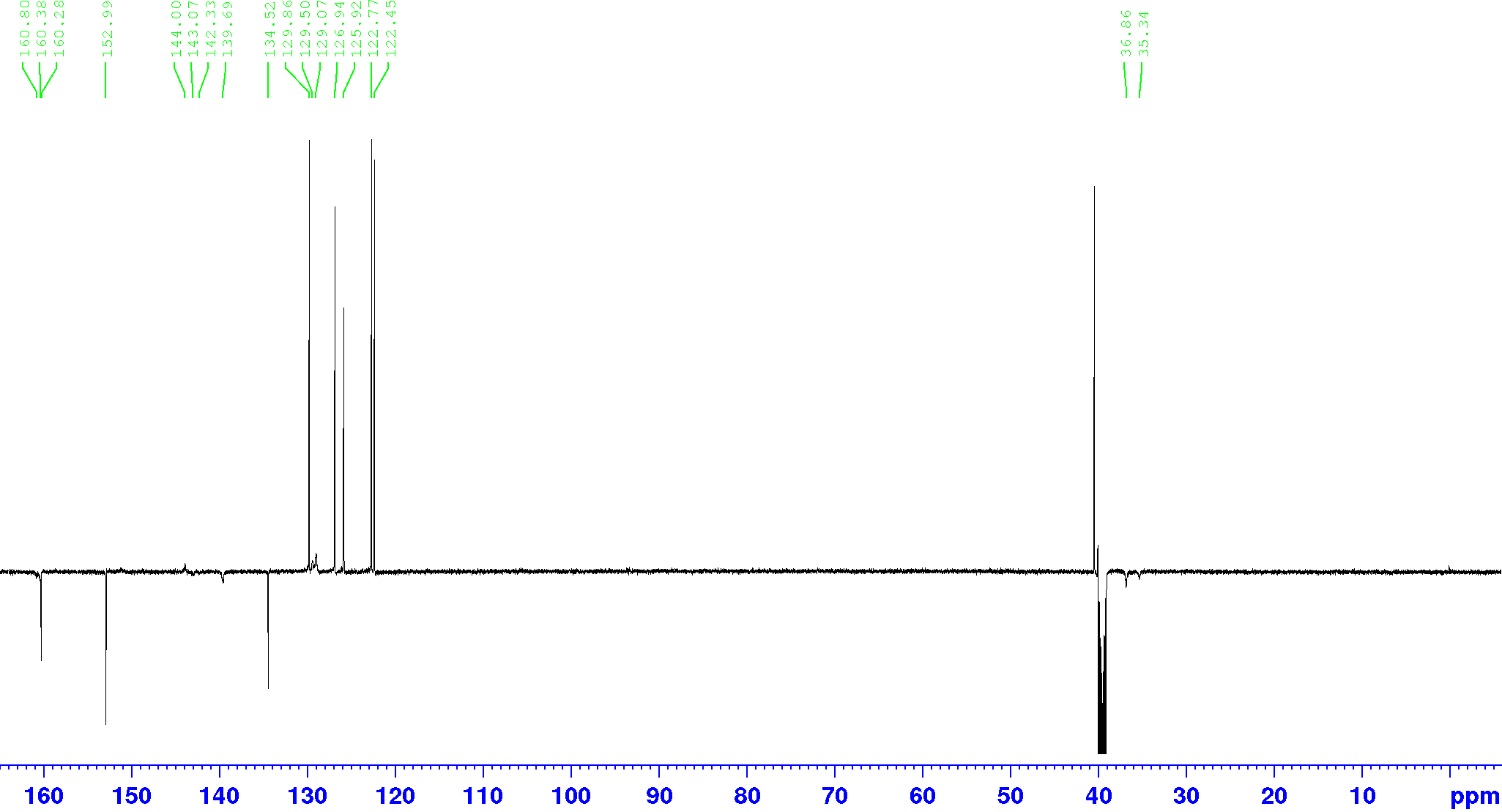

Compound **Z1783799713**, ^1^H-NMR, DMSO-d_6_, 600 MHz

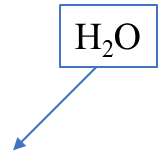

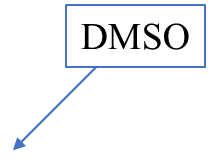

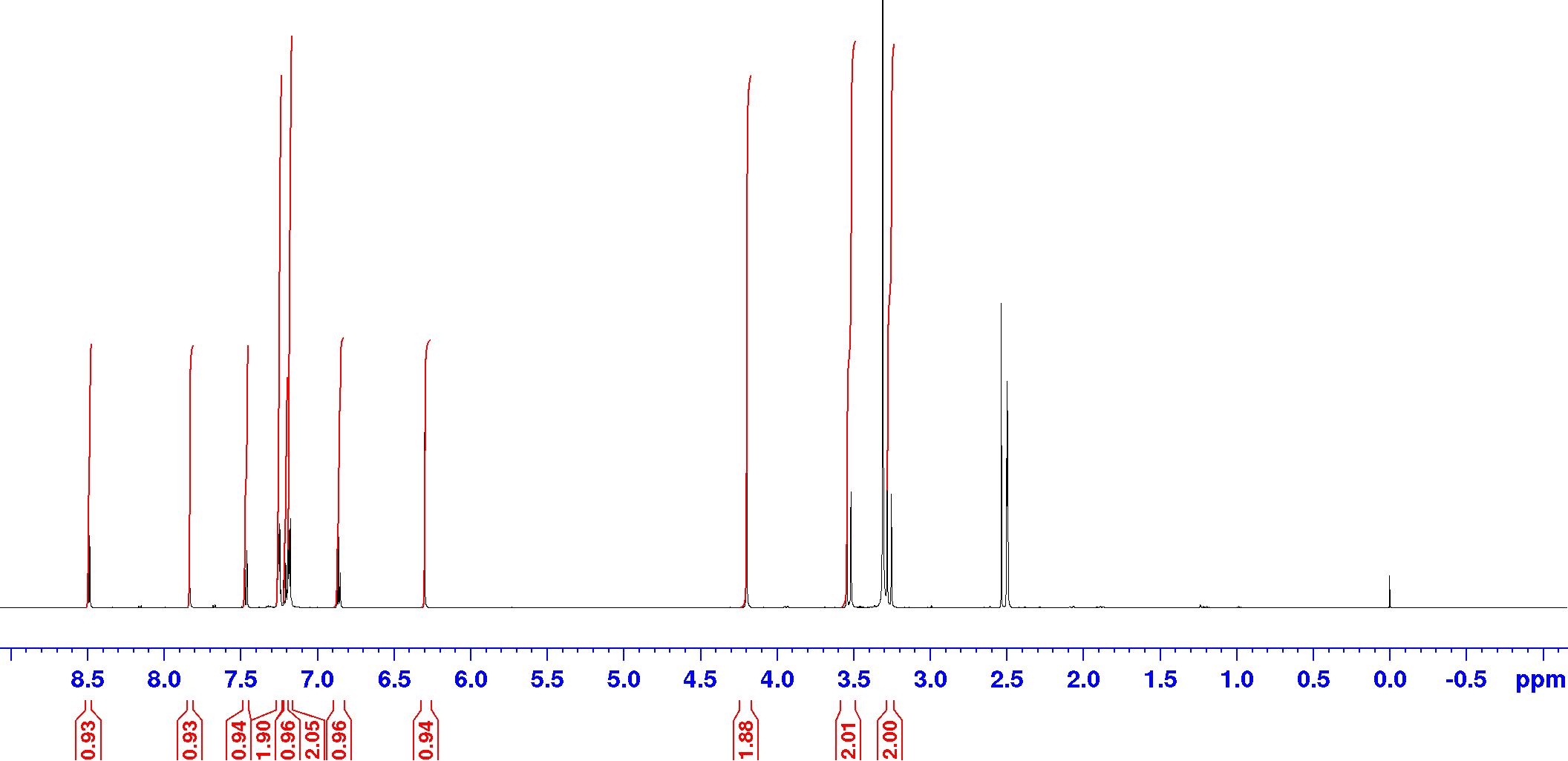

Compound **Z1783799713**, ^13^C-NMR (DEPTQ), DMSO-d_6_, 151 MHz

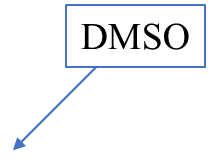

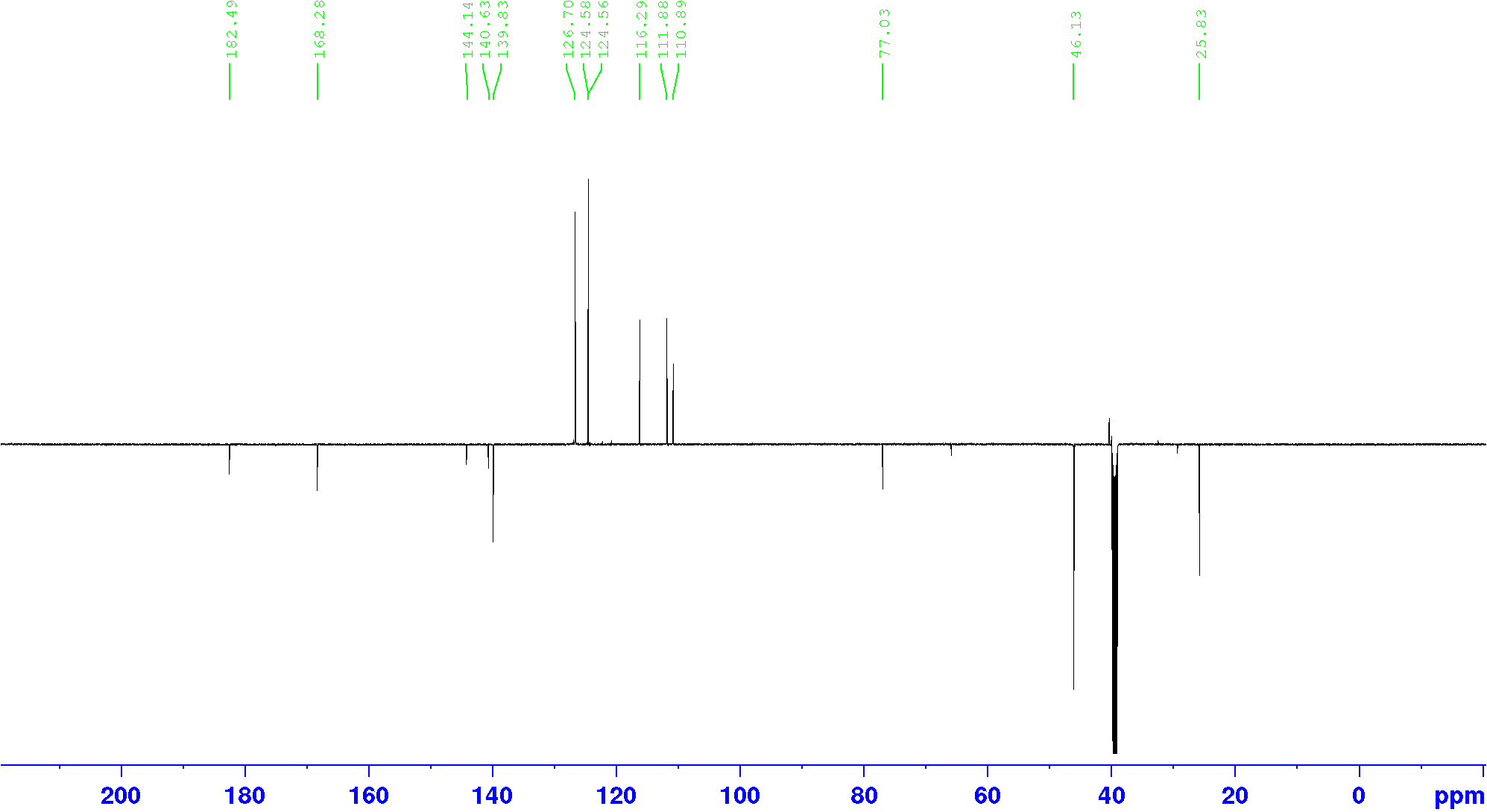

Compound **Z899051432**, ^1^H-NMR, DMSO-d_6_, 600 MHz

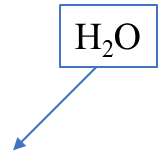

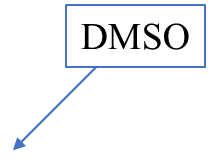

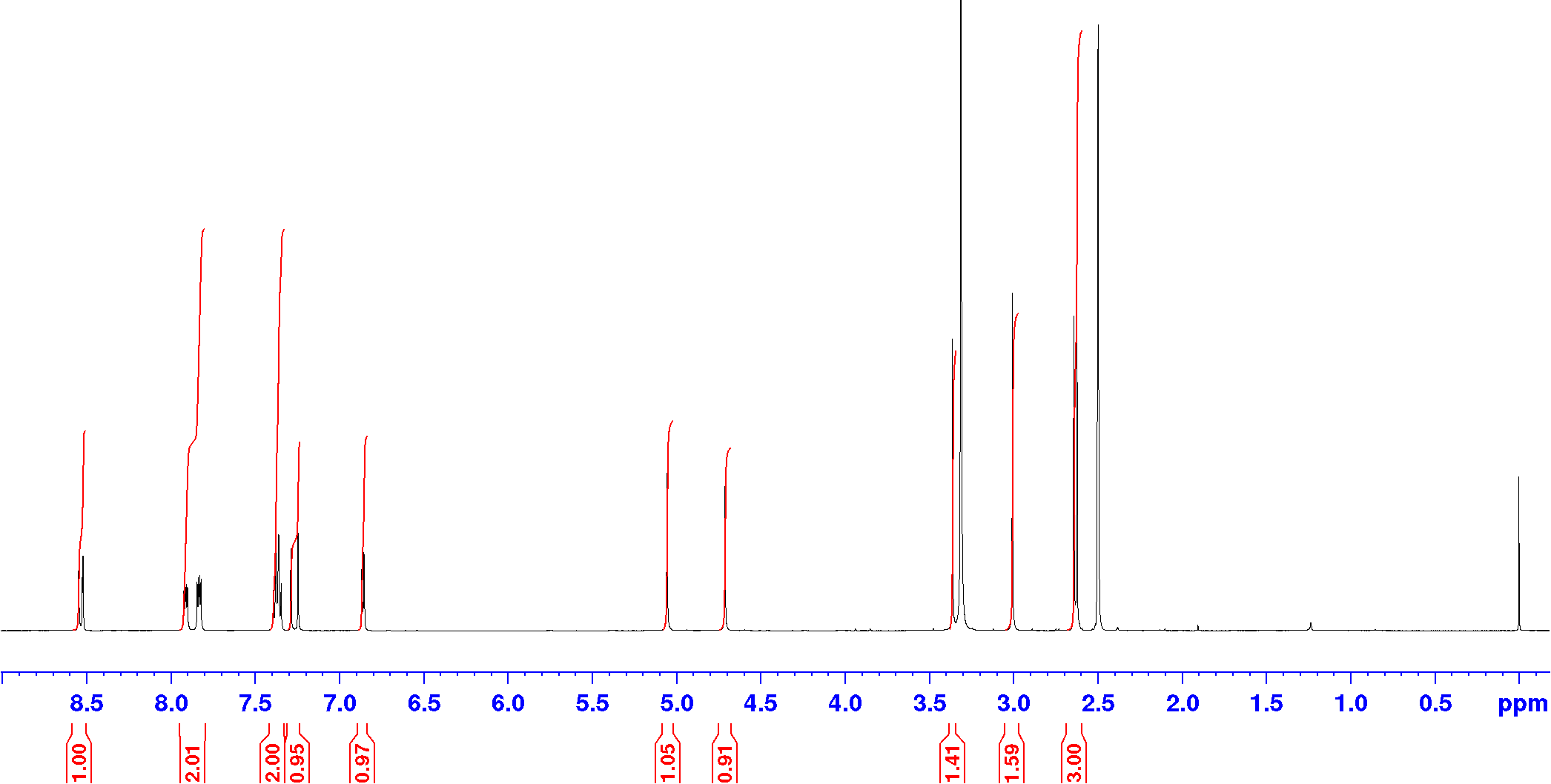

Compound **Z899051432**, ^13^C-NMR (DEPTQ), DMSO-d_6_, 151 MHz

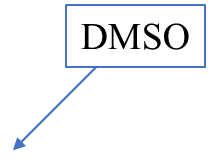

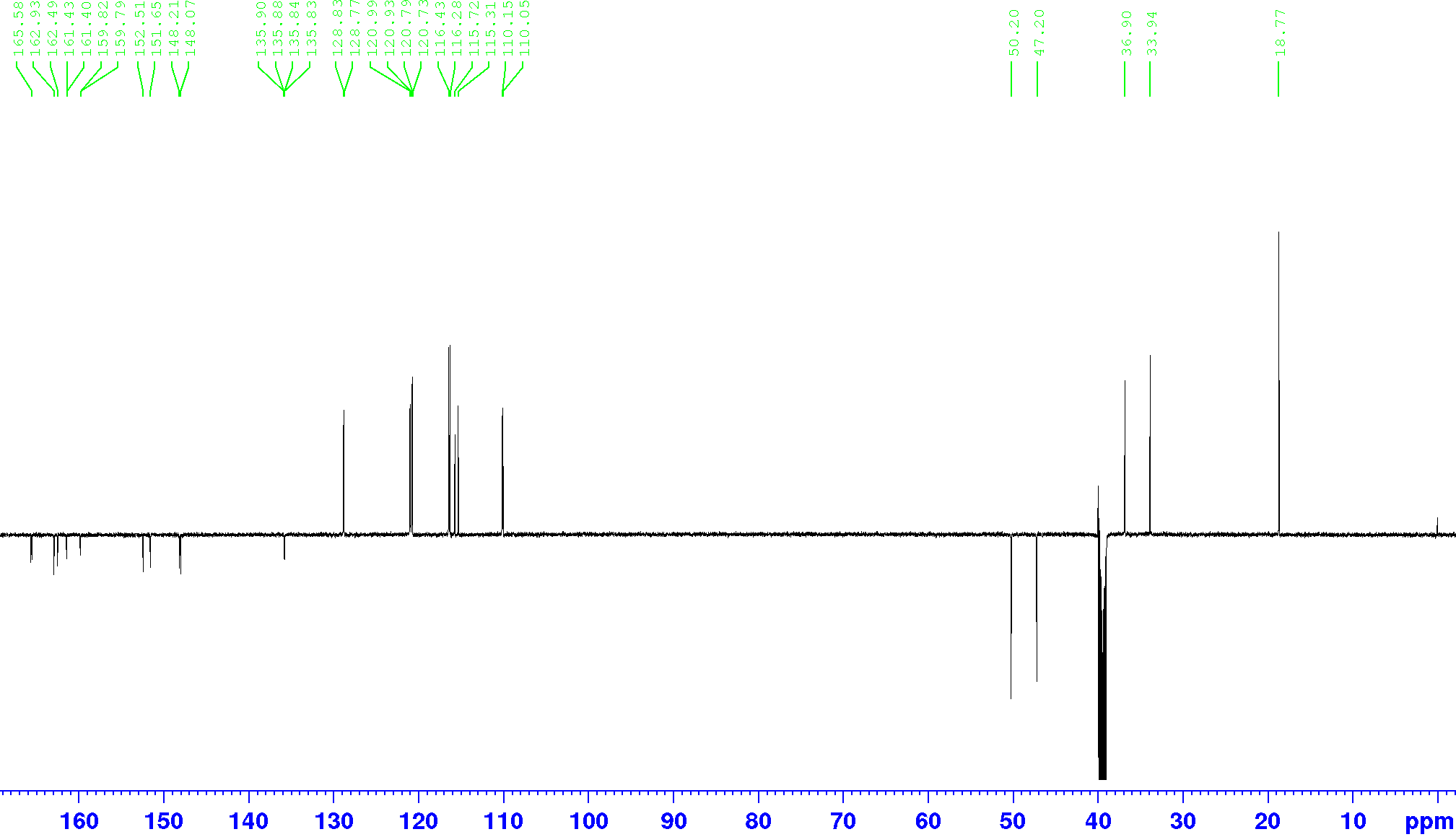

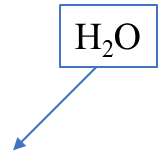

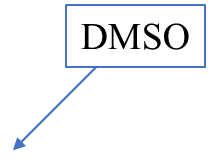
Compound **Z1262422554**, ^1^H-NMR, DMSO-d_6_, 600 MHz

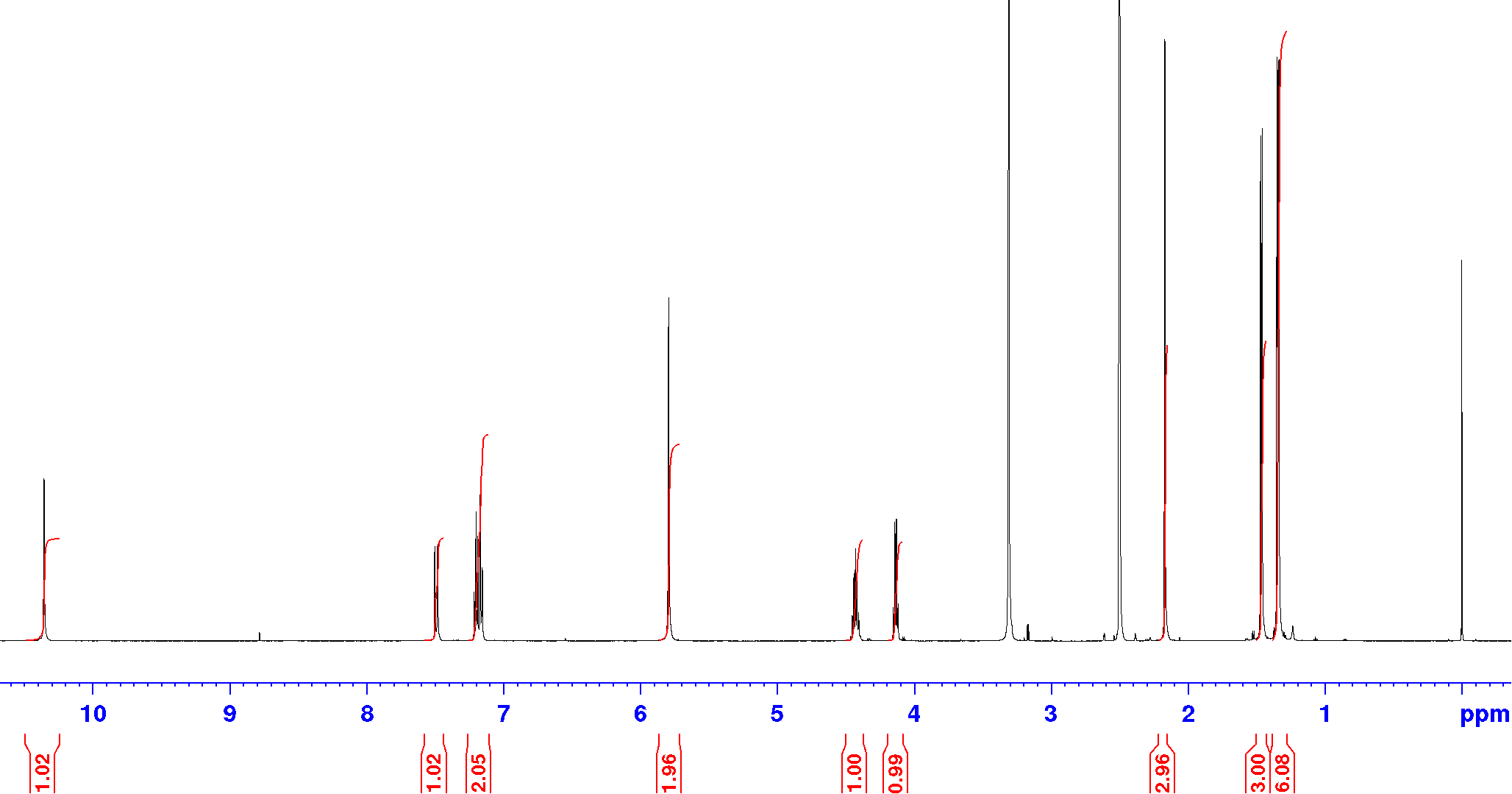

Compound **Z1262422554**, ^13^C-NMR (DEPTQ), DMSO-d_6_, 151 MHz

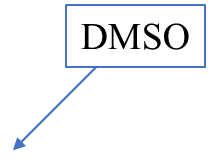

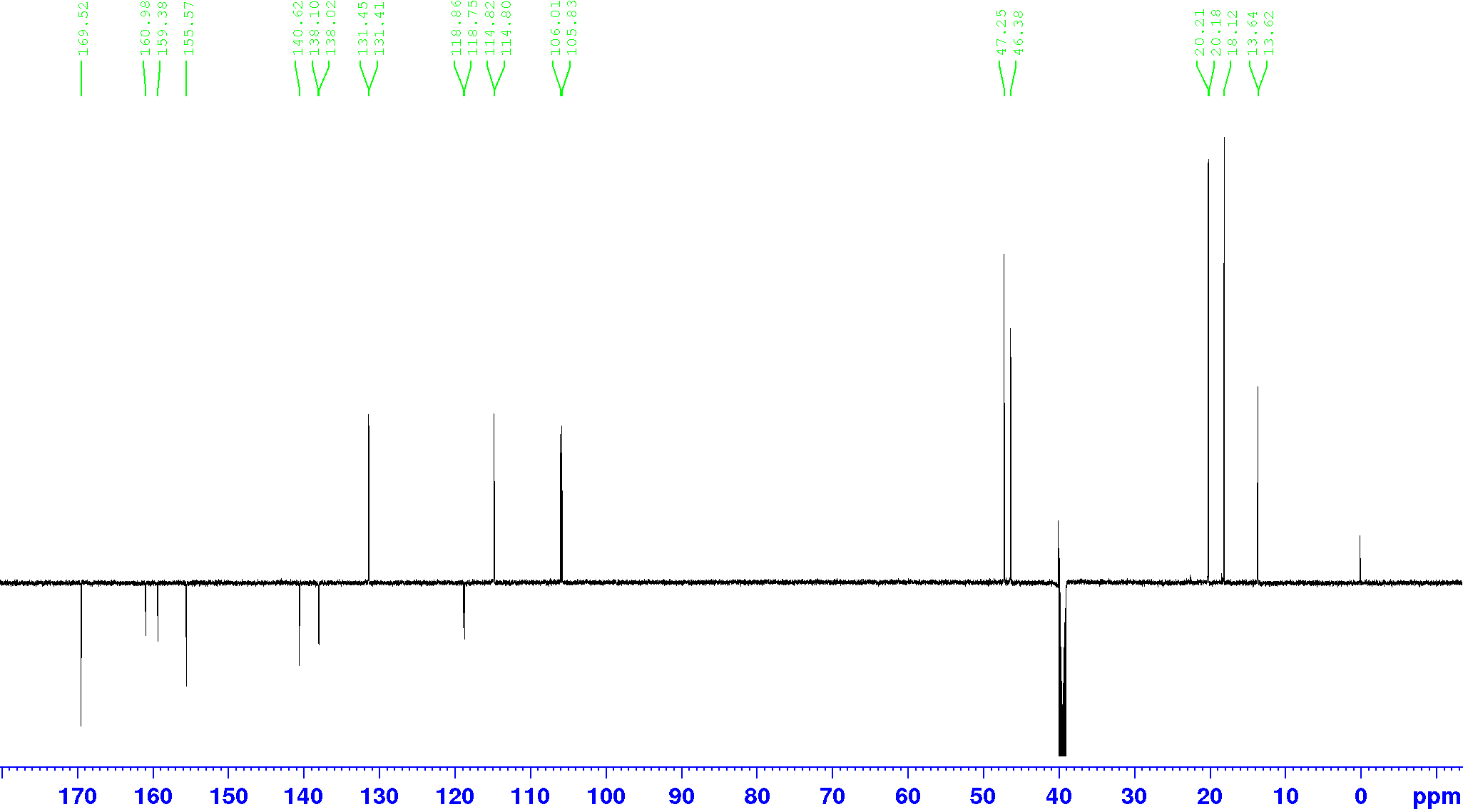

Compound **Z2171315755**, ^1^H-NMR, DMSO-d_6_, 600 MHz

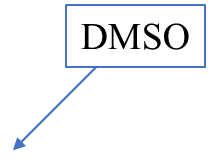

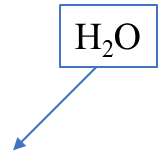

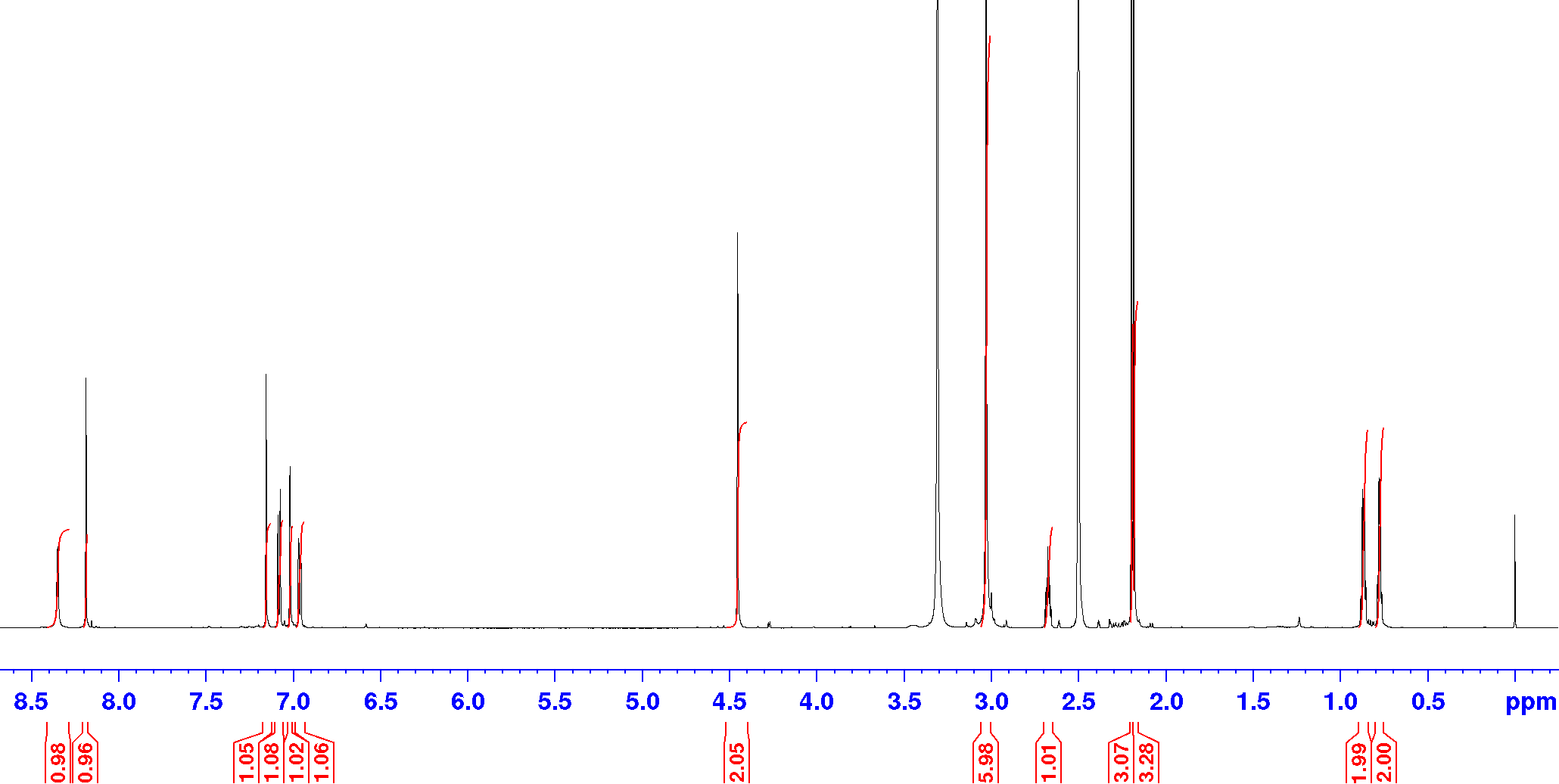

Compound **Z2171315755**, ^13^C-NMR (DEPTQ), DMSO-d_6_, 151 MHz

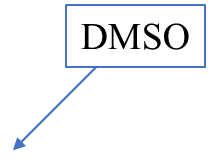

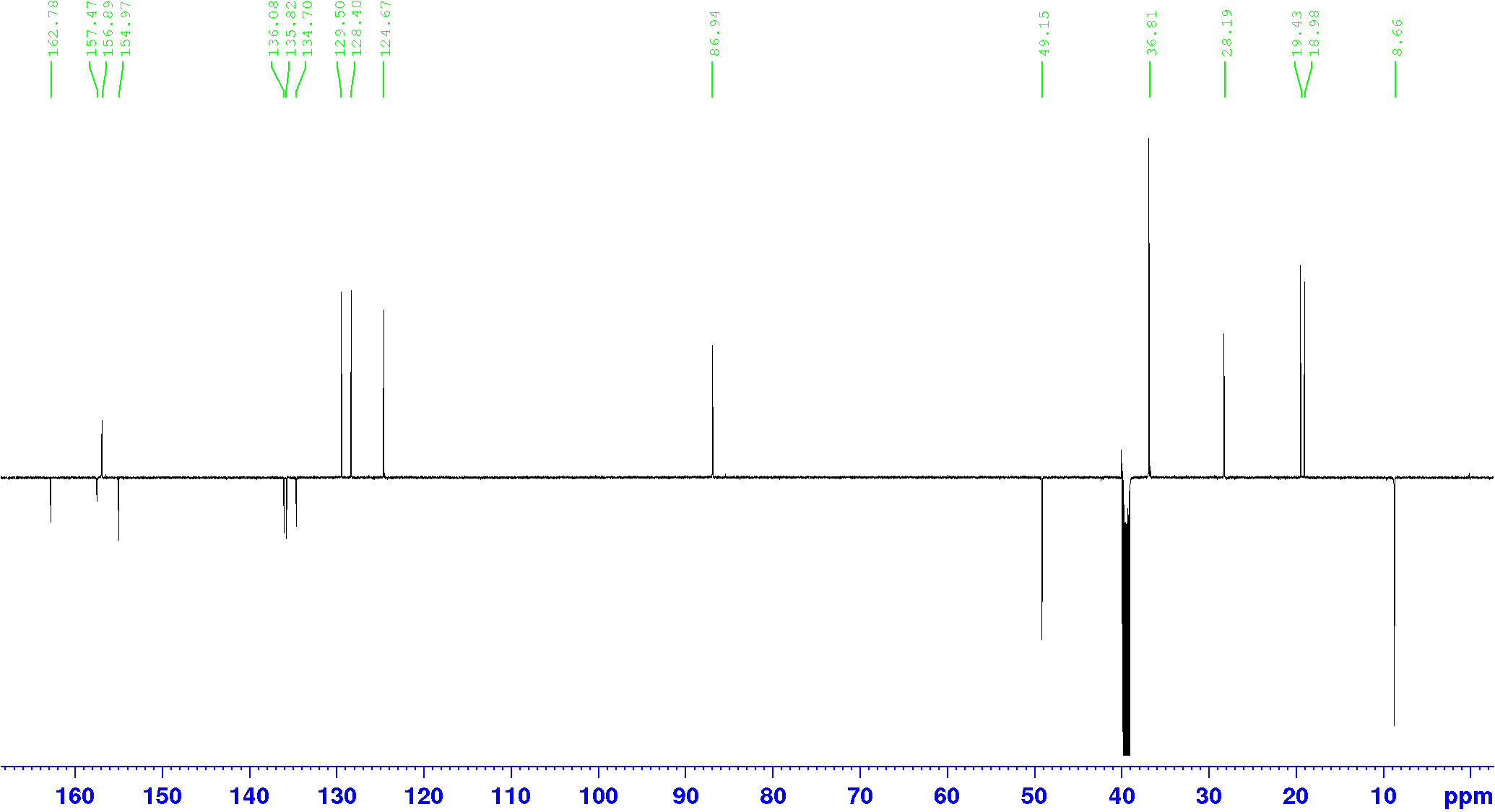

Compound **Z995908944**, ^1^H-NMR, DMSO-d_6_, 600 MHz

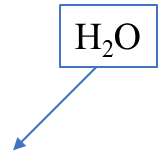

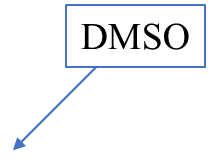

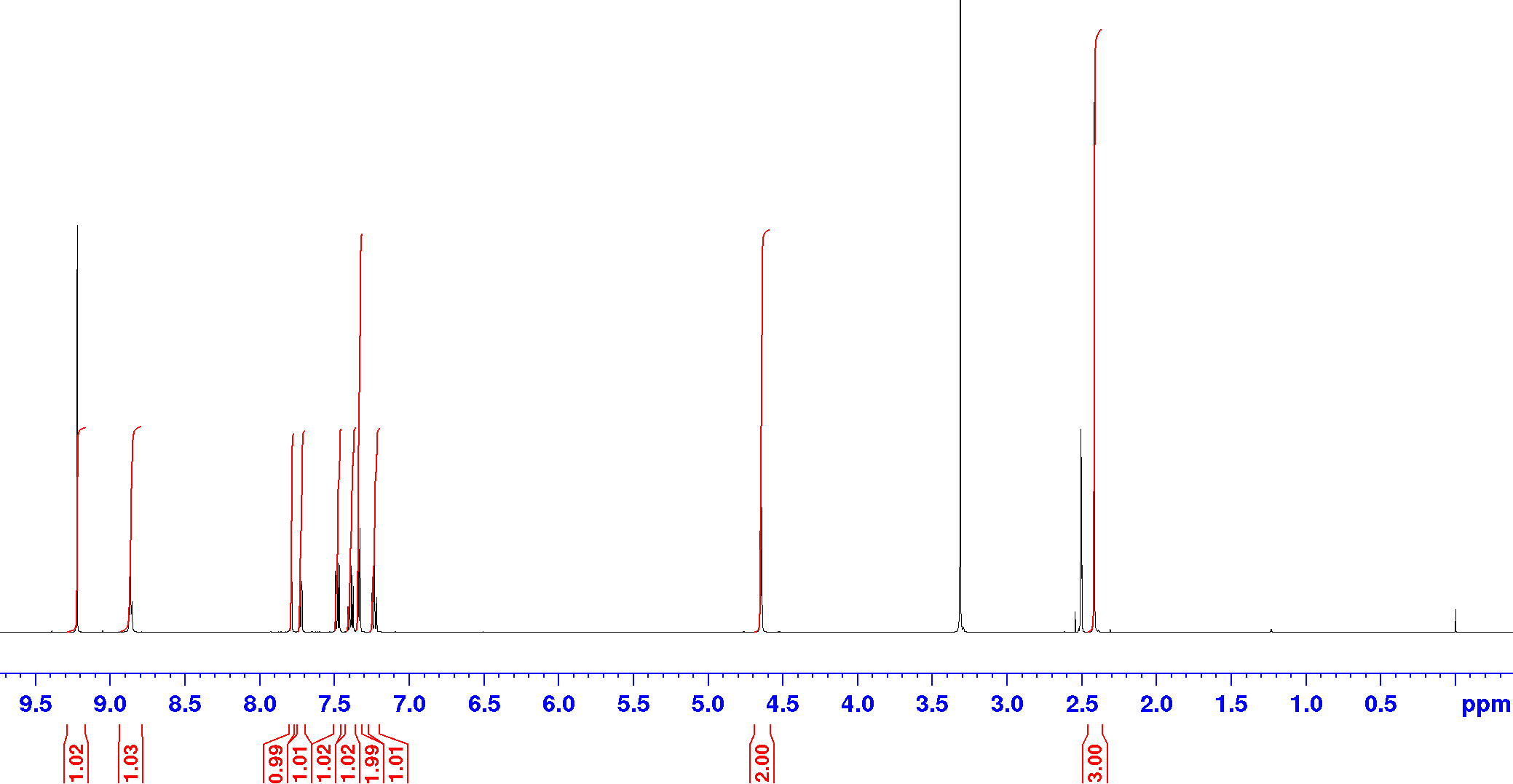

Compound **Z995908944**, ^13^C-NMR (DEPTQ), DMSO-d_6_, 151 MHz

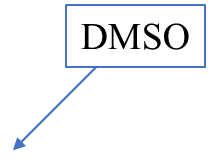

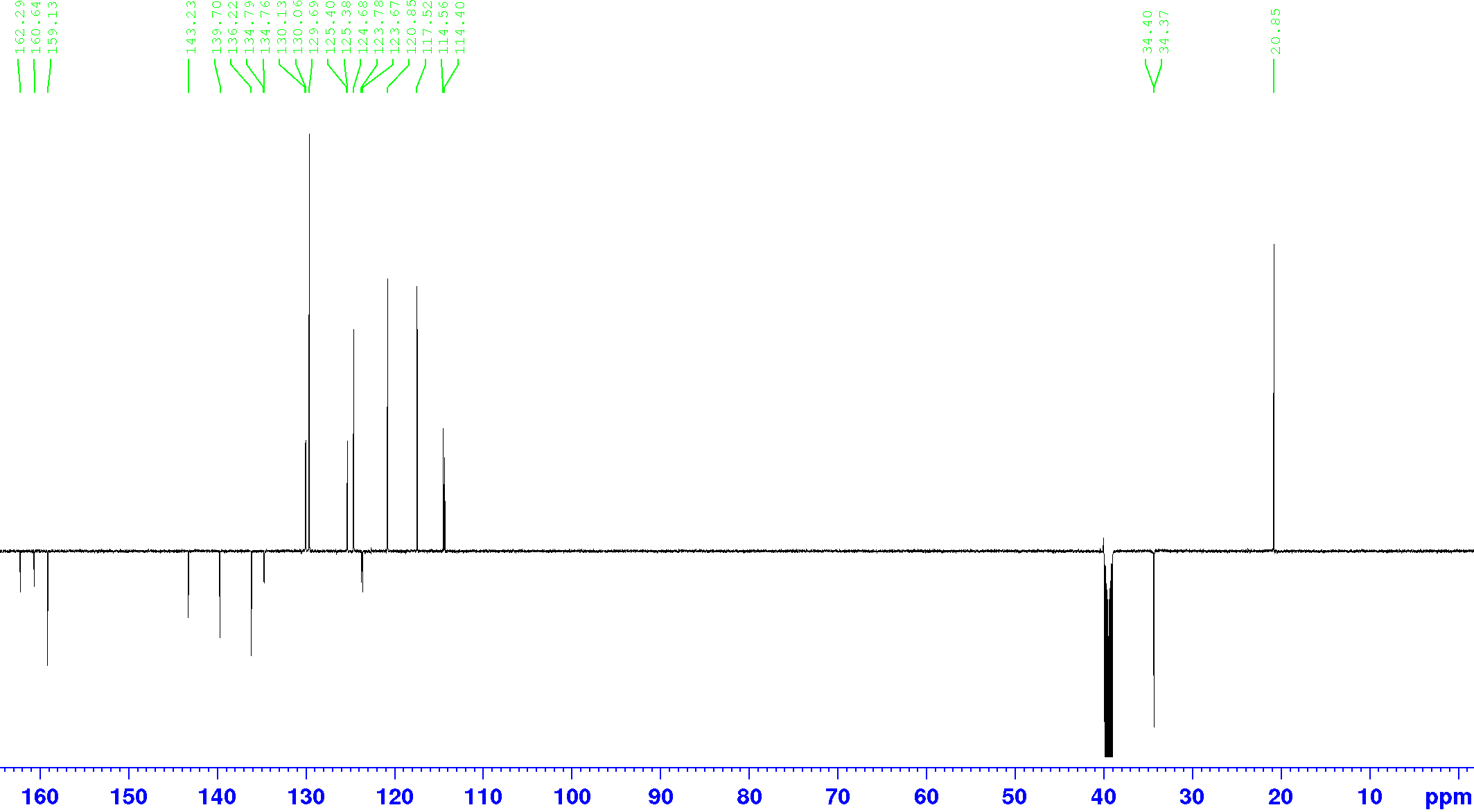

**^1^H and ^13^C NMR Spectra of the Compounds compounds of type 1-10**

Compound **1a**, ^1^H-NMR, DMSO-d_6_, 600 MHz

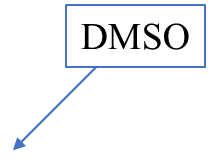

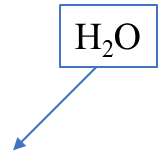

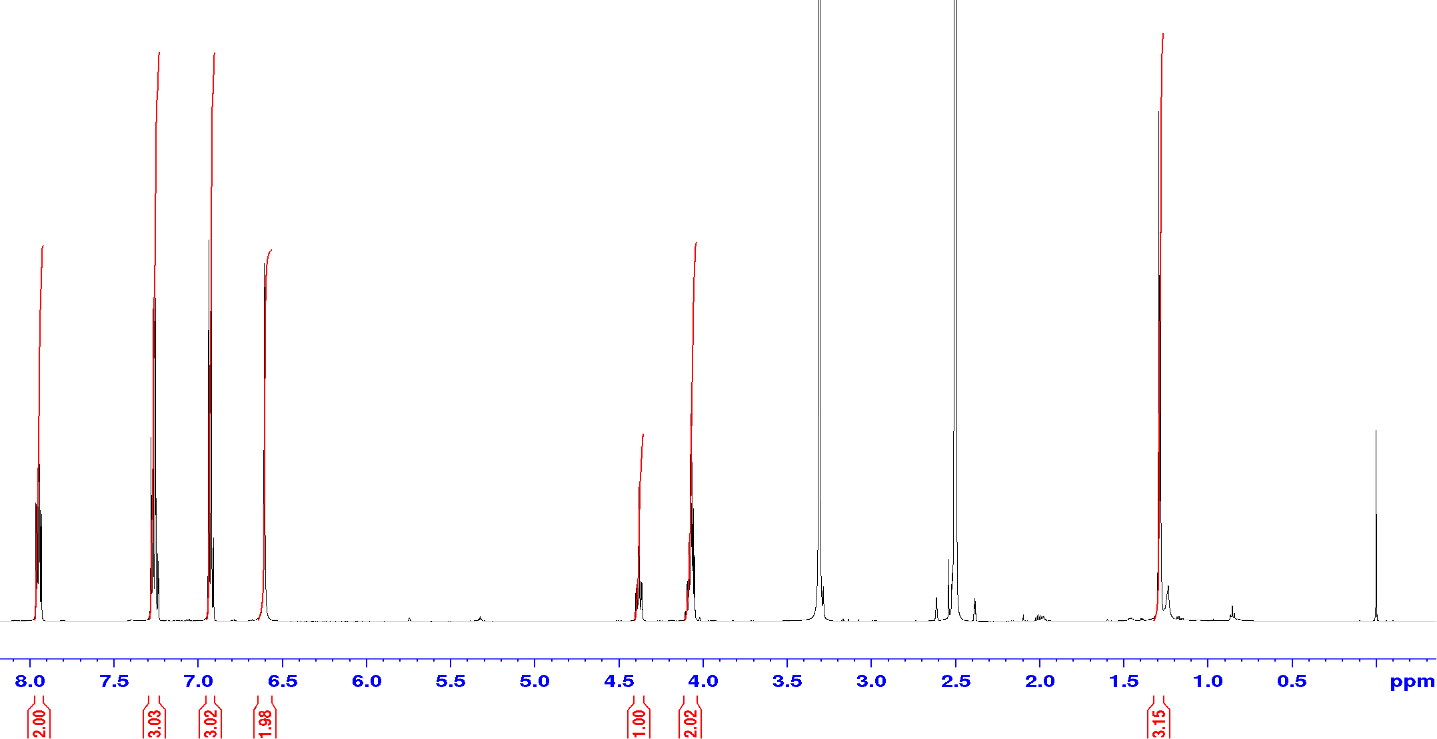

Compound **1a**, ^13^C-NMR (DEPTQ), DMSO-d_6_, 151 MHz

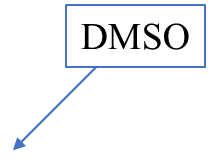

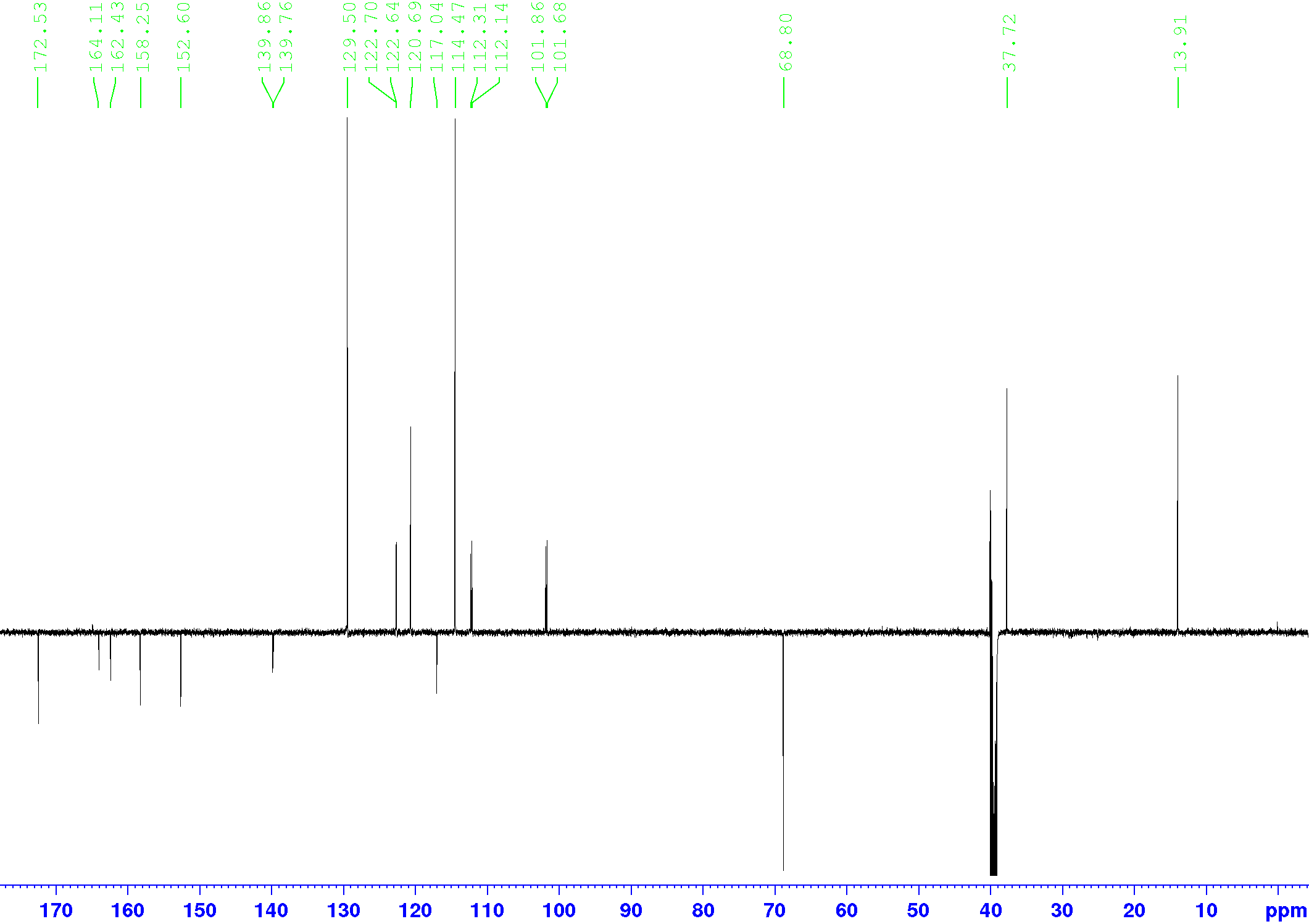

Compound **1b**, ^1^H-NMR, DMSO-d_6_, 600 MHz

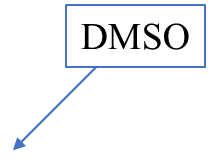

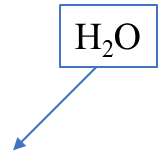

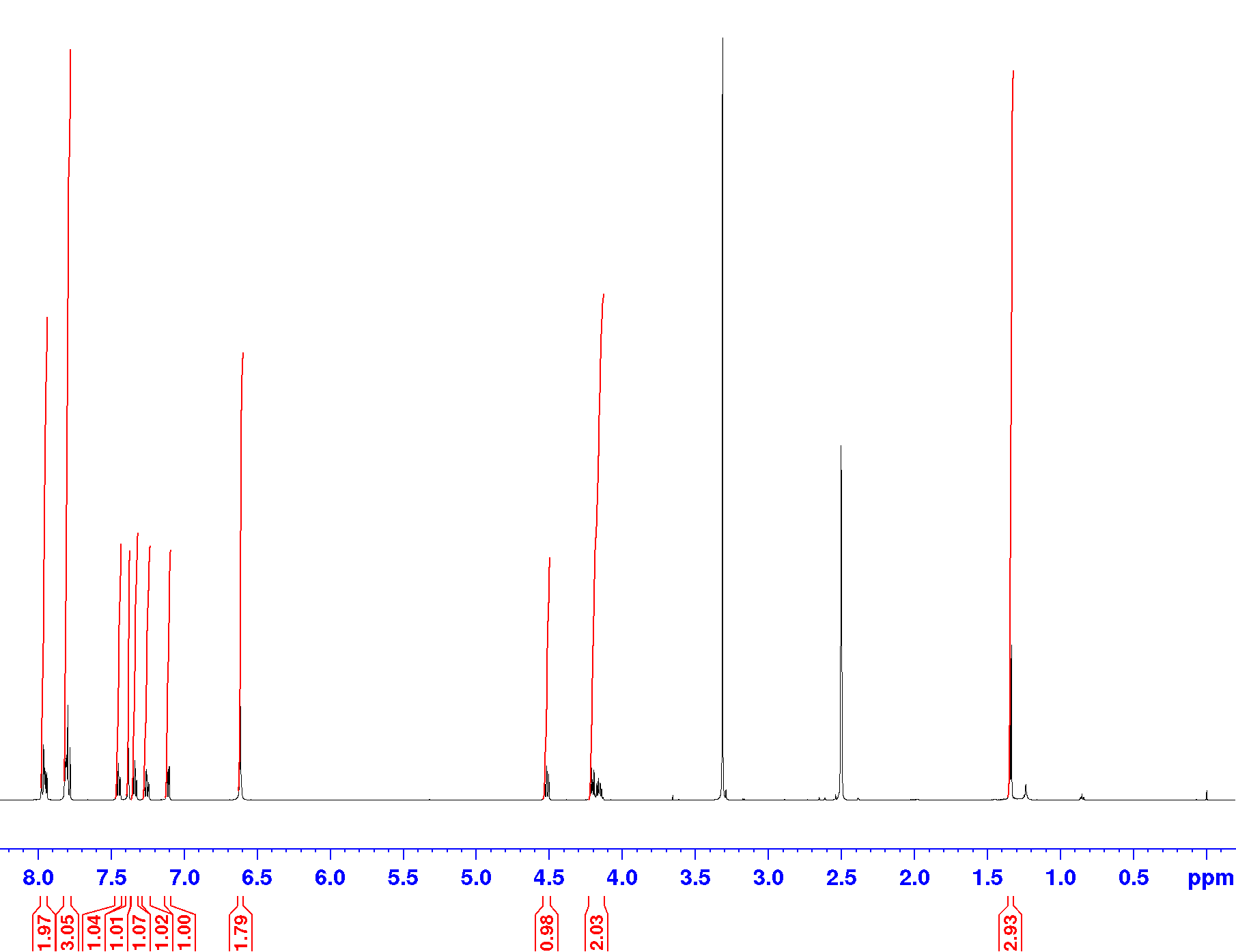

Compound **1b**, ^13^C-NMR (DEPTQ), DMSO-d_6_, 151 MHz

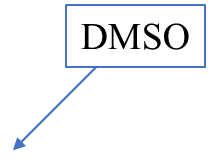

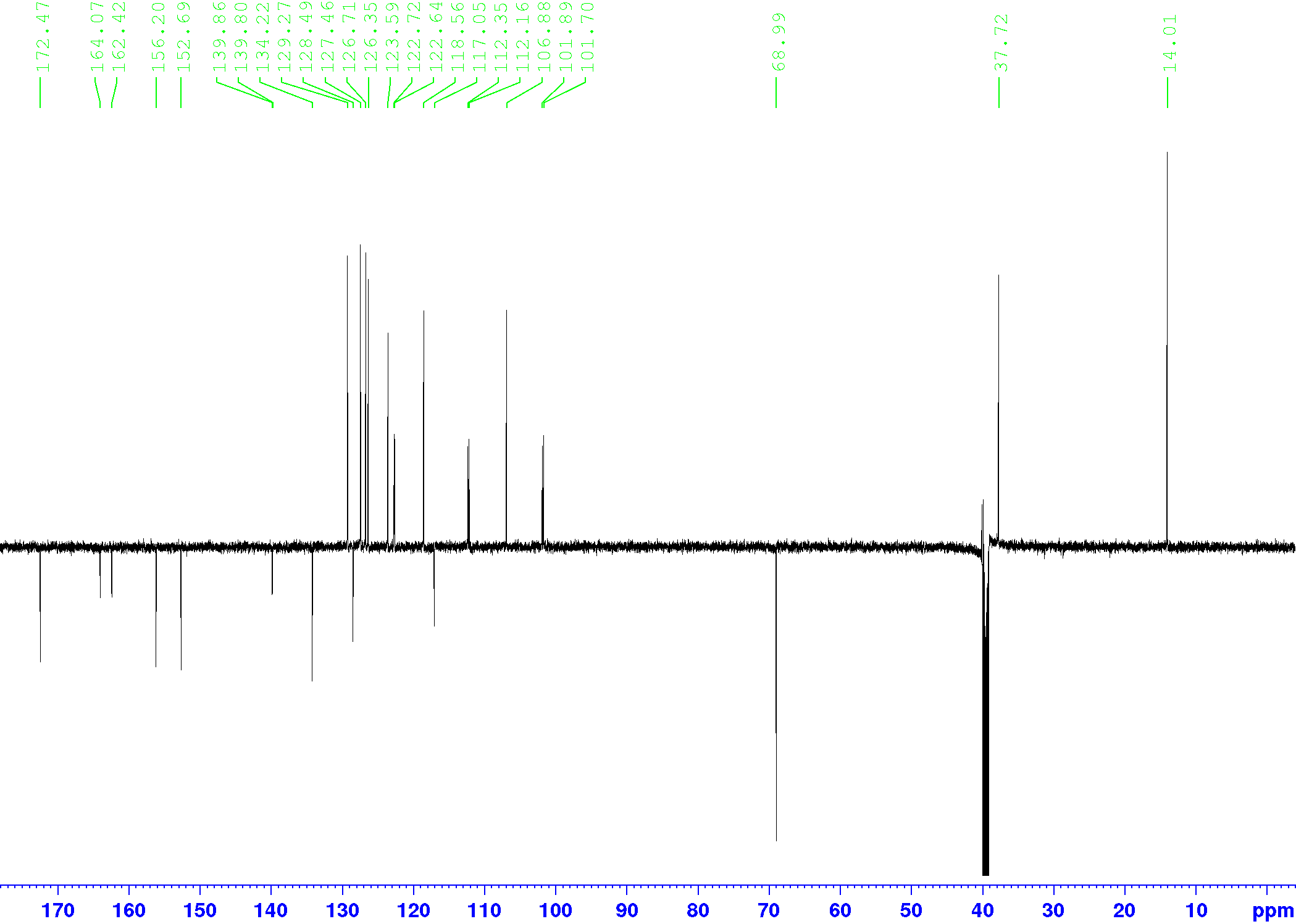

Compound **1c**, ^1^H-NMR, DMSO-d_6_, 600 MHz

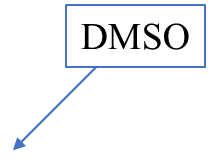

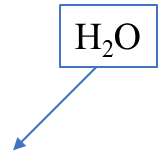

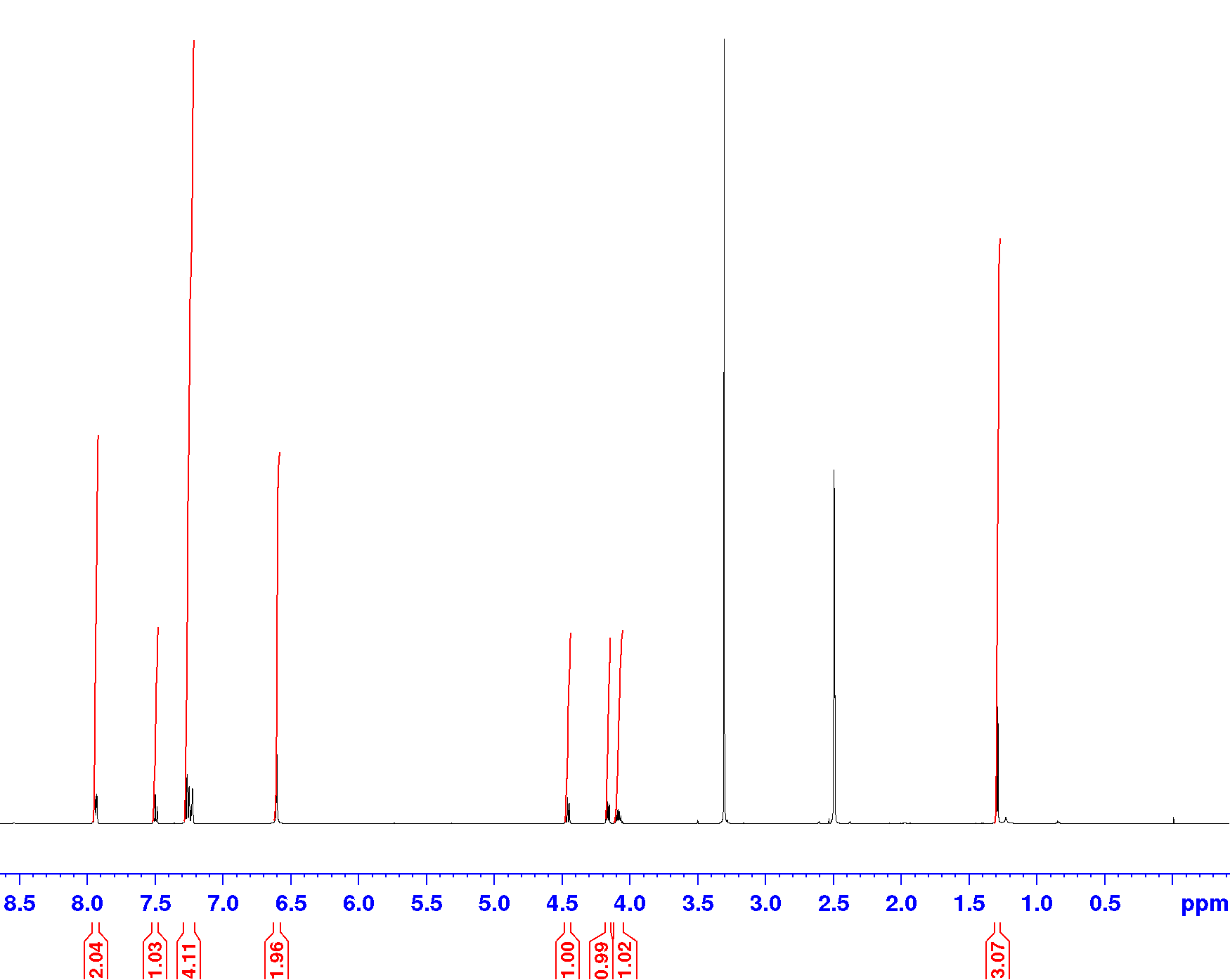

Compound **1c**, ^13^C-NMR (DEPTQ), DMSO-d_6_, 151 MHz

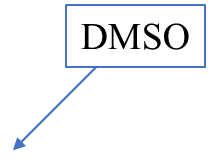

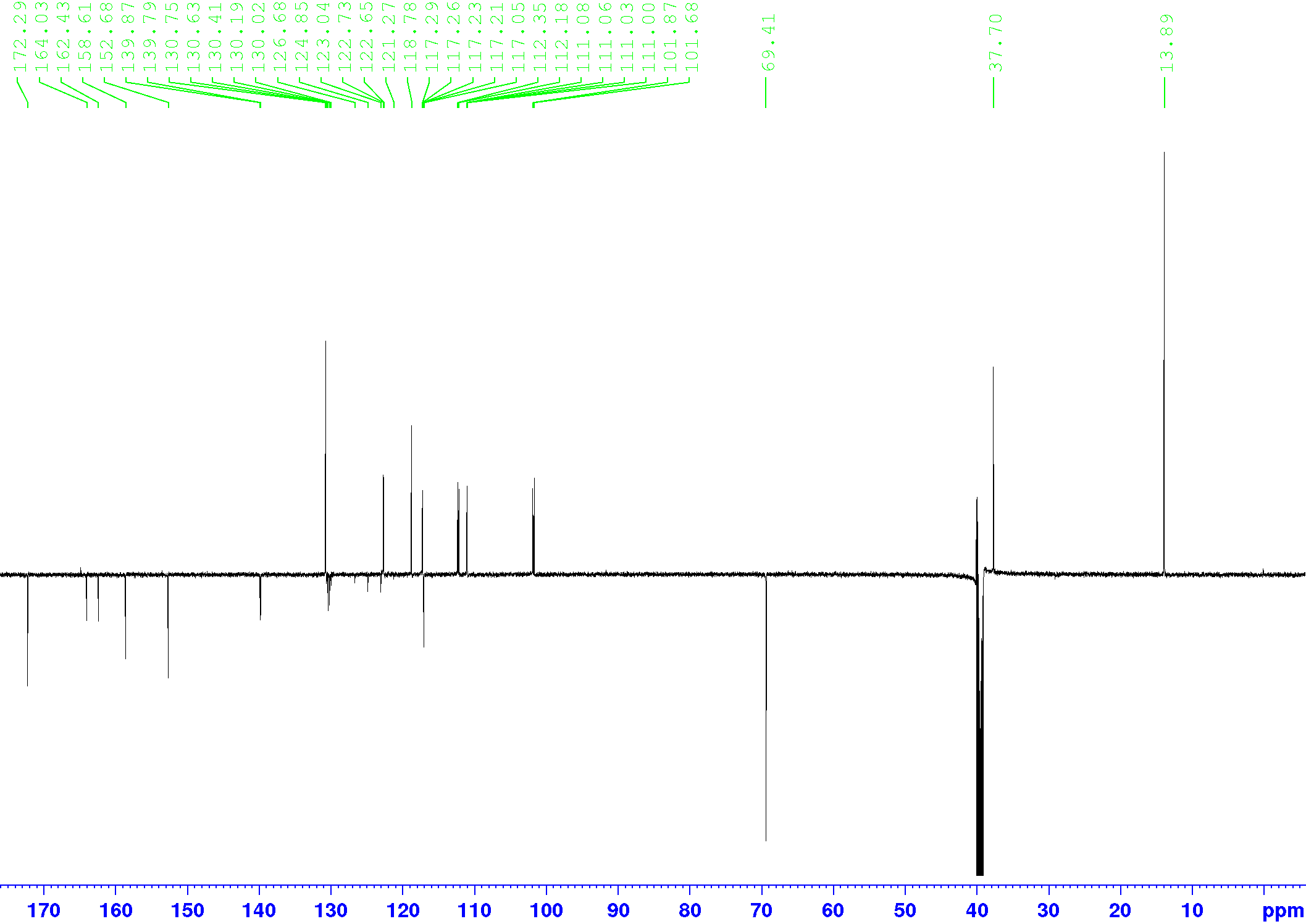

Compound **1d**, ^1^H-NMR, DMSO-d_6_, 600 MHz

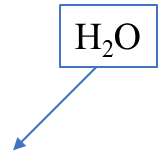

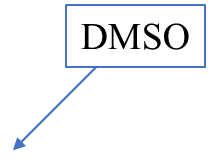

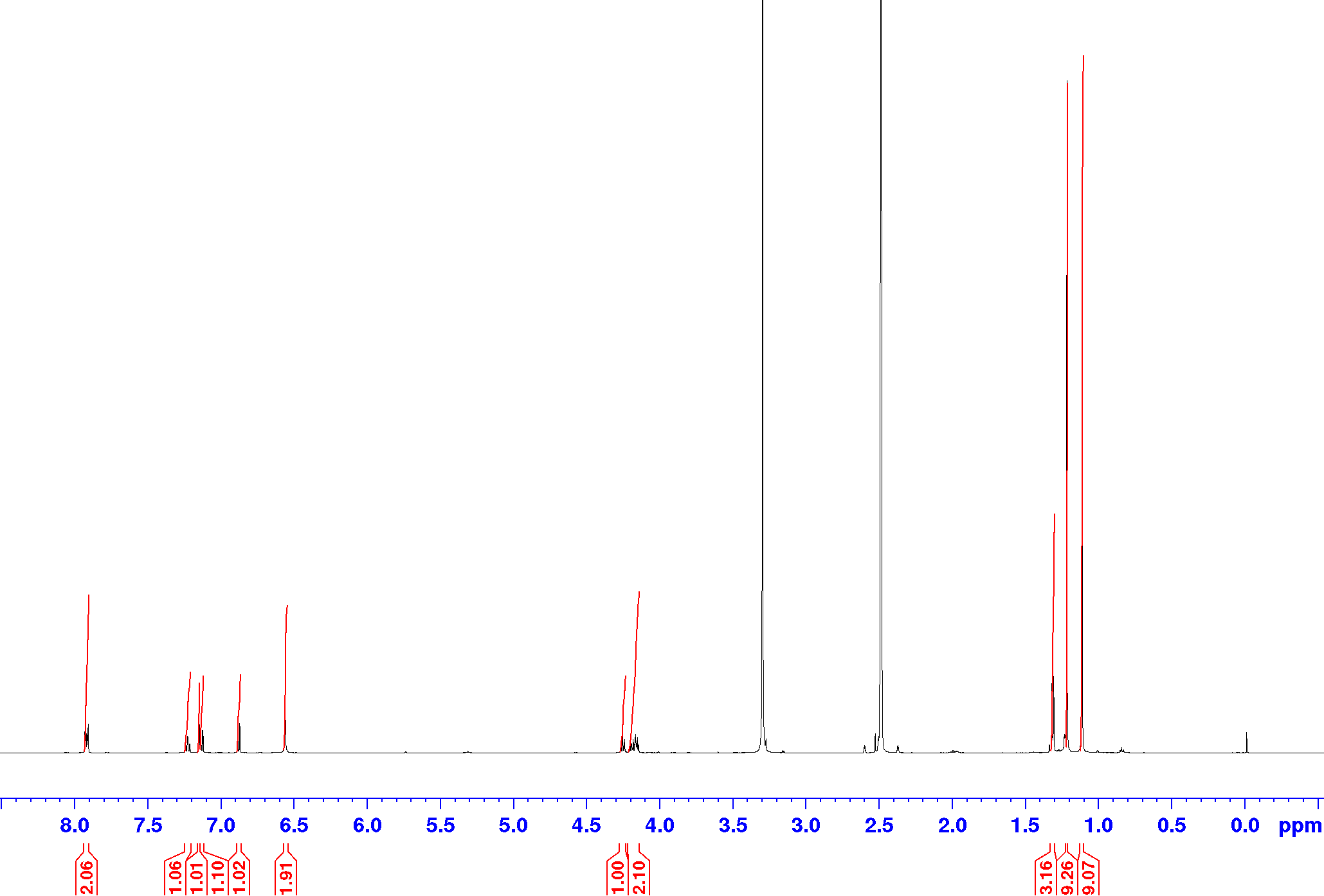

Compound **1d**, ^13^C-NMR (DEPTQ), DMSO-d_6_, 151 MHz

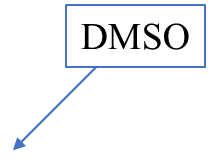

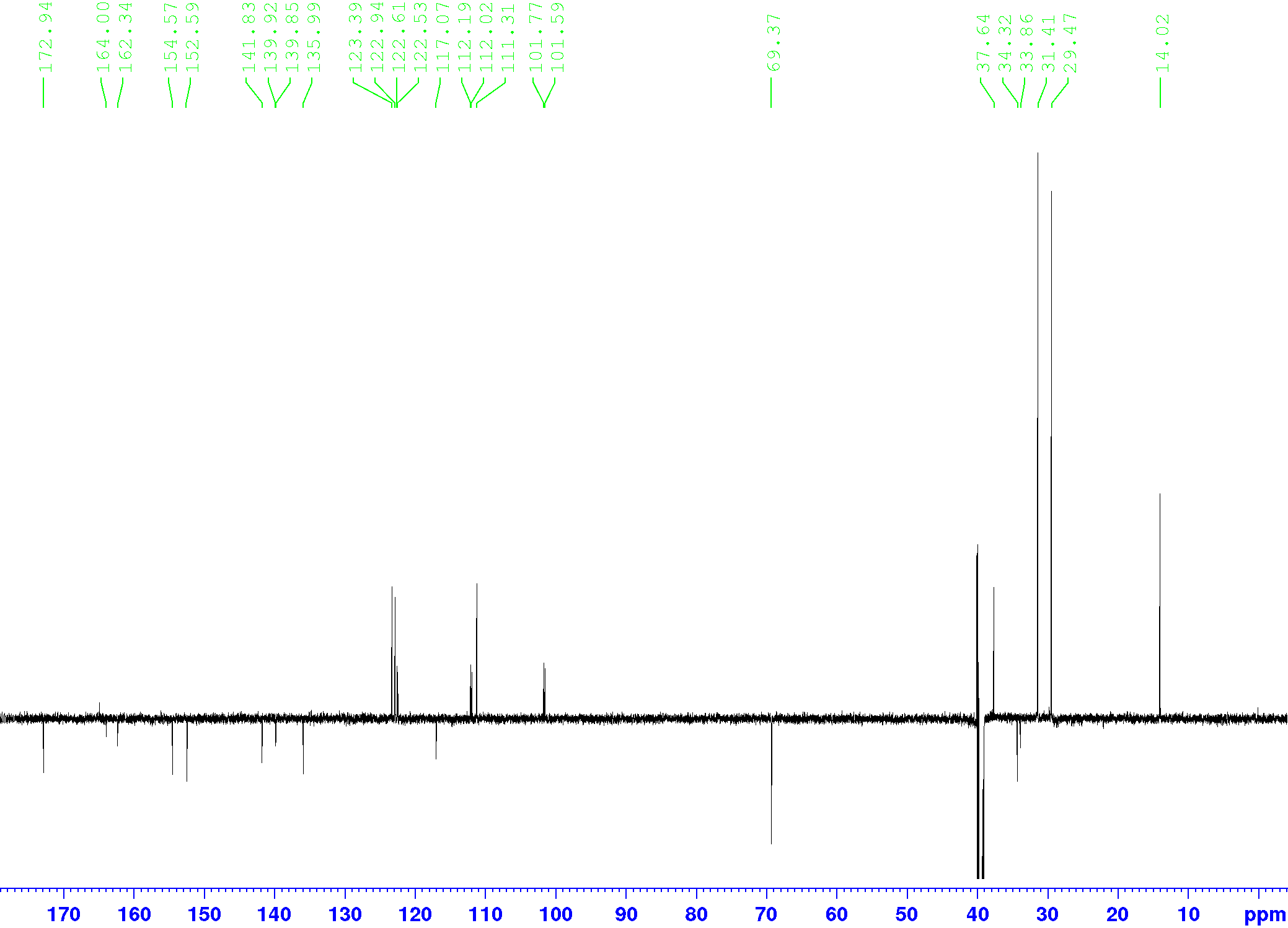

Compound **1e**, ^1^H-NMR, DMSO-d_6_, 600 MHz

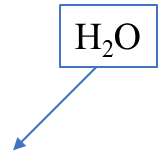

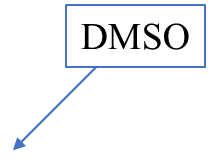

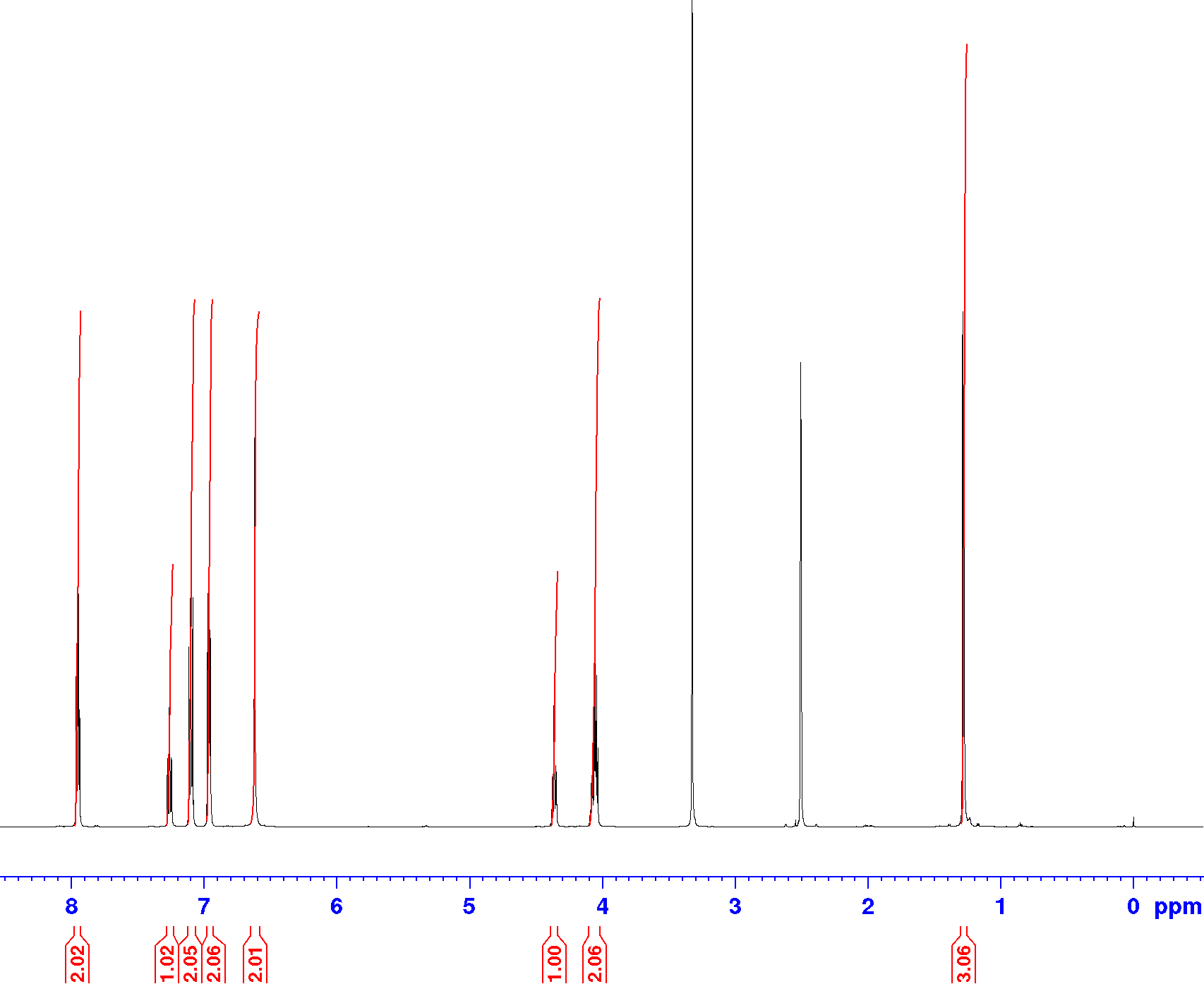

Compound **1e**, ^13^C-NMR (DEPTQ), DMSO-d_6_, 151 MHz

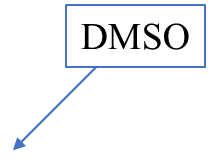

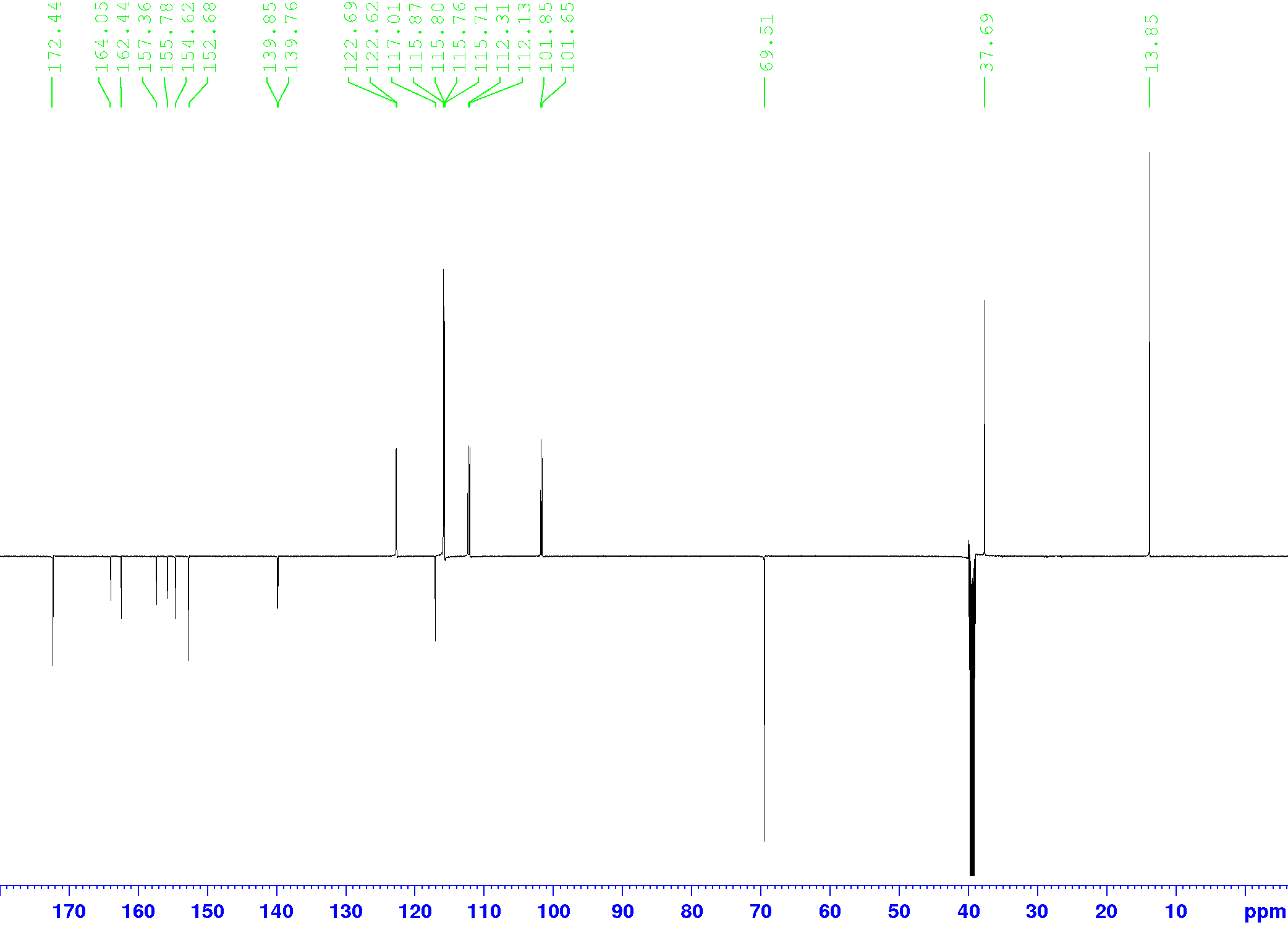

Compound **1f**, ^1^H-NMR, DMSO-d_6_, 600 MHz

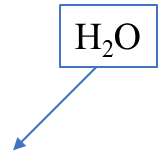

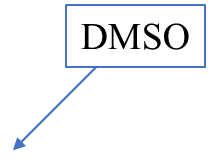

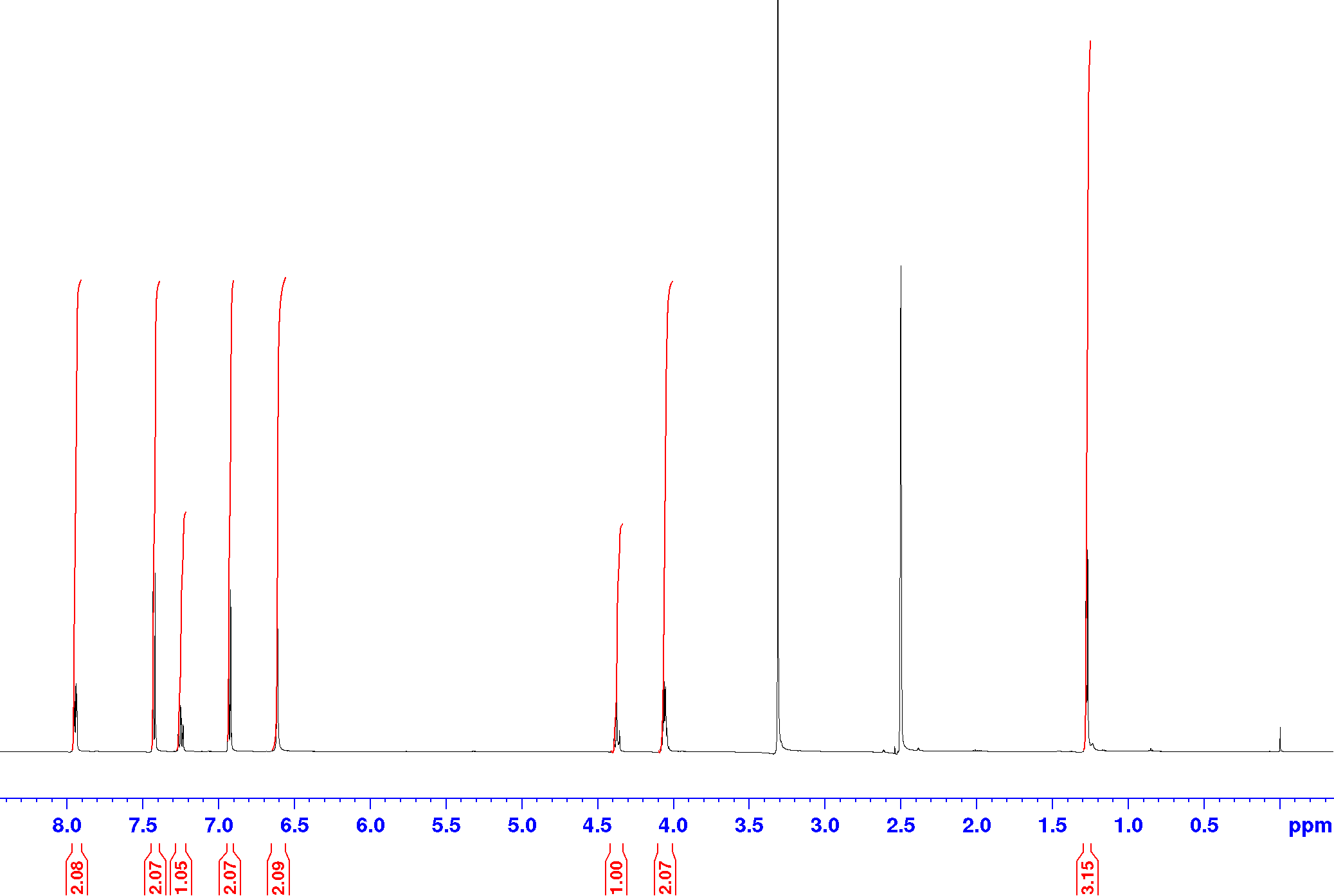

Compound **1f**, ^13^C-NMR (DEPTQ), DMSO-d_6_, 151 MHz

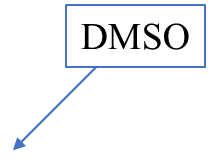

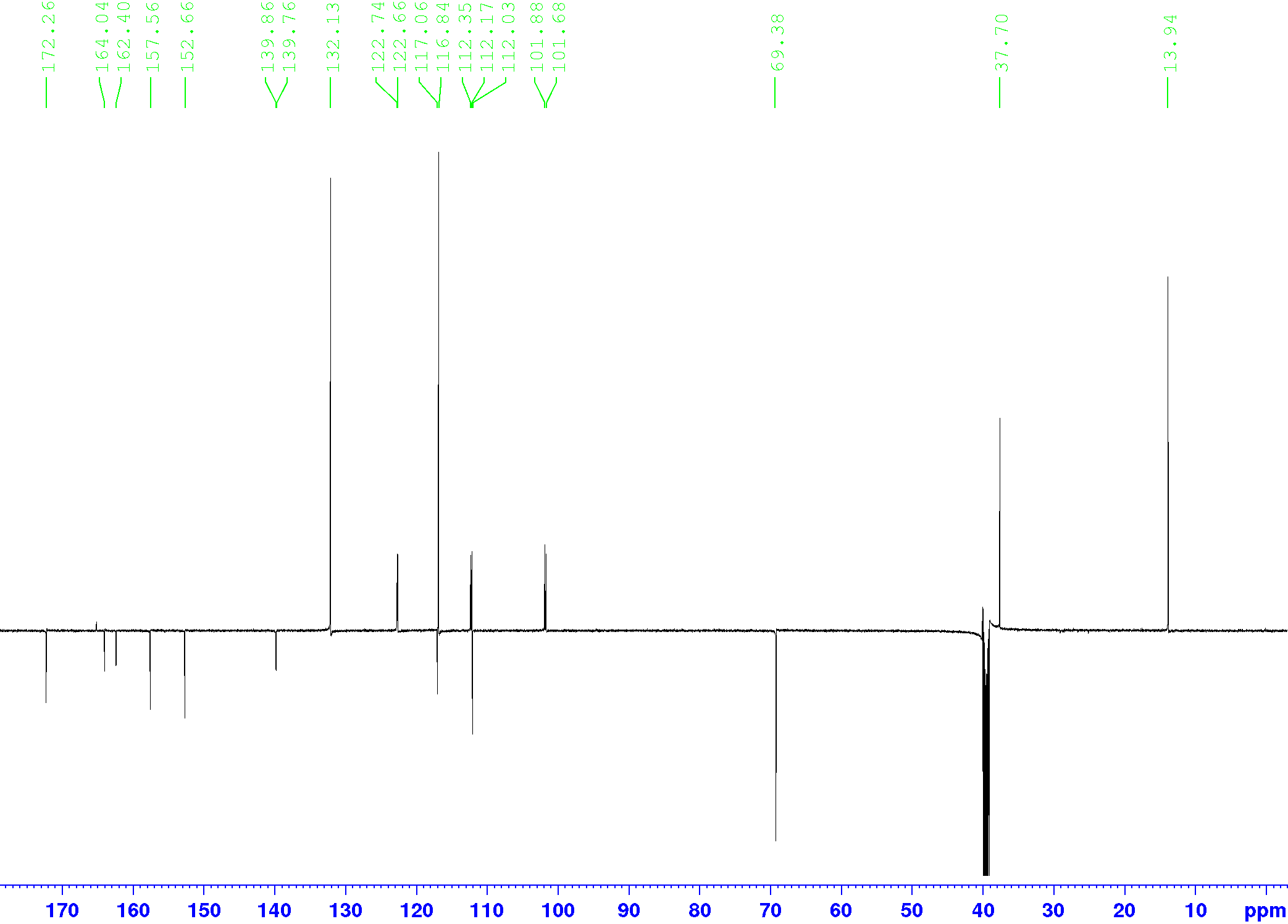

Compound **1g**, ^1^H-NMR, DMSO-d_6_, 600 MHz

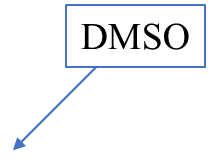

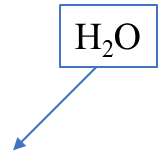

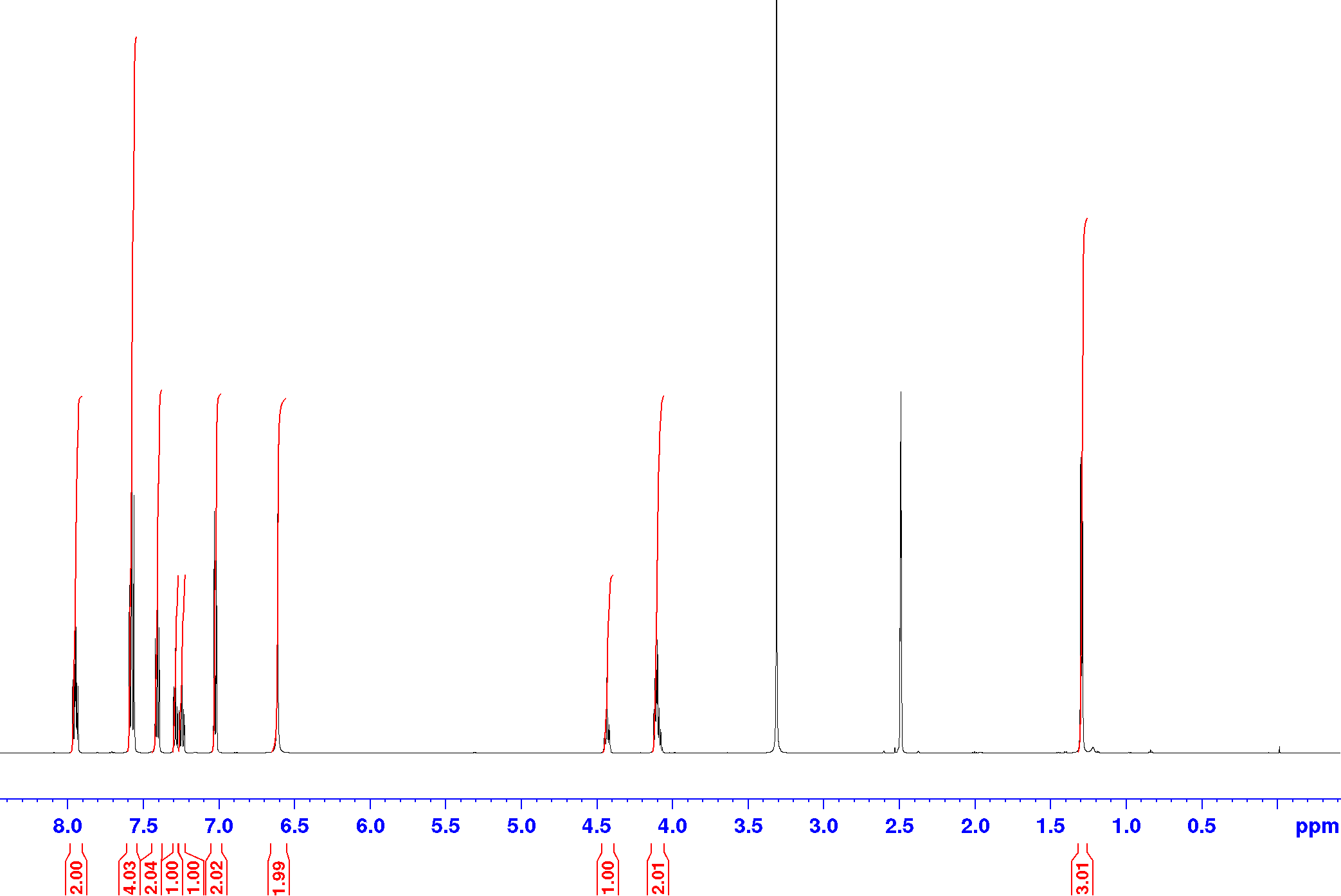

Compound **1g**, ^13^C-NMR (DEPTQ), DMSO-d_6_, 151 MHz

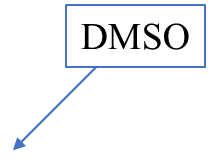

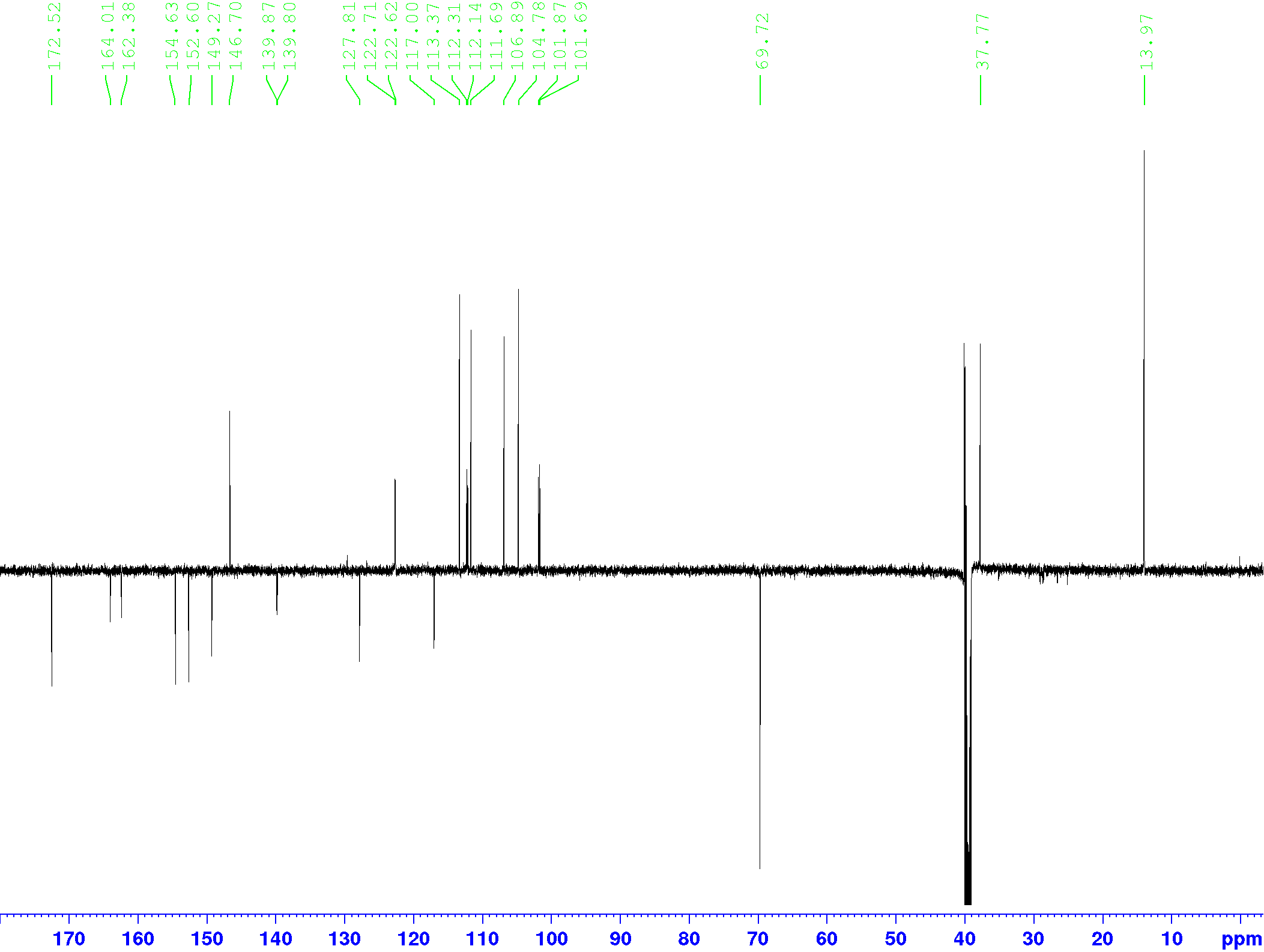

Compound **1h**, ^1^H-NMR, DMSO-d_6_, 600 MHz

Compound **1h**, ^13^C-NMR (DEPTQ), DMSO-d_6_, 151 MHz

Compound **2**, ^1^H-NMR, DMSO-d_6_, 600 MHz

Compound **2**, ^13^C-NMR (DEPTQ), DMSO-d_6_, 151 MHz

Compound **3a**, ^1^H-NMR, DMSO-d_6_, 600 MHz

Compound **3a**, ^13^C-NMR (DEPTQ), DMSO-d_6_, 151 MHz

Compound **3b**, ^1^H-NMR, DMSO-d_6_, 600 MHz

Compound **3b**, ^13^C-NMR (DEPTQ), DMSO-d_6_, 151 MHz

Compound **16**, ^1^H-NMR, DMSO-d_6_, 600 MHz

Compound **16**, ^13^C-NMR (DEPTQ), DMSO-d_6_, 151 MHz

Compound **17**, ^1^H-NMR, DMSO-d_6_, 600 MHz

Compound **17**, ^13^C-NMR (DEPTQ), DMSO-d_6_, 151 MHz

Compound **18**, ^1^H-NMR, DMSO-d_6_, 600 MHz

Compound **18**, ^13^C-NMR (DEPTQ), DMSO-d_6_, 151 MHz

Compound **3c**, ^1^H-NMR, DMSO-d_6_, 600 MHz

Compound **3c**, ^13^C-NMR (DEPTQ), DMSO-d_6_, 151 MHz

Compound **4**, ^1^H-NMR, DMSO-d_6_, 600 MHz

Compound **4**, ^13^C-NMR (DEPTQ), DMSO-d_6_, 151 MHz

Compound **5a**, ^1^H-NMR, DMSO-d_6_, 400 MHz

Compound **5a**, ^13^C-NMR (DEPTQ), DMSO-d_6_, 151 MHz

Compound **5b**, ^1^H-NMR, DMSO-d_6_, 400 MHz

Compound **5b**, ^13^C-NMR (DEPTQ), DMSO-d_6_, 101 MHz

Compound **5c**, ^1^H-NMR, DMSO-d_6_, 600 MHz

Compound **5c**, ^13^C-NMR (DEPTQ), DMSO-d_6_, 101 MHz

Compound **5d**, ^1^H-NMR, DMSO-d_6_, 600 MHz

Compound **5d**, ^13^C-NMR (DEPTQ), DMSO-d_6_, 151 MHz

Compound **5e**, ^1^H-NMR, DMSO-d_6_, 600 MHz

Compound **5e**, ^13^C-NMR (DEPTQ), DMSO-d_6_, 151 MHz

Compound **27**, ^1^H-NMR, DMSO-d_6_, 600 MHz

Compound **27**, ^13^C-NMR (DEPTQ), DMSO-d_6_, 151 MHz

Compound **5f**, ^1^H-NMR, DMSO-d_6_, 600 MHz

Compound **5f**, ^13^C-NMR (DEPTQ), DMSO-d_6_, 151 MHz

Compound **29**, ^1^H-NMR, DMSO-d_6_, 600 MHz

Compound **29**, ^13^C-NMR (DEPTQ), DMSO-d_6_, 151 MHz

Compound **5g**, ^1^H-NMR, DMSO-d_6_, 600 MHz

Compound **5g**, ^13^C-NMR (DEPTQ), DMSO-d_6_, 151 MHz

Compound **F-5g**, ^1^H-NMR, DMSO-d_6_, 600 MHz

Compound **F-5g**, ^13^C-NMR (DEPTQ), DMSO-d_6_, 151 MHz

Compound **6a**, ^1^H-NMR, DMSO-d_6_, 600 MHz

Compound **6a**, ^13^C-NMR (DEPTQ), DMSO-d_6_, 151 MHz

Compound **6b**, ^1^H-NMR, DMSO-d_6_, 600 MHz

Compound **6b**, ^13^C-NMR (DEPTQ), DMSO-d_6_, 151 MHz

Compound **7a**, ^1^H-NMR, DMSO-d_6_, 600 MHz

Compound **7a**, ^13^C-NMR (DEPTQ), DMSO-d_6_, 151 MHz

Compound **7b**, ^1^H-NMR, DMSO-d_6_, 600 MHz

Compound **7b**, ^13^C-NMR (DEPTQ), DMSO-d_6_, 151 MHz

Compound **8**, ^1^H-NMR, DMSO-d_6_, 600 MHz

Compound **8**, ^13^C-NMR (DEPTQ), DMSO-d_6_, 151 MHz

Compound **36**, ^1^H-NMR, CDCl_3_, 400 MHz

CH_2_Cl_2_

Compound **36**, ^13^C-NMR (DEPTQ), CDCl_3_, 151 MHz

CDCl_3_

Compound **37**, ^1^H-NMR, DMSO-d_6_, 600 MHz

Compound **37**, ^13^C-NMR (DEPTQ), DMSO-d_6_, 151 MHz

Compound **9**, ^1^H-NMR, DMSO-d_6_, 600 MHz

Compound **9**, ^13^C-NMR (DEPTQ), DMSO-d_6_, 151 MHz

Compound **10a**, ^1^H-NMR, DMSO-d_6_, 600 MHz

Compound **10a**, ^13^C-NMR (DEPTQ), DMSO-d_6_, 151 MHz

Compound **10b**, ^1^H-NMR, DMSO-d_6_, 600 MHz

Compound **10b**, ^13^C-NMR (DEPTQ), DMSO-d_6_, 151 MHz

Compound **10c**, ^1^H-NMR, DMSO-d_6_, 600 MHz

Compound **10c**, ^13^C-NMR (DEPTQ), DMSO-d_6_, 151 MHz

Compound **10d**, ^1^H-NMR, DMSO-d_6_, 600 MHz

Compound **10d**, ^13^C-NMR (DEPTQ), DMSO-d_6_, 151 MHz

Compound **10e**, ^1^H-NMR, DMSO-d_6_, 600 MHz

Compound **10e**, ^13^C-NMR (DEPTQ), DMSO-d_6_, 151 MHz

**HPLC Charts of the library compounds Z2075279358 - Z995908944**

Compound **Z2075279358**, flow rate, 0.5 mL/min, linear gradient in 3–85% CH_3_CN in H_2_O (+ 0.1% TFA) in 0-26 min, purity: 99.6% *(t_R_* = 20.6 min)

**Compound Z2194302854**, flow rate, 0.5 mL/min, linear gradient in 3–85% CH_3_CN in H_2_O (+ 0.1% TFA) in 0-26 min, purity: 98.8% (*t_R_* = 13.4 min)

Compound **Z1343848401** flow rate, 0.5 mL/min, linear gradient in 3–85% CH_3_CN in H_2_O (+ 0.1% TFA) in 0-26 min, purity: 99.7% (*t_R_* = 18.8 min)

**Compound Z2776419998**, two diastereomers, flow rate, 0.5 mL/min, linear gradient in 3–85% CH_3_CN in H_2_O (+ 0.1% TFA) in 0-26 min, purity: 99.1% (diastereomers, *t_R1_* = 10.0 min and *t_R2_* = 10.1 min)

Compound **Z1224795288**, flow rate, 0.5 mL/min, linear gradient in 3–85% CH_3_CN in H_2_O (+ 0.1% TFA) in 0-26 min, purity: 97.7% (*t_R_* = 16.9 min)

**Compound Z19702639,** flow rate, 0.5 mL/min, linear gradient in 3–85% CH_3_CN in H_2_O (+ 0.1% TFA) in 0-26 min, purity: 91.2% (*t_R_* = 22.7 min)

Compound **Z1096199008**, flow rate, 0.5 mL/min, linear gradient in 3–85% CH_3_CN in H_2_O (+ 0.1% TFA) in 0-26 min, purity: 97.0% (*t_R_* = 10.0 min)

**Compound Z873519648,** flow rate, 0.5 mL/min, linear gradient in 3–85% CH_3_CN in H_2_O (+ 0.1% TFA) in 0-26 min, purity: >99% (*t_R_* = 16.3 min)

Compound **Z1783799713**, flow rate, 0.5 mL/min, linear gradient in 3–85% CH_3_CN in H_2_O (+ 0.1% TFA) in 0-26 min, purity: 97.8% (*t_R_* = 14.0 min)

**Compound Z899051432,** flow rate, 0.5 mL/min, linear gradient in 3–85% CH_3_CN in H_2_O (+ 0.1% TFA) in 0-26 min, purity: >99% (*t_R_* = 17.8 min)

Compound **Z1262422554**, flow rate, 0.5 mL/min, linear gradient in 3–85% CH_3_CN in H_2_O (+ 0.1% TFA) in 0-26 min, purity: >99% (*t_R_* = 16.3 min)

**Compound Z2171315755,** flow rate, 0.5 mL/min, linear gradient in 3–85% CH_3_CN in H_2_O (+ 0.1% TFA) in 0-26 min, purity: 95.0% (*t_R_* = 18.5 min)

Compound **Z995908944**, flow rate, 0.5 mL/min, linear gradient in 3–85% CH_3_CN in H_2_O (+ 0.1% TFA) in 0-26 min, purity: >99% *t_R_* = 21.7 min)

**HPLC Charts of the Key Compounds of type 1-10**

**Compound 1a, (*S*)-‘358**, flow rate, 0.5 mL/min, linear gradient a) 5–5% CH_3_CN in H_2_O (+ 0.1% HCO_2_H) in 0-3 min b) 5-95% in 3-18 min c) 95-95% in 18-24 min purity: 99% (*t_R_* = 18.2 min)

**Compound ent-1a, (*R*)-‘358**, purity: >99% (*t_R_* = 18.4 min)

**Compound 1b, (*S*)-SX240**, flow rate, 0.5 mL/min, linear gradient a) 10–10% CH_3_OH in H_2_O (+ 0.1% HCO_2_H) in 0-3 min b) 10-100% in 3-18 min c) 100-100% in 18-24 min purity: 95% (*t_R_* = 20.5 min)

**Compound ent-1b, (*R*)-SX240**, purity: >99% (*t_R_* = 20.9 min)

**Compound 1c, (*S*)-SX245**, flow rate, 0.5 mL/min, linear gradient a) 5–5% CH_3_CN in H_2_O (+ 0.1% HCO_2_H) in 0-3 min b) 5-95% in 3-18 min c) 95-95% in 18-24 min purity: 98% (*t_R_* = 19.6 min)

**Compound ent-1c, (*R*)-SX245**, purity: >99% (*t_R_* = 19.2 min)

**Compound 1d, (*S*)-SX244**, flow rate, 0.5 mL/min, linear gradient a) 5–5% CH_3_CN in H_2_O (+ 0.1% HCO_2_H) in 0-3 min b) 5-95% in 3-18 min c) 95-95% in 18-24 min purity: >99% (*t_R_* = 22.0 min)

**Compound ent-1d, (*R*)-SX244**, purity: >99% (*t_R_* = 22.0 min)

**Compound 1e, (*S*)-SX263**, flow rate, 0.5 mL/min, linear gradient a) 10–10% CH_3_OH in H_2_O (+ 0.1% HCO_2_H) in 0-3 min b) 10-100% in 3-18 min c) 100-100% in 18-24 min purity: 97% (*t_R_* = 21.0 min)

**Compound 1f, (*S*)-SX264**, flow rate, 0.5 mL/min, linear gradient a) 10–10% CH_3_OH in H_2_O (+ 0.1% HCO_2_H) in 0-3 min b) 10-100% in 3-18 min c) 100-100% in 18-24 min purity: >99% (*t_R_* = 20.8 min)

**Compound 1g, (*S*)-SX267**, flow rate, 0.5 mL/min, linear gradient a) 5–5% CH_3_CN in H_2_O (+ 0.1% HCO_2_H) in 0-3 min b) 5-95% in 3-18 min c) 95-95% in 18-24 min purity: >99% (*t_R_* = 19.6 min)

**Compound 1h, (*S*)-SX268**, flow rate, 0.5 mL/min, linear gradient a) 10–10% CH_3_OH in H_2_O (+ 0.1% HCO_2_H) in 0-3 min b) 10-100% in 3-18 min c) 100-100% in 18-24 min purity: >97% (*t_R_* = 20.1 min)

**Compound 2, I1381**, flow rate, 0.5 mL/min, linear gradient in 3–85% CH_3_CN in H_2_O (+ 0.1% TFA) in 0-26 min, purity: >99% (*t_R_* = 21.2 min)

**Compound 3a, I1384**, flow rate, 0.5 mL/min, linear gradient a) 3–85% CH_3_CN in H_2_O (+ 0.1% TFA) in 0-26 min b) 85-95% in 26-28 min c) 95%-95% in 28 – 30 min, purity: 95.7% (*t_R_* = 27.5 min)

**Compound 3b, I1402**, flow rate, 0.5 mL/min, linear gradient in a) 3–85% CH_3_CN in H_2_O (+ 0.1% TFA) in 0-26 min b) 85-95% in 26-28 min c) 95%-95% in 28 – 30 min, purity: 97.6% (*t_R_* = 28.7 min)

**Compound 3c, I1436**, flow rate, 0.5 mL/min, linear gradient a) 3–85% CH_3_CN in H_2_O (+ 0.1% TFA) in 0-26 min b) 85-95% in 26-28 min purity: >99% (*t_R_* = 26.3 min)

**Compound 4, BBG3**, flow rate, 0.5 mL/min, linear gradient in a) 3–85% CH_3_CN in H_2_O (+ 0.1% TFA) in 0-26 min, purity: >99% (*t_R_* = 24.6 min)

**Compound 5a, IK195**, flow rate, 0.5 mL/min, linear gradient in a) 3–85% CH_3_CN in H_2_O (+ 0.1% TFA) in 0-26 min, purity: >99% (*t_R_* = 24.6 min)

**Compound 5b, IK192**, flow rate, 0.5 mL/min, linear gradient in a) 3–85% CH_3_CN in H_2_O (+ 0.1% TFA) in 0-26 min b) 85-95% in 26-28 min, purity: 99.0% (*t_R_* = 26.7 min)

**Compound 5c, (*S*)-KD1**, flow rate, 0.5 mL/min, linear gradient a) 3–85% CH_3_CN in H_2_O (+ 0.1% TFA) in 0-26 min b) 85-95% in 26-28 min purity: >99% (*t_R_* = 27.0 min)

**Compound 5d, I1414**, flow rate, 0.5 mL/min, linear gradient 3–85% CH_3_CN in H_2_O (+ 0.1% TFA) in 0-26 min, purity: 96.3% (*t_R_* = 25.8 min)

**Compound 5e, SX270**, flow rate, 0.5 mL/min, linear gradient a) 10–10% CH_3_OH in H_2_O (+ 0.1% HCO_2_H) in 0-3 min b) 10-100% in 3-18 min c) 100-100% in 18-24 min purity: 98% (*t_R_* = 17.3 min)

**Compound 5f, I1424**, flow rate, 0.5 mL/min, linear gradient 3–85% CH_3_CN in H_2_O (+ 0.1% TFA) in 0-26 min, purity: 99.5% (*t_R_* = 18.9 min)

**Compound ent-5f, I1422** purity: 98.2%

**Compound 5g, I1408**, flow rate, 0.5 mL/min, linear gradient 3–85% CH_3_CN in H_2_O (+ 0.1% TFA) in 0-26 min, purity: 99.2% (*t_R_* = 20.2 min)

**Compound ent-5g, I1411** purity: 98.9%

**Compound F-5g, I1421**, flow rate, 0.5 mL/min, linear gradient 3–85% CH_3_CN in H_2_O (+ 0.1% TFA) in 0-26 min, purity: >99% (*t_R_* = 20.4 min)

**Compound 6a, (*S*)-SX276**, flow rate, 0.5 mL/min, linear gradient a) 5–5% CH_3_CN in H_2_O (+ 0.1% HCO_2_H) in 0-3 min b) 5-95% in 3-18 min c) 95-95% in 18-24 min purity: >99% (*t_R_* = 16.1 min)

**Compound 6b, (*S*)-SX272**, flow rate, 0.5 mL/min, linear gradient a) 10–10% CH_3_OH in H_2_O (+ 0.1% HCO_2_H) in 0-3 min b) 10-100% in 3-18 min c) 100-100% in 18-24 min purity: 96% (*t_R_* = 18.1 min)

**Compound 7a, I1409**, flow rate, 0.5 mL/min, linear gradient 3–85% CH_3_CN in H_2_O (+ 0.1% TFA) in 0-26 min, purity: 98.4% (*t_R_* = 21.7 min)

**Compound ent-7a, I1412**, purity: >99%

**Compound 7b, (*S*)-SX254**, flow rate, 0.5 mL/min, linear gradient a) 10–10% CH_3_OH in H_2_O (+ 0.1% HCO_2_H) in 0-3 min b) 10-100% in 3-18 min c) 100-100% in 18-24 min purity: 97% (*t_R_* = 19.2 min)

**Compound 8, (*S*)-SX258**, flow rate, 0.5 mL/min, linear gradient a) 10–10% CH_3_OH in H_2_O (+ 0.1% HCO_2_H) in 0-3 min b) 10-100% in 3-18 min c) 100-100% in 18-24 min purity: 97% *t_R_* = 19.6 min)

**Compound 9, I1380**, flow rate, 0.5 mL/min, linear gradient in a) 3–85% CH_3_CN in H_2_O (+ 0.1% TFA) in 0-26 min, purity: >99% (*t_R_* = 24.5 min)

**Compound 10a, KD2**, flow rate, 0.5 mL/min, linear gradient 3–85% CH_3_CN in H_2_O (+ 0.1% TFA) in 0-26 min, purity: >99% (*t_R_* = 23.6 min)

**Compound 10b-ent1, KD6en1**, flow rate, 0.5 mL/min, a) 3–85% CH_3_CN in H_2_O (+ 0.1% HCO_2_H) in 0-26 min, purity: >99% (*t_R_* = 25.9 min)

**Compound 10b-ent2, KD6en2**, purity: >99%

**Compound 10c-ent1, KD5en1**, flow rate, 0.5 mL/min, a) 3–85% CH_3_CN in H2O (+ 0.1% TFA) in 0-26 min, purity: 97.1% (*t_R_* = 22.5 min)

**Compound 10c-ent2, KD5en2**, purity: >99%

**Compound 10d-ent1, KD3en1**, flow rate, 0.5 mL/min, a) 3–85% CH_3_CN in H2O (+ 0.1% TFA) in 0-26 min b) 85-95% in 26-28 min, purity: 96.0% (*t_R_* = 26.0 min)

**Compound 10d-ent2, KD3en2**, purity: 96.7%

**Compound 10e, KD4**, flow rate, 0.5 mL/min, linear gradient 3–85% CH_3_CN in H_2_O (+ 0.1% TFA) in 0-26 min, purity: 97.2% (*t_R_* = 25.1 min)

**Exemplary HPLC Charts of the Investigation by Chiral HPLC**

**Compounds 1a, (*S*)-‘853** and **ent-1b, (*R*)-‘853** in comparison to the racemate lead structure **rac-’853** prepared analogously from racemic (*R*,*S*)-2-methyl-3-phenoxypropionic acid, chiral HPLC, hexane/isopropanol 9:1, 254 nm

**Compounds 1b, (*S*)-SX240** and **ent-1b, (*R*)-SX240**, chiral HPLC, hexane/ethanol 9:1, 254 nm

**Compounds 1c, (*S*)-SX245** and **ent-1c, (*R*)-SX245**, chiral HPLC, hexane/ethanol 98:2, 254 nm

**Compounds 1d, (*S*)-SX244** and **ent-1d, (*R*)-SX244**, chiral HPLC, hexane/ethanol 99:1, 254 nm

**Compound 1e, (*S*)-SX263**, chiral HPLC, hexane/ethanol 95:5, 254 nm

**Compound 1f, (*S*)-SX264**, chiral HPLC, hexane/ethanol 95:5, 254 nm

**Compound 1g, (*S*)-SX267**, chiral HPLC, hexane/ethanol 9:1, 254 nm

**Compound 1h, (*S*)-SX268**, chiral HPLC, hexane/ethanol 95:5, 254 nm

**Compound 3b, I1402**, chiral HPLC, hexane/isopropanol 99:1, 254 nm

**Compound 5c, (*S*)-KD1** in comparison to the racemate prepared analogously from racemic (*R*,*S*)-2-methyl-3-phenoxypropionic acid, chiral HPLC, hexane/isopropanol 98:2, 254 nm

**Compound 5d, I1414**, chiral HPLC, hexane/isopropanol 85:5, 254 nm

**Compound 5e, (*S*)-SX270**, chiral HPLC, hexane/isopropanol + 0.1% ethylene diamine 9:1, 254 nm

**Compounds 5g, I1408** and **ent-5g, I1411** in comparison to the racemate prepared analogously from racemic (*R*,*S*)-2-methyl-3-phenoxypropionic acid, chiral HPLC, hexane/ethanol 9:1, 254 nm

**Compound F-5g, I1421**, chiral HPLC, hexane/ethanol 9:1, 254 nm

**Compound 6a, (*S*)-SX276**, chiral HPLC, hexane/isopropanol 9:1, 254 nm

**Compound 6b, (*S*)-SX272**, chiral HPLC, hexane/isopropanol 4:1 + 0.1% ethylene diamine, 254 nm

**Compounds 7a, I1409** and **ent-7a, I1412** in comparison to the racemate prepared analogously from racemic (*R*,*S*)-2-methyl-3-phenoxypropionic acid, chiral HPLC, hexane/isopropanol 85:15, 254 nm

**Compound 7b, (*S*)-SX254**, chiral HPLC, hexane/isopropanol 9:1, 254 nm

**Compound 8, (*S*)-SX258**, chiral HPLC, hexane/isopropanol 9:1, 254 nm

**Compounds 10b (KD6)**, both enantiomers **10b-ent1** (**KD6en1)** and **10b-ent2 (KD6en2)** in comparison to the racemate, chiral HPLC, hexane/isopropanol 98:2, 254 nm

**Compounds 10c (KD5)**, both enantiomers **10c-ent1 (KD5en1)** and **10c-ent2 (KD5en2)** in comparison to the racemate, chiral HPLC, hexane/isopropanol 93:7, 254 nm

**Compounds 10d, KD3**, both enantiomers **10d-ent1 (KD3en1)** and **10d-ent2 (KD3en2)** in comparison to the racemate, chiral HPLC, hexane/isopropanol 98:2, 254 nm
